# PARP1 as a biomarker for early detection and intraoperative tumor delineation in epithelial cancers – first-in-human results

**DOI:** 10.1101/663385

**Authors:** Susanne Kossatz, Giacomo Pirovano, Paula Demétrio De Souza França, Arianna L. Strome, Sumsum P. Sunny, Daniella Karassawa Zanoni, Audrey Mauguen, Brandon Carney, Christian Brand, Veer Shah, Ravindra D. Ramanajinappa, Naveen Hedne, Praveen Birur, Smita Sihag, Ronald A. Ghossein, Mithat Gönen, Marshall Strome, Amritha Suresh, Daniela Molena, Moni A. Kuriakose, Snehal G. Patel, Thomas Reiner

## Abstract

Major determining factors for survival of patients with oral, oropharyngeal, and esophageal cancer are early detection, the quality of surgical margins, and the contemporaneous detection of residual tumor. Intuitively, the exposed location at the epithelial surface qualifies these tumor types for utilization of visual aids to assist in discriminating tumor from healthy surrounding tissue. Here, we explored the DNA repair enzyme PARP1 as imaging biomarker and conducted optical imaging in animal models, human tissues and as part of a first-in-human clinical trial. Our data suggests that PARP1 is a quantitative biomarker for oral, oropharyngeal, and esophageal cancer and can be visualized with PARPi-FL, a fluorescently labeled small molecule contrast agent for topical or intravenous delivery. We show feasibility of PARPi-FL-assisted tumor detection in esophageal cancer, oropharyngeal and oral cancer. We developed a contemporaneous PARPi-FL topical staining protocol for human biospecimens. Using fresh oral cancer tissues within 25 min of biopsy, tumor and margin samples were correctly identified with >95% sensitivity and specificity without terminal processing. PARPi-FL imaging can be integrated into clinical workflows, potentially providing instantaneous assessment of the presence or absence of microscopic disease at the surgical margin. Additionally, we showed first-in-human PARPi-FL imaging in oral cancer. In aggregate, our preclinical and clinical studies have the unifying goal of verifying the clinical value of PARPi-FL-based optical imaging for early detection and intraoperative margin assignment.

## Introduction

Cancers that originate in the epithelium of exposed or accessible body areas, such as oral and oropharyngeal cancers as well as esophageal cancer, should be easy to screen for by monitoring and identifying visible changes in the mucosal lining. However, cancer registry data reveal that only 29% of oral and oropharyngeal cancers and 19% of esophageal cancers are detected at early stages, when the cancer is still confined to its primary site. Survival rates steeply drop with detection at advanced tumor stages. The five-year survival rate in the US is 83.7% for oral cancer and 45.2% for esophageal cancer when the disease is still localized at the time of diagnosis, compared to 39.1% (oral cancer) and 4.8% (esophageal cancer) for metastatic disease (*1, 2*). Clinical outcome is also negatively affected by the presence of residual tumor at the surgical margins in the postsurgical histopathology exam of the resected specimen (*3–5*). Intuitively, this suggests that the current clinical practice of visual examination in combination with biopsy-based histopathological diagnosis is insufficient to improve early diagnosis and consequently, to improve treatment outcomes for patients with cancers that develop close to the epithelial surface.

For initial diagnosis, identifying early-stage morphological changes and distinguishing among a variety of inflammatory and benign conditions, given their potentially similar appearance, is difficult. During surgical interventions, success is typically dependent on the absence of positive margins and micro-metastasis. Current clinical practice, e.g., for oral cancer, calls for surgical excision with at least 5 mm margins of normal tissue on histopathologic examination to reduce the risk of residual microscopic disease. Wide margins, however, can lead to additional morbidity and significant, irreversible impairment of phonation, mastication, gustation, and/or swallowing. Therefore, the current surgical paradigm consists of the surgeon arbitrarily removing at least 10 mm of “normal” appearing tissue in all dimensions around the visible and palpable extent of the tumor. Following excision of the tumor, frozen section histopathology is often used to evaluate the presence of tumor cells at the surgical margins. However, this approach leads to significant time delays during surgery and has less accuracy than the gold standard permanent histopathology due to tissue artifacts caused by the freezing process. Frozen section histopathology is also afflicted by high false-negative rates—intuitively, a direct consequence of the small sampling volume used for sectioning. Lastly, action is delayed several days until the definitive histopathological diagnosis is available, often leading to secondary surgeries and/or adjuvant treatment with radiation and chemotherapy, which are all associated with compounded morbidity, further cost, and poorer oncologic outcomes.

Enhancing the visibility of malignant lesions in relation to normal tissues contemporaneously— i.e., adding contrast—therefore holds immense potential to address the above-mentioned challenges associated with diagnosis and intraoperative imaging. In pursuit of this goal, a host of methods have been explored as clinical diagnostic adjuncts, including tissue staining with vital dyes (e.g., Toluidine Blue, Methylene Blue, or Lugol’s Iodine), chemiluminescence imaging following an acetic acid wash, and optical-based imaging techniques, which rely on autofluorescence (*6–10*). To date, these methods have shown limited sensitivity and/or specificity and are generally not recommended (*11*).

A new class of emerging diagnostic adjuncts focuses on *in vivo* microscopic (IVM) techniques. These include optical coherence tomography (OCT), reflectance confocal microscopy (RCM), and high-resolution microendoscopy, which aim to noninvasively visualize the features typically assessed via histopathology during the examination. Currently, IVM techniques for epithelial cancers are in early-stage development and their diagnostic accuracy remains to be determined. Thus, existing approaches possess limitations that have prevented their clinical adoption. Those limitations are i) absence of a tumor-specific molecular target, ii) confinement to either microscopic or macroscopic resolution iii) diagnostic accuracy that is limited by either low sensitivity or low specificity (*6, 7, 11–13*). In contrast, employing the optically active, molecularly targeted approach presented in this work could not only improve sensitivity and specificity of tumor recognition, but also allow for imaging on both the cellular and macroscopic level, which, if combined, could increase the breadth and depth of the molecular information used for early detection and intraoperative margin delineation.

The molecular target for our optical imaging approach is the DNA repair enzyme Poly(ADP-ribose)Polymerase 1 (PARP1). In addition to its many functions in cell cycle regulation and transcription, high levels of PARP1 expression have been observed in many different tumor types, indicating its potential for diagnostic applications (*14–20*). Higher expression has moreover been linked to worse prognosis (*21–24*). While PARP1 also appears in the nuclei of normal, proliferating cells, expression levels and especially cellularity tend to be lower than in rapidly growing tumors, supporting the rationale that high-contrast imaging can be achieved with PARP-targeted imaging approaches.

The optical PARP1-targeted imaging agent PARPi-FL is a fluorescently labeled analog of the FDA-approved PARP inhibitor olaparib with high affinity and specificity for PARP1 (*25*). PARPi-FL is a cell penetrating imaging agent that rapidly clears in the absence of target binding. Importantly, compared to other nuclear staining vital dyes, PARPi-FL does not intercalate into the DNA, but instead reversibly binds to PARP1, which locates to DNA strand breaks, and is not mutagenic, enabling *in vivo* applications in addition to *ex vivo* applications. The efficacy of high-contrast xenograft imaging with PARPi-FL after intravenous application has been shown on the whole body as well as at the cellular level (*25, 26*). Importantly, the tissue penetrating nature of PARPi-FL enables its use as both a systemic and topical agent. Topical application, as an alternative to intravenous injection, enables the application of microdoses of PARPi-FL and allows for almost immediate imaging (*27*). However, this concept has neither been tested systematically nor was it tested in humans. The versatile features of PARPi-FL make it a tool for use in early detection as well as surgical margin assignment applications, both of which could strongly benefit from a diagnostic adjunct.

Oral and oropharyngeal cancer as well as esophageal cancers typically grow at the epithelial surface. For all three, tumors are currently inspected by visual examination, followed by incisional biopsy without visual aids. We hypothesized that these malignancies would benefit most from PARP1-targeted optical imaging to improve tumor detection and increase the accuracy of tumor excision. The goal of the latter is to increase surgical success by facilitating contemporaneous margin delineation and minimizing the morbidity of oral surgery.

In this study, we aimed to characterize the potential of PARP1 expression as a quantitative biomarker for early detection and intraoperative delineation in human biospecimens and to subsequently establish PARPi-FL-based imaging strategies using human tissues and preclinical models. Due to differences in anatomical accessibility of esophageal, oropharyngeal, and oral cancer, we did not seek to create a one-size-fits-all optical imaging approach, but rather pursued individually adapted PARPi-FL imaging approaches to better inform clinical development.

In esophageal cancer, we validated PARPi-FL as a quantitative marker for PARP1 expression levels and showed that it could be applied topically to the esophagus. In oropharyngeal cancer, we tested whether combining brush biopsy (a less invasive approach to obtaining diagnostic tissue) with PARPi-FL staining is a feasible approach to identifying tumor cells in a complex tissue sample. In oral cancer, we developed a staining and imaging method for freshly excised biopsies that allows for the identification of positive margins within minutes while preserving the fresh tissue for other downstream applications. We then tested the diagnostic accuracy of this detection method in human biopsies and evaluated its translational potential. In additional support of the translational potential of PARPi-FL, we show first-in-human *in vivo* imaging after topical application (NCT03085147). Finally, we explore the potential to expand PARPi-FL imaging to *in situ* surgical guidance applications with a large surgical window after intravenous delivery in a large animal. The presented studies are consistent in demonstrating the clinical relevance of PARPi-FL-based optical imaging methods for the early detection and intraoperative delineation of cancers that develop at the epithelial surface. An overview of all conducted studies can be found in **Fig. 1**.

**Fig. 1.**
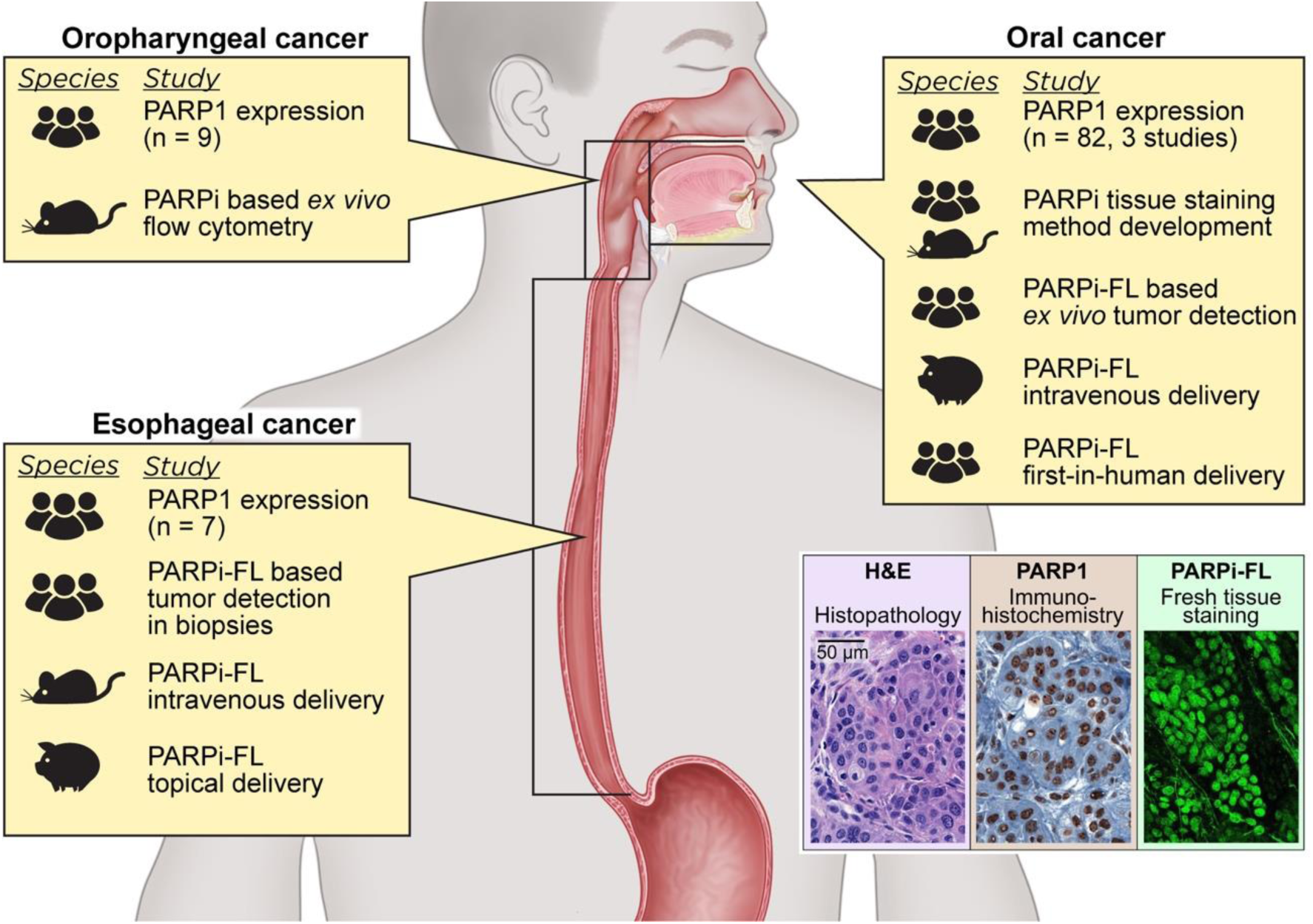
Study overview. We analyzed PARP1 expression via IHC staining in esophageal, oropharyngeal, and oral cancer samples and carried out different PARP1 imaging studies. The figure illustrates the type of imaging-based study and species used for each tumor type.

## Results

### PARP1 biomarker validation and imaging in esophageal cancer

To explore and confirm the clinical value of a PARP1-targeted imaging approach in esophageal cancer, we quantified PARP1 expression in human surgical biospecimens (n=7). In all specimens, three regions of interest were analyzed: tumor, normal epithelium, and deep margin (comprising submucosa and muscle tissue). We found strongly increased PARP1 expression in tumor compared to deep margin as well as epithelium, which had limited physiological PARP1 expression in its basal layer (**Fig. 2A, fig. S1A**). Quantification, measured as PARP1-positive area (details in Materials and Methods and **fig. S2**), revealed a mean value of 26.4% ± 10.8% PARP1-positive area in the tumor, which was significantly higher than the epithelium (5.0% ± 1.9%, p=0.03, Wilcoxon test) and the deep margin (0.9% ± 0.5%, p=0.02, Wilcoxon test) (**Fig. 2B** **and fig. S1, B and C**).

**Fig. 2.**
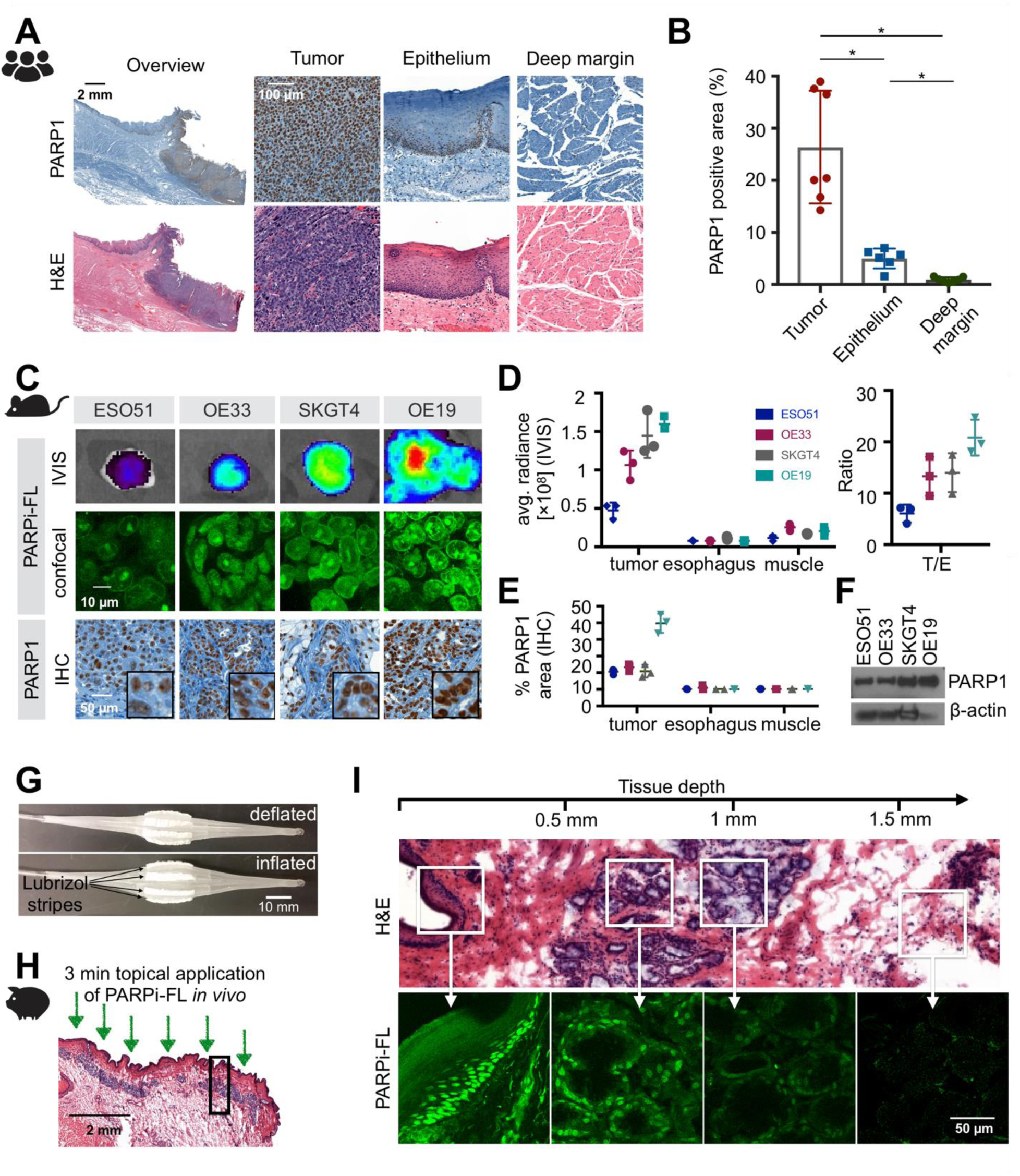
PARP1 expression and imaging in esophageal cancer. **(A)** Representative PARP1 IHC and H&E histology obtained from human biospecimens. **(B)** Quantification of PARP1 expression in IHC samples (n=7 patients). Statistical significance was determined using the Wilcoxon matched pairs signed rank test (all values present for n=6 patients). * p<0.05. **(C)** Fluorescence imaging of EAC in xenograft mouse models after intravenous injection of 75 nmol PARPi-FL. Excised tumors were imaged using epifluorescence (IVIS) and confocal microscopy. PARP1 expression was confirmed via IHC staining. **(D)** Quantification of the fluorescence signal 90 min post-injection of PARPi-FL and tumor/esophagus (T/E) contrast ratios. **(E)** Quantification of PARP1 expression of tumors, esophagus, and muscle assessed by PARP1 IHC. **(F)** Quantification of PARP1 expression in EAC cell lines via Western blot. **(G)** Inflatable balloon applicator used to topically apply PARPi-FL to a Yucatan mini swine esophagus. **(H)** 2 mL of a 1000 nM PARPi-FL solution were loaded into the balloon applicator and topically applied for 3 min onto the esophagus of the anaesthetized mini swine. **(I)** Penetration of topically applied PARPi-FL was visualized in perpendicular cryosections of the esophagus using a confocal microscope. H&E staining on adjacent sections was used for anatomical reference.

### PARP1-targeted optical imaging in esophageal cancer

The ability of PARPi-FL to identify high and low PARP1-expressing tumors was tested in four mouse xenograft models (ESO51, OE33, SKGT3, OE19) of esophageal adenocarcinoma (EAC). Tumor-bearing animals were sacrificed 90 min after intravenous injection of 75 nmol of fluorescent PARPi-FL. Macroscopic epifluorescence imaging of the excised tumors revealed increasing fluorescence intensities (ESO51<OE33<SKGT4<OE19), which was corroborated via confocal microscopy, where the nuclear accumulation of PARPi-FL could be clearly seen (**Fig. 2C**). Subsequent PARP1 immunohistochemistry (IHC) confirmed that increasing PARPi-FL uptake correlates with cellular PARP1 expression (**Fig. 2C**). Quantification of the fluorescence intensity in tumor, esophagus, and thigh muscle showed a three-fold increase from the lowest (ESO51) to highest (OE19) PARP1-expressing xenograft model, while fluorescence intensities in normal esophagus and muscle tissue were low and comparable in all animals (**Fig. 2D**). Tumor-to-esophagus ratios were high in all models, ranging from 6.1 ± 1.7 (ESO51) to 20.8 ± 3.5 (OE19), providing adequate contrast to discriminate tumor from normal esophagus tissue in both high and low PARP1-expressing tumors. PARP1 expression levels of the xenograft models were also quantified via PARP1 IHC (**Fig. 2E**) and Western blot (**Fig. 2F**). In addition, uptake of the radioactive, fluorine-labeled PARPi [^18^F]PARPi further corroborated that PARPi-FL uptake was quantitative, showing a similar increase in *in vivo* tumor uptake of the xenograft models (ESO51<OE33<SKGT4<OE19) with high tumor-to-muscle ratios (**fig. S3**).

### Validation of topical administration route for PARPi-FL in the esophagus

The feasibility of PARPi-FL topical application has been shown in oral cancer (*26*). However, given the limited accessibility of the esophagus, we explored an alternative topical application strategy and measured penetration depth of PARPi-FL in the esophagus of a pig using a balloon applicator with sponge-like Lubrizol stripes (**Fig. 2G**) soaked with 2 mL of a 1 µM PARPi-FL solution. After insertion into the pig’s esophagus in a deflated state, the balloon was inflated to expose the Lubrizol stripes and induce release of PARPi-FL onto the esophagus wall of the anesthetized pig for a duration of 3 min; 10 min after the application, the pig was sacrificed, followed by necropsy and cryopreservation of the esophagus in 10×10 mm tiles. Cryosections were cut perpendicular to the esophagus surface (**Fig. 2H**). On fresh slides, without further staining, we detected nuclear PARPi-FL staining in the basal layer of the epithelium (100-200 µm depths) and mucus glands in the submucosa, up to 800 µm below the tissue surface (**Fig. 2I**). Deeper-lying mucus glands and submucosal tissue (>1000 µm depth) showed very low to non-detectable fluorescence. We found that PARPi-FL was able to penetrate approximately 800 µm deep into the esophagus within 3 min of topical application and bind to PARP1 in cell nuclei without noticeable background staining.

### PARP1 expression in oropharyngeal cancer

Expanding on whether PARP1 could be a relevant general biomarker to tumors of the oral cavity, we investigated PARP1 expression in a clinical data set of oropharyngeal cancer, which arises at the base of the tongue, tonsils, soft palate, or pharynx wall (n=9). All biospecimens contained invasive carcinomas with high levels of PARP1 expression (**Fig. 3A** **and fig. S4A**). In the normal epithelium, PARP1 expression was found in the basal layer and at very low levels in the deep margin (**Fig. 3A** **and fig. S4A**). Analogous to esophageal biospecimens, quantification of the PARP1-positive area confirmed significantly higher expression levels in the tumor (46.5% ± 16.7% PARP1-positive area) than the epithelium (9.4% ± 4.9%; p=0.02, Wilcoxon test) and deep margin (2.0% ± 1.1%; p<0.01, Wilcoxon test) (**Fig. 3B** **and fig. S4B**).

**Fig. 3.**
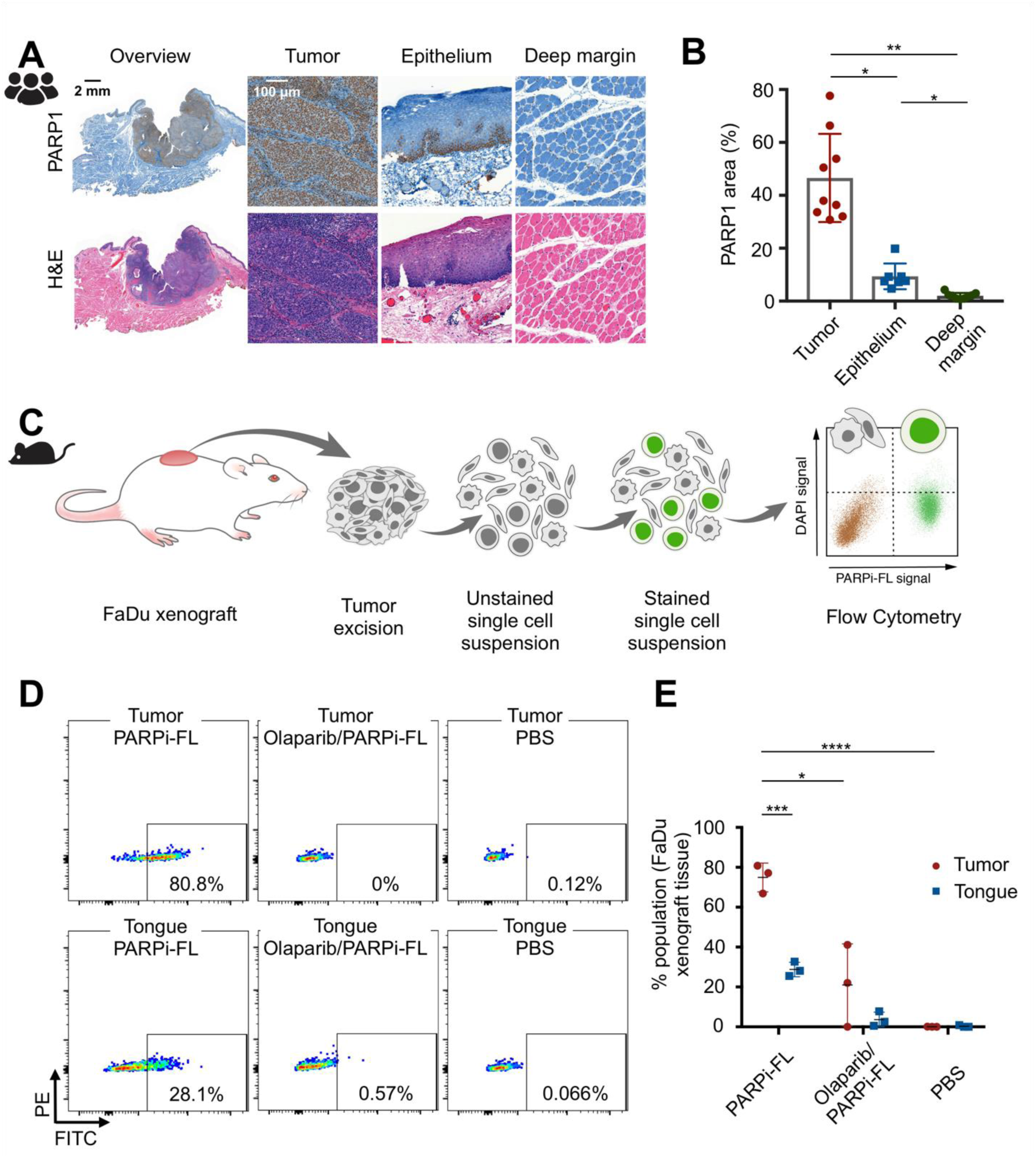
PARP1 expression and PARP cytometry in oropharyngeal cancer. **(A)** Representative PARP1 IHC from human biospecimens of oropharyngeal cancer, displaying PARP1 expression in the epithelium, deep margin, and tumor, with corresponding H&E images. **(B)** Quantification of PARP1 expression in IHC samples (n=9 patients). Statistical significance was determined using the Wilcoxon matched pairs signed rank test (all values present for n=7 patients). * p<0.05, ** p<0.01. **(C)** Workflow of PARP cytometry. Tissue samples (FaDu xenograft, tongue) were dissociated into single cell suspensions, stained with 100 nM PARPi-FL, and subjected to flow cytometry to determine the percentage of PARPi-FL stained cells in the sample. **(D)** Dot plots of the gated samples showing PARPi-FL-positive cells (FITC channel) vs. live/dead stain (DAPI). Representative dot plot for tumor and tongue stained with 100 nM PARPi-FL or controls, which were used to assess specificity of the PARPi-FL binding (Olaparib/PARPi-FL) and specificity of the signal (PBS). **(E)** Quantification of PARPi-FL flow cytometry (n=3 mice processed in 3 separate experiments). Significance was determined using an unpaired t-test. * p<0.05, *** p<0.001, ****p<0.0001.

### A brush biopsy approach to PARP1-targeted optical imaging

Brush biopsy techniques have been investigated as a less invasive alternative to standard punch and scalpel biopsies. We conducted a feasibility study to determine if brush biopsies in combination with PARPi-FL staining could enable rapid detection of tumor cells in cell suspensions derived from solid tissues. In a proof-of-principle approach, we developed an *ex vivo* staining protocol and flow cytometric analysis of single-cell suspensions derived from pharynx cancer xenografts (FaDu cells) and healthy mouse tongues (**Fig. 3C**). We analyzed PARPi-FL staining of tumor and tongue tissue after applying appropriate gating procedures (**fig. S5A**). We found a significantly higher number of PARPi-FL-positive cells in tumor-derived suspensions than in tongue-derived suspensions (**Fig. 3, D and E**; p<0.001, unpaired t-test). Specificity of PARPi-FL staining in cell suspensions was confirmed by blocking PARPi-FL binding sites with an excess of the non-fluorescent PARP inhibitor olaparib, which reduced the percentage of positive cells from 74.9% ± 7.2% to 21.0% ± 20.0% (p<0.05, unpaired t-test using the Holm-Sidak method without assuming a consistent SD). A PBS-stained control group showed no PARPi-FL-positive cells. In addition, we were able to confirm these results in esophageal cancer cells (OE19), which also showed a higher percentage of PARPi-FL-positive cells in tumor compared to tongue tissue and a significant uptake reduction after olaparib blocking (**fig. S5B**). Thus, this proof-of-principle experiment showed that PARPi-FL staining of a cell suspension derived from a complex tissue can help to distinguish between tumor cell containing and normal samples, encouraging further investigation into combining PARPi-FL staining with brush biopsy approaches.

### PARP1 expression during malignant transition and at tumor margins in oral cancer

In oral cancer, we confirmed the gradual increase of PARP1 expression during malignant transition. We further analyzed tumor margins to gain insight into the clinical potential of PARP1-based optical imaging for early detection and intraoperative guidance.

In healthy epithelial tissues, physiological PARP1 expression was localized predominantly to the basal layer of the epithelium. In benign, dysplastic, and malignant biospecimens (n=60), we found an increase in PARP1 expression, particularly for cases with severe dysplasia (=carcinoma in situ) and in invasive tumors (**Fig. 4A**). This was reflected in the quantification of the PARP1-positive area, which was significantly higher in severe dysplasia and tumors compared to benign cases, mild dysplasia, and moderate dysplasia (p<0.05, Mann-Whitney test; **Fig. 4B**). Combining cases that are predominantly treated non-surgically (benign, mild dysplasia) and cases that require surgical resection with adjuvant radiation/chemotherapy (severe dysplasia/tumors), we found a pronounced, statistically significant difference in PARP1 expression (5.6% ± 2.6% vs. 13.1% ± 6.8%, p<0.001, Mann-Whitney test) (**Fig. 4C**). We determined a sensitivity of 81% and specificity of 83% at the optimal cutoff of 8.1% PARP1-positive area for separation of these two groups, with an area under the curve (AUC) of 0.908 (**Fig. 4D**). To substantiate these results, we also employed a manual scoring method for PARP1 IHC, based on the percentage of PARP1-positive cells and staining intensity (**fig. S6A**). Manual scoring yielded similar results, resulting in a sensitivity of 82% and specificity of 86% for the differentiation between benign/mild dysplasia and severe dysplasia/tumor (**fig. S6B**).

**Fig. 4.**
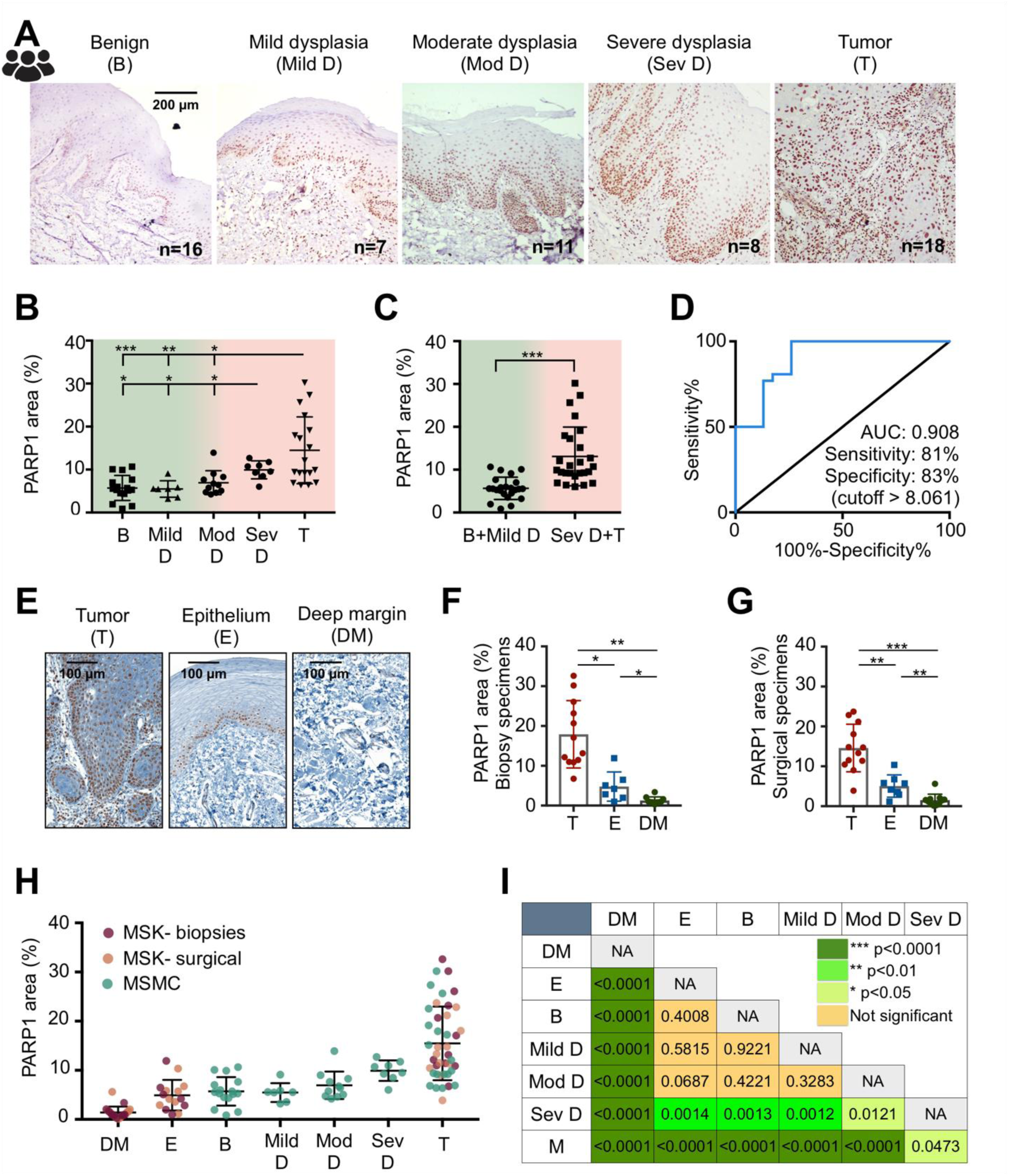
PARP1 expression during malignant transition and at tumor margins in oral cancer. **(A)** Representative PARP1 IHC images in different stages of oral cancer progression (benign, mild dysplasia, moderate dysplasia, severe dysplasia, malignant) of patient biospecimens from Mazumdar Shaw Medical Center (MSMC). **(B)** Quantification of PARP1 expression in IHC samples (n=60 patients). The PARP1-positive area (in relation to the entire tissue area) was quantified in high-resolution images of the epithelium or tumor area (mean of n ≥ 4 images per case). Statistical significance was determined using an unpaired Mann-Whitney rank sum test. **(C)** Comparison of PARP1 expression in low-grade, potentially malignant lesions (benign vs. mild dysplasia) vs. severe dysplasia (carcinoma in situ) and malignant cases. Statistical significance was determined using an unpaired Mann-Whitney rank sum test. **(D)** ROC curve for the data represented in (C). **(E)** Representative images of PARP1 expression in samples from two populations: presurgical biopsies of tumor and benign tissues (divided in two areas: epithelium and deep margin) and surgical specimens, which included tumor, adjacent benign epithelium, and deep margin on each specimen. **(F)** Quantification of PARP1 expression in IHC samples of biopsy specimens (n=12 patients). Statistical significance was determined using the Wilcoxon matched pairs signed rank test (all values present for n=6 patients). **(G)** Quantification of PARP1 expression in IHC samples of surgical specimens (n=12 patients). Statistical significance was determined using the Wilcoxon matched pairs signed rank test (all values present for n=8 patients). A different analysis of the same dataset has been published previously (*26*). **(H)** Pooling of all three datasets of oral biospecimens investigated for PARP1 expression via IHC. **(I)** Statistical significance of the datasets in (H) was determined between all groups using the unpaired Mann-Whitney rank sum test. T-Tumor. E-Epithelium, DM-deep margin, D-dysplasia, mod-moderate, sev-severe, b-benign. Significance levels: *p<0.05, **p<0.01, ***p<0.001.

In presurgical biopsies (tumor: n=12, benign tissues: n=10 subdivided into epithelium and deep margin from n=12 patients) and surgical specimens (n=12), we quantified the difference in PARP1 expression between the tumor area, epithelium, and deep margin (**Fig. 4E**). While PARP1 staining was high in the tumor area of both biopsies (17.9% ± 8.5% PARP1-positive area; **Fig. 4F**) and surgical specimens (14.6% ± 6.0% PARP1-positive area; **Fig. 4G**), it was significantly lower in the epithelium (4.8% ± 3.6% and 5.1% ± 2.8%; p=0.03 and p=0.008, respectively, Wilcoxon test) as well as in the deep margin (1.3% ± 0.9% and 1.5% ± 1.5%; p=0.04 and p=0.001, respectively, Wilcoxon test). Importantly, when only looking at paired datasets, PARP1 expression was always higher in the tumor than in the epithelium or deep margin, with no case of overlap, which was also the case in biospecimens of oropharyngeal and esophageal tumors (**fig. S7**).

Combining all oral cancer IHC datasets (three datasets, n=84 patients total) confirmed consistently elevated PARP1 expression in tumors and severe dysplasia, whereas a much lower expression was found in the epithelium and deep margin of normal oral tissue and benign and early dysplastic cases (**Fig. 4H**). Analyses of statistically significant differences in this dataset (Mann-Whitney test, **Fig. 4I**) support PARP1 expression as viable biomarker for both early detection and intraoperative margin delineation of oral cancer. Overall, PARP1 levels increased significantly with disease stage, from a median of 1.0% in deep margin to 13.2% in tumor (Kendall’s tau = 0.67, p<0.0001).

### Rapid, tissue-preserving PARPi-FL staining of fresh biopsies

Motivated by the abrupt drop of PARP1 expression in benign tissues adjacent to the tumor, we wanted to determine whether PARPi-FL uptake can serve as a diagnostic tool in fresh biopsy tissue. We aimed to develop a protocol that does not interfere with tissue integrity or morphology (e.g., by freezing or fixing) and can produce results faster than standard frozen section histopathology. In protocol optimization studies, we tested different staining concentrations, staining times, and washing times on freshly excised FaDu xenograft tissue toward high nuclear staining intensity and low cytosolic and non-specific background staining (**fig. S8**). The selected fresh tissue staining protocol consisted of 5 min PARPi-FL staining followed by 10 min of washing, and was translated to fresh human biopsy samples. A similar protocol can be used for non-fixed cryosections, leading to a nuclear staining pattern that reflects PARP1 expression of the same OE19 xenograft specimen (**fig. S9**). Staining specificity was confirmed by PARP1 IHC and H&E staining (**Fig. 5A**).

**Fig. 5.**
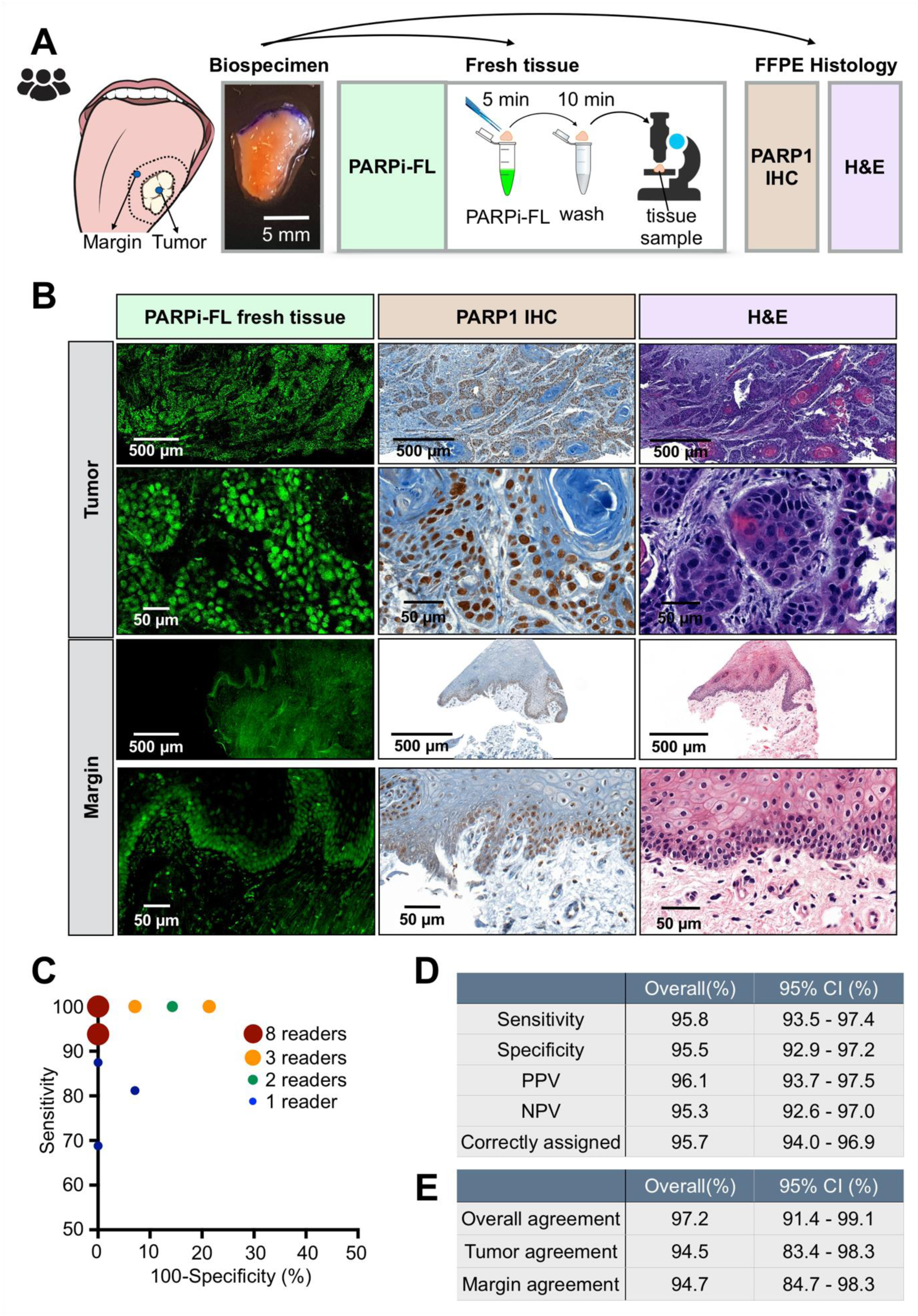
Rapid PARPi-FL staining on fresh biospecimens. **(A)** Workflow of rapid PARPi-FL staining of fresh human biospecimens. Biospecimens were split and separately processed for fresh tissue staining as well as histopathology. **(B)** Examples of PARPi-FL staining of a tumor and margin sample and corresponding histopathology of the same sample. We aimed to scan the entire fresh tissue in a high-resolution tile scan. Lower and higher magnification images showcase PARPi-FL staining and corresponding PARP1 expression in the specimens. A total of 22 tissues (n=12 tumors and n=10 benign tissues adjacent to tumor from 13 individual patients) were stained and analyzed. **(C)** In a blinded study, readers (n=27) scored 30 cases (n=12 tumors, n=10 margins, n=8 duplicates (4 tumors, 4 margins)) as tumor or margins (see **fig. S9** for study design). Pairs of sensitivity and specificity for each reader are represented in the graph. **(D)** Average values for sensitivity (95.8%), specificity (95.5%), positive predictive value (96.1%), negative predictive value (95.3%), and correctly assigned cases (95.7%). **(E)** Overall tumor agreement was 94.5% and overall margin agreement was 94.7%. Overall agreement was 97.2%.

Confocal microscopy images of tumor samples showed abundant nuclear PARPi-FL in areas identified as PARP1-expressing tumor cells based on PARP1 IHC and H&E staining (**Fig. 5B**). In benign samples, nuclear PARPi-FL staining was confined to the thin PARP1-expressing basal layer of the epithelium (**Fig. 5B**). Endogenous collagen-related autofluorescence, which was observed in the submucosa of the clinical samples, was discriminated from the exogenous PARPi-FL signal by two-channel imaging. PARPi-FL signals only appeared after 488 nm excitation, while autofluorescence was present after both the 488 nm and 543 nm excitations (**fig. S10A**). Confocal images were acquired from fresh biopsies of 12 patients and compiled into a study set of 30 cases (n=12 tumors, n=10 benign tissues adjacent to tumor, n=8 duplicates (4 tumors, 4 benign tissues adjacent to tumor)).

The images, acquired at 488 nm and 543 nm excitations, were read by volunteers (n=27) in a blinded study; each specimen was scored as tumor or benign tissue adjacent to tumor (see **fig. S10B** for study design and **Supplementary Materials PDF files 1 and 2** for the blinded study training set and data set). Of a total of 810 ratings, 361/378 negative margins and 414/432 tumors were assigned correctly (**fig. S10, C and D**). Pairs of sensitivity and specificity for each reader are reported in **Fig. 5C**. We found an overall sensitivity of 95.8%, an average specificity of 95.5%, a positive predictive value of 96.1% and a negative predictive value of 95.3% (**Fig. 5D**). To assess intrareader agreement, 8 cases were presented as duplicates without the readers’ knowledge (images were mirrored and rotated). Overall intrareader agreement was 97.2% (**Fig. 5E** **and fig. S10E**). We confirmed the non-destructive nature of the staining to biopsy tissue by carrying out regular, permanent H&E histopathology following PARPi-FL staining and imaging and found no evidence of perturbed tissue integrity or quality (**fig. S11**). Thus, PARPi-FL allowed for rapid staining and identification of tumor cells in fresh biopsies with high sensitivity and specificity, and without impeding the use of the sample for regular processing afterwards.

The purpose of the presented dataset was to validate our PARPi-FL staining method and identify if samples containing tumor (in the clinical setting: positive margins) can be reliably distinguished from samples not containing tumor (negative margins). Most importantly, considering an intraoperative setting, our data also suggests that focal tumor invasion into a suspected negative margin or a predominantly negative biopsy can be clearly identified and demarcated from surrounding normal tissue (**fig. S12**).

The simplicity of the PARPi-FL staining method, paired with its contemporaneous readout, makes for a technology well-suited to clinically relevant settings, including the operating room. Admittedly, the rate-limiting step of this technology would be image acquisition with a state-of-the-art confocal microscope. Reasoning that a quicker, more user-friendly readout would be preferable, we tested a confocal scanner, which is designed to rapidly analyze fresh tissue specimens by creating a high-resolution strip-mosaic (Vivascope 2500, Caliber ID, Andover, MA). We stained fresh FaDu xenograft tissue with PARPi-FL (0, 100, 250 nM) for 10 min, placed the tissue on the scanner, and scanned the field of view of 15×15 mm in about 150 s. The high-resolution images allowed for clear identification of PARPi-FL-stained nuclei (**fig. S13A**). This was confirmed using human biospecimens, which were also used for regular confocal microscopy (**compare** **Fig. 5B**). Tumor cell nuclei in the human biospecimen samples were clearly identified based on their strong PARPi-FL staining, which was in alignment with H&E and PARP1 IHC (**fig. S13B**). The Vivascope 2500 could be used as a back-table instrument in a surgical suite, allowing for ad hoc PARPi-FL images of freshly excised tissues, before tissue is passed on for other in-depth analyses such as cryosections, IHC, or sequencing.

We also explored the feasibility of macroscopic and microscopic *in vivo* imaging. Here, we validated an *in vivo* imaging device, the View*n*Vivo (Optiscan, Mulgrave, Australia), a miniaturized handheld confocal endomicroscope with an adjustable focal plane, suitable for *in vivo* or *ex vivo* imaging (**Fig. 6A**). The device is equipped with a 488 nm laser—the optimal wavelength for PARPi-FL detection. The choice of different longpass and bandpass filters allows for discrimination between specific PARPi-FL fluorescence and autofluorescence. The z-mechanism was able to detect PARPi-FL at up to 50 µm tissue depths (**fig. S14, A and B**). Images of the same human biospecimens as presented in fig. S13B revealed comparable image quality for tumor detection, including clear delineation of tumor cell nuclei and the PARP1-expressing basal layer in margin tissue (**Fig. 6B**). Intuitively, the View*n*Vivo has the potential to be used for intraoperative, PARPi-FL-based *in vivo* imaging without tissue excision. For this specific application – imaging in the surgical cavity – intravenous instead of topical delivery would be required, since distribution of PARPi-FL in the entire tumor mass would enable surgical guidance across the whole surgical bed. Key questions in this regard are the necessary injected mass and surgical window for imaging. Intravenous injection of 0.05 mg/kg PARPi-FL in a pig (**fig. S15A**) resulted in a blood half-life of 1.6 ± 0.48 min and simultaneously revealed that it was possible to detect PARPi-FL specifically in PARP1 expressing nuclei in the epithelial basal layer as soon as 5 min post injection and up to 120 min post injection (**fig. S15, B and C**). This suggests feasibility of PARPi-FL injection during surgery to provide visual guidance, to identify tumor margins and to detect focal lesions in the surgical cavity.

**Fig. 6.**
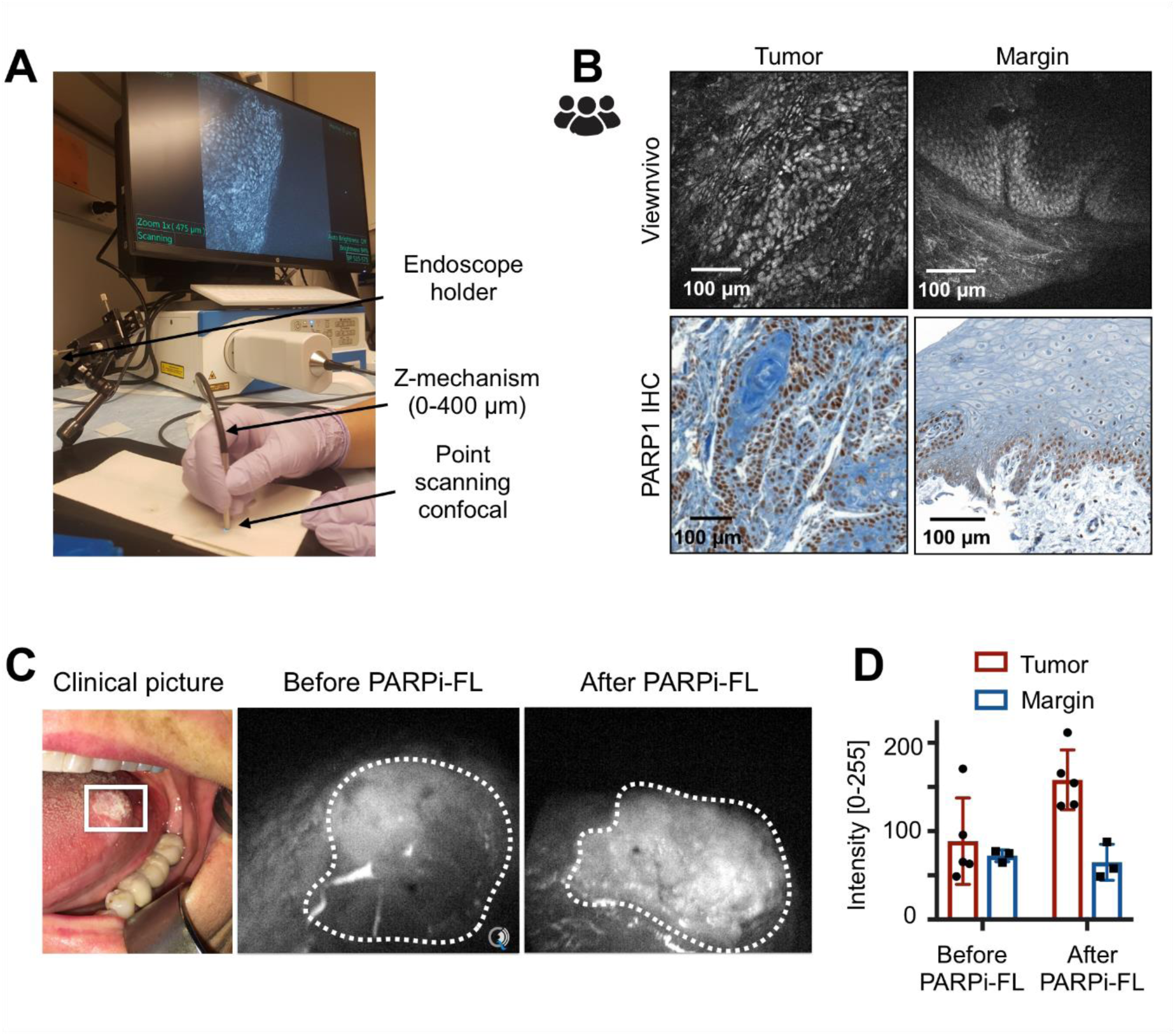
Feasibility of microscopic and macroscopic in-human imaging of PARPi-FL. (**A**) Imaging setup for the View*n*Vivo point scanning confocal endomicroscope suitable for *in vivo* imaging. (**B**) Fluorescence images from a patient sample stained with 100 nM PARPi-FL and corresponding PARP1 IHC. (**C**) PARPi-FL first-in-human imaging. An oral squamous cell carcinoma patient was imaged as part of clinical evaluation of PARPi-FL (NCT03085147). Using a Quest Spectrum imaging device with a laparoscopic camera and PARPi-FL optimized laser/filter system the tumor area of the patient was imaged before and after gargling a 250 nM PARPi-FL solution. (**D**) Fluorescence intensities in the tumor and surrounding non-tumor area were analyzed using ImageJ.

Lastly, while the first-in-human clinical trial using PARPi-FL (NCT03085147) is ongoing, feasibility of translation to in-human imaging has been shown using a topically applied mouthwash (**Fig. 6C**). Compared to images before contrast application, a one-minute gargle with 250 nM PARPi-FL led to a 1.9-fold increase in tumor fluorescence, resulting in a tumor-to-non-tumor ratio of 2.4 (**Fig. 6D**).

## Discussion

In this study, supported by both clinical and preclinical specimens, we designed and validated a fluorescence-driven imaging approach for oral, oropharyngeal, and esophageal cancer. In human biospecimens, and across all three investigated cancer types, we confirm that malignant tissues express significantly higher levels of PARP1 than normal epithelium and submucosal tissue, supporting our central hypothesis that PARP1 will yield high-contrast imaging as a diagnostic and intraoperative biomarker. We experimentally demonstrate how PARPi-FL staining and high-resolution imaging has the potential to improve early diagnosis and surgical removal of tumors. Of particular translational importance, PARPi-FL-based differentiation of tumor and normal tissue in fresh human oral cancer biopsies could be achieved with >95% sensitivity and specificity. PARPi-FL imaging uses fresh tissue for staining and is non-destructive and contemporaneous (5-min staining time), all features that promote effortless clinical integration. Intuitively, the absence of terminal processing preserves the tissue for further applications following PARPi-FL imaging, such as histopathological, proteomic, or genomic analyses.

We investigated PARP1 expression in esophageal cancer, consisting of adeno- and squamous cell carcinomas (stage T1-T3). We found that all tumors had higher levels of PARP1 expression (14.3 to 37.6% PARP1-positive area) than normal epithelium and submucosal (deep margin) tissue (0.3% to 7.0% PARP1-positive area), representing a clear quantitative separation between normal and tumor tissue. Similarly, we found a range of different PARP1 expression levels in our preclinical models, where increasing PARP1 expression resulted in higher PARPi-FL uptake and tumor-to-esophagus ratios of up to 20. Even the mouse model with the lowest PARP1 expression, however, had a six-fold higher PARPi-FL signal in the tumor than in the normal esophagus (**Fig. 2, C-F**). These ratios suggest it might be possible to delineate EAC in the esophagus with high contrast using a non-invasive endoscopic imaging approach.

Considering current clinical practice in EAC, including screening and surveillance protocols in patient populations with an increased risk to develop the disease—specifically in patients with Barrett’s esophagus (BE)—a non-invasive imaging approach could have a significant impact on screening and early detection. In addition to white light endoscopy, current surveillance protocols require invasive four-quadrant biopsies every 2-5 years. Sampling errors complicate this approach, since progression to cancer usually occurs in only a small fraction of the area presenting Barrett’s mucosa, while excisional quadrant biopsies cover only about 4% of the area affected by BE (*28*). Conceivably, complementing white light endoscopy with PARPi-FL imaging could guide biopsy site selection and enhance non-invasive identification of suspicious areas, including flat, small, and developing superficial lesions.

Existing diagnostic aids, including acetic acid, methylene blue (*29, 30*), or Lugol’s iodine (*31*) do not provide a biomarker-specific signal. As their benefit for biopsy site selection is limited, their clinical adoption rate is low. Clinical translation and integration of PARPi-FL imaging into regular white light endoscopy workflows can be optimized by using a topical application approach, as has been suggested for oral cancer imaging (*26*). To reach the esophagus, we tested an alternative route of administration using a balloon applicator. Recently, balloon-based devices have been introduced to retrieve cells (*32*) or DNA (*33*) from the esophagus in order to detect BE and EAC. In our case, we deployed a balloon-based applicator with highly absorbent Lubrizol stripes, which are collapsed inside of folds when the balloon is deflated but exposed and pressed onto the esophageal surface upon inflation. The pig model we used generally resembles the human esophagus’s anatomy and size. However, Yucatan pigs were found to have a considerable number of esophageal mucus glands, which do nevertheless express PARP1. By measuring PARPi-FL uptake in relation to gland depth, we were able to show penetration and specific binding of PARPi-FL up to 800 µm into the tissue within 3 min of application. These results corroborate the small molecule’s ability to penetrate well beyond the epithelial layers where EAC originates. Thus, our results indicate that specific, PARPi-FL-based detection of superficially presenting cancerous lesions in the esophagus could not only be feasible, but also clinically beneficial.

Novel methods for early detection, surveillance, and surgical guidance are also urgently needed in oral and oropharyngeal cancer. The incidence of oral and oropharyngeal cancer is increasing, while survival has shown no or only modest improvement over the last three decades (*34*). A main contributor to disease-related mortality is tumor stage at the time of diagnosis. Although the oral cavity is easily accessible for regular inspection, about two-thirds of all patients in the US present with advanced-stage disease, resulting in an overall five-year survival rate of about 50%, as compared to over 80% if diagnosed at an early stage. Another complicating factor for improving the standard of care is that socioeconomic groups with high oral cancer incidence are also the least likely to have dental or health insurance, further delaying diagnosis of their often asymptomatic disease (*35, 36*). The global situation is now more dire than ever, with respect to both incidence and number of associated deaths (300,000 and 145,000, respectively (*37*)). In India, where oral cancer is the third most common malignancy in men, the disease often remains untreated for extended amounts of time. Here, it is less HPV or smoking/drinking that drives the disease, but rather the consumption of paan/gutka (chewing tobacco), both common mild stimulants that contain betel leaf and areca nuts. Very few patients in India survive if their disease is discovered with regional or distant metastases (18% and 1.1%, respectively).

Currently, no diagnostic adjuncts have shown a proven clinical benefit for detection or surveillance of oral cancer. However, considering the various emerging optical imaging methods (*11*), we believe that molecularly specific fluorescence contrast, as provided by PARPi-FL, combined with novel analytical devices and methodologies, has the clear potential to improve delineation and detection of oral and oropharyngeal cancer. One such novel combination is PARPi-FL brush cytology. Brush cytology, also known as exfoliative cytology or brush biopsy, is a minimally invasive technique in which cells are collected between the surface and basement membrane of the epithelium, rather than using invasive incisional biopsy, followed by cytopathological evaluation (*7, 38*). Despite extensive clinical evaluation, adoption into clinical practice is still limited, partially due to the high cost of cytopathology as well as mixed reports about its sensitivity and specificity (*39*).

We evaluated whether a combination of brush biopsy and PARPi-FL staining is feasible in oropharyngeal cancer to identify and quantify tumor cells in a tissue sample. Analysis of PARP1 expression via IHC quantification in oropharyngeal cancer biospecimens revealed consistently high PARP1 expression—even higher than in oral and esophageal cancer—while normal epithelial expression resembled other physiological backgrounds. Located behind the oral cavity, oro- and hypopharyngeal cancers are less accessible to visual inspection than other oral cancer types, such as tongue cancer. Brush cytology, however, could be easily performed in this location. To establish a PARPi-FL staining method for brush cytology-like samples, we prepared a single-cell solution, as it would be derived during the brush cytology process, from solid xenograft tumor samples with known high PARP1 expression (FaDu (*26*); OE19, **Fig. 2**). Instead of cytopathological evaluation, we then stained this complex tissue sample with PARPi-FL and analyzed the percentage of PARPi-FL-positive cells using flow cytometry. We found a significantly higher number of PARPi-FL-stained cells in both xenograft models than in non-neoplastic tongue tissue samples. Importantly, due to the homology of human and murine PARP1, PARPi-FL targets both human and murine tissues. Considering PARP1 IHC results, a baseline level of PARPi-FL-positive cells in normal tongue samples was not surprising, since PARP1 expression in the epithelium’s basal layer is well-established.

These proof-of-principle results warrant further studies on combining PARPi-FL with brush cytology to establish baselines for PARPi-FL-positive cells in preclinical and human samples. Furthermore, a combination of several readouts, including the PARPi-FL-positive population, staining intensity, and cell size and granularity (forward and side scatter) could further increase the accuracy of this approach. A flow-based technology has the advantage of investigating suspicious lesions in a minimally invasive, quantitative, and automatable approach. In addition to pharyngeal cancer, application of PARPi-FL brush cytology is promising in all locations where brush cytology is currently explored, including other sites inside the oral cavity (*40*) and esophagus (*41, 42*).

While PARPi-FL brush cytology is a technology tailored for the screening and surveillance setting, PARPi-FL tissue staining is a promising approach for both pre- and intraoperative diagnostics. In several patient datasets from the US (Memorial Sloan Kettering Cancer Center; MSK) and India (Mazumdar Shaw Medical Center, Narayana Health; MSMC), we were able to establish that PARP1 expression at the epithelium undergoes statistically significant increases during disease progression. Considering the potential clinical impact, our most essential finding was that in severe dysplasia/carcinoma in situ and invasive carcinomas, PARP1 expression was significantly increased when compared to all other degrees of dysplasia and normal epithelium. Our results indicate that PARP1-based diagnosis could differentiate benign and potentially malignant lesions from normal epithelium and cancerous lesions that require surgical treatment, allowing for surveillance and chemoprevention to reduce the risk of cancer development (*43*).

Such results suggest measurement of PARP1 expression via PARPi-FL has the potential to identify different disease stages, including early stages. Despite a strong statistical separation of advanced disease from normal tissue, a certain overlap can be observed between groups, which we attribute to i) inter-patient heterogeneity of PARP1 expression and ii) imperfect histopathologic diagnostic separation of different stages of dysplasia. This is supported by our data where only paired data on PARP1 expression are analyzed – here, no intra-patient overlap between tumor and normal epithelium was observed. PARP1 IHC quantification also showed that surgical tumor margins were clearly delineated by PARP1 expression. This is significant, since almost all oral cancers are surgically treated. Complete removal, defined by clear margins, is a major determining factor for local recurrence (*3, 44, 45*). The current gold standard for negative tumor margins is clearance greater than 5 mm on permanent histopathology (*46*), which is not available until days after surgery. The only method currently available to surgeons for intraoperative feedback before surgical closure and reconstruction is frozen section analysis of surgical margins, which are sampled during surgery and sent to a pathologist. However, there is no standardized practice for frozen sections in head and neck cancer (*47*). Moreover, many view frozen sections critically, since accuracy is reduced as compared to paraffin sections, positive effects on outcome have not been reported, and the cost:benefit ratio is 20:1 (*48*). The PARPi-FL fresh tissue staining method reported here offers a promising alternative or addition to frozen section analysis. We have shown that PARPi-FL staining patterns of tumor biopsies and normal epithelium are distinct. We also identified several cases of focal invasion of tumor cells into an otherwise normal area based on the PARPi-FL signal. PARPi-FL staining patterns strongly represent PARP1 staining on IHC but were achieved within minutes of biopsy (as quickly as one minute). The rapid nature of the staining, in combination with back-table imaging, could provide feedback much faster than with frozen sections. If positive margins are found, the surgeon could perform re-excision and margin evaluation without delay. The vicinity to the surgery would also improve accuracy in anatomic sample orientation, a frequent source of error in frozen sections. We determined an excellent sensitivity and specificity of >95% for the identification of tumor and margin samples in fresh human biospecimens using blinded readers who received minimal training (10 min), supporting our claim that this method could be implemented in the clinic without the need for highly specialized personnel. A unique feature of our PARPi-FL staining method is the tissue-preserving nature of the staining, which would allow for integration into existing workflows, since PARPi-FL stained tissue would remain available for either frozen or permanent pathology or other techniques, including sequencing.

In addition to *ex vivo* imaging and microscopic evaluation, PARPi-FL holds promise for an array of further applications. We have shown feasibility of PARPi-FL *in vivo* diagnostics, based on our findings that intravenous injection provides a broad surgical window of at least 5-120 min post injection and can be detected with a handheld confocal endomicroscope. In a clinical scenario, this could enable a targeted search for a positive margin within the tumor bed. Currently, correlating the exact location of a positive margin, identified histologically *ex vivo*, to its counterpart in the surgical cavity is challenging due to significant shrinking and distortion that the tissue undergoes upon resection. In addition, we show feasibility of macroscopic first-in-human PARPi-FL staining and imaging with a topically applied PARPi-FL solution, which is currently under clinical evaluation (NCT03085147).

It is frequently suggested that a near infrared version of PARPi-FL would be advantageous over the green fluorescent variant. However, the fact that PARP1 is an intranuclear target puts several design constraints on a functional PARP1 imaging agent, which have been discussed previously (*49*). Longer wavelength PARP imaging agents were introduced *in vitro*, but the available data suggests pronounced effects on nuclear penetration and clearance affecting their *in vivo* performance (*50, 51*). In our opinion, overall pharmacokinetics of PARPi-FL and its ability to serve as a sensor for PARP1 outweighs the wavelength trade-offs. This is supported by numerous clinical studies involving green fluorescent dyes for surgical guidance, including the first first-in-human study of a molecularly targeted optical imaging agent (*52*).

Together, the versatility of PARPi-FL with respect to application, imaging settings, platforms, and technologies enables a combination of micro- and macroscopic evaluation techniques and thus a large range of applications, including screening, surveillance, biopsy guidance, fresh biopsy staining, *in vivo* diagnostic capability, and intraoperative margin delineation. Thus, PARPi-FL offers a whole host of diagnostic and intraoperative applications that can also be combined with each other, which could not be achieved with previously introduced contrast agents or label free diagnostic methods.

In the future, more extensive studies will be necessary to further confirm the clinical value of PARPi-FL, following guidelines specifically developed for fluorescent contrast agents to characterize their procedural benefit over existing practices (*27, 53*). Our next logical step toward standard of care clinical use of PARPi-FL will be blinded sampling of fresh tissue specimens at multiple clinical sites, with a focus on recruiting patients from different backgrounds and of varying disease stages. We do know that PARPi-FL staining is a highly sensitive and specific marker for discriminating tumor from non-tumor margins in cases with pathologically confirmed malignancy. While PARPi-FL could therefore conceivably be used in a surgical setting, its ultimate clinical value as a field diagnostic cannot be confirmed until a larger multi-national, multi-institutional study has been completed. Overall, this study represents a significant milestone towards clinical use of PARPi-FL, and it is critical for characterizing fluorescence-guided workflow and outcome benefits over existing practices.

## Online Methods

### PARP1 expression analysis

#### Sample collection of formalin-fixed, paraffin-embedded biospecimens

The analysis of PARP1 expression was conducted on banked or newly collected paraffin embedded biospecimens at MSK and MSMC. Specimen selection and collection was approved by the Institutional Review Boards (IRB) at MSK and MSMC and the Independent Ethics Committee (MSMC). All diagnoses were conducted by the pathology department on H&E stained sections from the same samples. We analyzed specimens of oral cancer (n=84 patients, 3 separate studies), oropharyngeal cancer (n=9) and esophageal cancer (n=7) patients (a more detailed description can be found in **Table S1**).

#### PARP1 staining

PARP1 IHC staining was carried out using a primary polyclonal rabbit anti-PARP1 antibody (sc-7150, 0.4 µg/mL, Santa Cruz Biotechnology). At MSK, the staining was carried out using the automated Discovery XT processor (Ventana Medical Systems) at the Molecular Cytology Core Facility as described previously (*26*). At MSMC, the same anti-PARP1 antibody was used (0.4 µg/mL, 1 h) in combination with the DAKO Real Envision kit (K5007, DAKO). Since the PARP1 antibody was discontinued by Santa Cruz, we tested and validated PA5-16452 (Invitrogen) as potential alternative.

#### Quantification of PARP1 in automated protocol

For PARP1 protein quantification, PARP1 IHC slides were digitalized using a MIRAX Slide Scanner (3DHISTECH, Budapest, Hungary). In surgical specimens, we separately analyzed tumor, normal epithelium and deep margin if present. On biopsy specimen (tumor or margin) we analyzed tumor on the tumor biopsy and normal epithelium and deep margin on the margin biopsy, if present. On at least 3 fields of view per area of interest (tumor, epithelium, deep margin) (20x magnification), PARP1 presence was quantified using ImageJ/FIJI. Diaminobenzidine (DAB) and Hematoxylin stainings were separated using the Color Deconvolution algorithm and appropriate threshold levels were set to measure the area of specific DAB staining (=PARP1 area) as well as hematoxylin staining (=tissue area) to calculate the relative PARP1 positive area on each image. The thresholds were kept constant within each dataset. We report means and standard deviation for each sample. For grouped analysis, we pooled the means of all samples. ROC curves were generated using GraphPad Prism 7.0.

#### Quantification in manual protocol

Additional manual scoring was conducted on PARP1 IHC cases from MSMC. Therefore, the staining intensity was evaluated on a scale of 1-4 (**Fig. S6A**) and multiplied with the % PARP1-positive cells (1-100%, visual evaluation), resulting in a final score of 1-400. All slides were scored by three people. We report the mean score for each slide and the ROC curve comparing benign/mild dysplasia to severe dysplasia/malignant cases.

### Synthesis of PARP1-targeted imaging agents

The synthesis of PARPi-FL and [^18^F]PARPi have been carried out as described previously (*54*). PARPi-FL was prepared as 1.5 mM stock solution in PEG300 and was diluted to its final concentration in 30%PEG300/PBS. [^18^F]PARPi was formulated in 0.9% sterile saline with 10% EtOH.

### Small animals and xenografting

All animal experiments were done in accordance with protocols approved by the Institutional Animal Care and Use Committee (IACUC) of MSK and followed the National Institutes of Health guidelines for animal welfare. For *in vivo* and *ex vivo* experiments, we grew subcutaneous xenografts in athymic nude mice (NCr-Foxn1nu, Taconic) using human pharynx squamous cell carcinoma (FaDu) and esophageal adenocarcinoma (EAC) cell lines (ESO51, OE33, SKGT4, OE19) cancer cells by injecting 2×10^6^ cells in a mixture of culture medium (50 µl) and Matrigel™ (BD Biosciences) (50 µl). Experiment were conducted 12-16 days after xenografting. We used 12 nude mice for imaging of EAC models, 9 nude mice for brush biopsy experiments, 10 nude mice for PARPi-FL fresh tissue staining optimization and 3 nude mice for rapid fresh tissue imaging on a strip-mosaic confocal scanner (Vivascope 2500). FaDu cells were ordered from ATCC, EAC cell lines were ordered from DSMZ. To prepare cells for xenografting, they were cultivated in a monolayer culture at 37°C in a 5% CO_2_ humidified atmosphere, following standard procedures. They were maintained in their respective growth medium (Roswell Park Memorial Institute medium for ESO51, OE33, SKGT4 and OE19) and minimum essential medium for FaDu), containing 10% (v/v) fetal bovine serum and 1% penicillin/streptavidin.

### Imaging of PARP1 expression in EAC models via PARPi-FL

To determine PARPi-FL uptake in EAC xenograft models, we intravenously injected PARPi-FL (75 nmol in 167 µl, 30%PEG300/PBS) in mice bearing ESO51, OE33, SKGT4 or OE19 xenografts (n=3/group). 90 min post-injection, animals were sacrificed and tumors, the esophagus and thigh muscle were excised and were imaged immediately in the epifluorescence system IVIS (PerkinElmer, Waltham, MA) using the standard filter set for GFP imaging. Autofluorescence removal and quantification were carried out as described previously (*26, 55*).

After epifluorescence imaging, the freshly excised whole tumors were imaged using a confocal microscope. Tissues were placed on a cover slip with a freshly cut surface facing the cover slip and all images were taken using identical settings and 488 nm laser excitation (LSM880, Zeiss, Jena, Germany). Subsequently, tumors were fixed in 4% Paraformaldehyde for 24 h and then paraffin embedded. H&E staining and PARP1 IHC staining and quantification were carried out as described above.

### Western Blot (EAC)

PARP1 protein expression was measured in ESO51, OE19, SKGT4 and OE33 cell lysates using Western blot analysis as described before (*56*). Briefly, proteins were isolated from cells and 30 µg of protein per sample were separated with SDS/PAGE gel electrophoresis and transferred to a Nitrocellulose membrane. Proteins were detected using antibodies specific for PARP1 (1:1000; sc-7150, Santa Cruz) and b-actin (1:2000; A3854, Sigma-Aldrich) with a corresponding horseradish peroxidase (HRP) conjugated secondary antibody (1:10,000, sc-2004, Santa-Cruz). Detection was performed using a chemiluminescent substrate (Thermo Scientific #34077, SuperSignal West Pico). Since distribution of the primary anti-PARP1 antibody was discontinued, we tested and validated PA5-16452 (Invitrogen) as potential alternative.

### Topical application and tissue penetration of PARPi-FL in the esophagus

A topical application strategy of PARPi-FL for the human esophagus was developed using a Yucatan mini swine (*Sus scrofa domesticus*), which reflects the human anatomy more closely than small animals. Pig experiments were carried out at CBSet, Inc. (Lexington, MA) and were approved by their IACUC. Topical application was realized using a balloon applicator (developed by Aero-Di-Namics, New York, NY; produced by Medical Murray, North Barrington, IL). The balloons are made of silicone with 12 Lubrizol stripes (electrospun Lubrizol tecophilic TN TG 500) and have a nominal diameter of 7 mm, which extends to 20-25 mm upon inflation (**Fig. 2G**).

The Lubrizol strips are protected inside of pleats during device delivery and removal and are only exposed when inflated. The Lubrizol stripes were loaded with 2 mL of a 1 µM PARPi-FL solution. The balloon was inserted into the anesthetized pigs esophagus in a deflated state under white light endoscopic guidance. Inflation led to exposure of the Lubrizol stripes and a pressure induced release of PARPi-FL onto the esophagus wall of the anesthetized pig for a duration of 3 min. 10 min after the application, the pig was sacrificed followed by necropsy and cryopreservation of the esophagus in 10×10 mm tiles. Cryosections were cut perpendicular to the esophagus surface and imaged using a confocal microscope to detect penetration depths and binding of PARPi-FL from the topical application. Adjacent slides were H&E stained to identify morphological structures. We used one pig and applied PARPi-FL to three different sites by re-loading and re-insertion of the balloon.

### PARPi-FL staining and flow cytometry of tissue derived cell suspensions

Brush biopsy samples, obtained from tissue derived cell suspensions were used for PARPi-FL staining. Specifically, FaDu (n=9) and OE19 (n=12) xenografts were grown in female athymic nude mice. Tumors and tongues were harvested, and single cell suspensions were obtained via tissue dissociation using the gentleMACS Octo Dissociator with Heaters (Miltenyi Biotech) in combination with a tissue dissociation kit (#130-095-929, Miltenyi Biotech), following the manufacturer’s instructions. Cells were then resuspended and stained with either 100 nM PARPi-FL, 100 µM olaparib mixed with 100 nM PARPi-FL or PBS for 1 hour at room temperature with occasional shaking, followed by a 1 hour wash in PBS at room temperature with occasional shaking. After resuspension in fresh PBS, cells were analyzed on a Fortessa flow cytometer (BD Biosciences). DAPI was added to exclude dead cells from the analysis. Side scatter and forward scatter were used to select single cells, excluding doublets, aggregates and debris. Fluorescence was measured in the DAPI channel (DAPI live/dead stain), FITC channel (PARPi-FL) and PE channel (to control for near FITC laser bleeding). The appropriate gates were selected on unstained and single stained controls (**Fig. S5A**). The cell populations were analyzed for the percentage of PARPi-FL positive cells.

### Fresh biospecimen staining

Optimization of the PARPi-FL fresh tissue staining protocol towards short staining time and specific nuclear accumulation was carried out using FaDu xenografts (n=5) (**Fig. S8**). The final protocol was used to stain fresh biopsies from oral cancer patients. 2 biopsies per patient (one biopsy was taken from the visible tumor and one biopsy at the 5 mm margin - see fig. 5A) were obtained pre-surgically under an IRB approved protocol. 13 patients were included in the study (see **Table S2** for histopathological diagnoses; all margins were tumor negative). One tumor sample was retrospectively excluded due to a non-malignant, non-dysplastic histopathological diagnosis (verrucous hyperplasia). For 3 patients, margin samples were not available, resulting in a total of n=12 tumors and n=10 margins. Upon receipt, tissues were photographed and divided into three equal parts, each containing mucosa and submucosa. One part was used for immediate fresh tissue imaging, one part was cryoconserved and one part was fixed in 4% PFA for H&E and PARP1 IHC. The fresh tissue staining protocol consisted of staining for 5 min in 100 nM PARPi-FL (in 30%PEG300/PBS, room temperature) and a 10 min wash in 30%PEG300/PBS. Samples were then transferred to ice-cold PBS containing 10 µg/mL Hoechst 33342 until confocal imaging (LSM880, Zeiss, Germany). Tissues were placed between two cover glasses (48×60 mm, Brain Research Laboratories) and a three-channel tile scan of the entire tissue (at least 2×2 mm) was captured (405 nm to detect Hoechst 33342, 488 nm to detect PARPi-FL and 543 nm to detect autofluorescence) using a 20x objective at 0.6x zoom in a scan time under 10 min.

The diagnostic value of PARPi-FL fresh tissue staining was determined in a blinded reader study (n=27 readers) (study design in **Fig. S10B**). Readers received a ∼10 min training on characteristic features of PARPi-FL staining in tumor and margin samples (**Supplementary Material X**). They were then shown the study set, consisting of 30 cases (n=12 tumors, n=10 margins, n=8 duplicates (4 tumors, 4 margins) to determine sensitivity, specificity and intrareader variability. For each case, readers were presented 1-5 images for 5 sec/image, followed by a 10 sec decision window to classify a tissue as tumor or margin (full study set in **Supplementary Material Y**; each case was assigned a random ID (random.org) and cases were presented in ascending numbers to the readers). Images were presented at 488 nm and 488/543 nm overlay (to facilitate autofluorescence identification), where available (**Fig. S10A**). Statistical values were calculated in R.

Biopsy samples from esophageal cancer patients (n = 5), acquired under an IRB approved protocol, were stained with 100 nM PARPi-FL for 5 min, subjected to confocal microscopy scanning and were then fixed in 4% PFA, followed by paraffin embedding and H&E staining. H&E sections were evaluated by a pathologist.

### Fresh biospecimen imaging with a strip-mosaic scanning confocal fresh tissue scanner

To test the feasibility of tissue scanning at a faster speed than standard confocal microscopy, we tested a confocal scanner that was specifically designed to create high-resolution strip-mosaics of whole tissues and features a 488 nm laser excitation (Vivascope 2500, Caliber ID, Andover, MA). Following PARPi-FL staining of FaDu xenografts (0, 100, 250 nM PARPi-FL for 10 min) and human biospecimens (100 nM PARPi-FL for 10 min), tissues were placed on the specimen holder and images of a 15×15 mm field-of-view were acquired in about 150 sec. Images were compared to regular confocal microscopy and PARP1 IHC of the same samples.

### Fresh biospecimen imaging with a point scanning miniaturized confocal endomicroscope

In addition to scanning of excised biospecimens, imaging with a confocal endomicroscope would enable imaging within the surgical cavity/wound bed in real time during surgery. Therefore, we tested if a miniaturized point-scanning confocal endomicroscope (Optiscan, Mulgrave, Australia) can distinguish PARPi-FL stained tumor and normal biospecimens. The system uses a 488 nm laser with several bandpass and emission filters, which allow for identification of autofluorescence (**Fig. S14A**). The focal plane is adjustable between 0-400 µm depth (**Fig. S14B**). The same human biospecimens that were used for regular confocal imaging were also imaged using the View*n*Vivo to determine if PARPi-FL stained nuclei can be clearly identified. The images were compared to PARP1 IHC images from the same patient.

### PARPi-FL blood half-life and imaging after intravenous injection in a pig

In a Yorkshire pig (50-55 kg), anesthesia was induced and the pig was placed on a table in the supine position. PARPi-FL (0.05 mg/kg, 10 mL, 30% PEG300 in PBS) was administered intravenously and, subsequently, blood samples (1.0 mL) and tongue punch biopsies (3 mm) were taken at predetermined time points (5, 10, 30, 60, 90, and 120 minutes). The pig was euthanized with an intravenous overdose of pentobarbital sodium and phenytoin sodium (440 mg/ml). Tongue punch biopsies were stored in ice-cold PBS containing 10 µg/mL Hoechst 33342 until confocal imaging. The whole, fresh punch biopsies were imaged using a LSM880 confocal microscope (Zeiss, Germany) to identify PARPi-FL staining in the epithelial basal layer. Ice-cold acetonitrile (1.5 mL) was added to blood samples (1.0 mL) and vortexed for 30 s. After centrifugation for 10 min (4000 rpm), the supernatant (1.8 mL) was lyophilized overnight. Acetonitrile (200 uL) was added, vortexed for 15 s, and spun down for 5 min (4000 rpm). Then, supernatant (100 uL) of each time was analyzed in a black bottom-less 96-well plate using a UV/VIS plate reader (Spectramax M5, Molecular Devices).

### PARPi-FL first-in-human imaging

To conduct PARPi-FL first in-human imaging, we obtained IRB approval. Informed consent was obtained from each patient. The presented data are part of a clinical phase I/II trial (NCT03085147). The patient had histopathologically confirmed oral squamous cell carcinoma. Videos were acquired before and post contrast application using the Quest Spectrum platform (Quest Medical Imaging, Middenmeer, Netherlands) in combination with a laparoscope and a customized PARPi-FL optimized laser/filter system. The same instrument settings were used throughout the imaging procedure (30 ms exposure time, 100% laser power, gain: 25.5 dB). For contrast application, the patient gargled 15 mL of a 250 nM PARPi-FL solution (in 15% PEG300/PBS) for 60 sec, followed by a washing solution (15%PEG300/PBS) for 60 sec. No adverse events or discomfort were observed. Image processing did not involve autofluorescence removal. For quantification, still frames were selected from a pre- and postcontrast video and in each image 5 regions of interested were placed on tumor and non-tumor regions using ImageJ.

### PET Imaging with [^18^F]PARPi

Animals bearing EAC xenografts were injected intravenously with 160-230 µCi of [^18^F]PARPi 2 hours before PET/CT imaging. To acquire PET/CT images (Inveon PET/CT, Siemens) animals were anesthetized with 2% isoflurane and positioned on the scanner bed. PET data were collected for 5-10 minutes, followed by CT. Data analysis was carried out using the Inveon Research Workplace Software.

### H&E Histopathology after PARPi-FL staining

We conducted formalin fixed paraffin embedded (FFPE) histopathology evaluation following PARPi-FL staining. In this case, PARPi-FL staining and imaging was conducted in approximately 60-90 min from receiving the fresh human biopsy. Then, the biopsy was fixed in 4% PFA for 24 hours at 4°C and processed for paraffin embedding, sectioning and H&E staining.

### Preclinical staining optimization protocols

PARPi-FL staining was optimized on fresh FaDu xenograft tissue. Freshly excised tumor tissue was cut into small pieces (2-3 mm diameter) before staining. We tested different staining protocols, varying PARPi-FL concentration (50, 100, 250 nM PARPi-FL in 30%PEG300/PBS), staining times (1, 5, 10 min) and washing steps (0, 2, 10 min in 30%PEG300/PBS). Samples were then transferred to ice-cold PBS containing 10 µg/mL Hoechst 33342 until confocal imaging (LSM880, Zeiss, Germany). To collect images, tissues were placed on top of a cover glass (48×60 mm, Brain Research Laboratories) and instrument settings were identical for all images.

### Statistical analysis

Statistical analysis was conducted using Graphpad Prism 7, except for the blinded study readings, which were analyzed with R. The confidence intervals for the diagnostic values were computed using a robust variance estimate accounting for a reader effect using a GEE approach. We used the Wilcoxon test for analysis of paired samples, e.g., PARP1-positive area of tumor, epithelium, and deep margin. We used the Mann-Whitney test for analysis of unpaired samples, e.g., PARP1-positive area of different disease stages. We used an unpaired t-test for analyzing flow cytometry data and corrected for multiple comparison using the Holm-Sidak method, without assuming a consistent standard deviation (SD). Statistical significance was determined with alpha = 0.05. We specify which test was used and which level of significance was found for each result (*p<0.05, **p<0.01, ***p<0.001, ****p<0.0001). ROC curves were generated with Graphpad Prism 7. All *in vivo* and *ex vivo* experiments subjected to statistical analysis consisted of group sizes of at least three. If not stated otherwise, data are presented as mean ± standard deviation.

### Data availability statement

The authors declare that all data supporting the findings of this study are available within the paper and its supplementary information files. The associated raw data and step-by-step protocols can be made available from the corresponding author upon reasonable request. The full code of the ImageJ macro for automated analysis of PARP1 expression on IHC slides is available through request by the corresponding author.

## Supporting information

Blinded Study_Training Set

Blinded Study_Data Set

## Supplementary Data

### Supplementary figures

Fig. S1: PARP1 IHC of all human EAC biospecimens.

Fig. S2: PARP1 quantification method via color thresholding.

Fig. S3: [^18^]F-PARPi imaging of EAC xenograft-bearing animals.

Fig. S4: PARP1 IHC of all human oropharyngeal biospecimens.

Fig. S5: Flow cytometry gating and OE19 staining.

Fig. S6: Manual PARP1 IHC scoring.

Fig. S7: Paired analysis of PARP1 expression in human biospecimens.

Fig. S8: Optimization of PARPi-FL fresh tissue staining.

Fig. S9: Rapid staining of cryosections.

Fig. S10: Diagnostic value of rapid PARPi-FL staining on fresh biospecimens.

Fig. S11: Histopathology following PARPi-FL fresh tissue staining is unaltered.

Fig. S12: Focal margin detection and dysplasia detection using PARPi-FL.

Fig. S13: Biospecimen imaging with a backtable scanner optimized for whole tissues.

Fig. S14: Imaging using a miniaturized handheld confocal endomicroscope.

Fig. S15: Intravenous delivery of PARPi-FL in a pig.

### Supplementary tables

Table S1: Overview of n numbers for PARP1 IHC studies.

Table S2: Histopathological diagnosis of fresh biopsy tissues.

### Other supplementary data

Supplementary PDF file 1: Blinded study training set.

Supplementary PDF file 2: Blinded study data set.

## Acknowledgments

We thank Aditi Sahu and Rachel Giese for supporting the work at MSMC. We thank Jay Budrewicz for his support of the experiments at CBSet, and Nora Katabi for providing their expertise in histopathology. We also thank Violeta Dokic for assistance during clinical work with PARPi-FL. We thank the Molecular Cytology Core Facility, Radiochemistry & Molecular Imaging Probes Core Facility, and Flow Cytometry Core Facility at Memorial Sloan Kettering Cancer Center. We also thank Garon Scott and Leah Bassity for editing the manuscript. Finally, we thank the participants of the blinded study, including (in alphabetical order): Adam Schulman, Aditi Sahu, Alexander Bolaender, Christian Mason, Edwin Pratt, Chrysafis Andreou, Fay Nicolson, Jack Berry, Jeroen Goos, Junior Gonzales, Kelly Henry, Luke Carter, Manu Jain, Marlena McGill, Navjot Guru, Nick Sobol, Patricia Ribeiro Pereira, Rustin Mirsafavi, Sheryl Roberts, Sophie Poty, Stephen Jannetti, Troy Crawford, Veronica Nagle, Xiancheng (Lewis) Wu, and Ahmad Sadique.

## Author contributions

S.K., G.P., M.A.K., S.P., and T.R. conceived the study and designed the experiments. S.K., G.P., A.L.S., S.P.S., P.D.S.F., D.K.Z., C.B., R.G. V.S., P.B., N.H., R.D.R., A.S., and T.R. carried out the experiments and collected and analyzed the data. S.K., S.P.S., P.D.S.F., D.K.Z., A.S., S.P., and T.R. wrote IRB protocols. A.M., M.G., G.P., and S.K. conducted statistical analysis of the data. M.S. and M.A.K. contributed experimental or analysis tools. S.K. and T.R. wrote the manuscript. All authors carefully reviewed and approved the manuscript.

## Data and materials availability

All data are in the main manuscript or in the Supplementary Materials.

