## Supplementary Figures and Tables for "PARP1 as a biomarker for early detection and intraoperative tumor delineation in epithelial cancers – first-in-human results"

### Content Supplementary Materials

#### Supplementary figures

Fig. S1: PARP1 IHC of all human EAC biospecimens.

Fig. S2: PARP1 quantification method via color thresholding.

Fig. S3: [$^{18}$]F-PARPi imaging of EAC xenograft-bearing animals.

Fig. S4: PARP1 IHC of all human oropharyngeal biospecimens.

Fig. S5: Flow cytometry gating and OE19 staining.

Fig. S6: Manual PARP1 IHC scoring.

Fig. S7: Paired analysis of PARP1 expression in human biospecimens.

Fig. S8: Optimization of PARPi-FL fresh tissue staining.

Fig. S9: Rapid staining of cryosections.

Fig. S10: Diagnostic value of rapid PARPi-FL staining on fresh biospecimens.

Fig. S11: Histopathology following PARPi-FL fresh tissue staining is unaltered.

Fig. S12: Focal margin detection and dysplasia detection using PARPi-FL.

Fig. S13: Biospecimen imaging with a backtable scanner optimized for whole tissues.

Fig. S14: Imaging using a miniaturized handheld confocal endomicroscope.

Fig. S15: Intravenous delivery of PARPi-FL in a pig.

#### Supplementary tables

Table S1: Overview of n numbers for PARP1 IHC studies.

Table S2: Histopathological diagnosis of fresh biopsy tissues.

### Supplementary figures

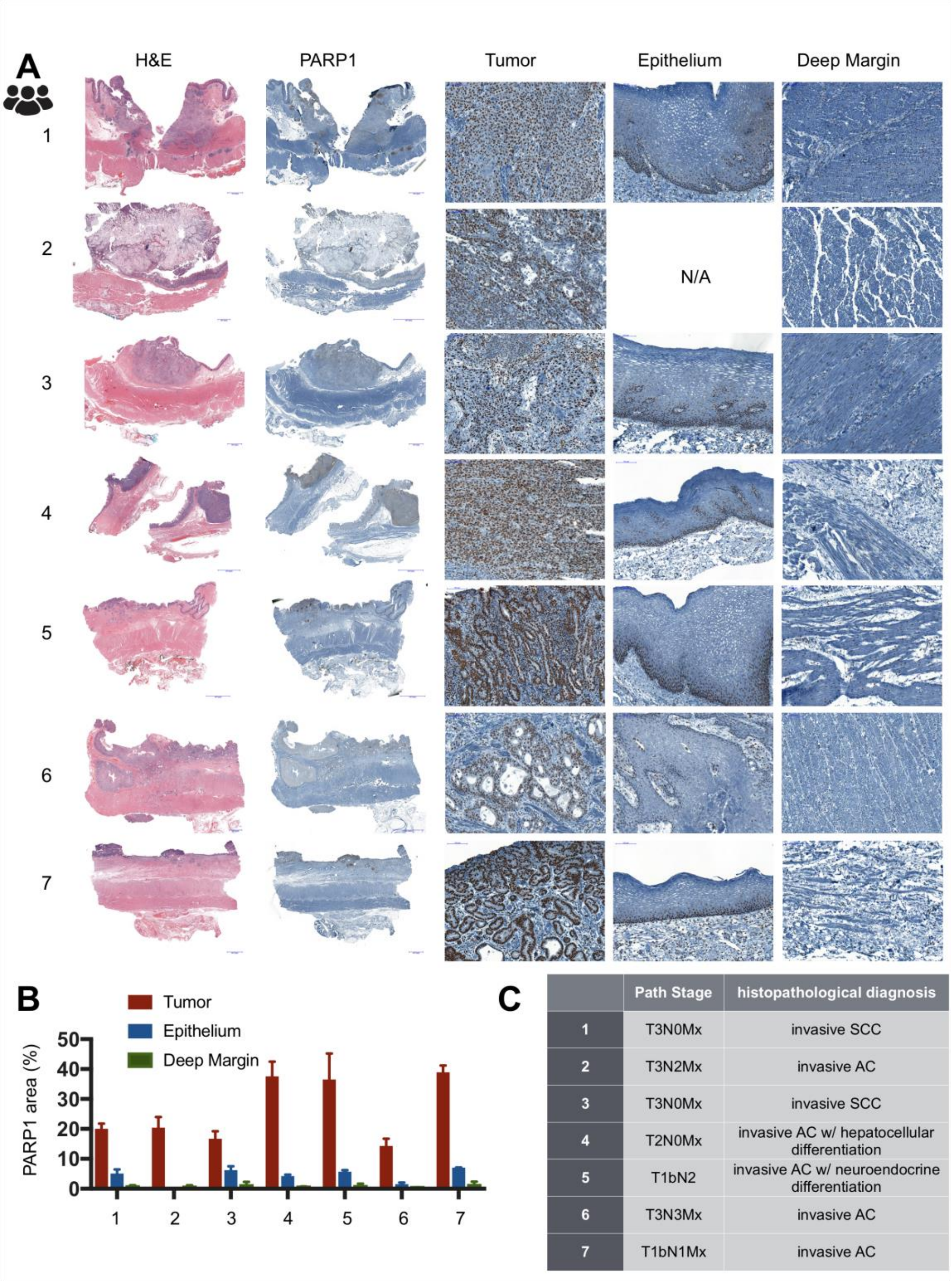


| | Path Stage | histopathological diagnosis |
|---|---|---|
| 1 | T3N0Mx | invasive SCC |
| 2 | T3N2Mx | invasive AC |
| 3 | T3N0Mx | invasive SCC |
| 4 | T2N0Mx | invasive AC w/ hepatocellular differentiation |
| 5 | T1bN2 | invasive AC w/ neuroendocrine differentiation |
| 6 | T3N3Mx | invasive AC |
| 7 | T1bN1Mx | invasive AC |

**Fig. S1. PARP1 IHC of all human EAC biospecimens. (A)** H&E and PARP1 IHC of human EAC biospecimens including an overview and high magnification images. **(B)** Quantification of the PARP1 positive area for each biospecimen. **(C)** Pathological stage and histological diagnosis for each patient

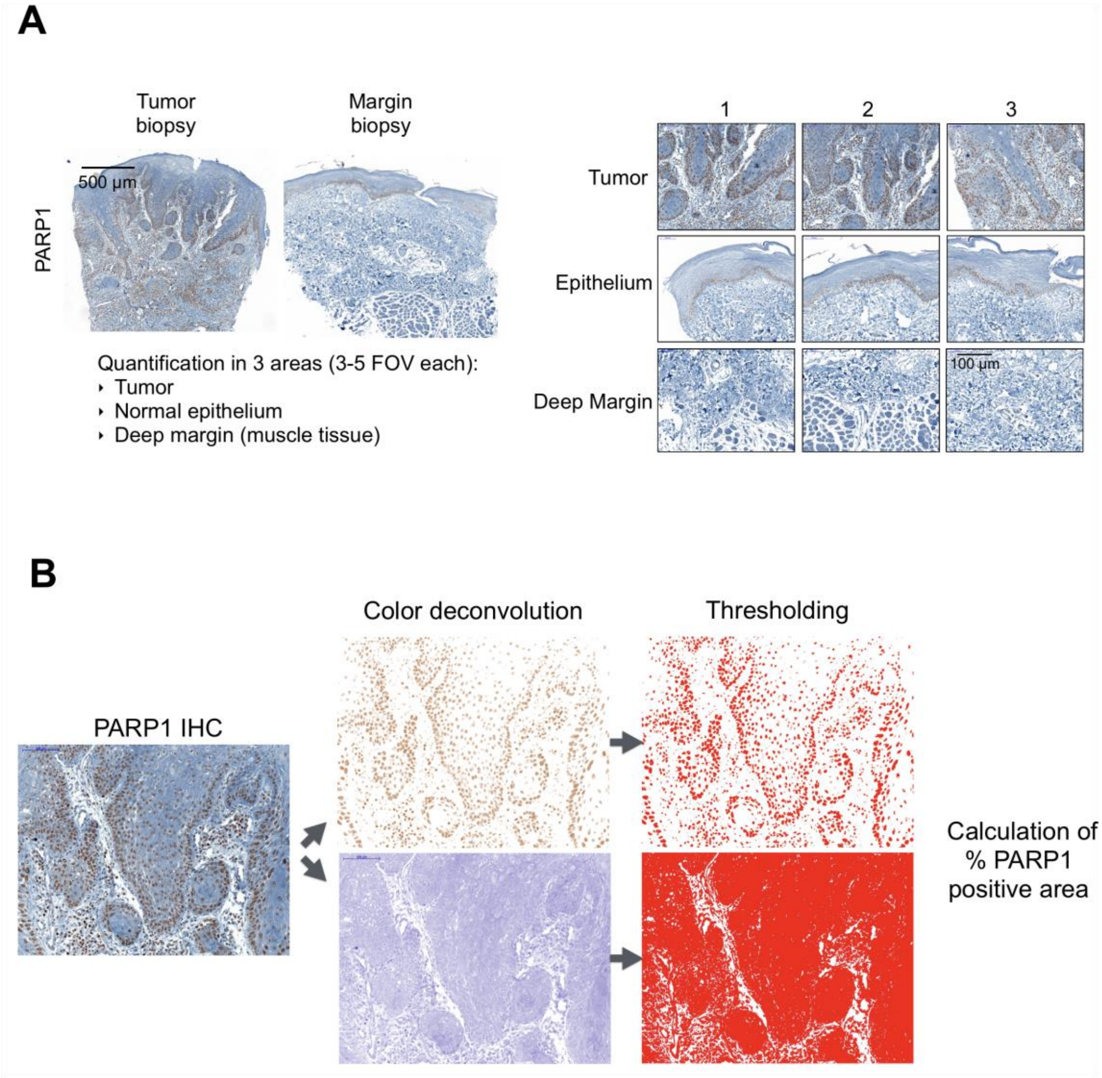


**Fig. S2. PARP1 quantification method via color thresholding. (A)** We quantified PARP1 expression in three areas for each patient (if available): tumor, normal epithelium and deep margin (submucosa, muscle tissue). From the scan of the entire tissue, 3-5 fields of view at 20x were selected for quantification. (B) Automated thresholding was conducted using an ImageJ macro. Here, following color deconvolution into brown and blue, the brown area and blue areas were measured and the relative PARP1 (=brown) positive area was calculated (PARP1 positive area= blue area/brown area*100).

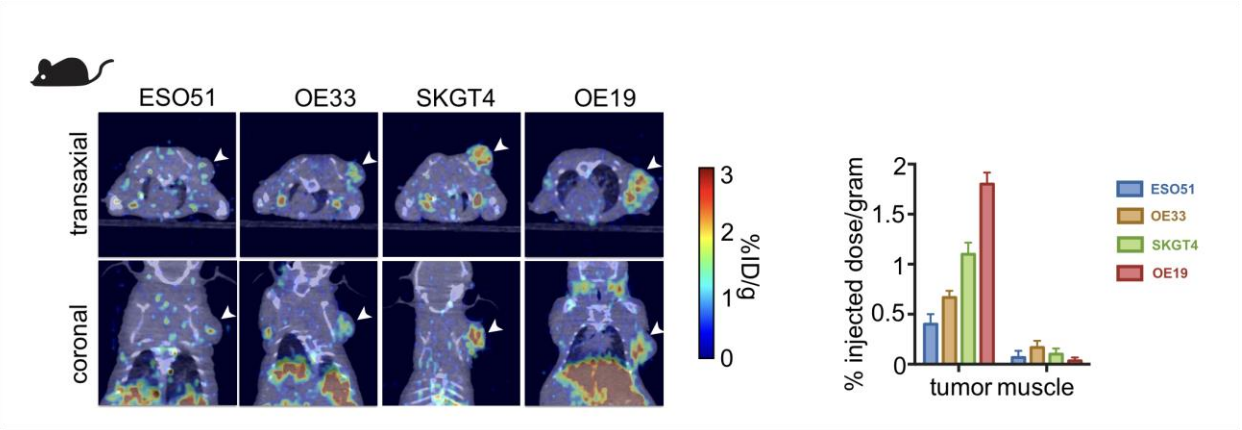


**Fig. S3. [$^{18}$]F-PARPi imaging of EAC xenograft-bearing animals.** 200 μCi [$^{18}$F]PARPi were intravenously injected into xenograft bearing animals (n=3/group) and imaged on an Inveon PET/CT 2 h post injection (arrows indicate tumor location). Quantification via ROIs from PET showed different levels of [$^{18}$F]PARPi uptake (ESO51<OE33<SKGT4<OE19) and low background uptake in muscle.

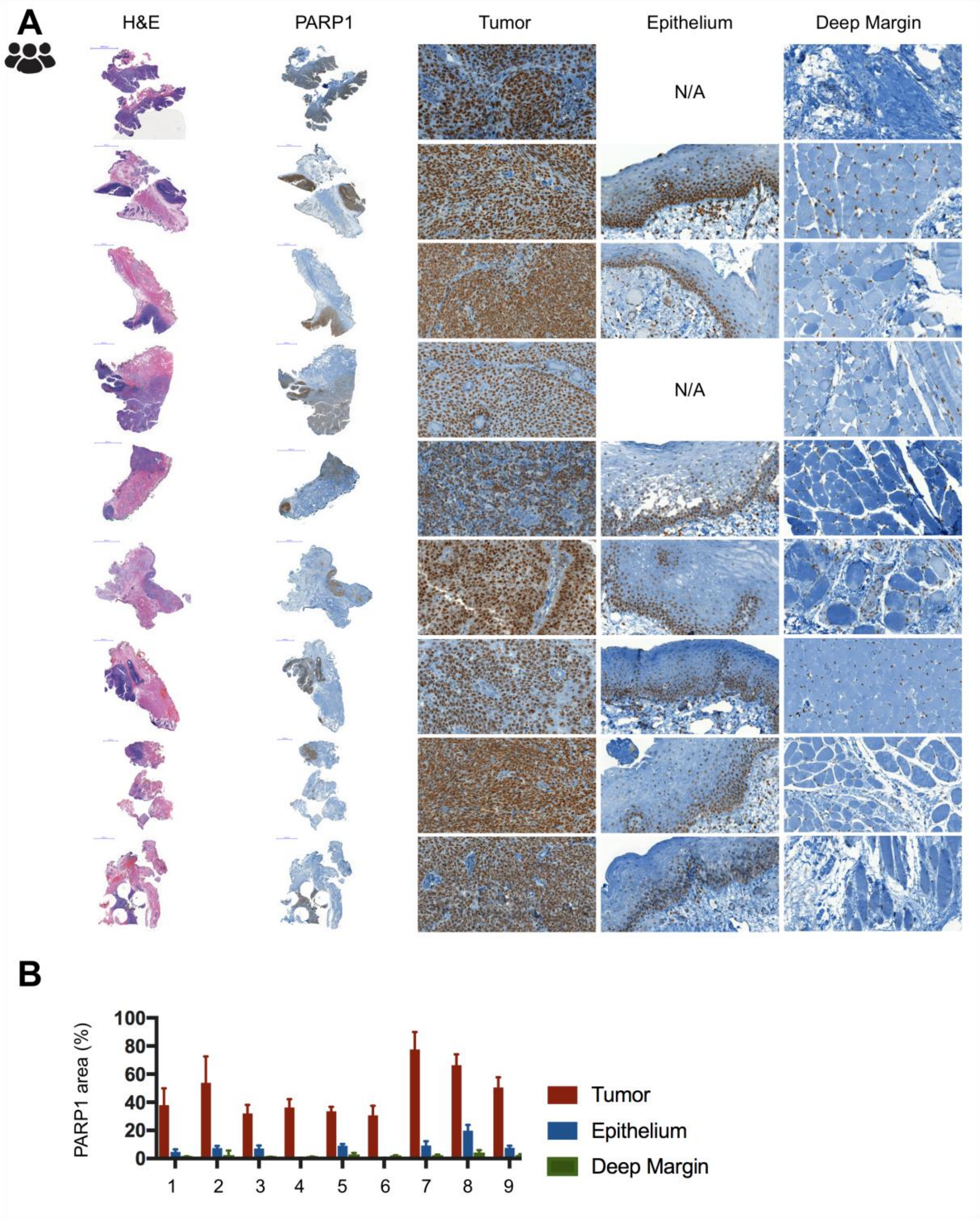


**Fig. S4. PARP1 IHC of all human oropharyngeal biospecimens. (A)** H&E and PARP1 IHC of human EAC biospecimens including an overview and high magnification images. **(B)** Quantification of the PARP1 positive area for each biospecimen.

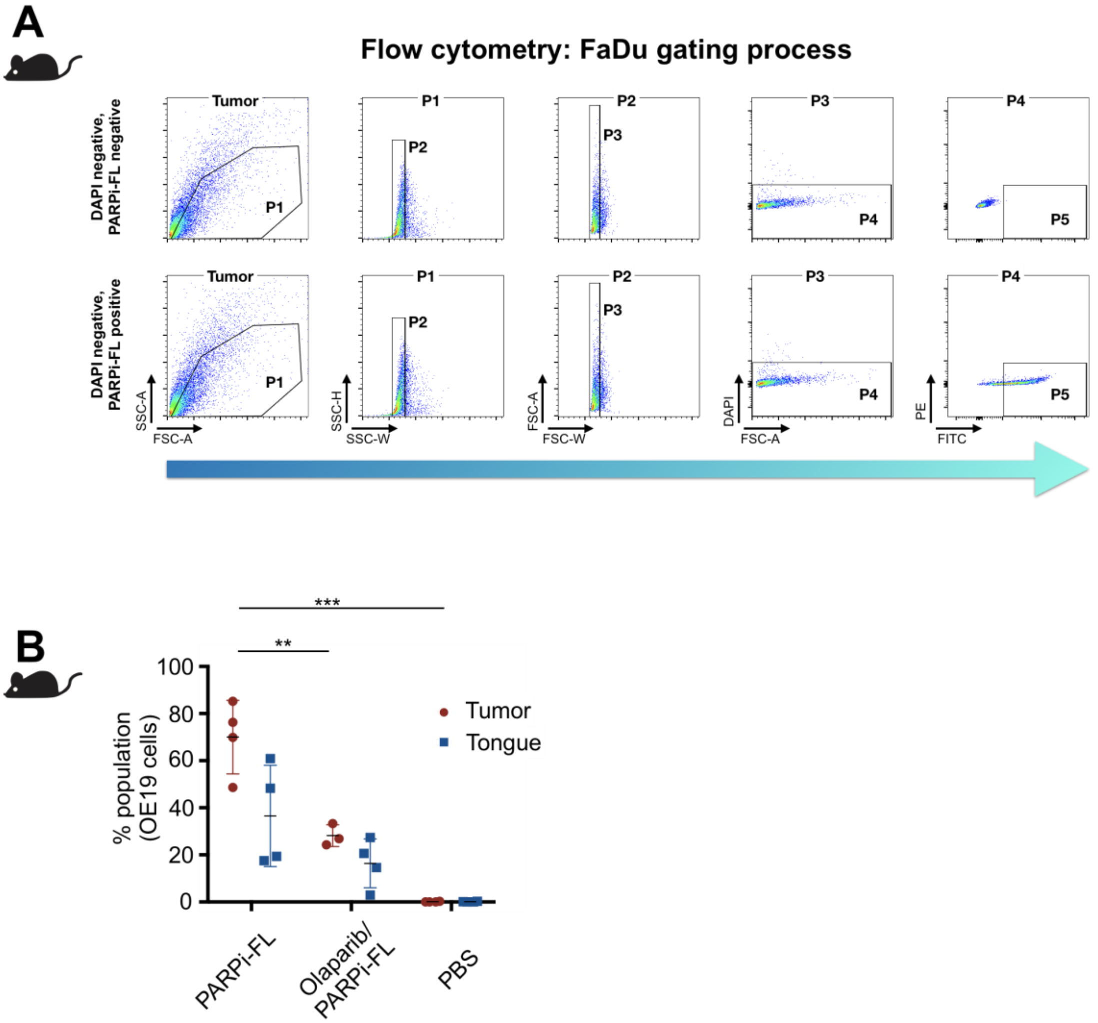


**Fig. S5. Flow cytometry gating and OE19 staining. (A)** Flow cytometry was performed on a BD LSRFortessa flow cytometer. Gates were determined before acquisition in a size-driven way. DAPI was used to exclude dead cells and the PE channel was used as a control for near-FITC laser bleeding. The percentage of PARPi-FL positive cells were used as readout. **(B)** Quantification of PARPi-FL flow cytometry of OE19 xenograft tissue (n=4 mice processed in 4 separate experiments). Significance was determined using an unpaired t-test. * p<0.05, *** p<0.001, ****p<0.0001. One data point in the Olaparib/PARPi-FL group (tumor) was identified as an outlier using the Grubbs' test (alpha=0.05) and was removed from the analysis.

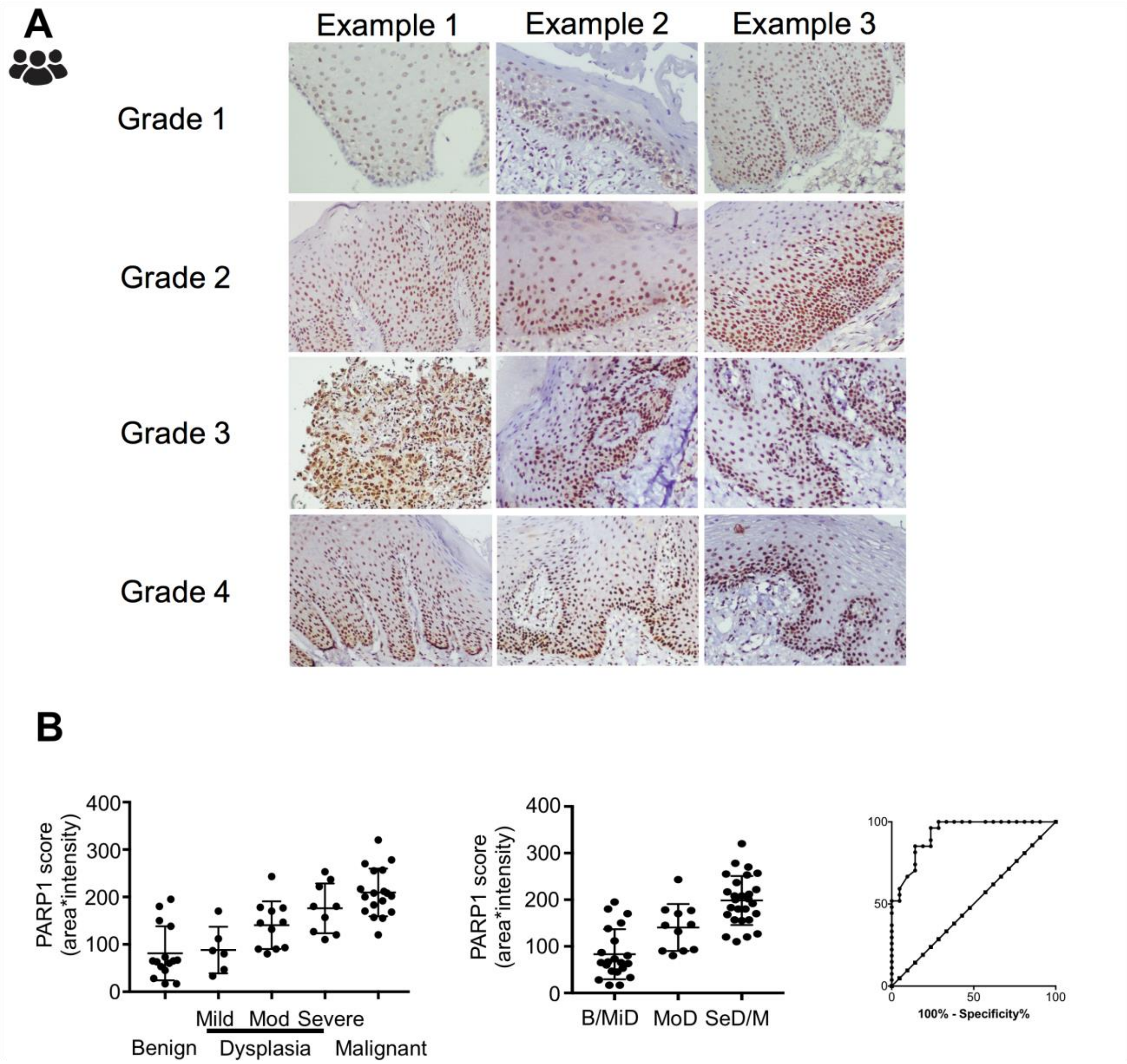


**Fig. S6. Manual PARP1 IHC scoring**. (A) Example of different intensity scores. The PARP1 score was calculated as: area (in %) × intensity grade. The area was estimated visually by each reader. (B) PARP1 scores for each category, combined categories and ROC curve for B/MiD versus SeD/M. B-benign, MiD-mild dysplasia, MoD-moderate dysplasia, SeD-severe dysplasia, M-malignant.

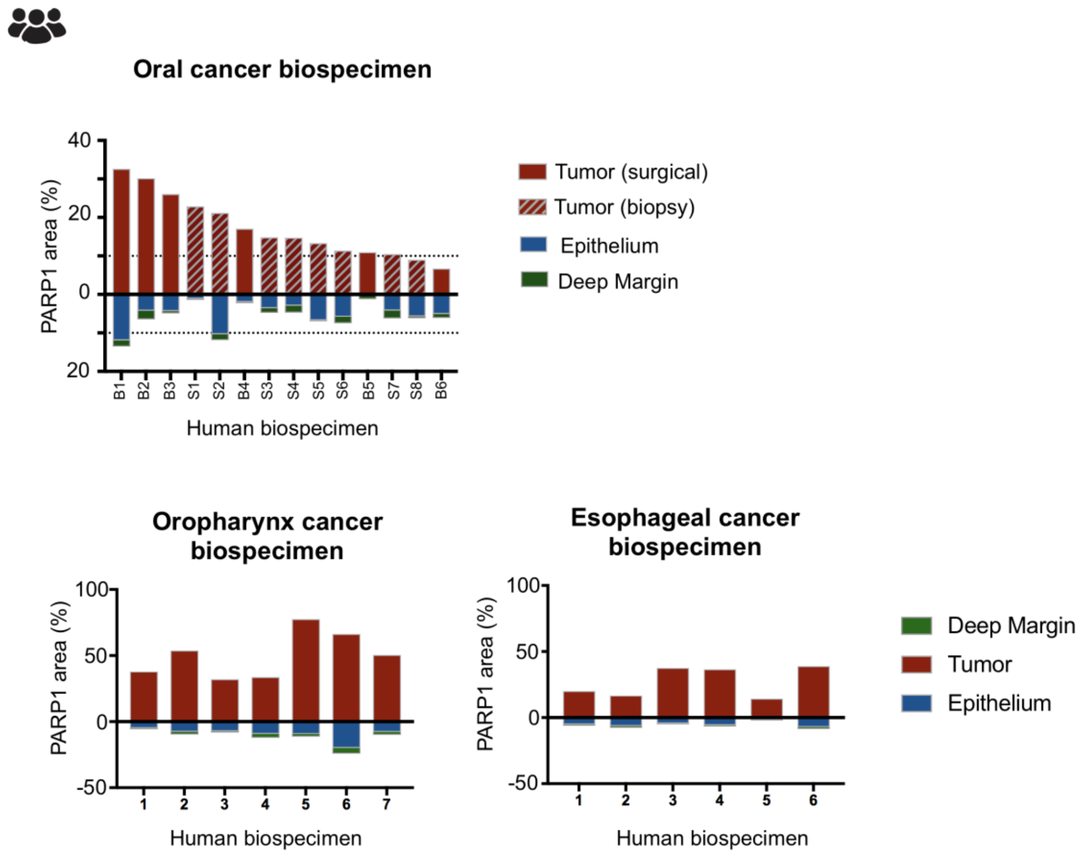


**Fig. S7. Paired analysis of PARP1 expression in human biospecimens.** Analysis of all biospecimen (oral, oropharyngeal and esophageal cancer) that had paired values (% PARP1 positive area, obtained through automated color thresholding) for tumor, epithelium and deep margin showing that expression in the tumor was higher than in epithelium or deep margin in every case. (A) oral cancer biospecimen, (B) Oropharynx cancer biospecimens, (C) Esophageal cancer specimens.

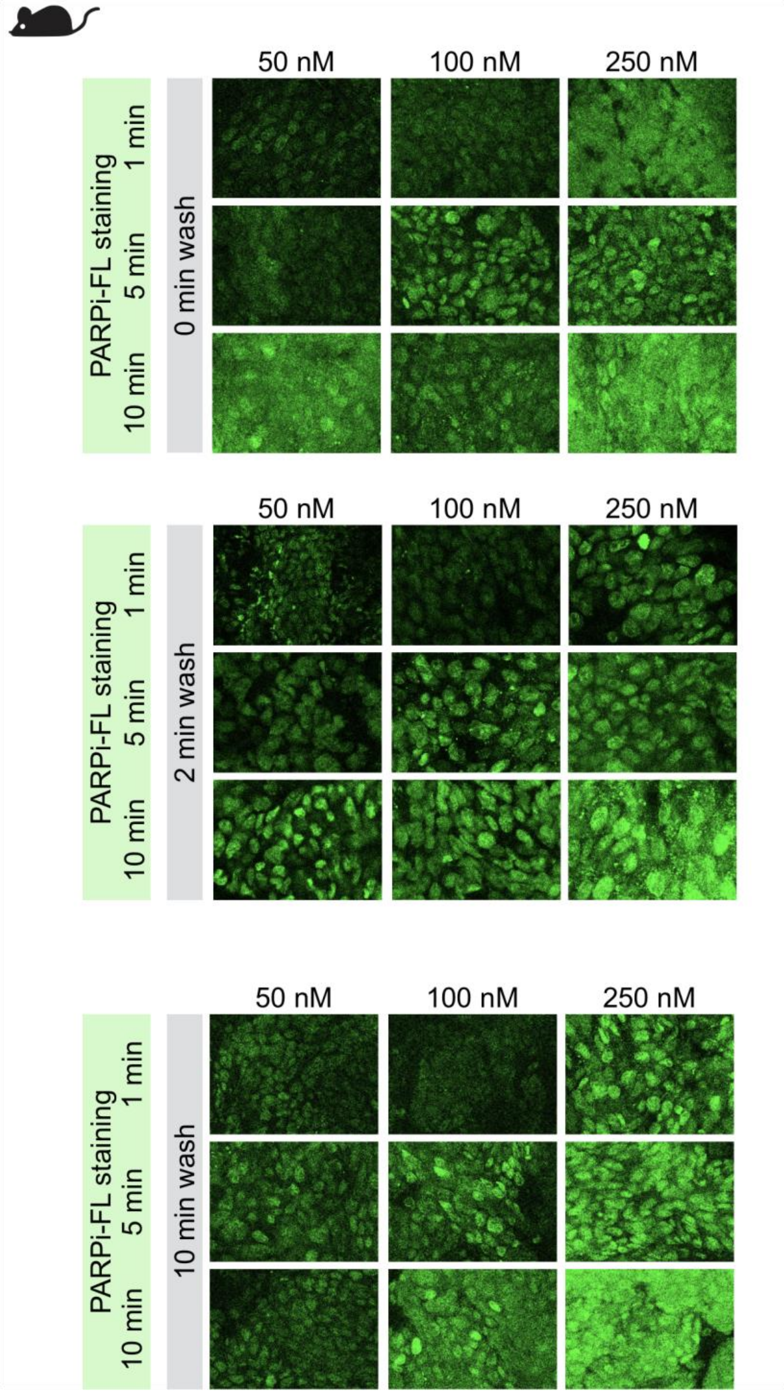


**Fig. S8. Optimization of PARPi-FL fresh tissue staining.** We tested different concentrations (50, 100, 250 nM), staining times (1, 5, 10 min) and washing steps (0, 2, 10 min) on FaDu xenograft tissue. We identified 100 nM staining for 5 min, followed by a 10 min wash as protocol that yielded desirable results (high nuclear staining intensity, low cytoplasmic and low background staining).

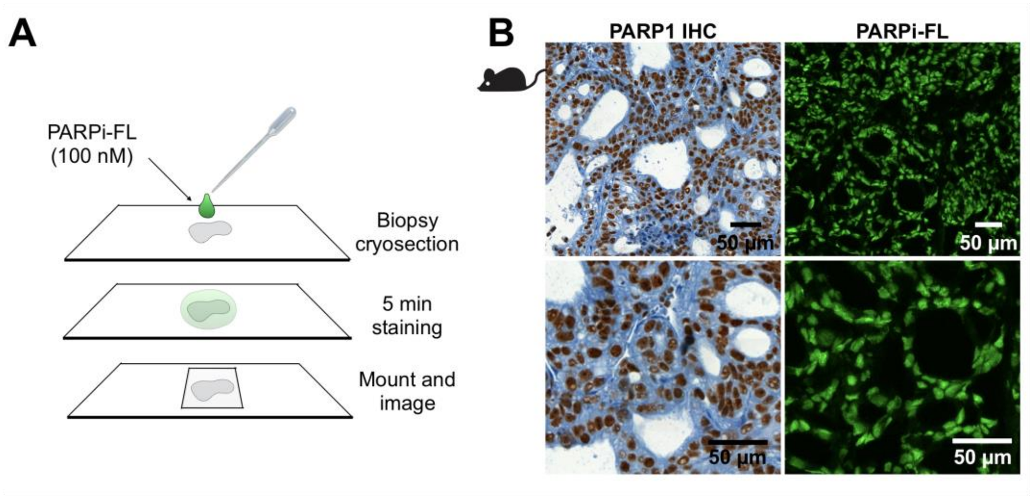


**Fig. S9. Rapid staining of cryosections.** We confirmed that rapid PARPi-FL staining, as developed for fresh tissue, also applies to cryosections. (A) On non-fixed cryosections of OE19 xenografts, we applied 100 nM PARPi-FL for 5 min, mounted the slide with Mowiol and conducted confocal microscopy within 24 hours. (B) PARPi-FL staining correlated with PARP1 IHC, confirming that rapid, specific staining of fresh tissue is also possible using cryosections.

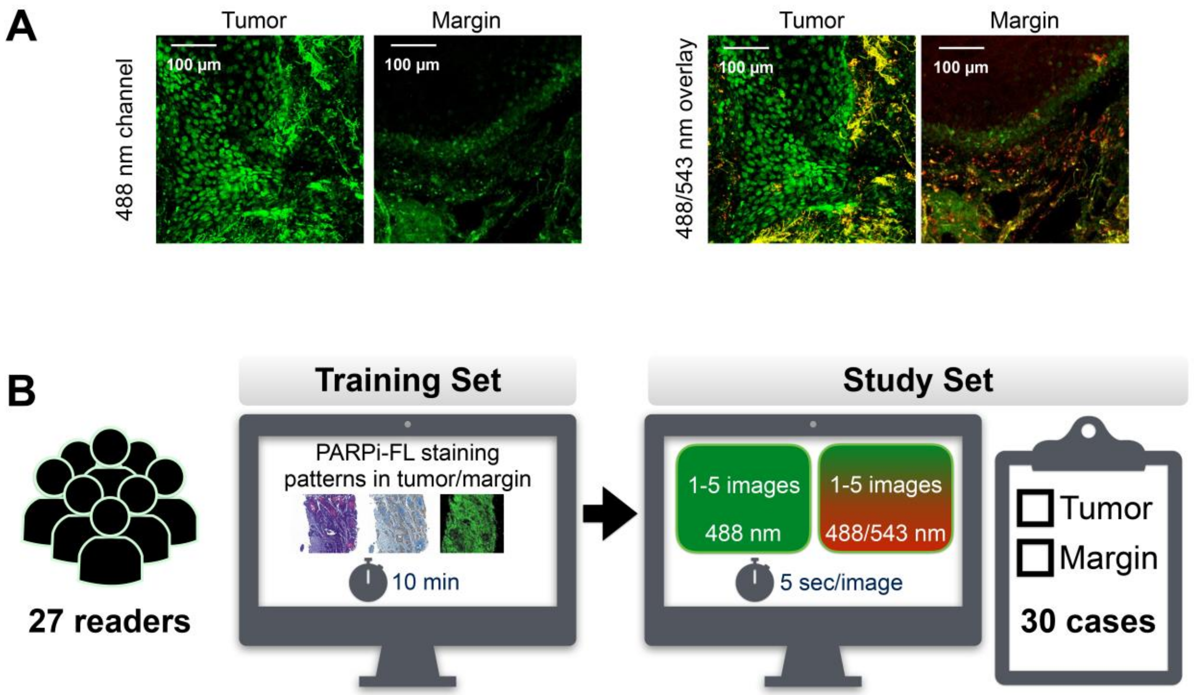


**C**

| Reference \ Readers | Margin | Tumor | Total |
|---|---|---|---|
| Margin | 361 | 17 | 378 |
| Tumor | 18 | 414 | 432 |
| Total | 379 | 431 | 810 |

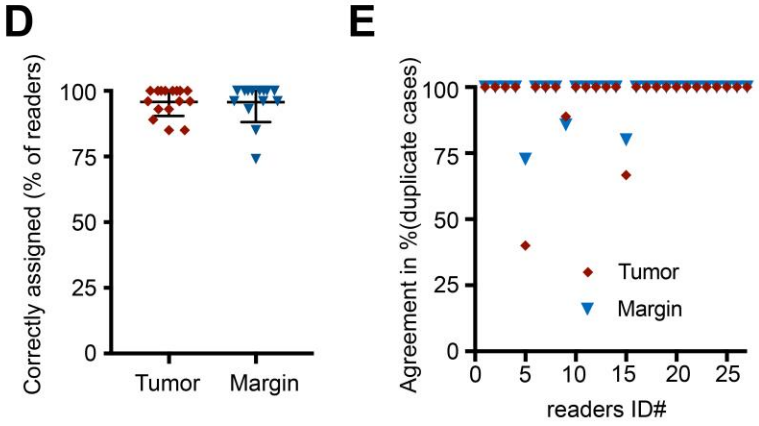


**Fig. S10. Diagnostic value of rapid PARPi-FL staining on fresh biospecimens. (A)** Examples of a tumor and margin image as green fluorescent only (488 nm excitation) and as overlay of 488/543 nm excitation to assess autofluorescence. Readers were presented both images (green only and green/red overlay) during the blinded reader study. **(B)** Study design: Readers (n=27) scored a total of 30 cases (n=12 tumors, n=10 margins, n=8 duplicates (4 tumors, 4 margins) as tumor or margins. Readers received a 10 min training slideshow with example stainings and corresponding histology. Then they were presented 1-5 images of each PARPi-FL stained biopsy as green fluorescent only (488 nm excitation) and as overlay of 488/543 nm excitation to assess autofluorescence. **(C)** Overall rating performance. A total of 810 scores (27 readers * 30 cases) were recorded. Readers correctly scored 414 of 431 tumor cases 361 of 379 margins. **(D)** Comparison of correctly assigned cases as % of readers for tumors and margin cases separately. **(E)** Intrareader agreement for pairs of original/duplicate cases (4 tumor and 4 margin tissues were presented to the readers mirrored and a 90° angle) without knowledge of the readers. Overall tumor agreement was 96.1% and overall margin agreement was 97.7%. Overall agreement was 97.2%.

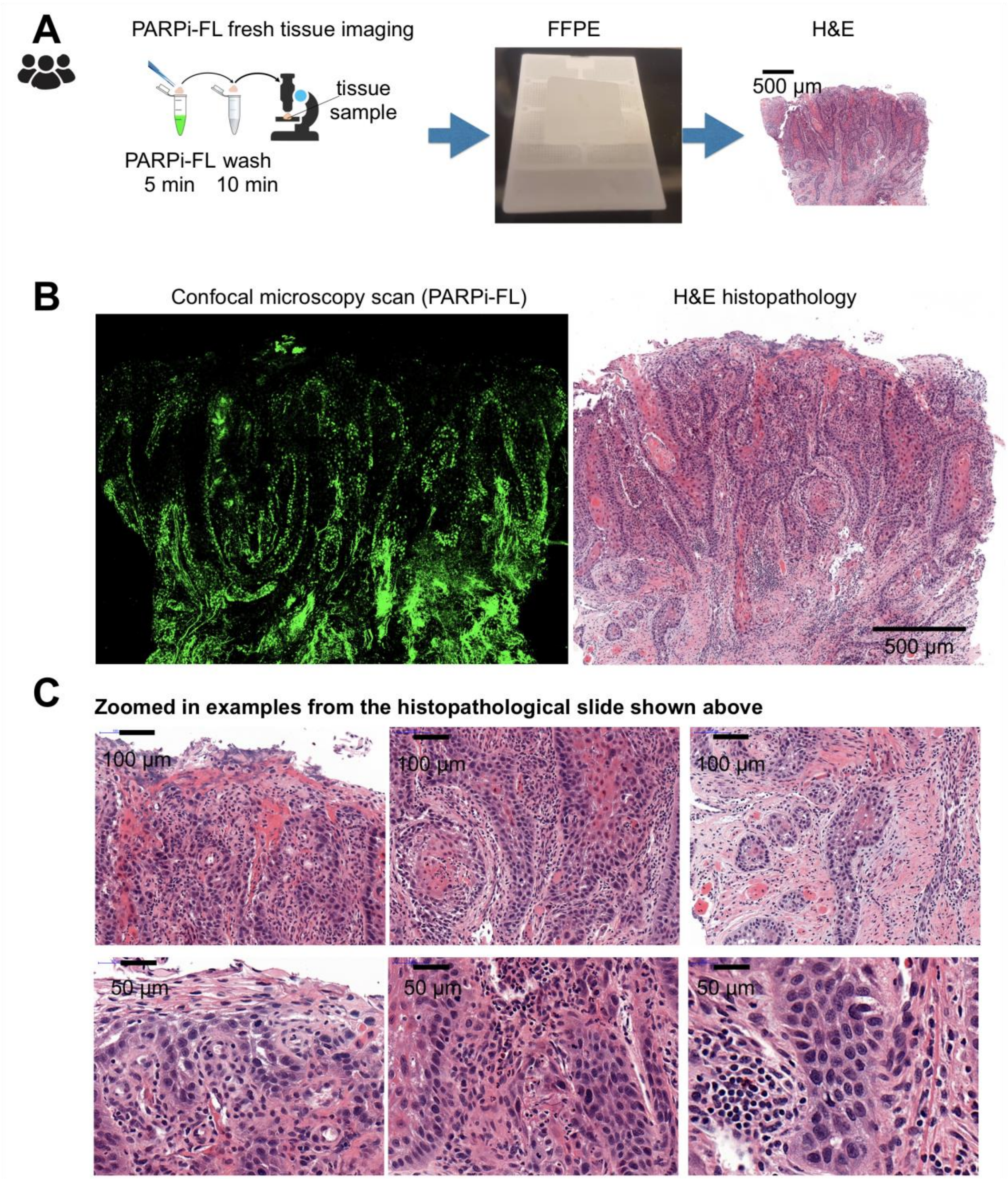


**Fig. S11. Histopathology following PARPi-FL fresh tissue staining is unaltered. (A)** To assess if PARPi-FL staining would influence the integrity or morphology of the fresh tissue, we conducted formalin fixed paraffin embedded (FFPE) histopathology evaluation following PARPi-FL staining. PARPi-FL staining and imaging was conducted in approximately 60-90 min from receiving the biopsy from the OR.

Then, the biopsy was fixed in 4% PFA for 24 hours at 4°C and processed for paraffin embedding, sectioning and H&E staining. **(B)** Comparison of the confocal scan of PARPi-FL stained fresh tissue and an FFPE H&E section of the same specimen. **(C)** High resolution zoomed in images show no signs of altered tissue morphology by conducting PARPi-FL staining and imaging before FFPE processing.

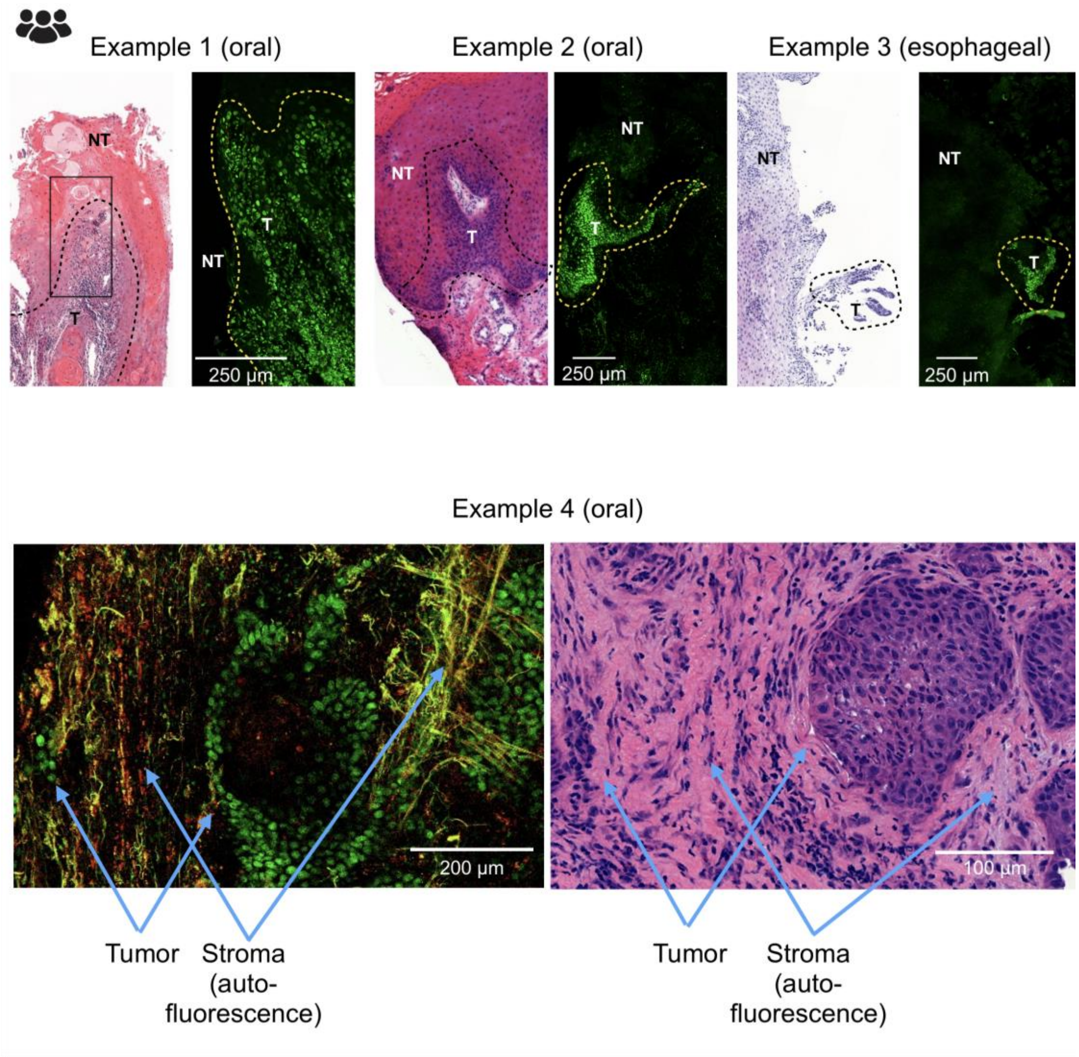


**Fig. S12. Examples for focal margin detection using PARPi-FL**. On several individual fresh tissue samples, we identified microinvasion of tumor cells into the epithelium and stroma, which was clearly delineated by PARPi-FL staining. Example 4 shows invasion into the stroma, which shows collagen related autofluorescence, which is easily distuingishable from the PARPi-FL signal by wavelength. green fluorescence-PARPi-FL, yellow/red: autofluorescence in tissue stroma.

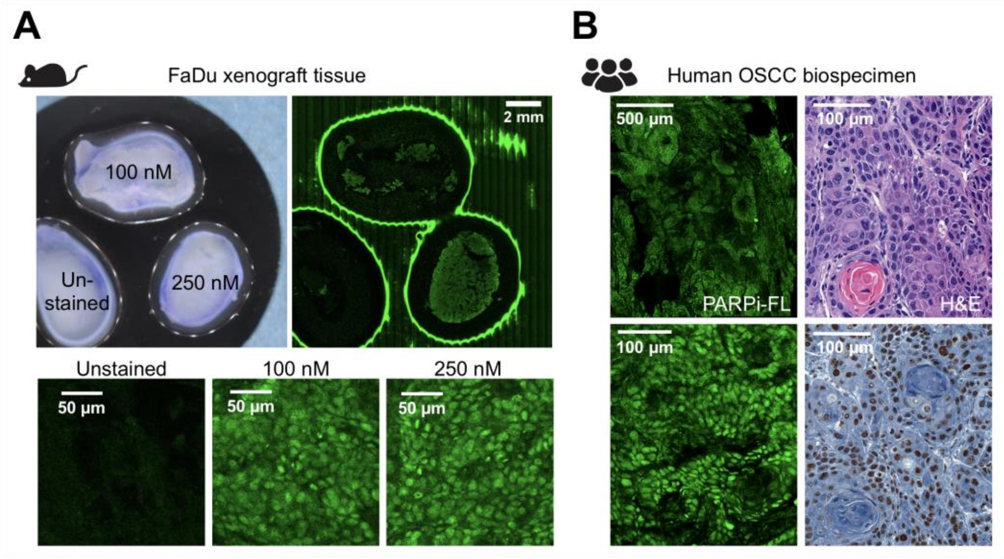


**Fig. S13. Biospecimen imaging with a backtable scanner optimized for whole tissues.** (**A**) Images of excised FaDu tumors, stained with 100 nM or 250 nM PARPi-FL, scanned with the Vivascope 2500 backtable scanner for whole tissues in comparison to unstained controls. (**B**) A patient sample stained with 100 nM PARPi-FL for 10 min and scanned on the Vivascope 2500 and corresponding H&E and PARP1 IHC from the same patient.

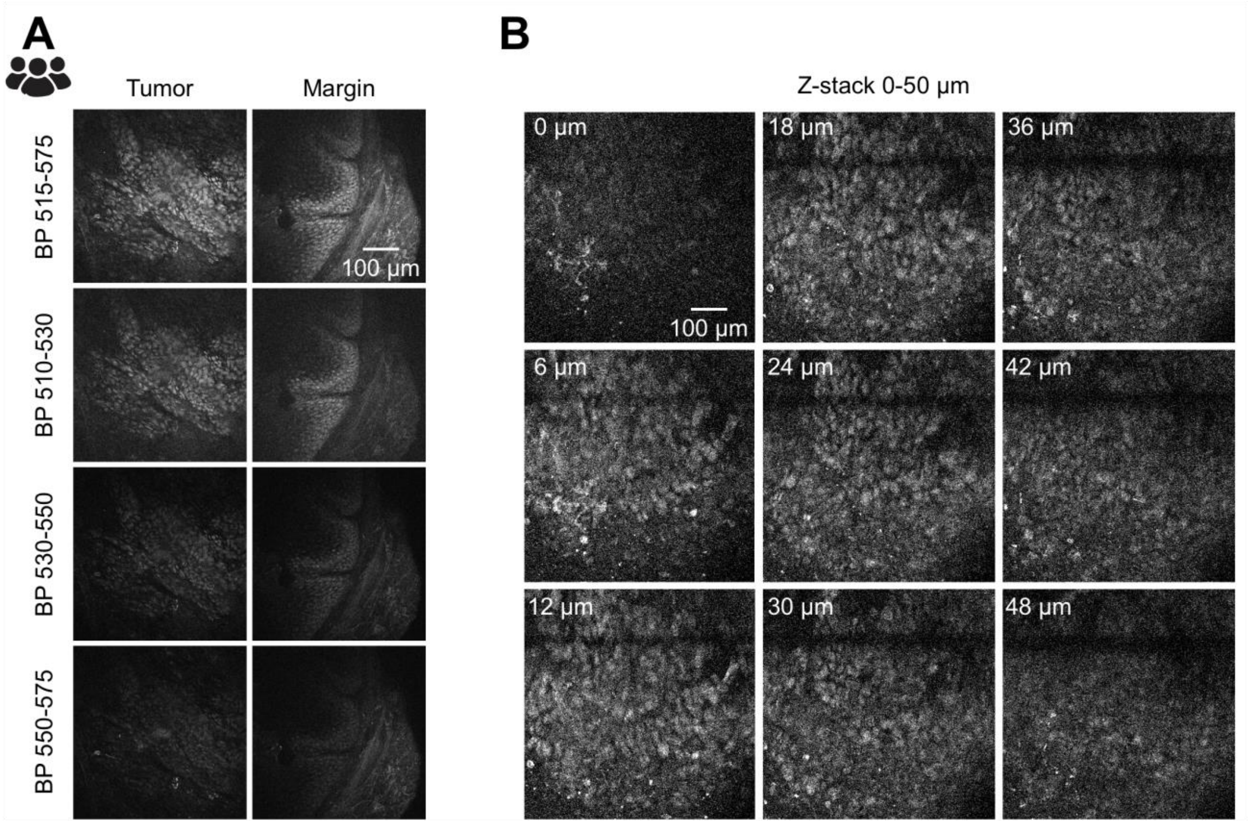


**Fig. S14. Imaging using a miniaturized handheld confocal endomicroscope.** In addition to a regular stand-alone confocal microscope, we also conducted imaging with a handheld confocal endomicroscope. **(A)** PARPi-FL is best visible in the bandpass (BP) filters 515-575 nm and 510-530 nm, while the signal is very low in BP 530-550 nm and BP 550-575 nm, corresponding to the emission maximum of PARPi-FL at 503 nm. **(B)** The point scanning technology of the confocal endomicroscope allows for adjustment of the focal plane between 0-400 µm. We were able to detect PARPi-FL in up to 50 µm depths after a 5 min topical staining of fresh tissue.

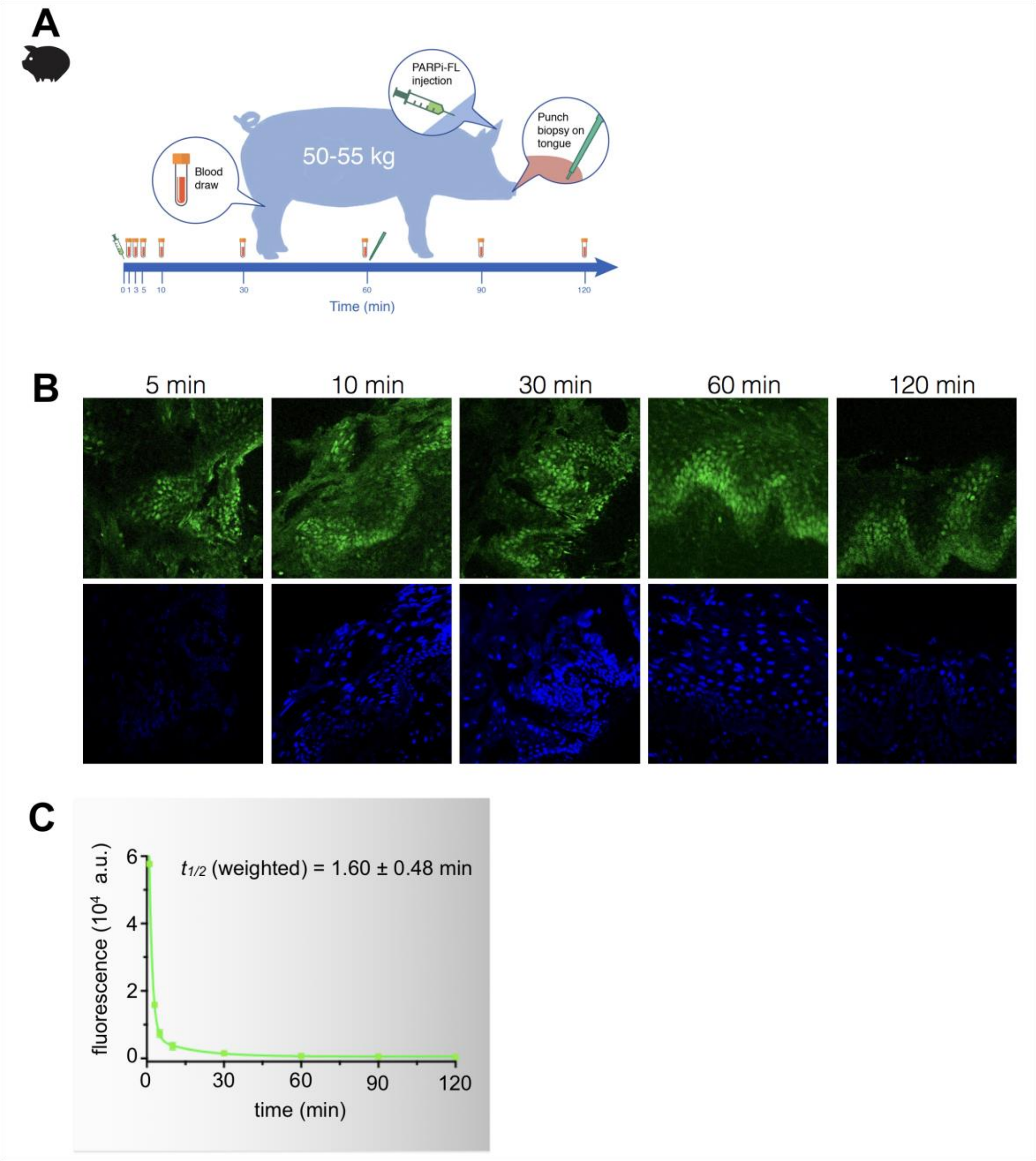


**Fig. 15. PARPi-FL detection after intravenous injection**. PARPi-FL was detected in the tongue epithelium of a pig at different time-points (5-120 min) post i.v. injection using punch biopsies. PARP1 expression was detected in the basal layer, where it occurs naturally. We also measured the blood half-life from blood draws taken at the same time-points. In absence of a tumor in the large animal, we used basal layer imaging as a surrogate for tumor associated PARP1 expression, which, according to our data, is

significantly higher compared to the basal layer, potentially leading to higher PARPi-FL uptake.

### ***Supplementary tables***

#### Table S1. Overview of n numbers for PARP1 IHC studies.

| | *Specimen type* | *Collection type* | *Patients* | *Number of specimen per patient* | *Number of complete data sets (tumor, epithelium, deep margin)* | *Number of data points* | *Results presented in Fig.* |
|---|---|---|---|---|---|---|---|
| *Esophageal cancer (MSK)* | Surgical | Retro-spective | 7 | 1 | 6 | 20 | Fig. 2A,2B; Suppl. Fig. 1A, 1B |
| *Esophageal cancer (MSK)* | presurgical | Pro-spective | 5 | 4-6 | NA | NA | Suppl. Fig. S12 |
| *Oropharyngeal cancer (MSK)* | Surgical | Retro-spective | 9 | 1 | 7 | 25 | Fig. 3A, 3B, Suppl. Fig. 4A, 4B |
| *Oral cancer (MSK)** | Surgical | Retro-spective | 12 | 1 | 8 | 31 | Fig. 4F, 4H |
| *Oral cancer (MSK)* | Presurgical biopsy | Pro-spective | 12 | 2 (tumor, margin) | 6 | 29 | Fig. 4G, 4H |
| *Oral cancer (MSMC)* | Diagnostic biopsy | Retro-spective | 60 | 1 | NA | 60 | Fig. 4A, 4B, 4C, 4D, Suppl. Fig. 6 |

*PARP1 IHC samples from this cohort were originally analyzed in (*1*). Samples were re-analyzed for this study, categorizing them into tumor/epithelium/deep-margin (compared to tumor/non-tumor) and different regions of interest were used for the quantification.

**Table S2. Histopathological diagnosis of fresh biopsy tissues.**

| | *Histopathological diagnosis* | ***Fresh tissue-Tumor*** | ***Fresh tissue - margin*** |
|---|---|---|---|
| 1 | Invasive SCC, keratinizing moderately differentiated | Y | NA |
| 2 | Invasive SCC, keratinizing moderately differentiated | Y | Y |
| 3 | Invasive well differentiated SCC, and moderate dysplasia in top | Y | Y |
| 4 | SCC in situ | Y | Y |
| 5 | Fragments of invasive SCC moderately differentiated, keratinizing | Y | Y |
| 6 | Fragment of invasive SCC, poorly differentiated, and granulation tissue | Y | Y |
| 7 | Invasive SCC, moderately differentiated, keratinizing, with PNI | Y | Y |
| 8 | Poorly differentiated SCC with spindle cell features | Y | NA |
| 9 | Verrucous hyperplasia | excluded | Y |
| 10 | Invasive SCC, well differentiated, keratinizing | Y | Y |
| 11 | Invasive SCC, moderately differentiated, keratinizing | Y | Y |
| 12 | Invasive SCC, moderately differentiated, keratinizing | Y | NA |
| 13 | Invasive SCC, moderately differentiated, keratinizing | Y | Y |
