## Supplementary material for "PARP1 as a biomarker for early detection and intraoperative tumor delineation in epithelial cancers – first-in-human results": Blinded Study_Training Set

### PARPi-FL staining of fresh biopsy tissues for identification of tumor and margin tissue

Blinded study

Training set

### Oral cancer - basic histology

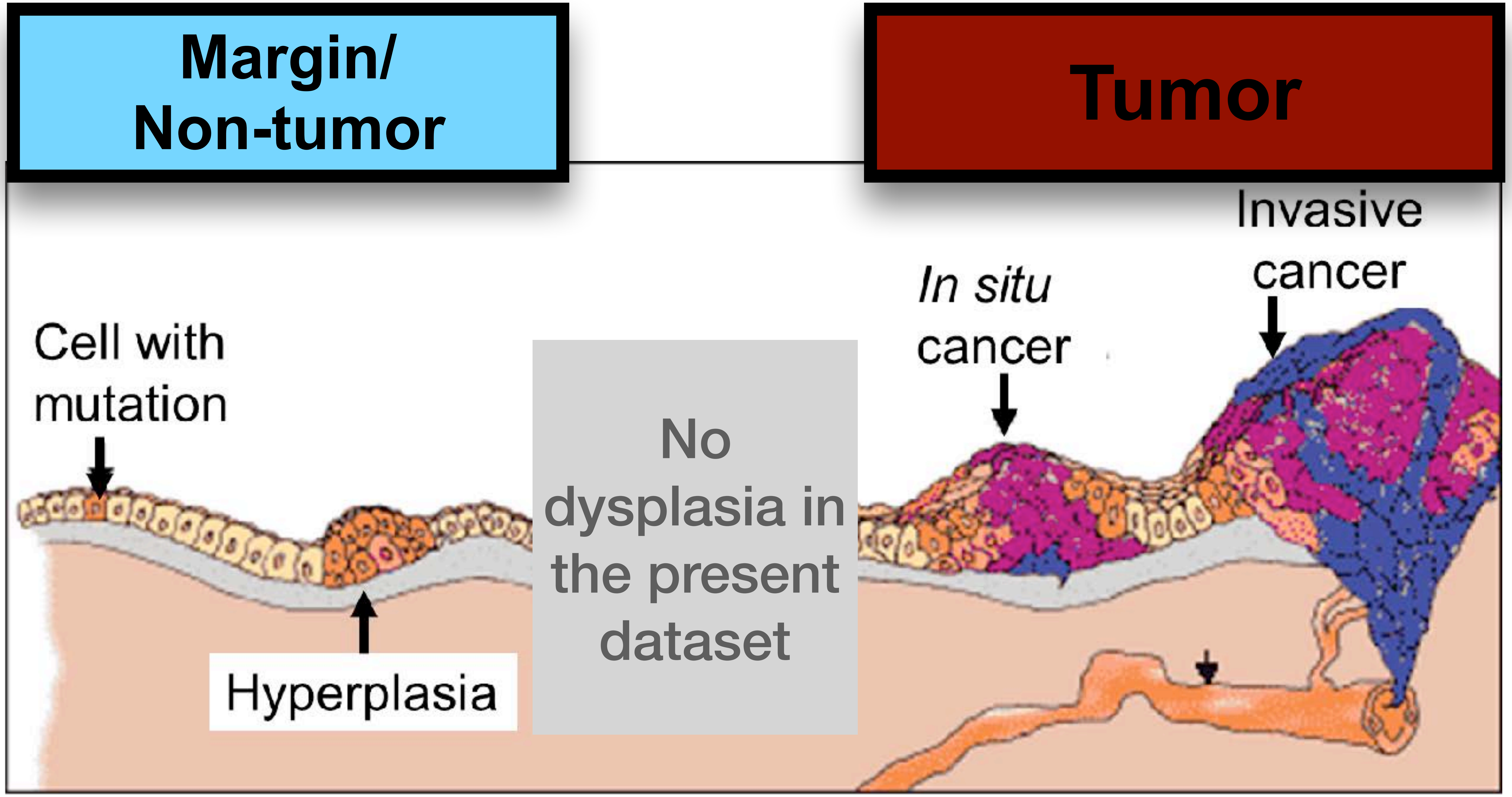


**Goal:** Identify tissues as tumor or margin based on confocal microscopy images (after staining with our fluorescent marker PARPi-FL)

➡ **what are the features that define tumor vs. margin tissues?**

### Margin [1]

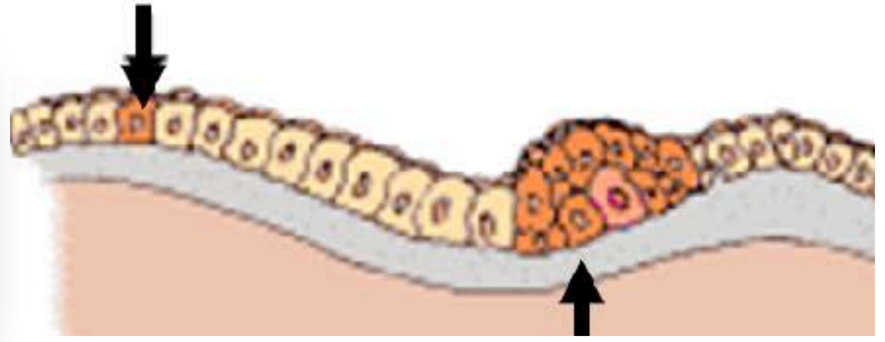

| | | Basal layer | Connective tissue |
|---|---|---|---|
| Morphological stain (H&E) | 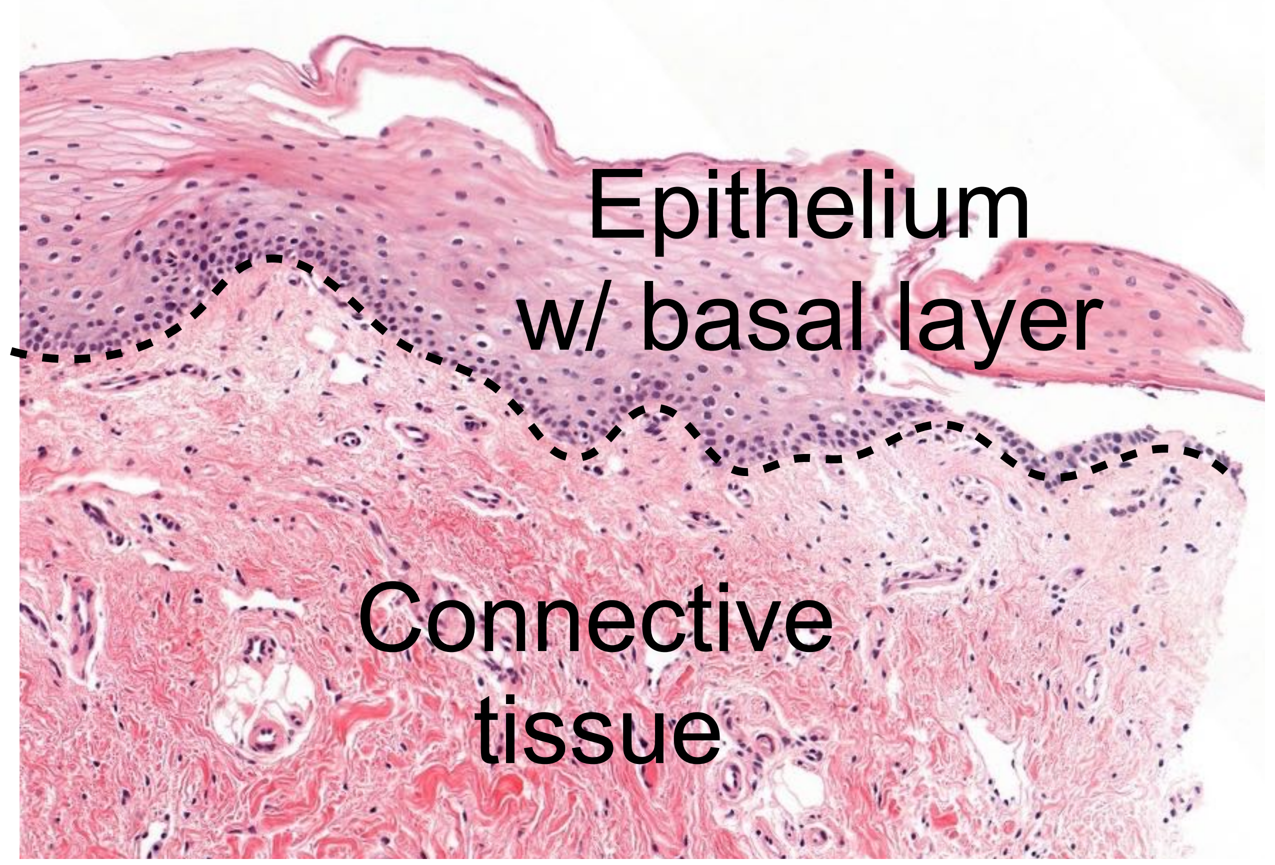  | 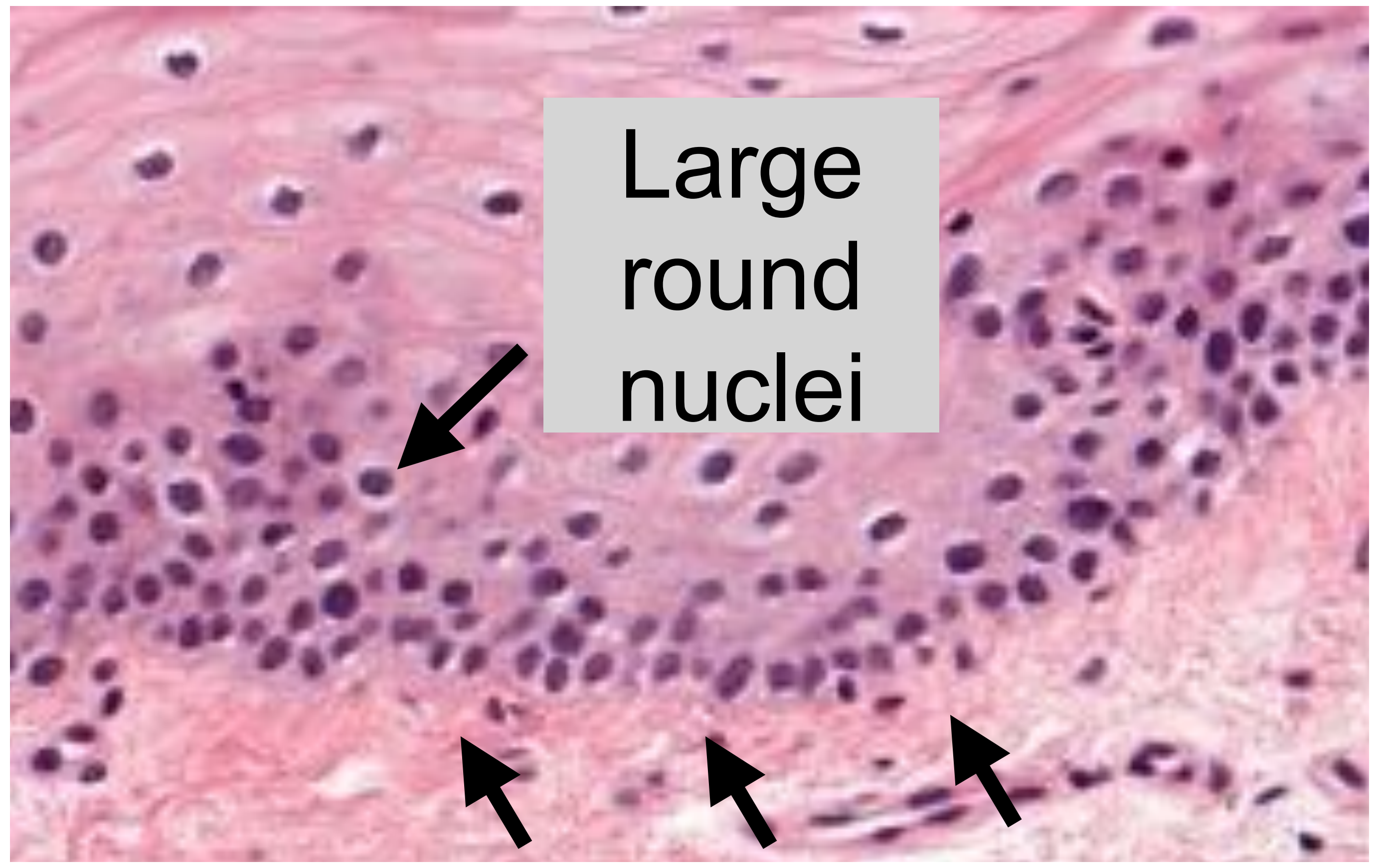  | 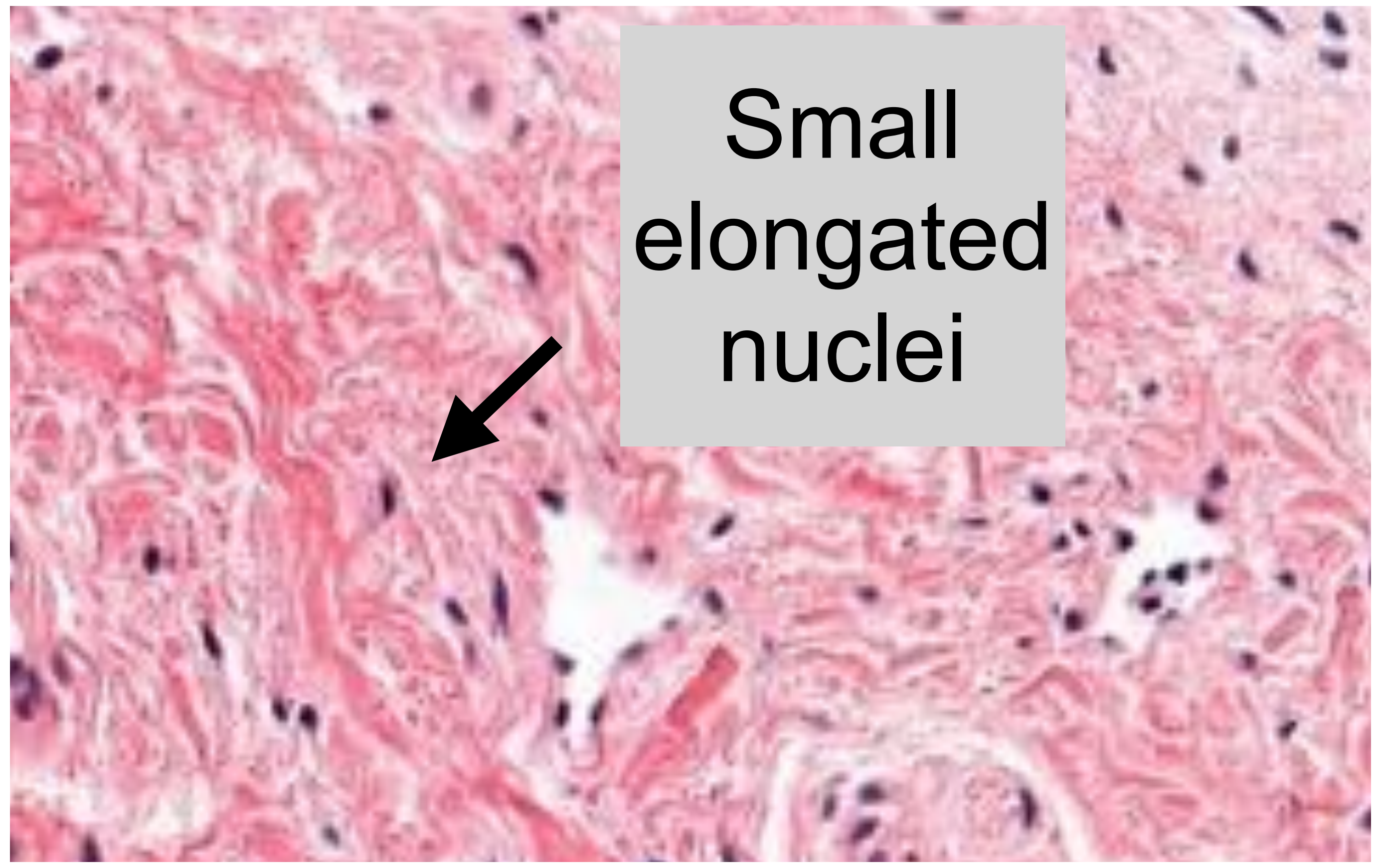  |
| PARP1 stain<br><br>**[PARP1 = brown staining]** | 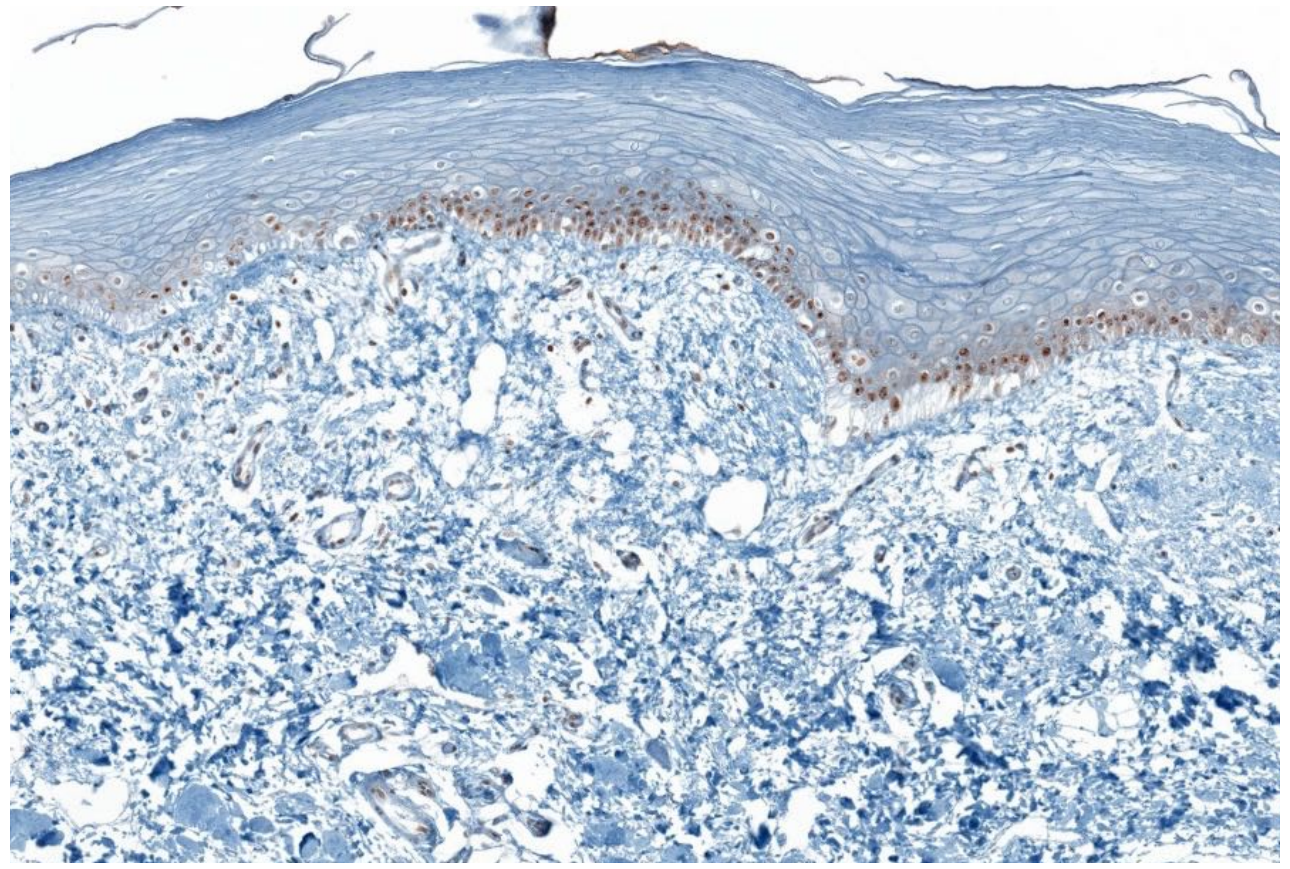 | 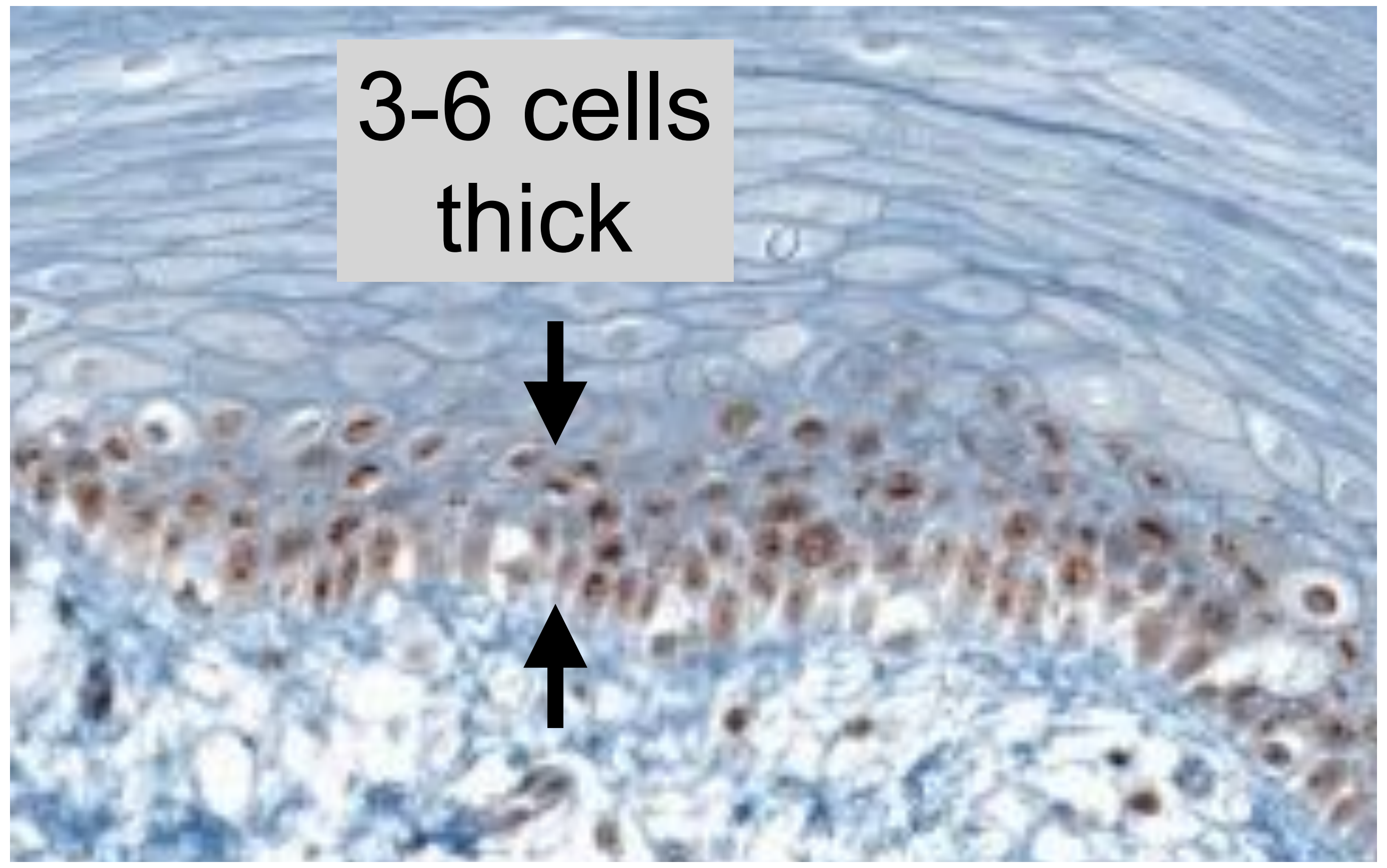  | 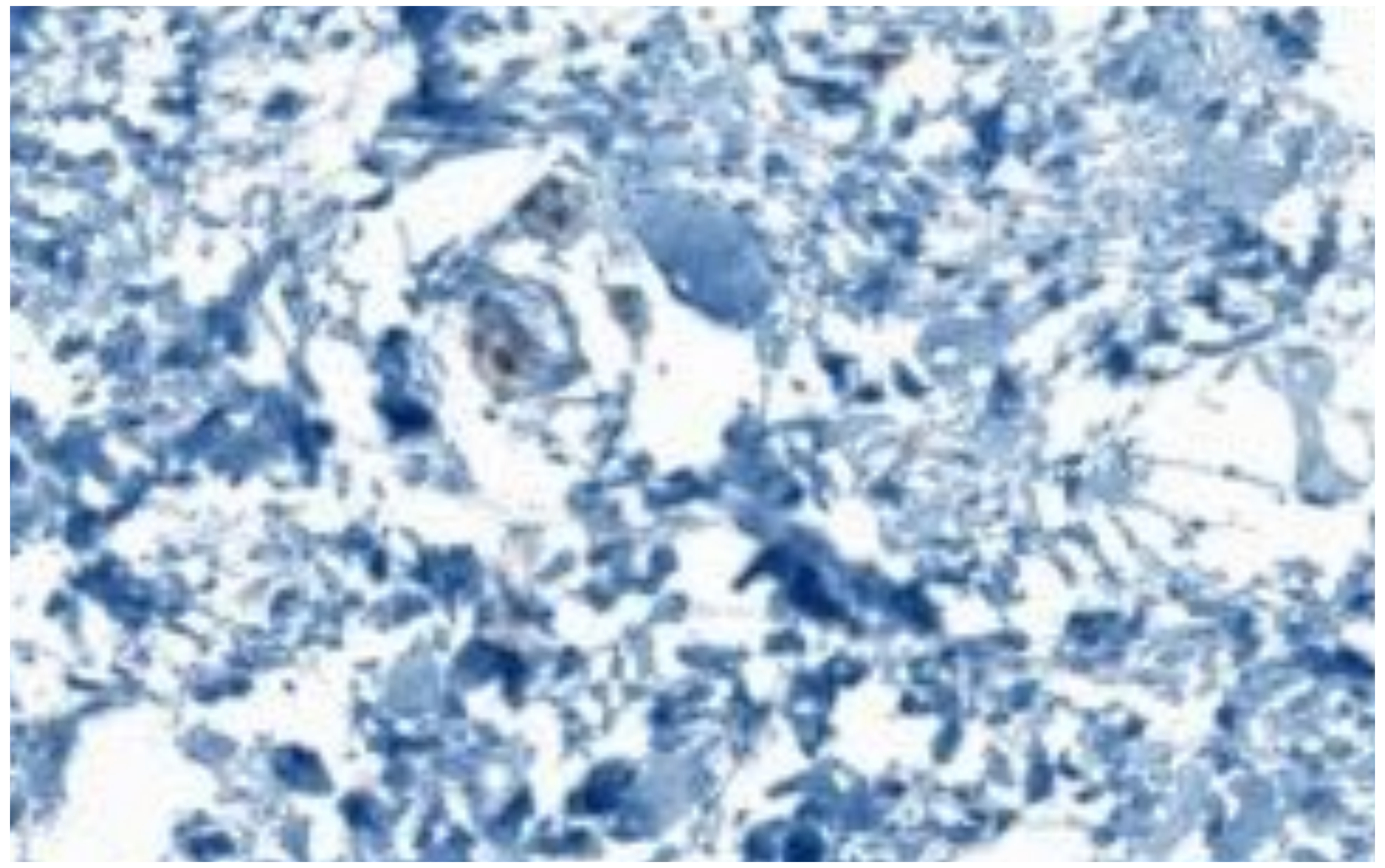 |
| | | • PARP1 in basal layer<br>• PARP1 in large, round nuclei<br>• Thin layer of cells (3-6 cells)<br>• straight or arched line | • Nuclei smaller and elongated compared to epithelium |

### Margin [2]

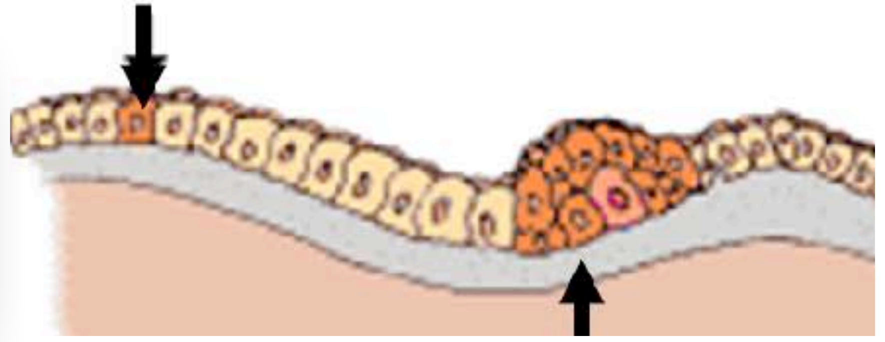

Basal layer

Connective tissue

PARP1 stain

**[PARP1 = brown staining]**

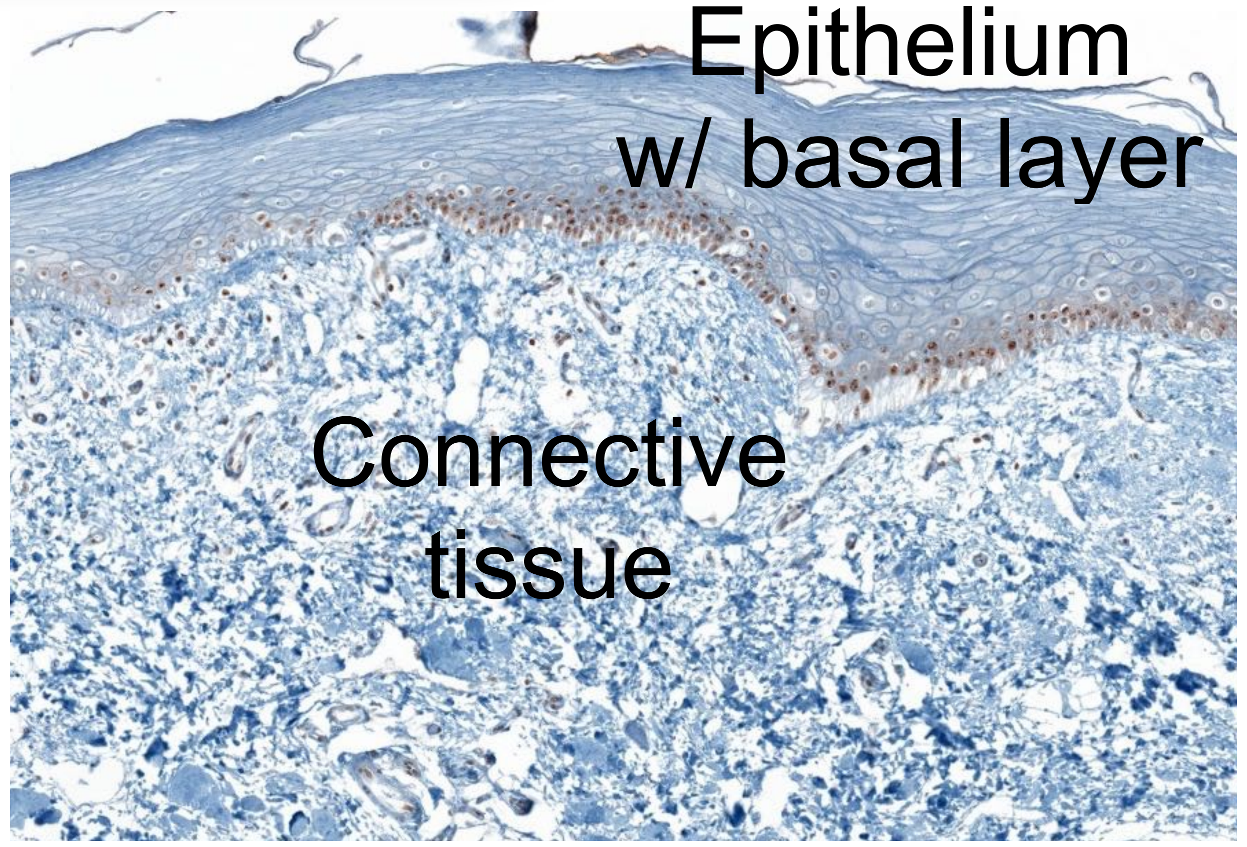


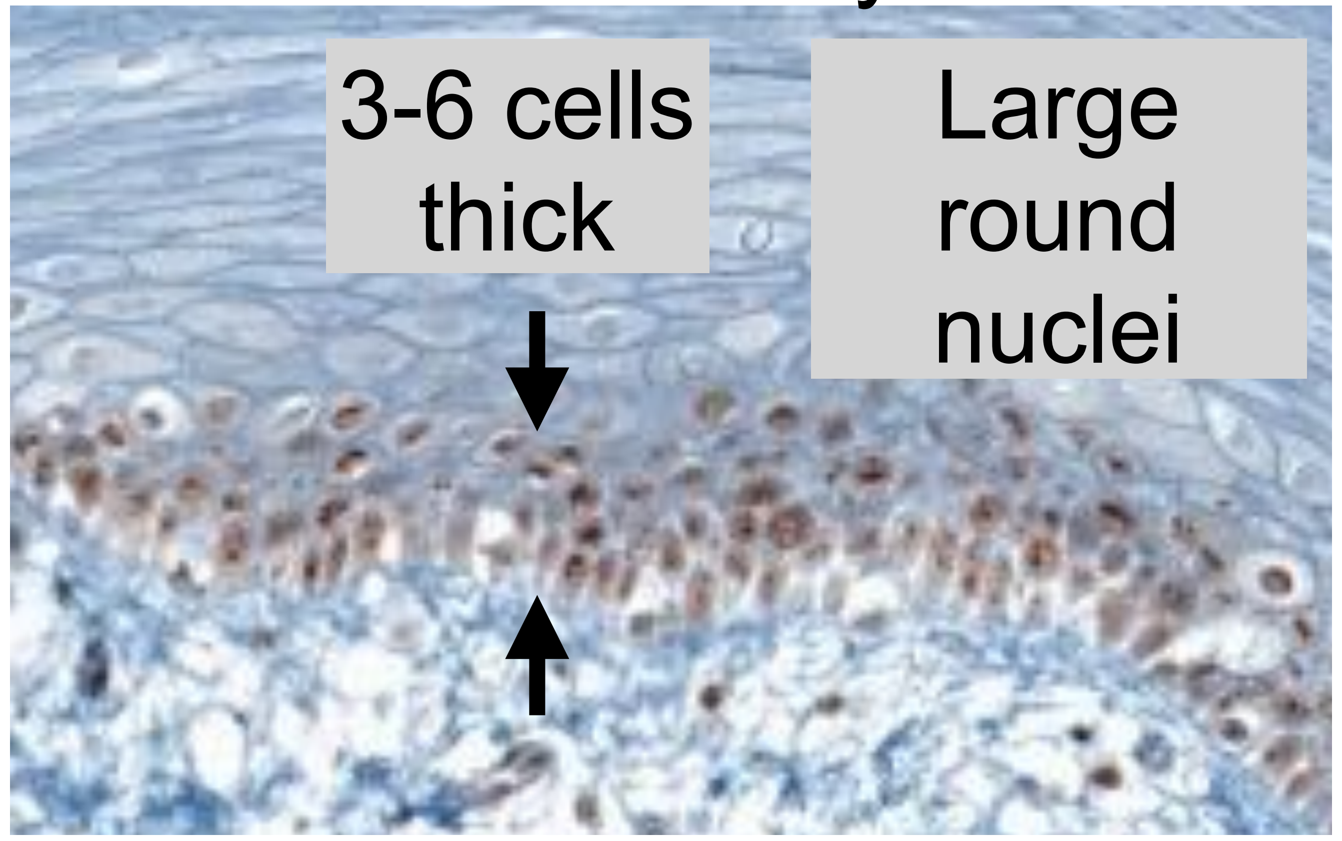


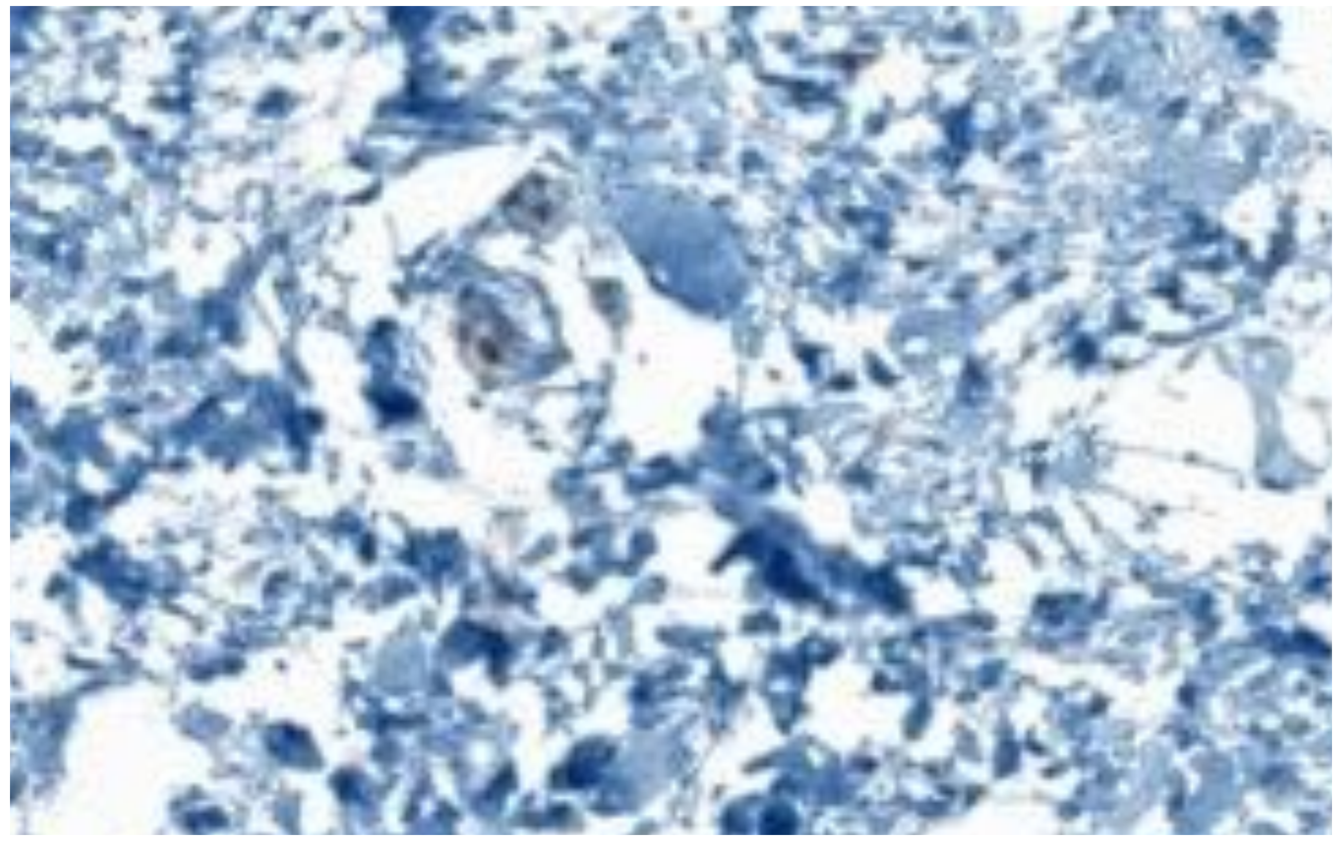

PARPi-FL

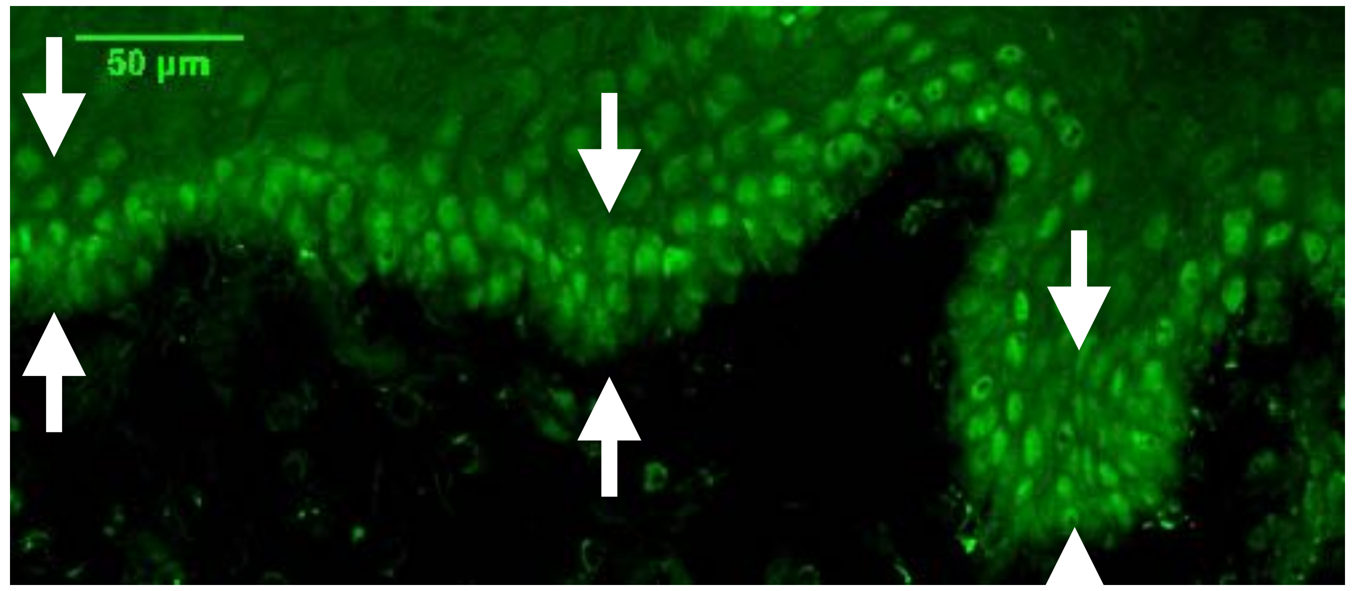


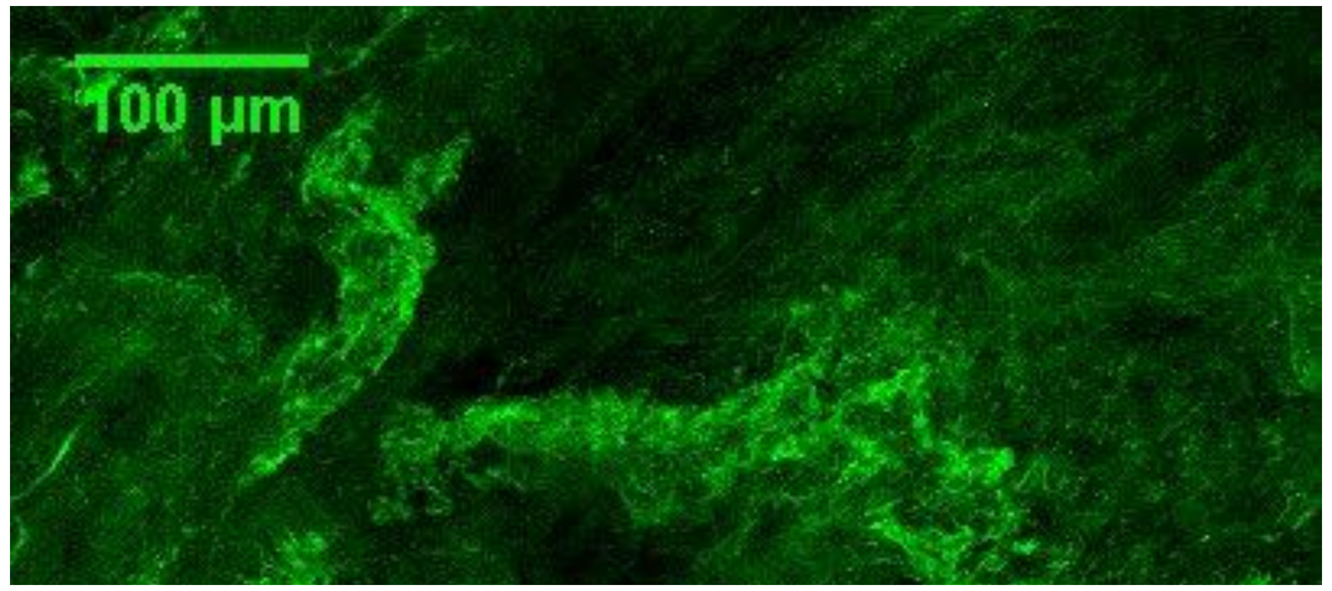


- PARP-FL related fluorescence confined to basal layer
- Fluorescence in large, round cell nuclei
- Straight or arched basal layer

- Connective tissue can have quite high autofluorescence (AF)
- AF is flat, uniform fluorescence with sometime very bright streaks
- No round shapes/cell nuclei!
- Not indicative if tissue is tumor or margin!

### Tumor

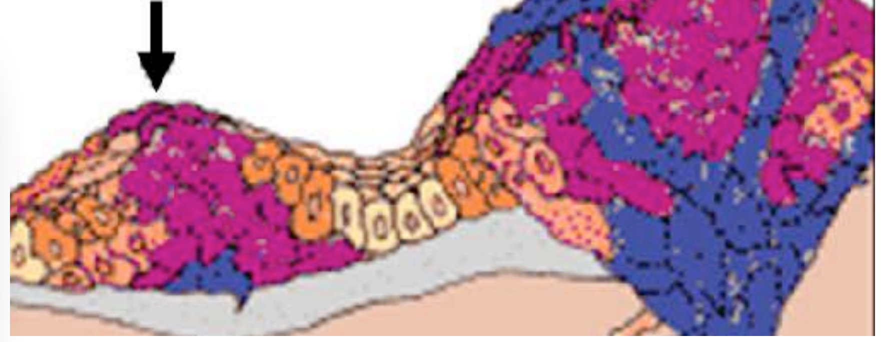

Basal layer | Connective tissue

Morphological stain (H&E)

PARP1 stain

**[PARP1 = brown staining]**

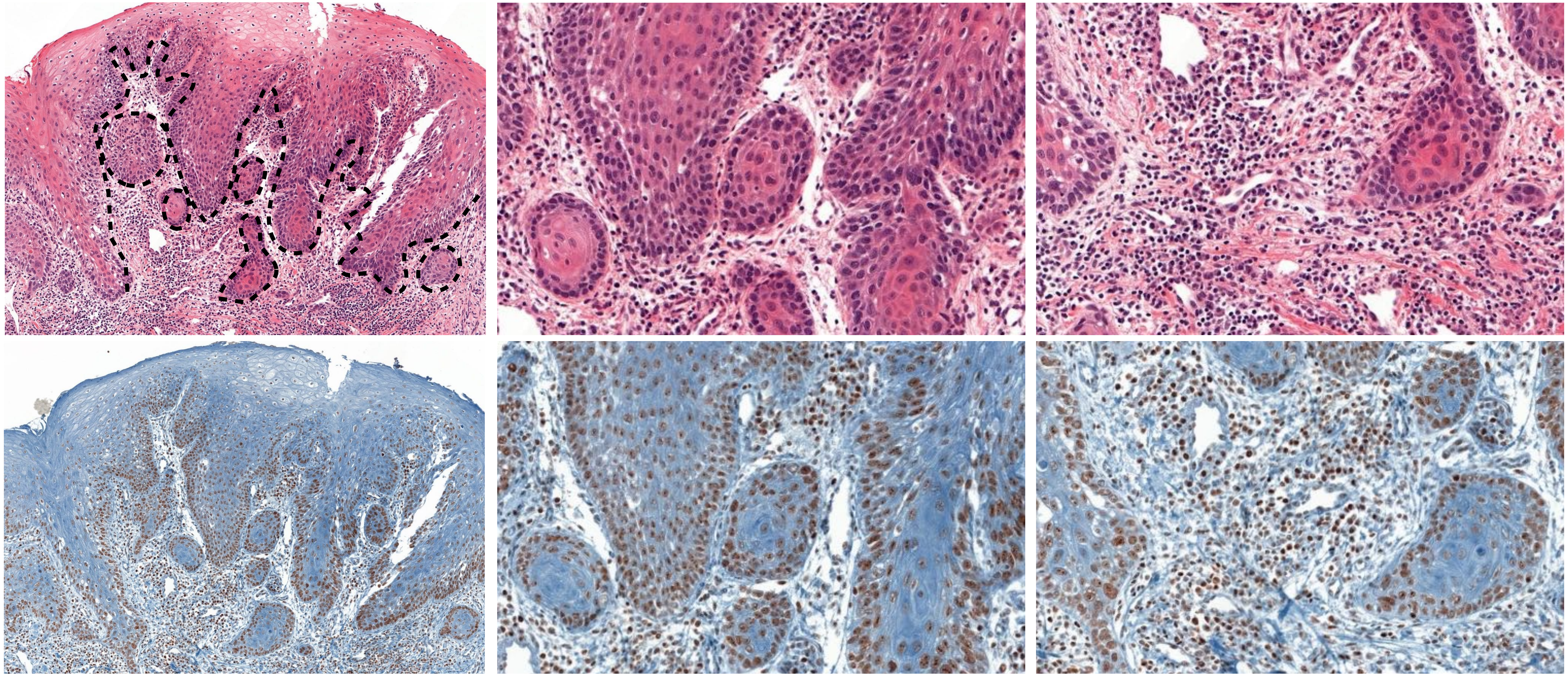

- Basal layer disorganized (extends into deeper layers)
- PARP1 in large, round nuclei outside of a basal layer like structure

- Tumor cells (round nuclei) can extend deep into the connective tissue

### Tumor

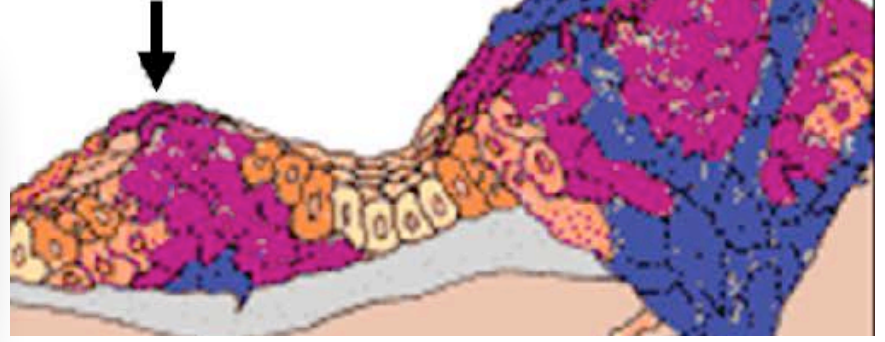

Basal layer | Connective tissue

PARP1 stain

**[PARP1 = brown staining]**

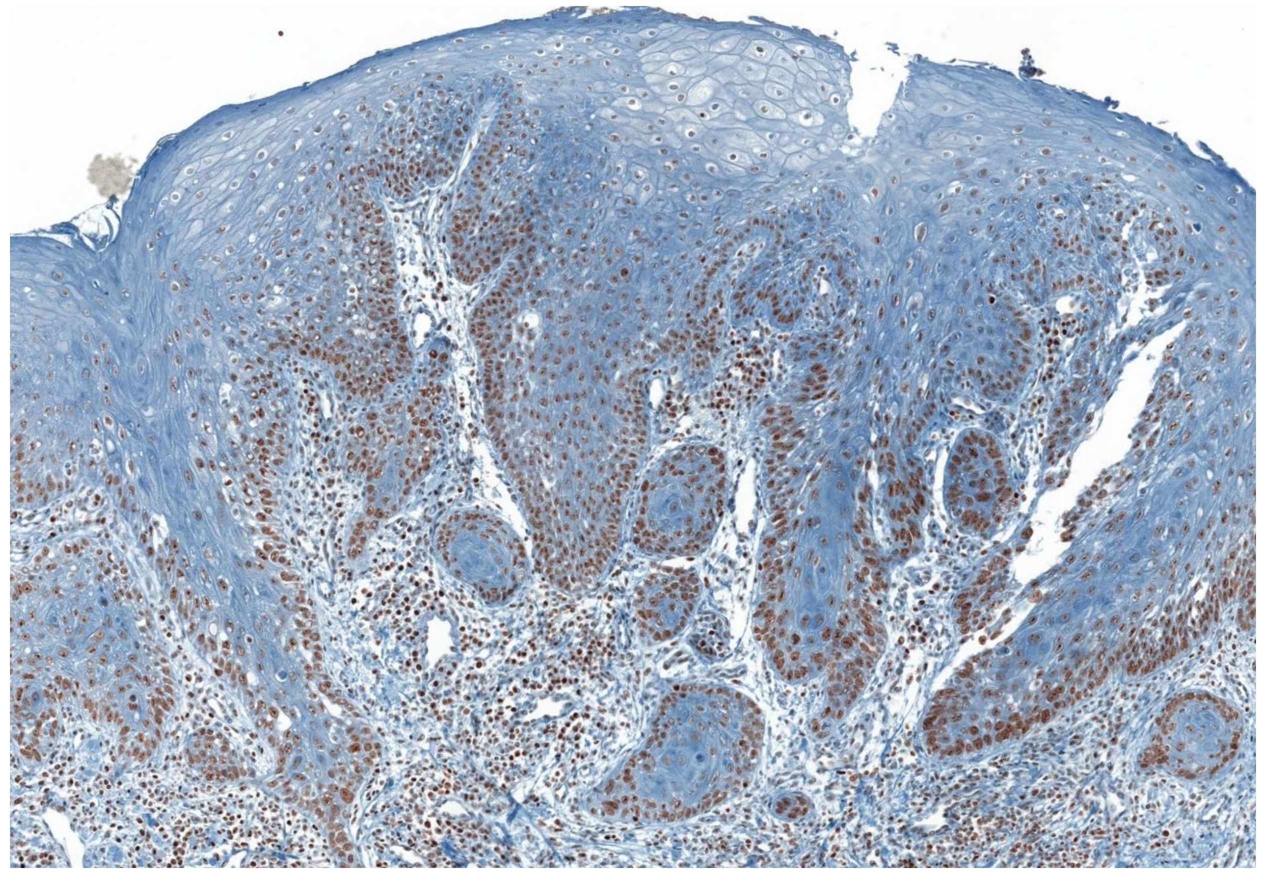

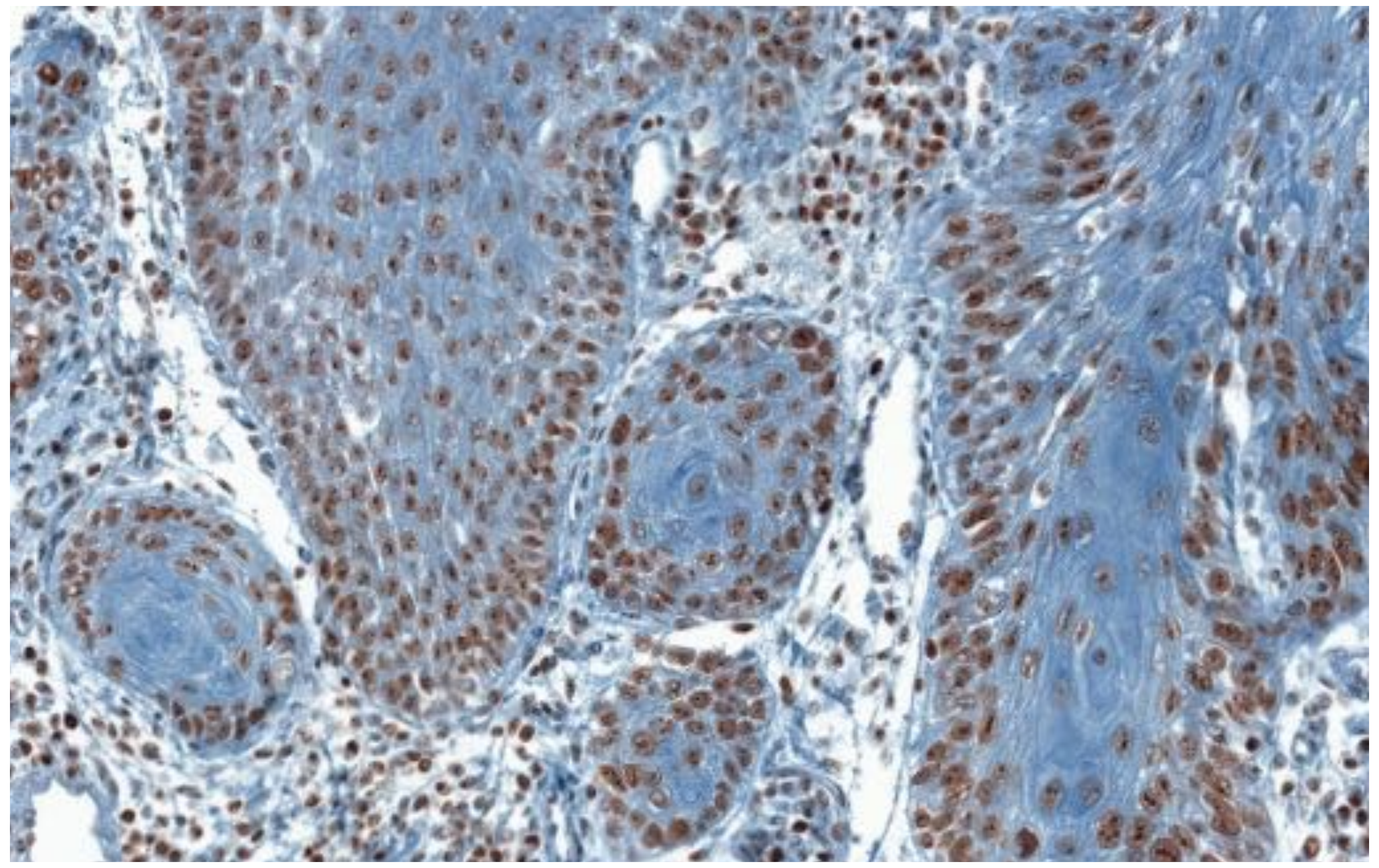

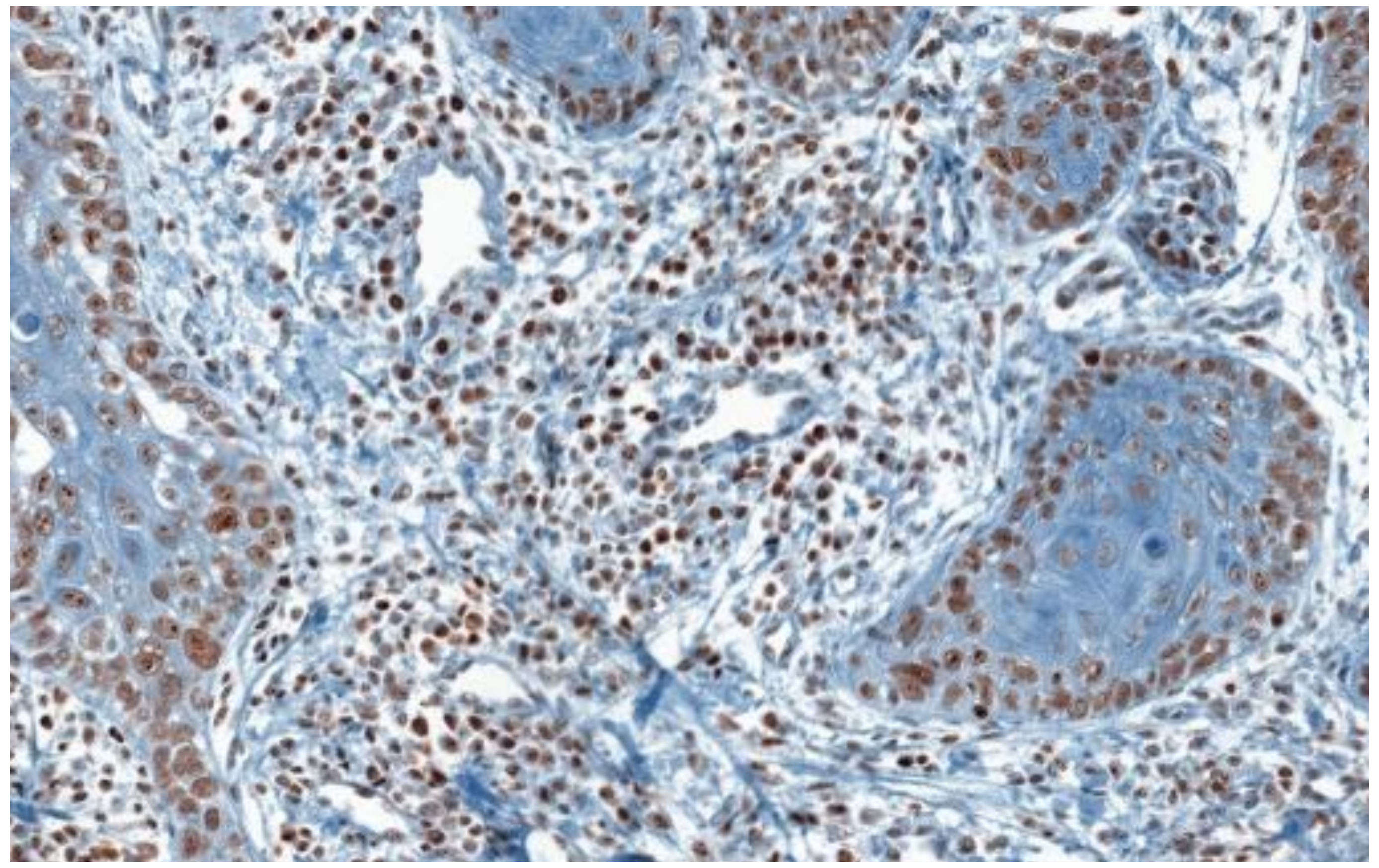

PARPi-FL

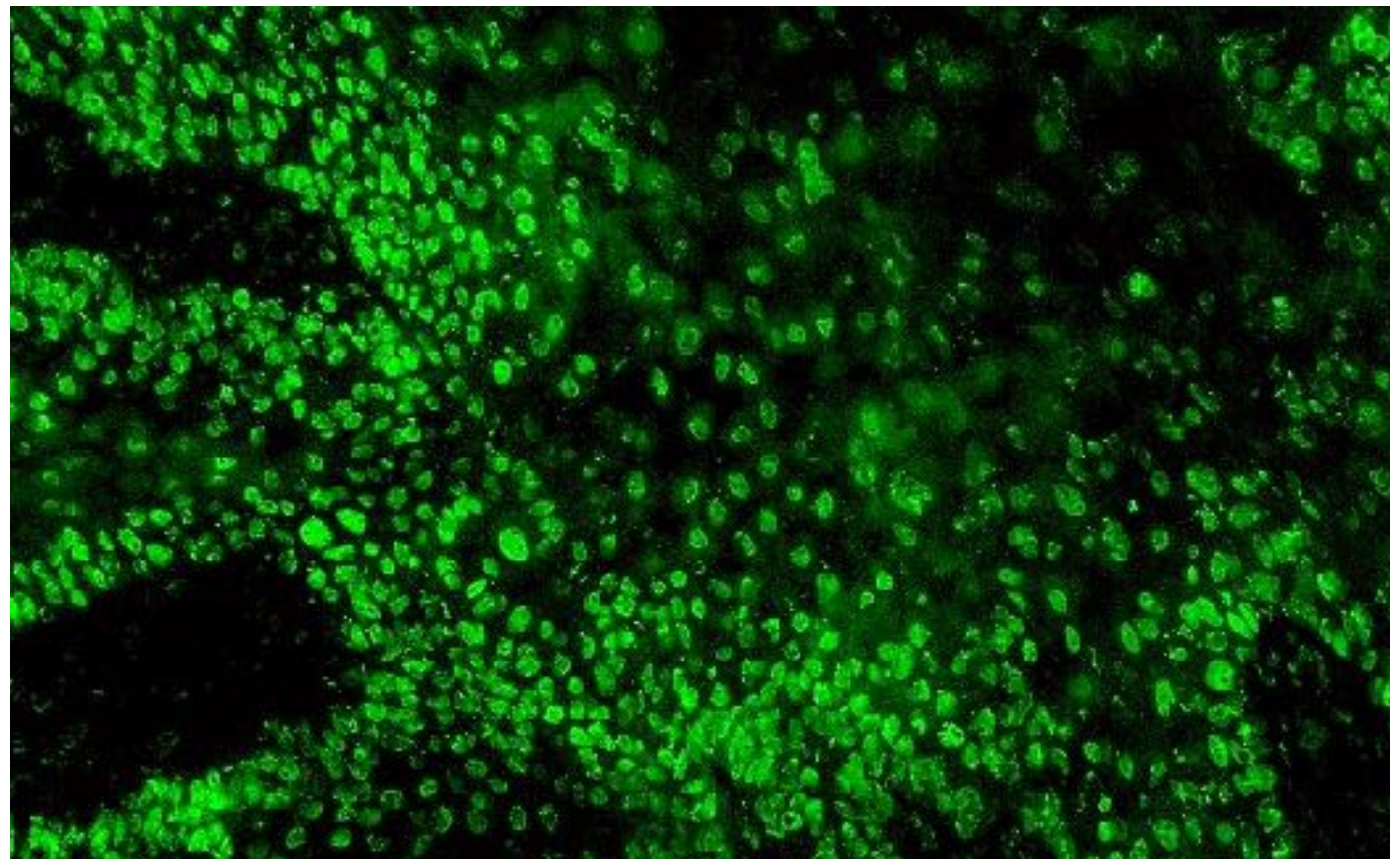

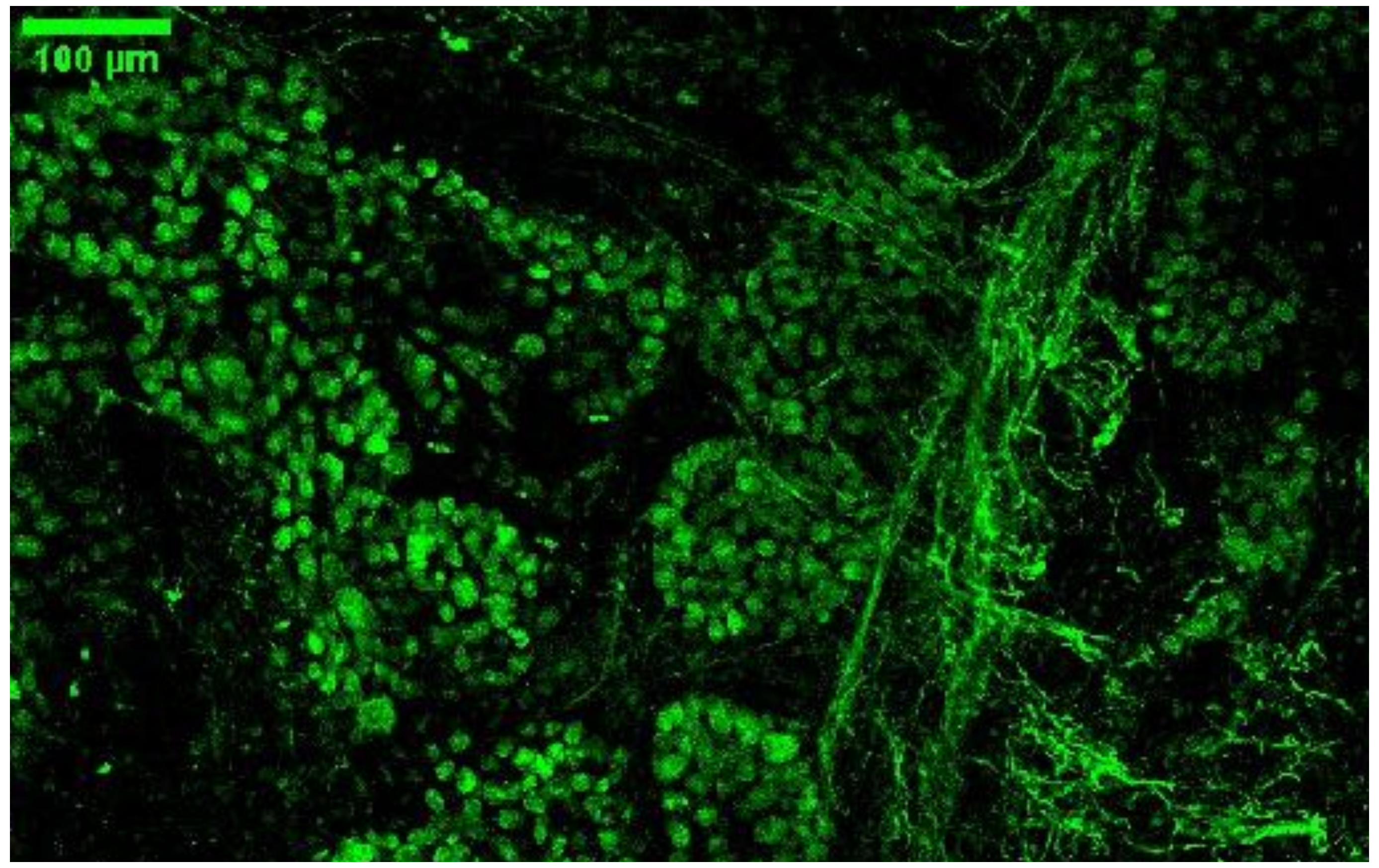


- PARPi-FL in round cells in more or less organized fashion
- Extension of green fluorescent cells beyond a thin basal layer indicates tumor

- Tumor cells (round nuclei) can extend deep into the connective tissue
- Structures with strong AF present (areal, flat fluorescence or bright streaks)

### Autofluorescence

**Green only:**
PARPi-FL: green
AF: green

**Overlay:**
PARPi-FL: green
AF: red/yellow

**Most important characteristics**

**Tumor**

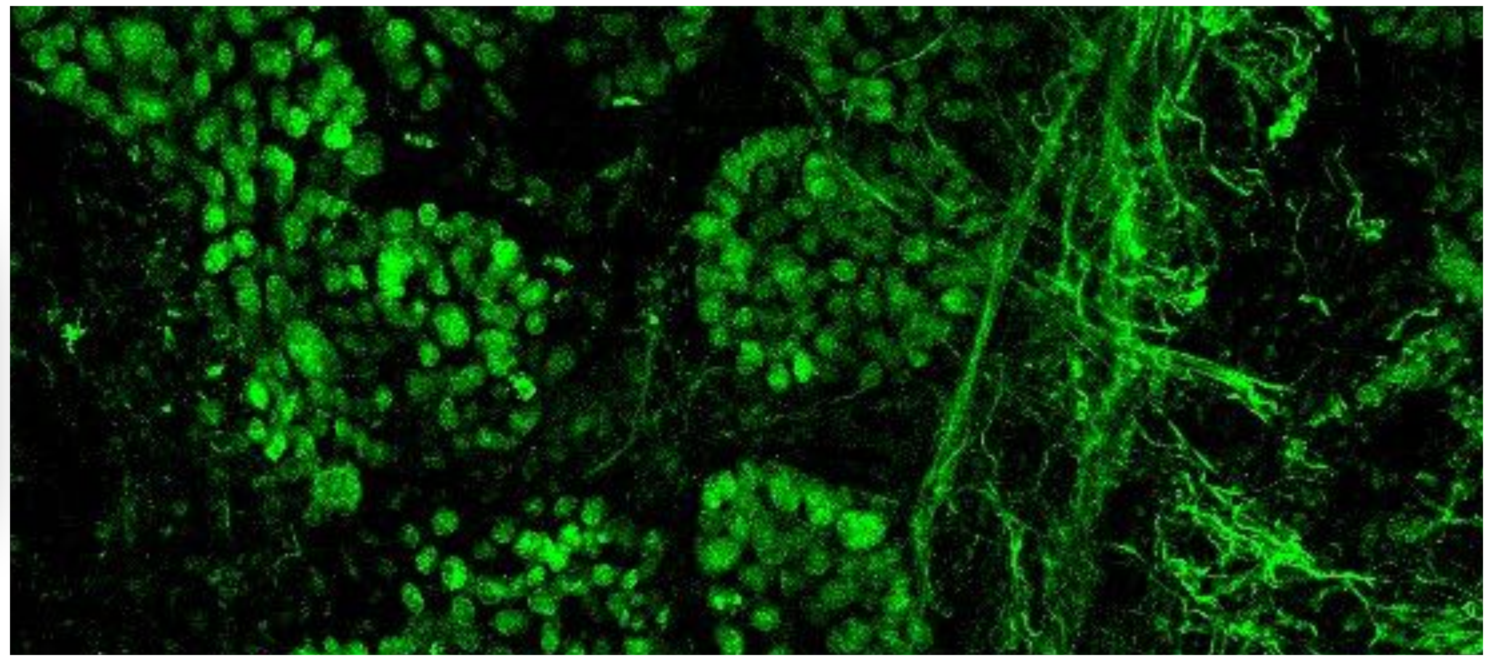

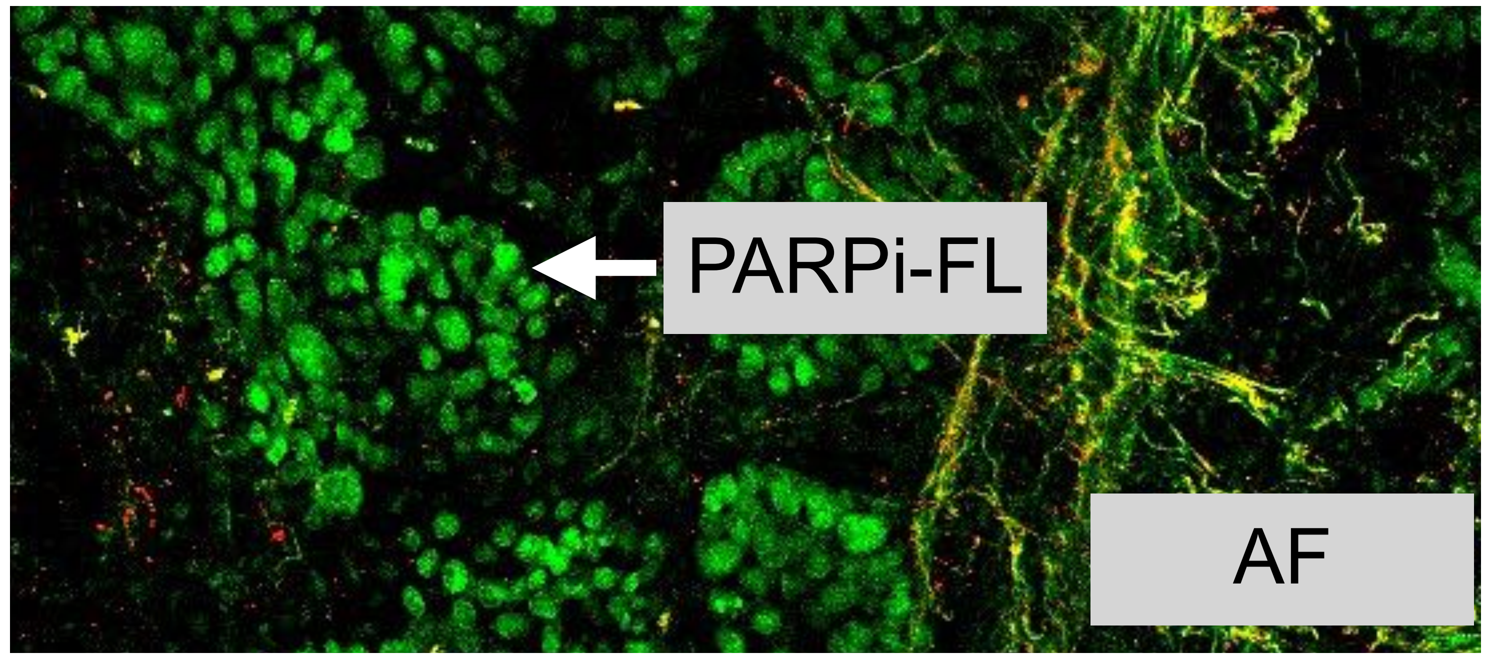


- PARPi-FL stained large, round cells beyond basal layer (**MUST** be present)
- AF **CAN** be present

**Margin**

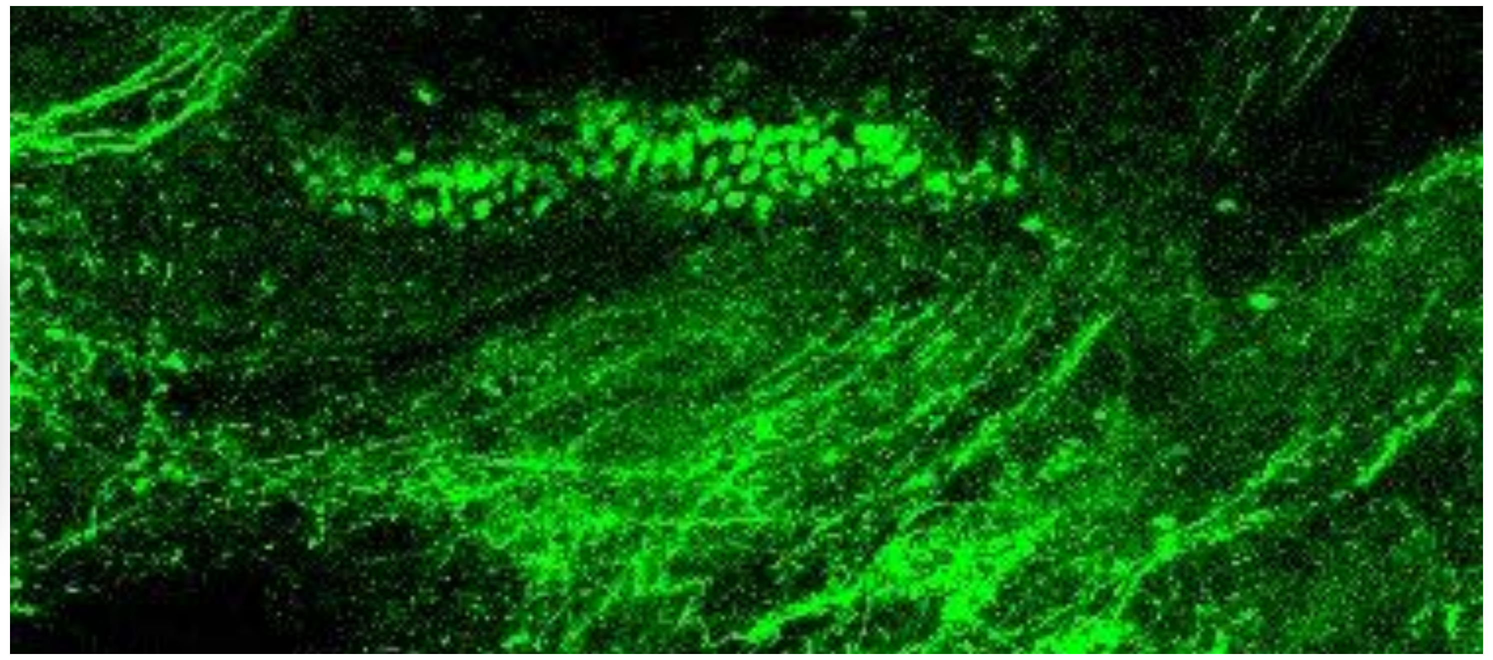

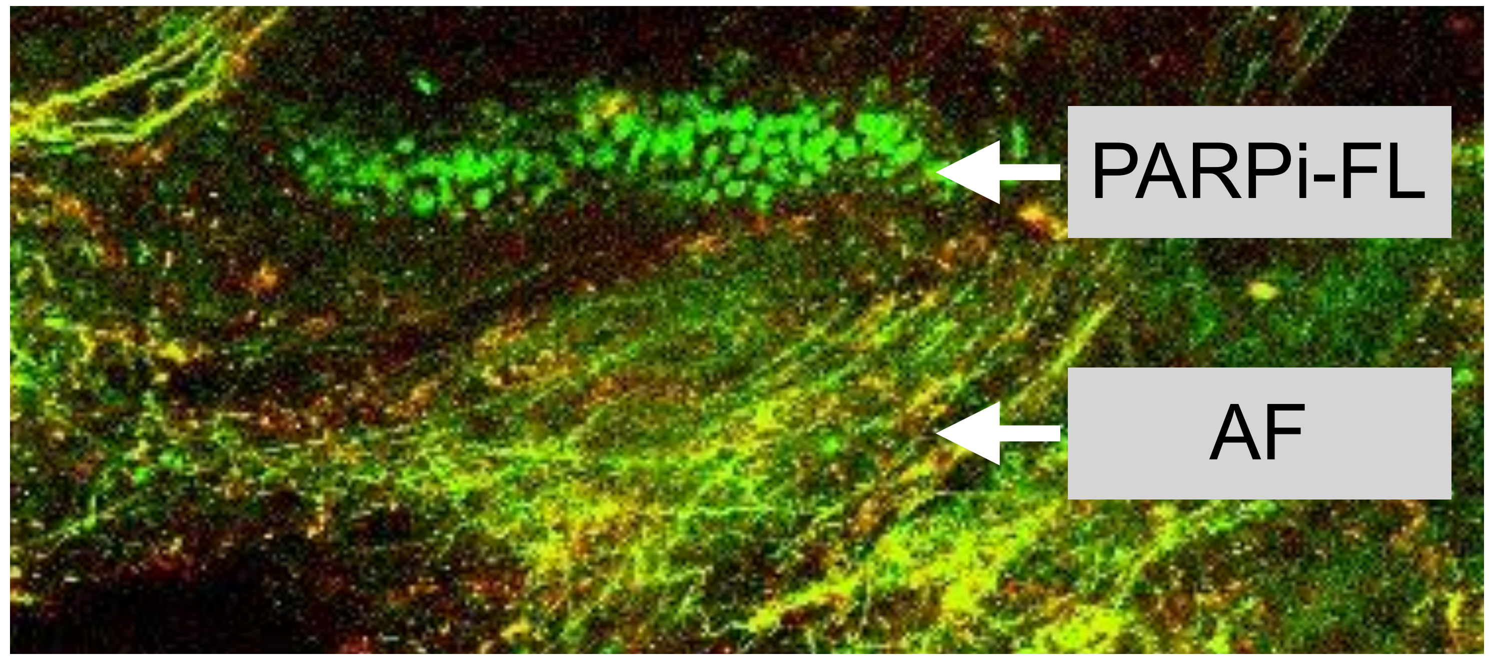


- Thin, regular basal layer with PARPi-FL stained cells (<6 cell layers) **CAN** be present
- AF **CAN** be present

➡ Round cells: PARPi-FL

➡ flat/area/streak stained: Autofluorescence

➡ Green round cells: PARPi-FL

➡ red/yellow signals: Autofluorescence (not diagnostic/ does not help to decide if a tissue is tumor or margin

### Tumor

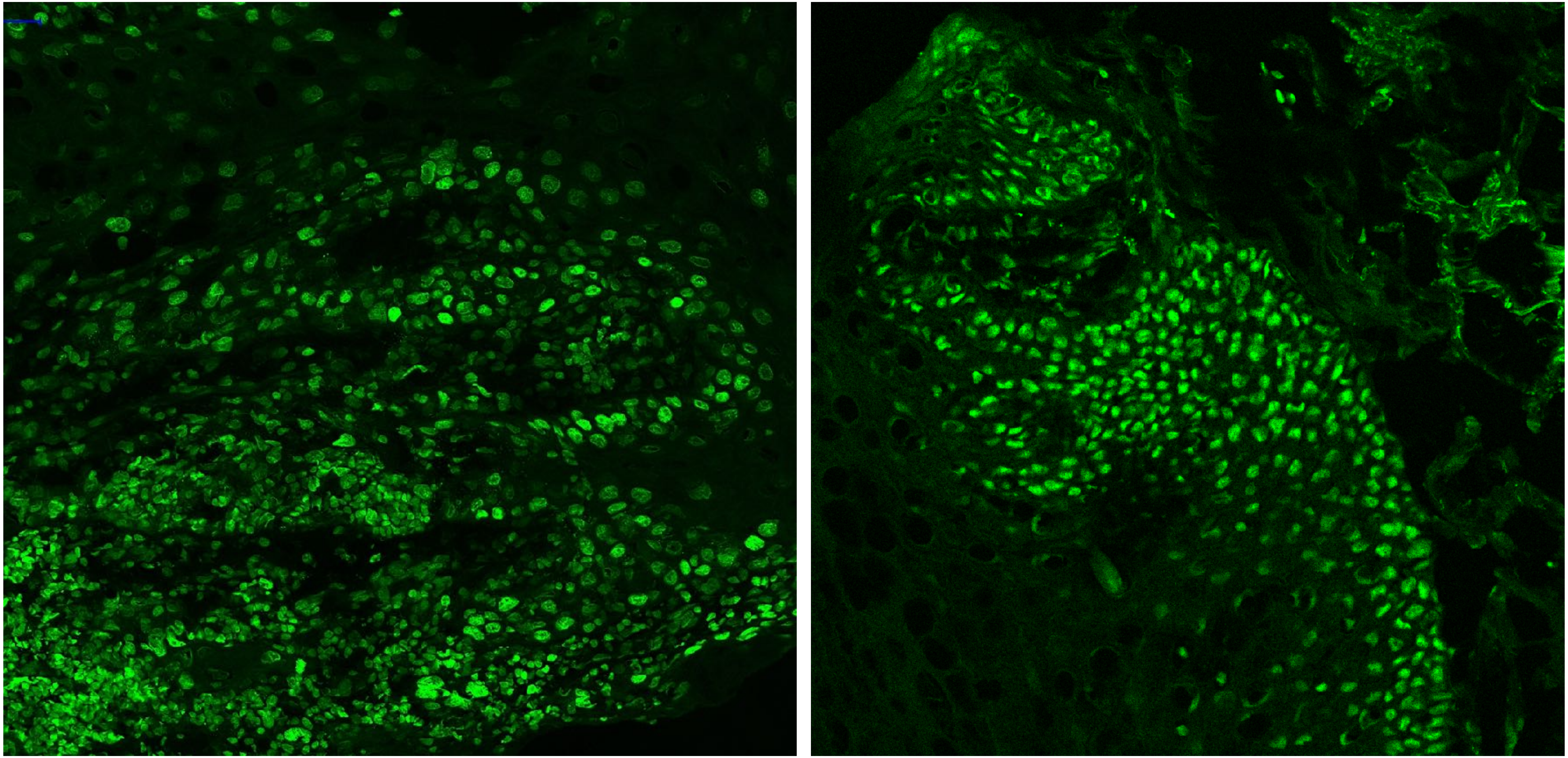

PARPi-FL only (green)

### Margin

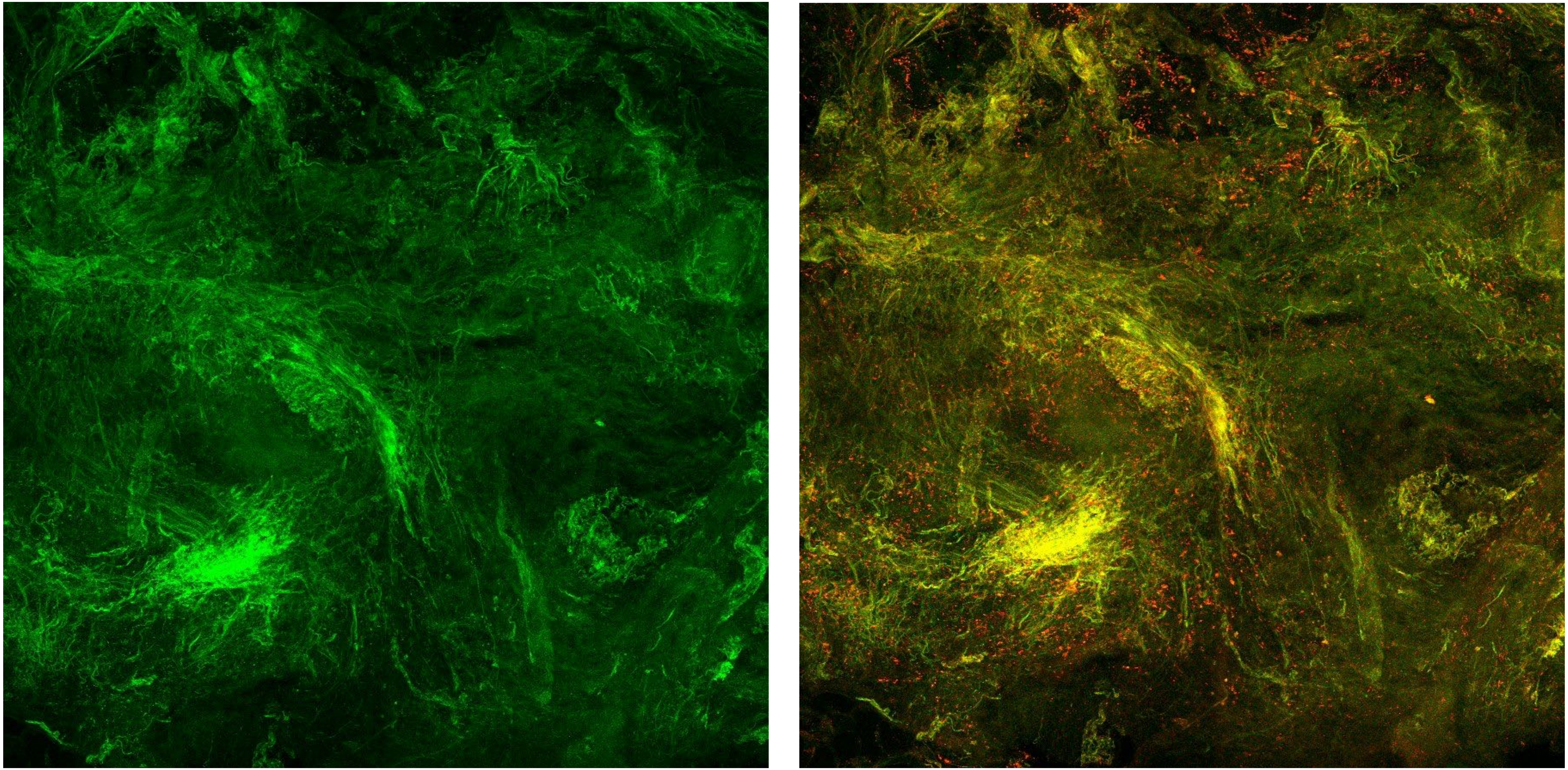

| Margin | Tumor |
|---|---|

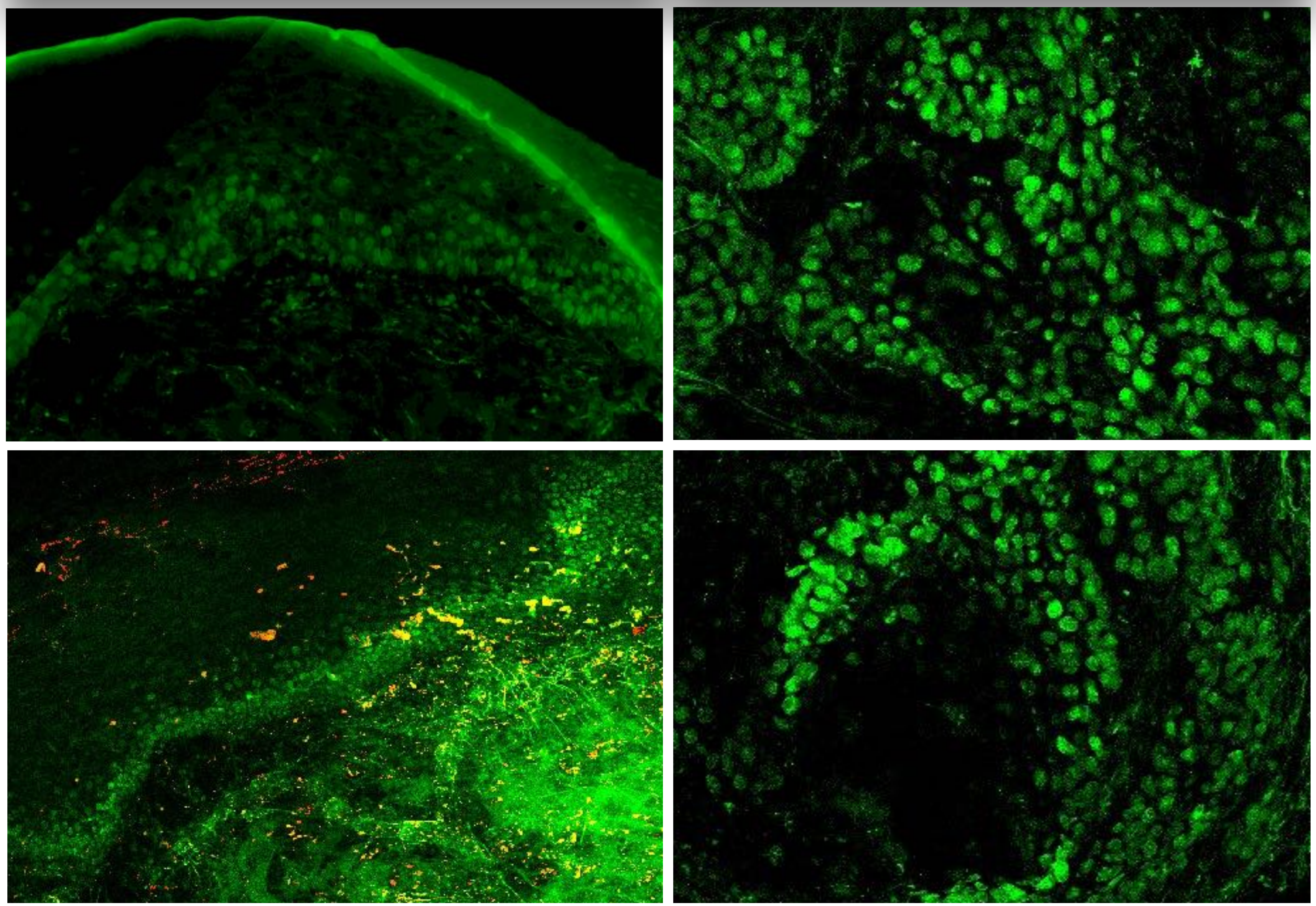

**Margin**

- PARP-FL related fluorescence confined to thin, regular basal layer (<6 cell layers)
- PARPi-FL signal in large, round cell nuclei **CAN** be present
- Autofluorescence **CAN** be present
- basal layer not always visible

**Tumor**

- PARPi-FL stained large, round cells in structures other than thin, regular basal layer **MUST** be present
- PARPi-FL signal in large, round cell nuclei
- AF **CAN** be present (check on overlay image)

#### *How to decide*

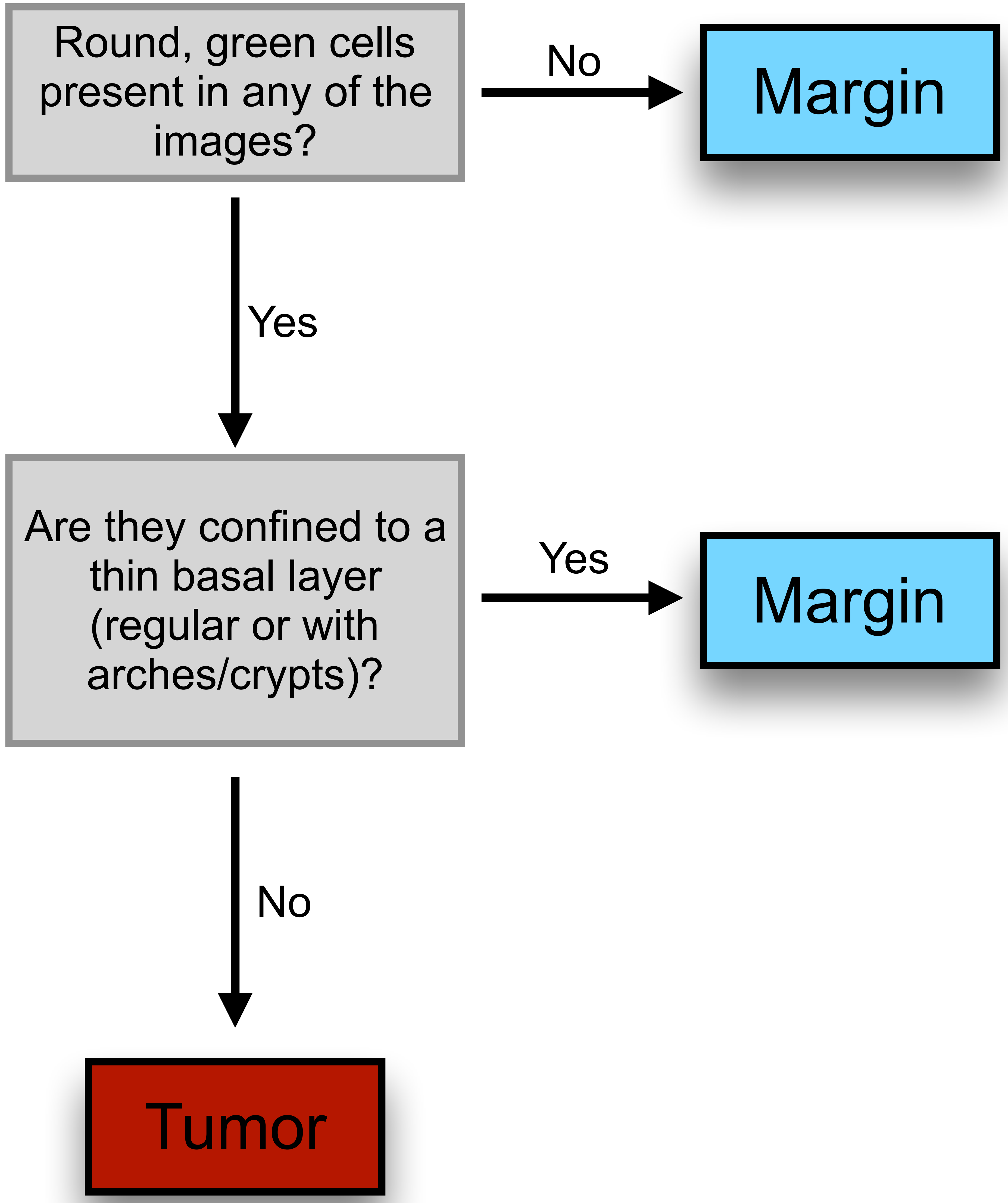
