## Supplementary material for "PARP1 as a biomarker for early detection and intraoperative tumor delineation in epithelial cancers – first-in-human results": Blinded Study_Data Set

### PARPi-FL staining of fresh biopsy tissues for identification of tumor and margin tissue

Blinded study

Data set

1472

### Overview

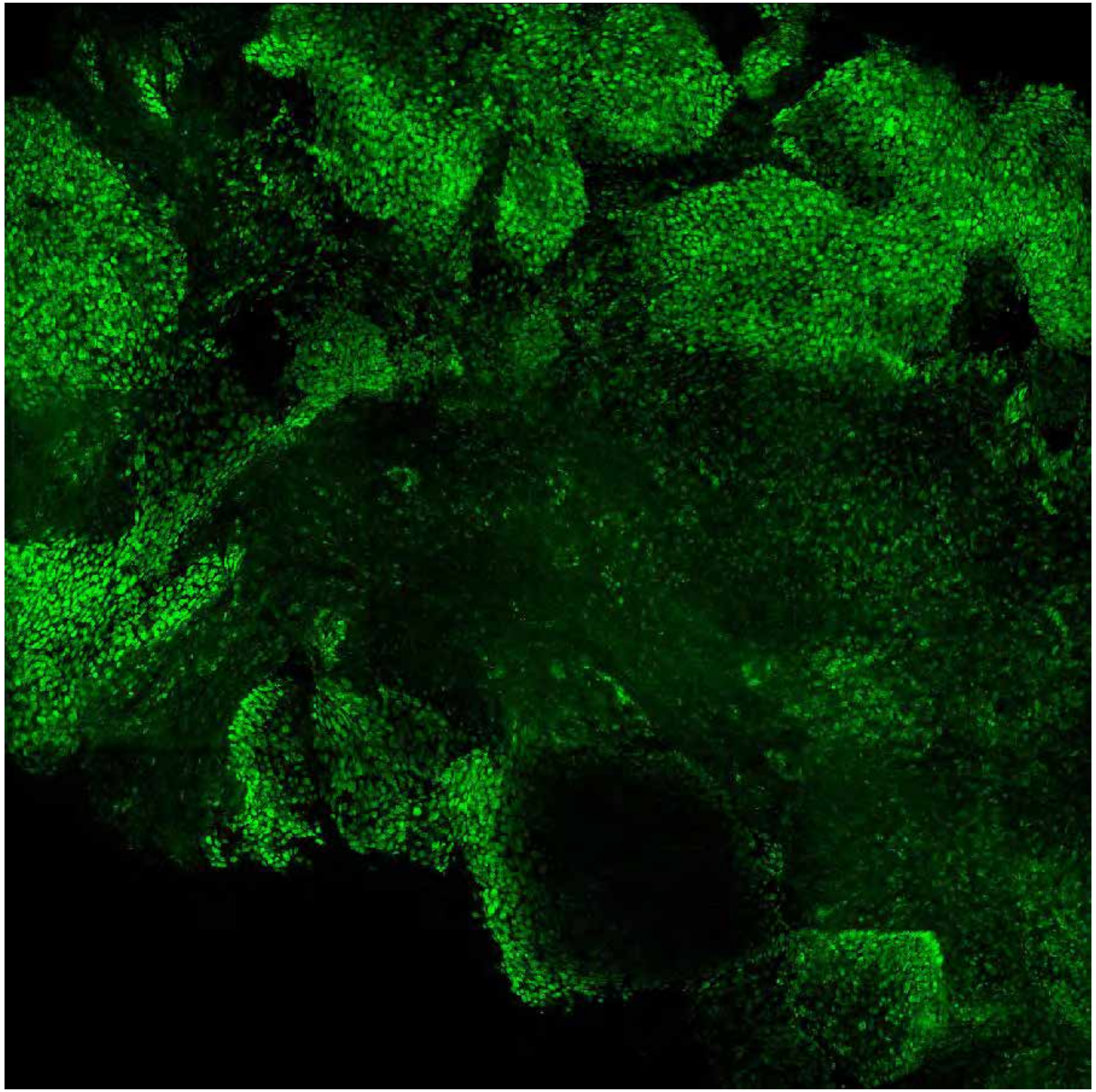

PARPi-FL only (green)

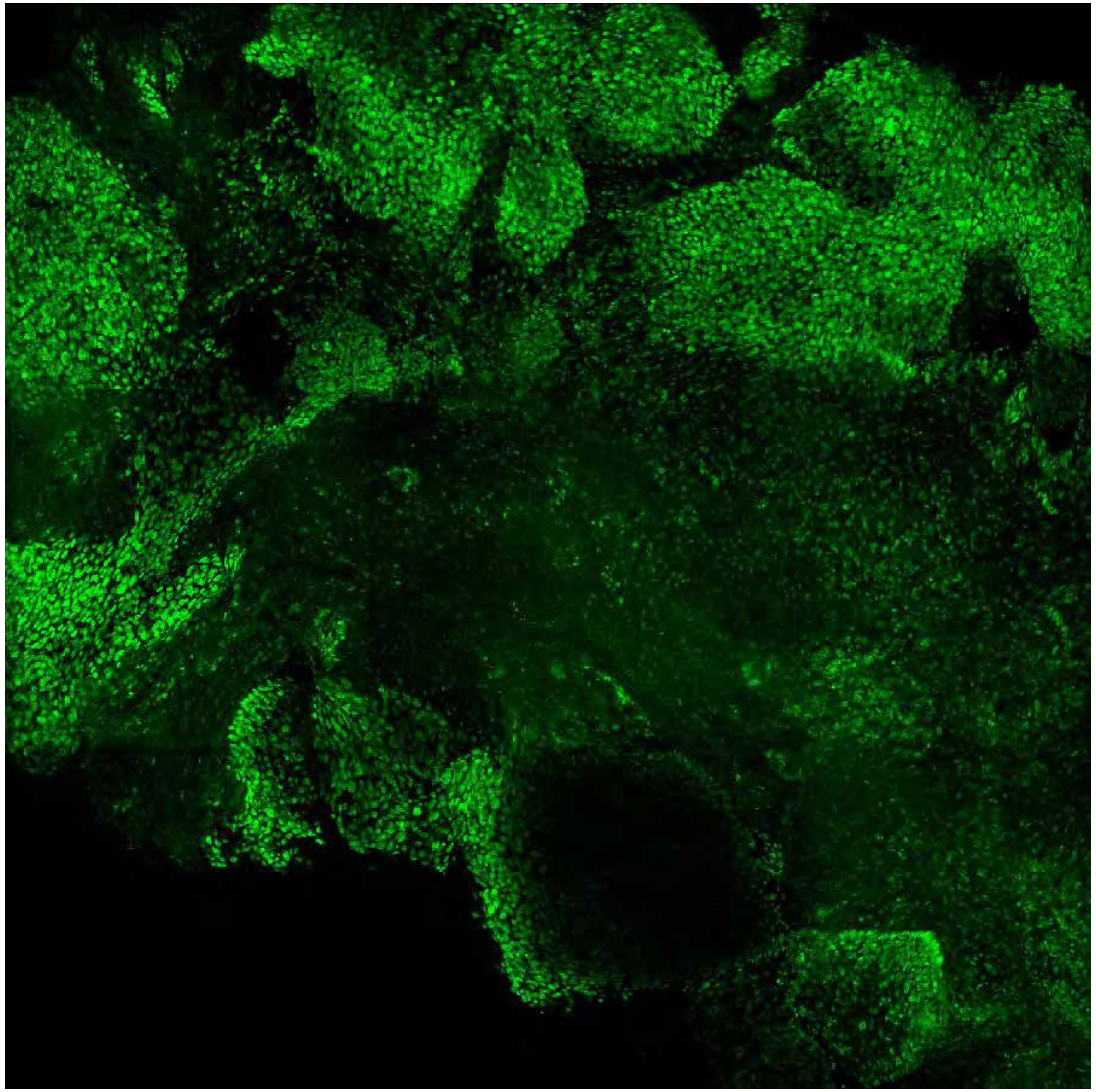

PARPi-FL (green) + Autofluorescence (red/yellow)

### Close-up

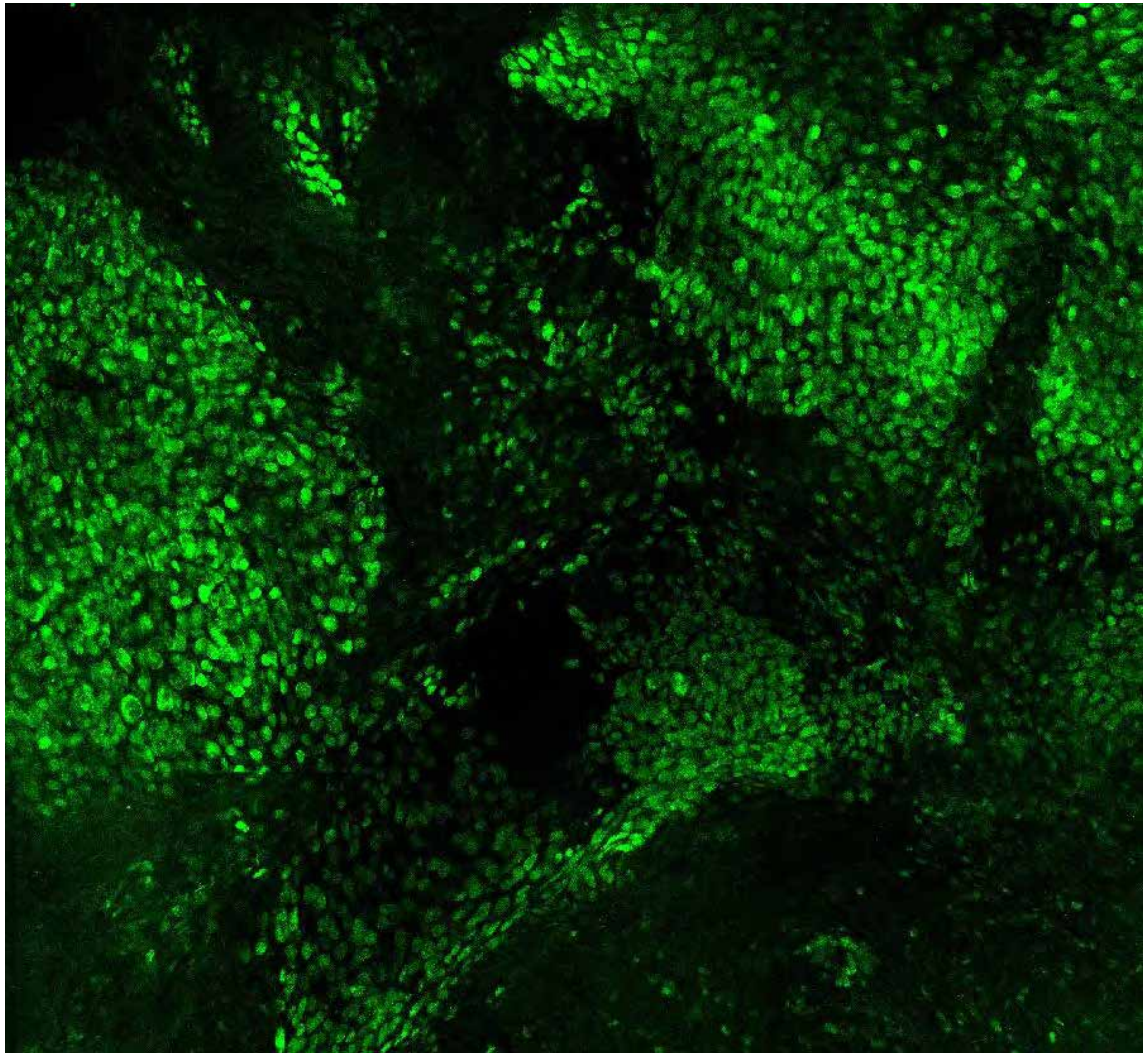

**PARPi-FL only (green)**

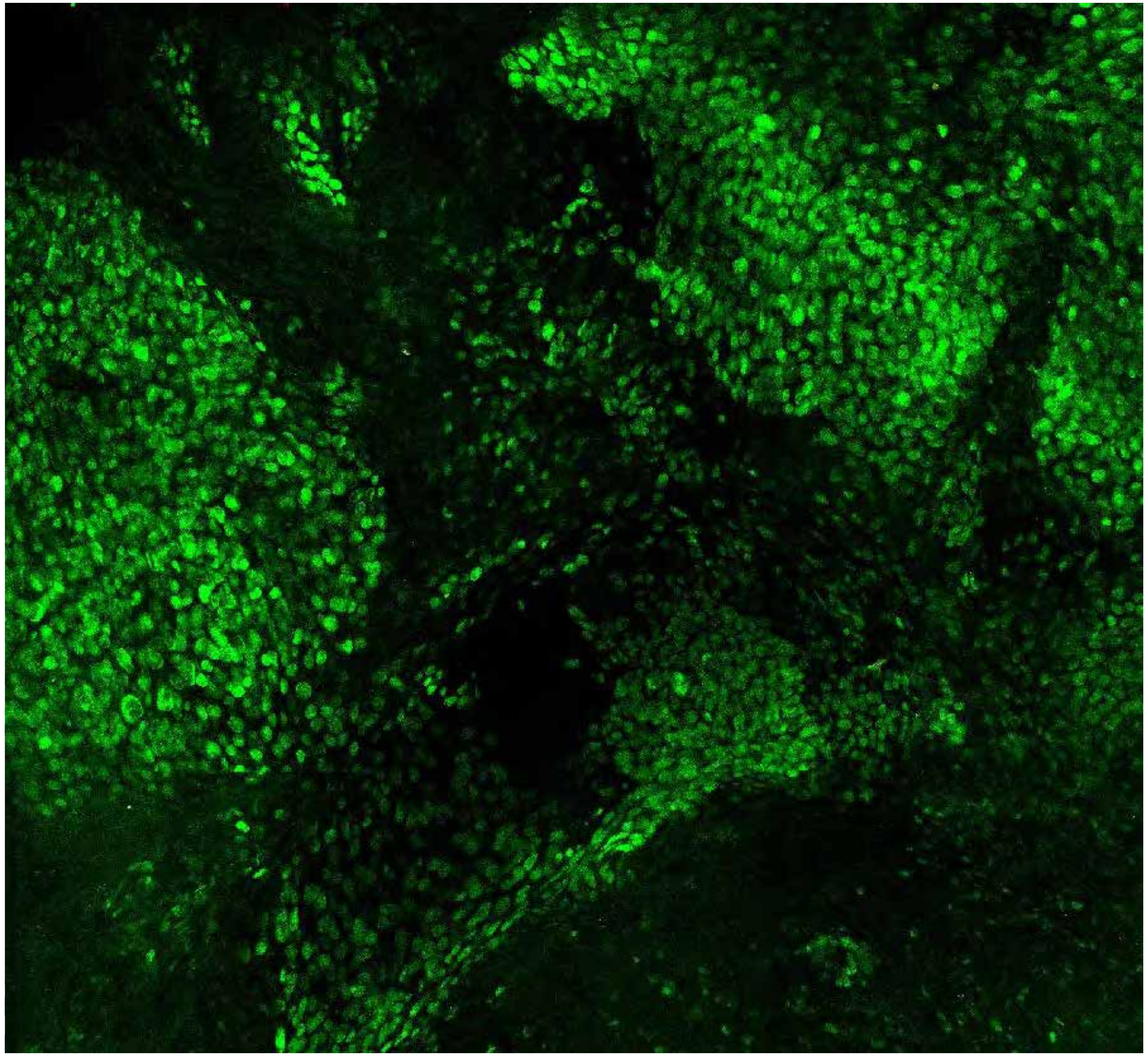

**PARPi-FL (green) + Autofluorescence (red/yellow)**

1472

### Close-up

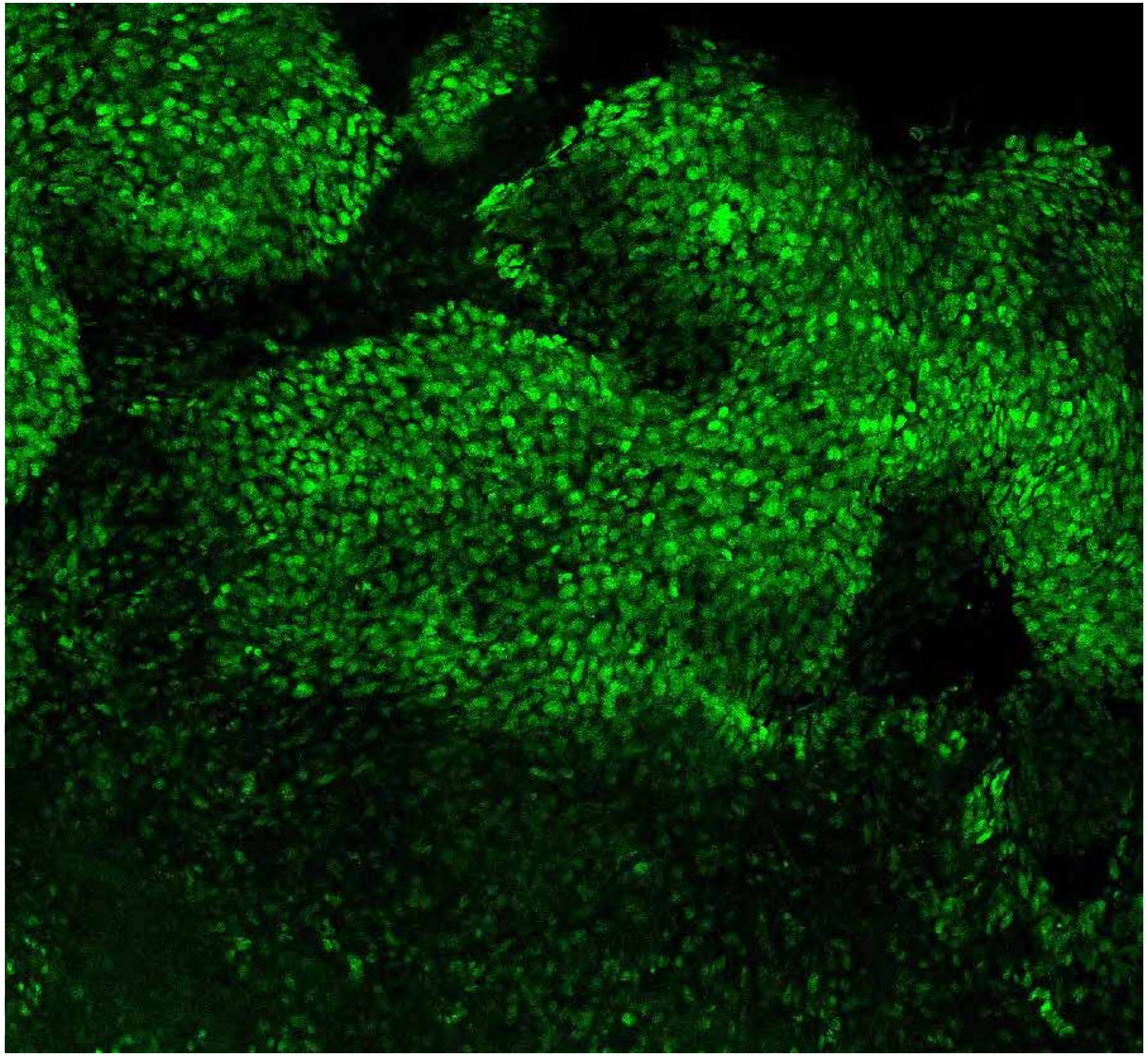

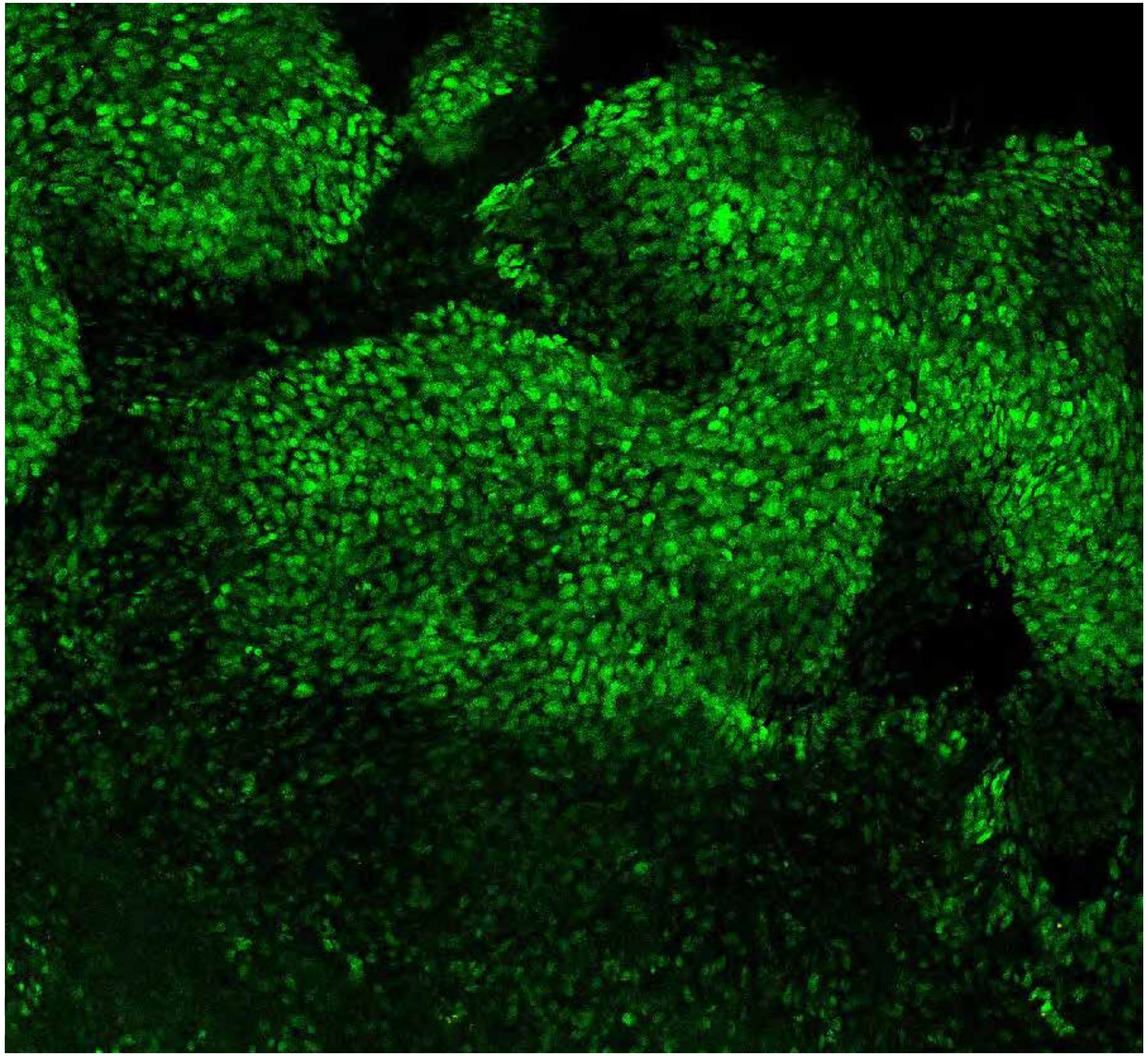

**PARPi-FL only (green)**

**PARPi-FL (green) + Autofluorescence (red/yellow)**

1472

### Close-up

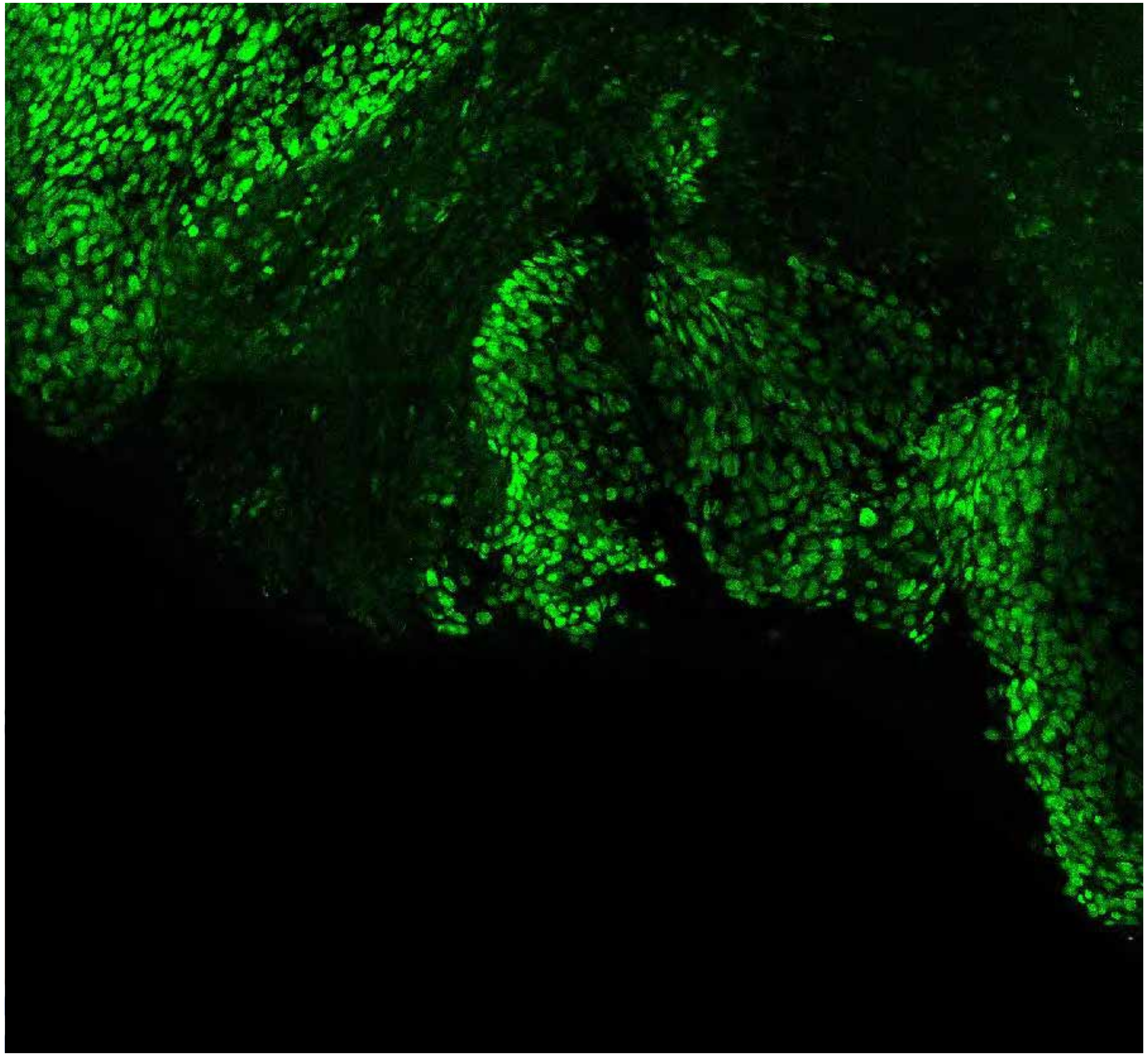

PARPi-FL only (green)

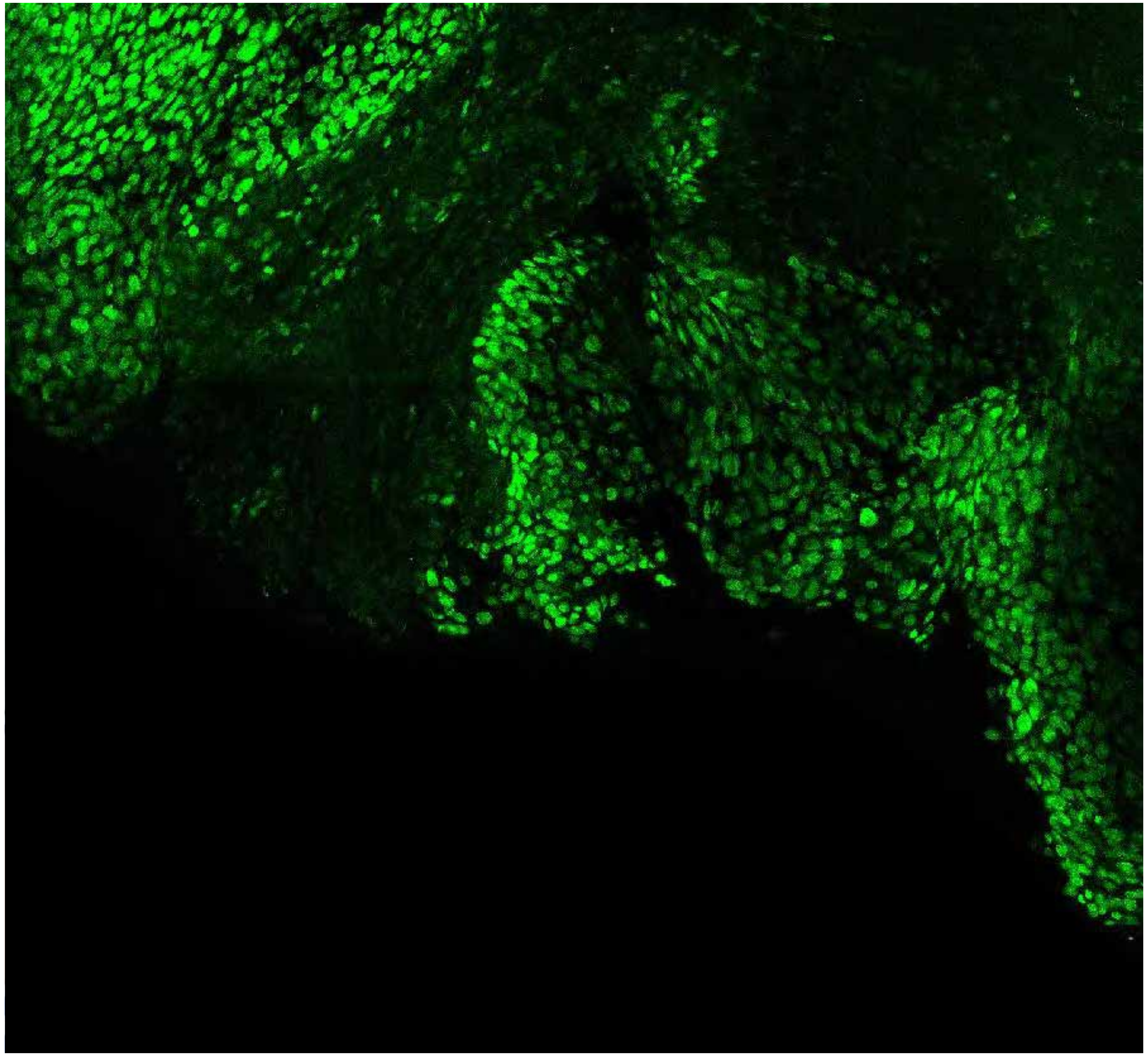

PARPi-FL (green) + Autofluorescence (red/yellow)

1472

### Close-up

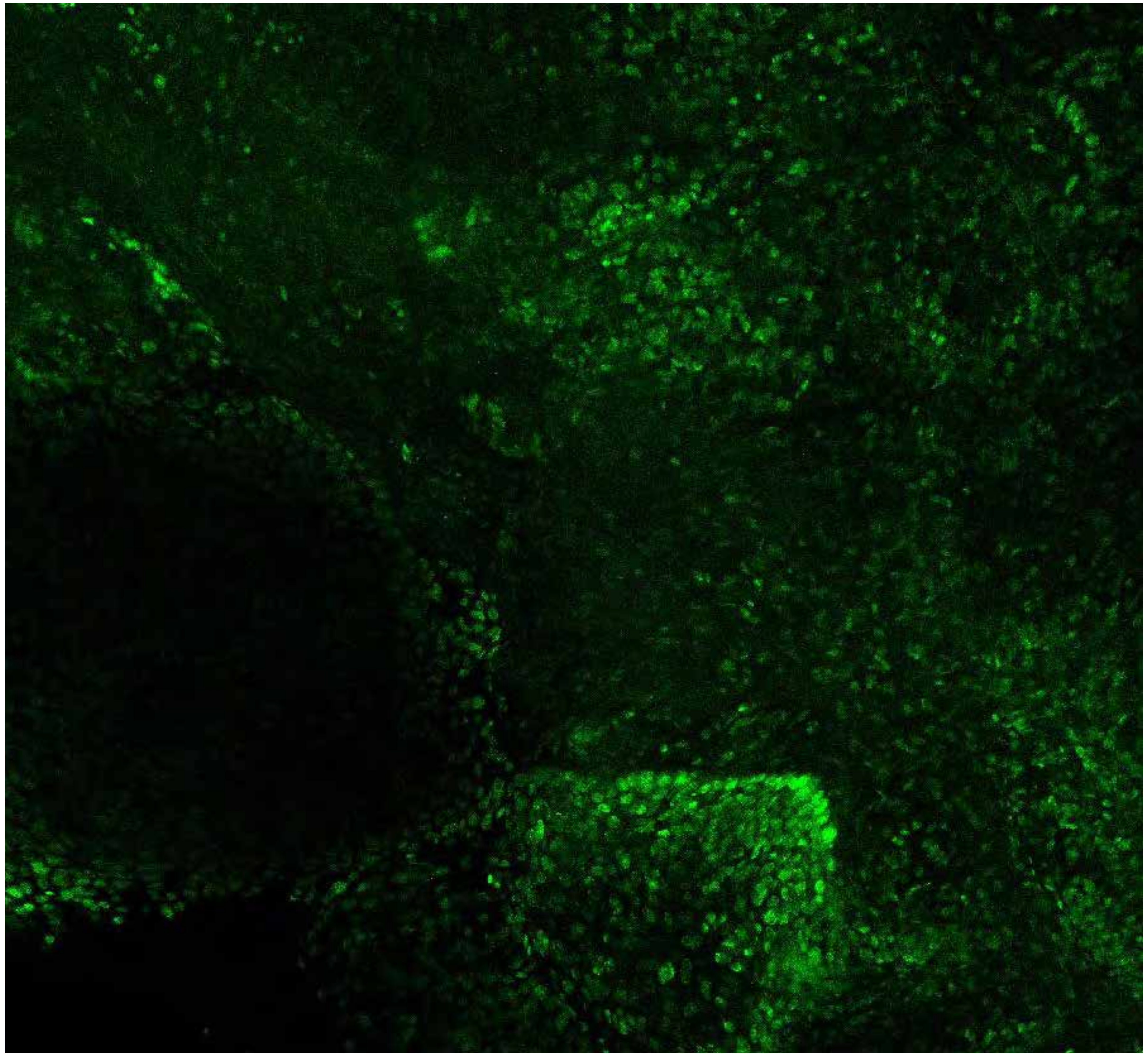

PARPi-FL only (green)

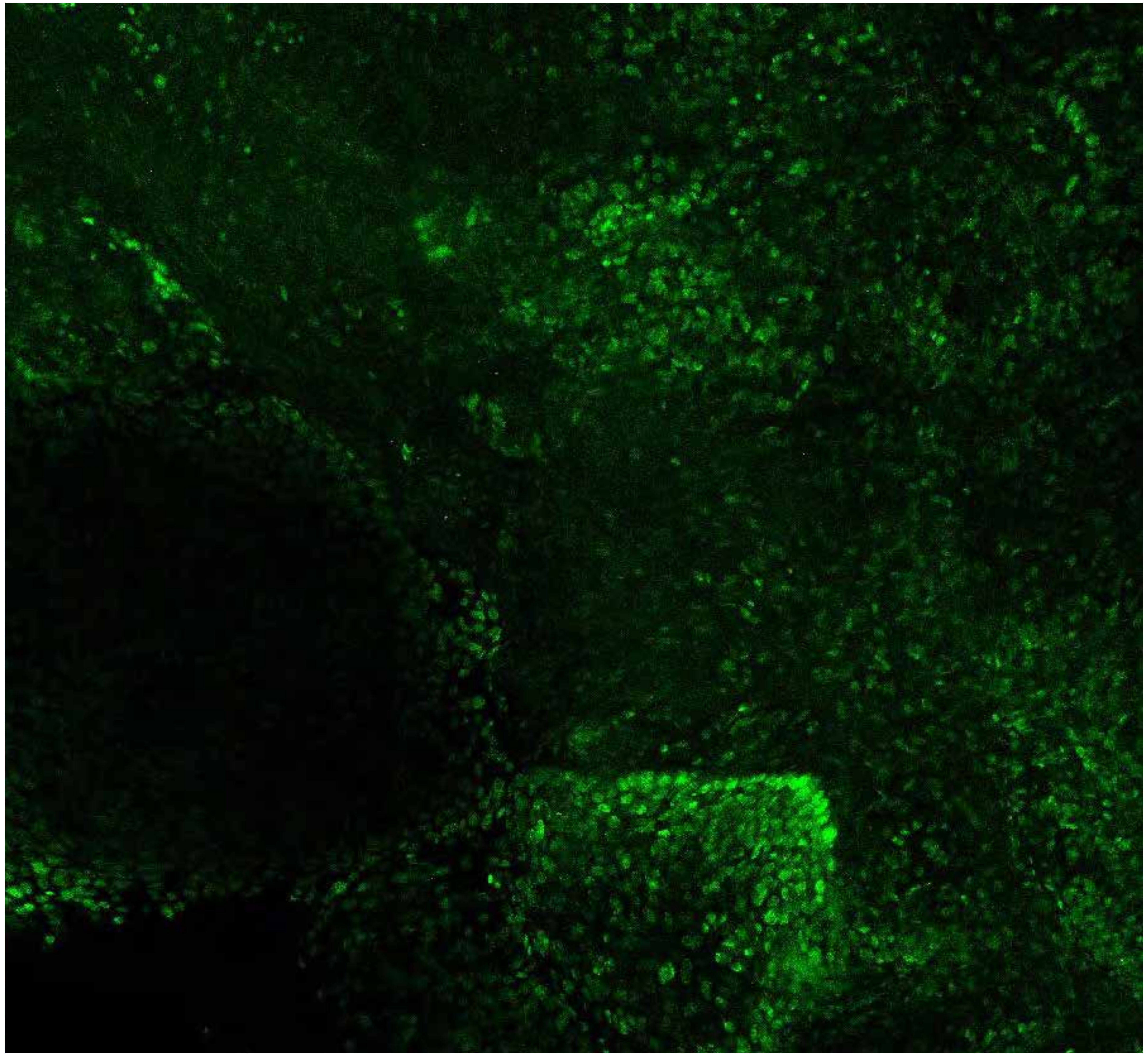

PARPi-FL (green) + Autofluorescence (red/yellow)

1472

10

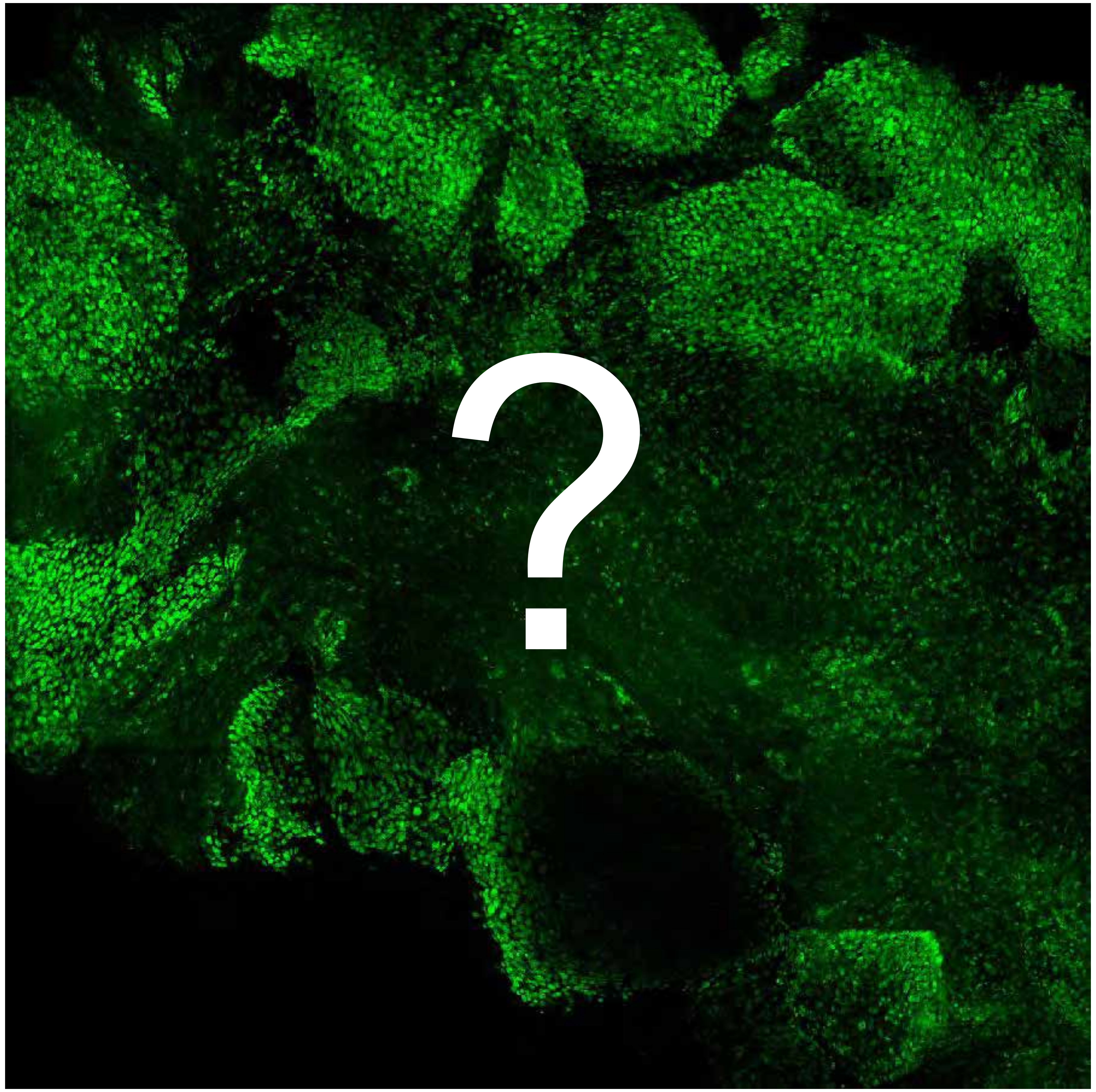


Tumor

Margin

1732

### Overview

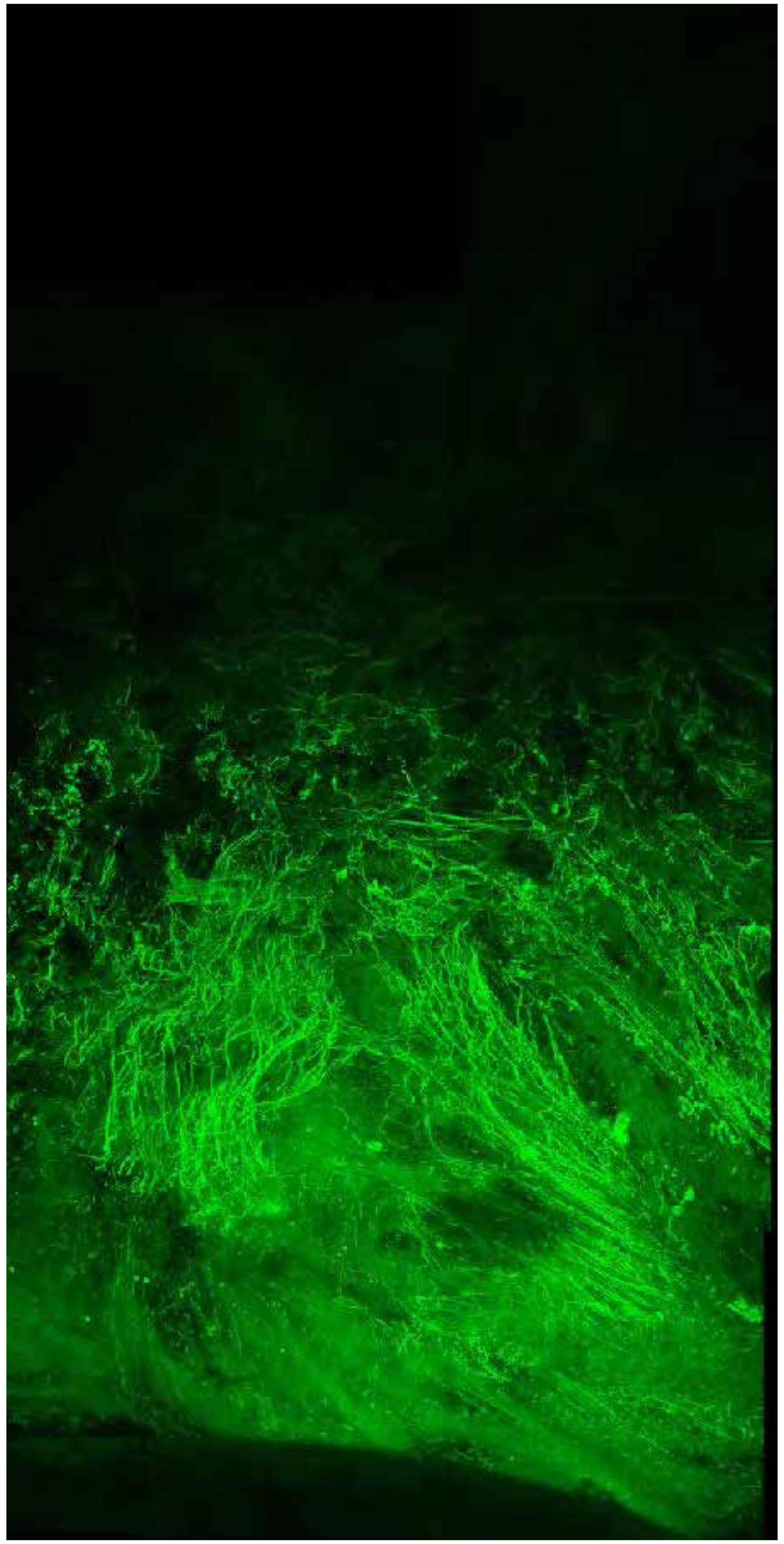

PARPi-FL only (green)

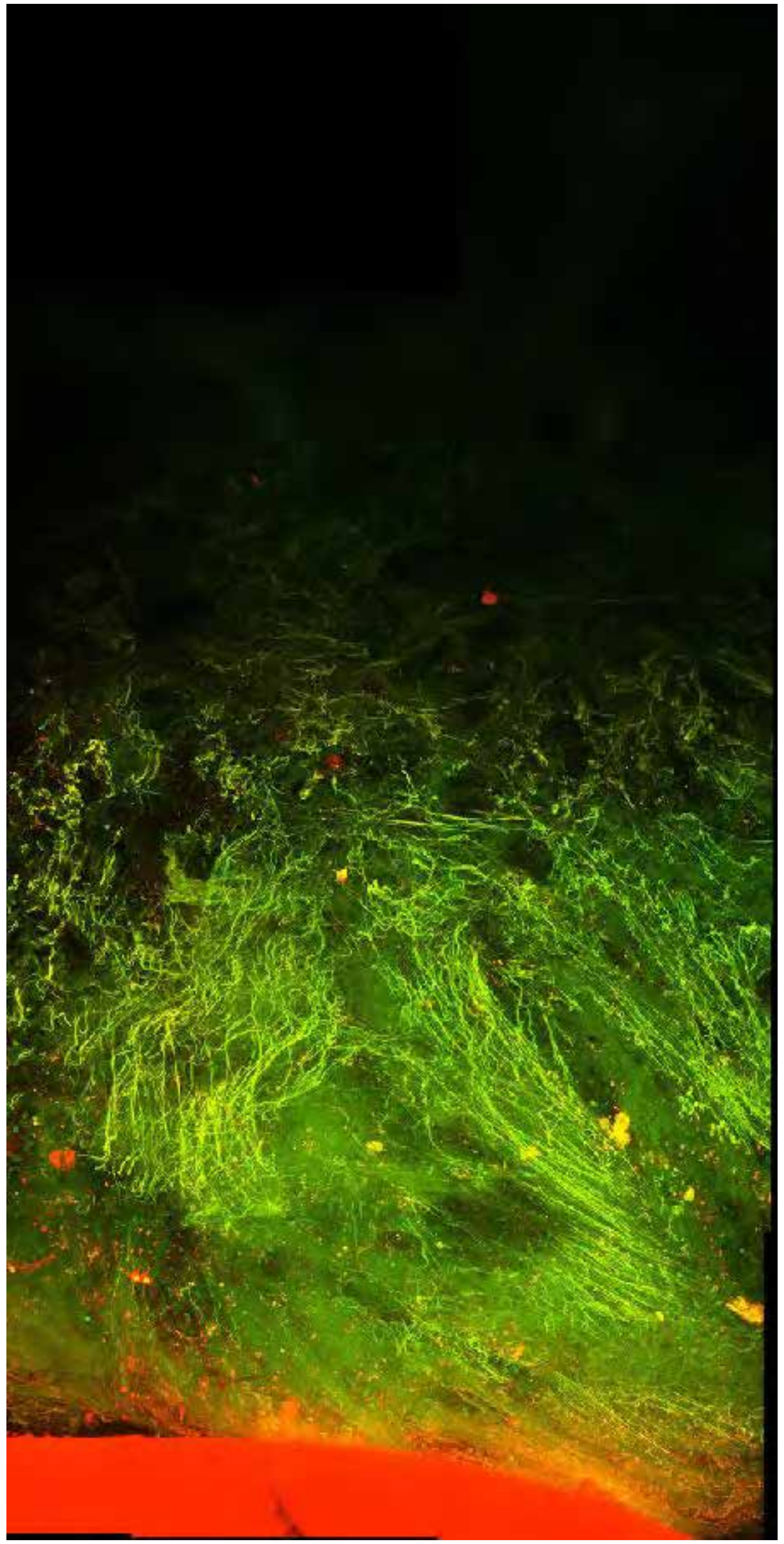

PARPi-FL (green) + Autofluorescence (red/yellow)

1732

### Close-up

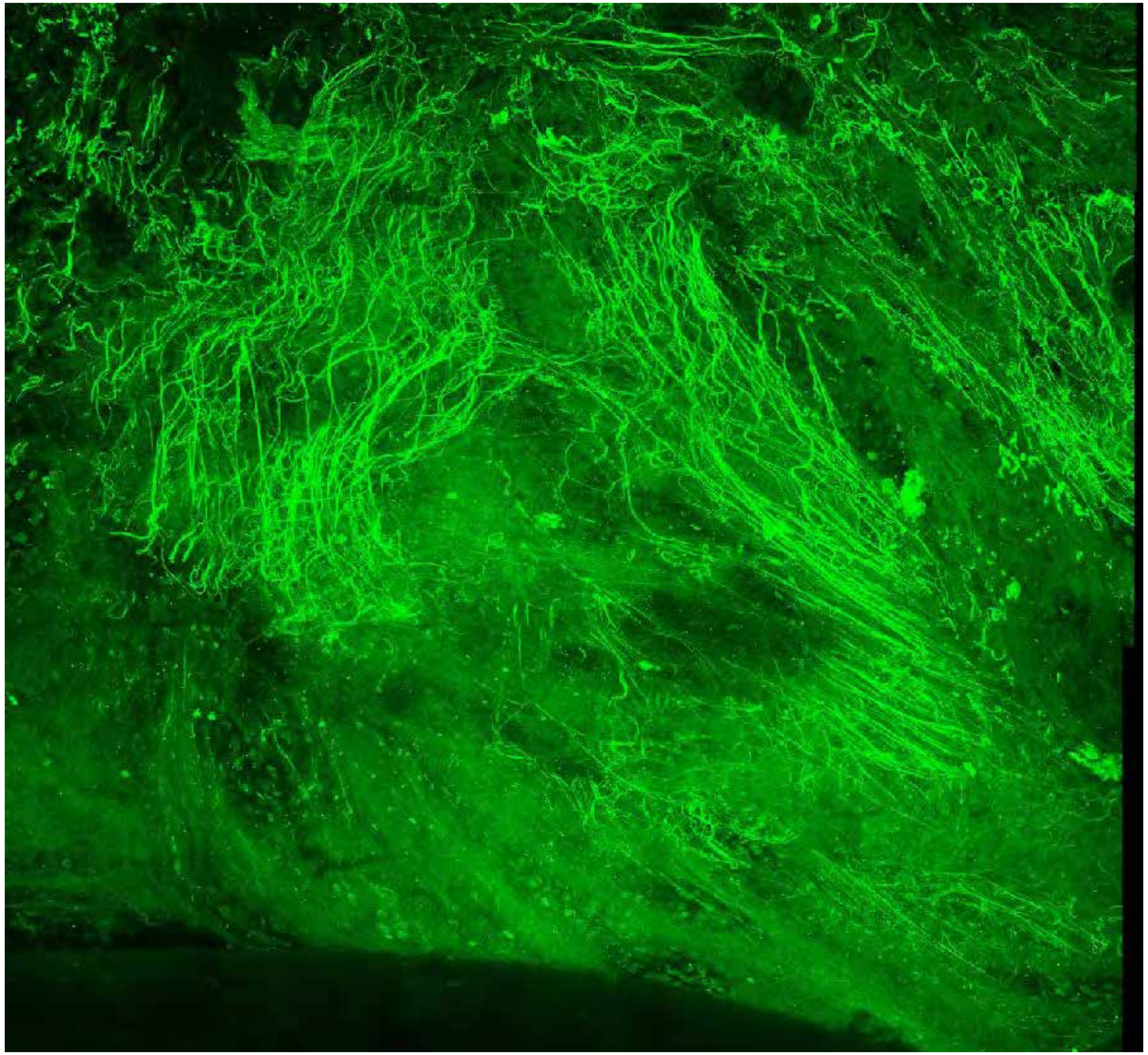

PARPi-FL only (green)

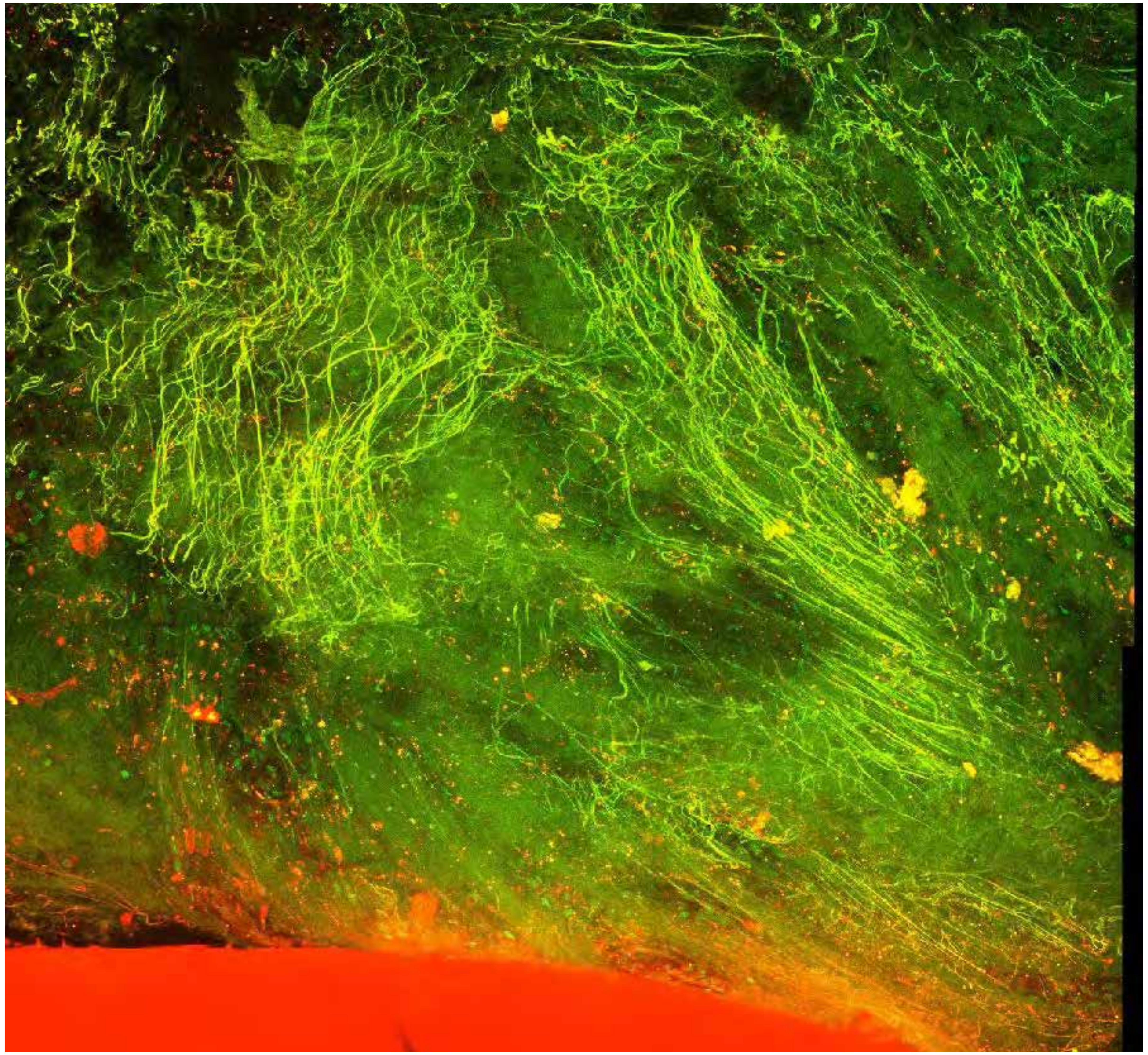

PARPi-FL (green) + Autofluorescence (red/yellow)

1732



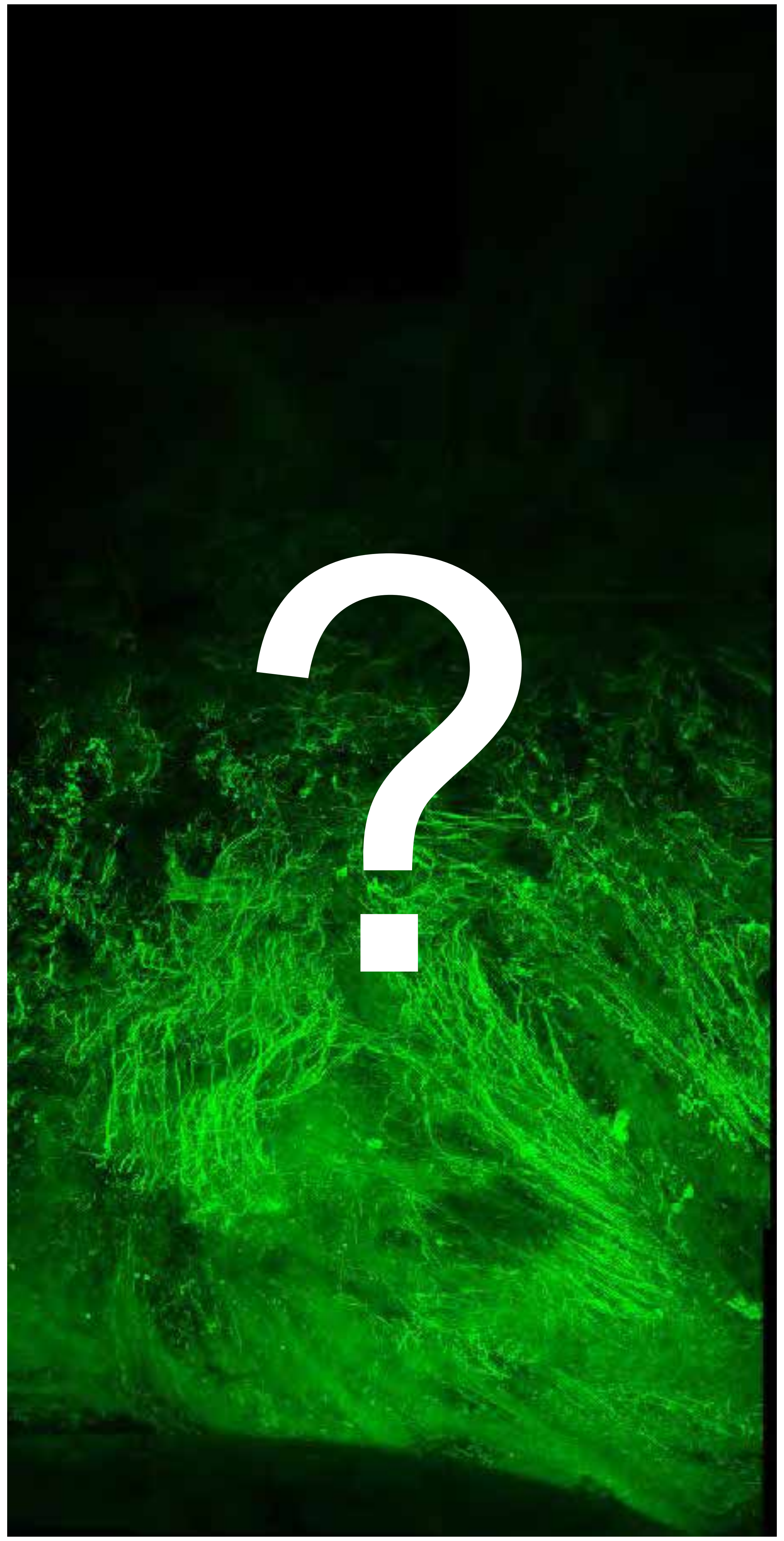


Tumor

Margin

2154

### Overview

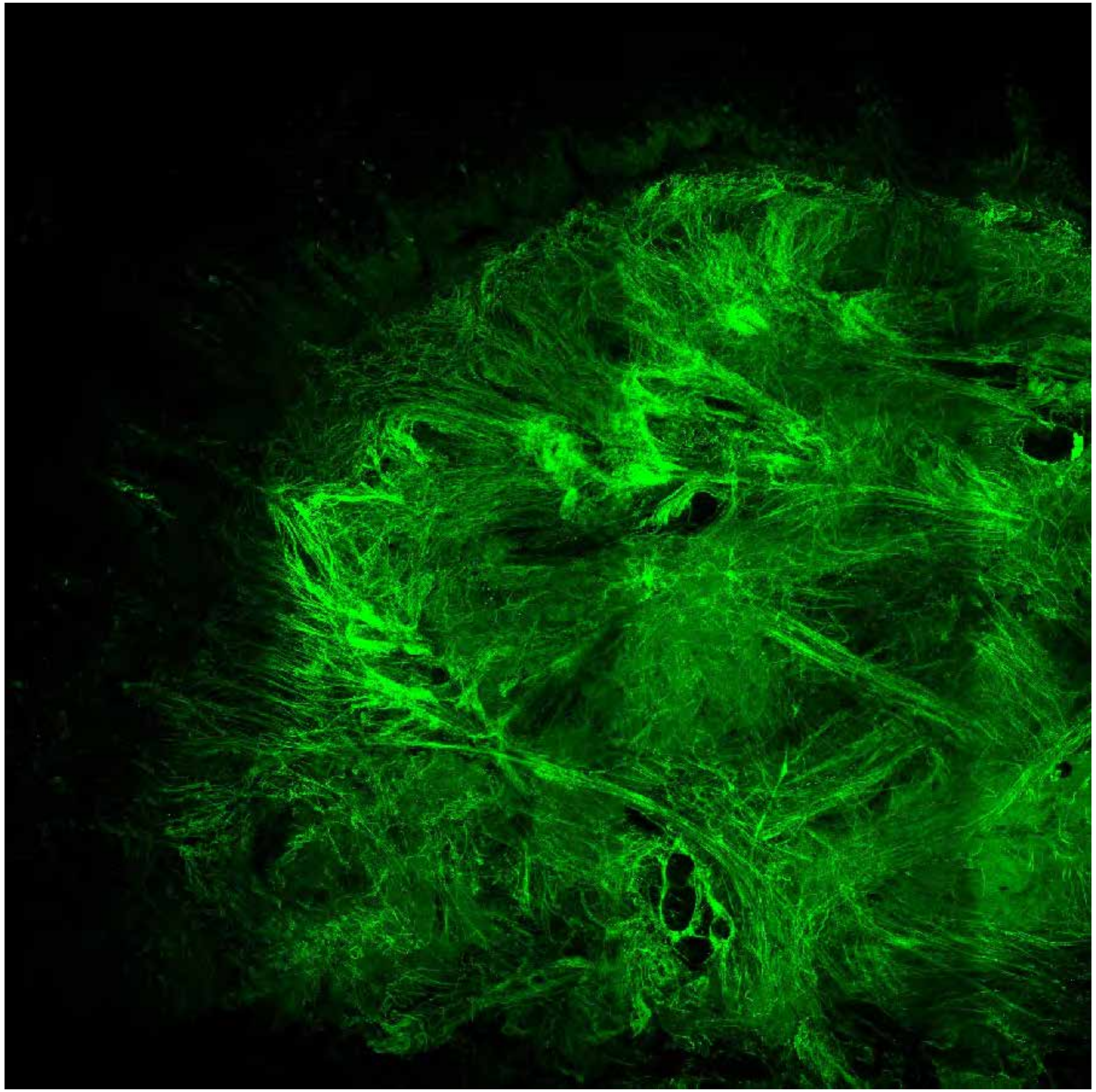

PARPi-FL only (green)

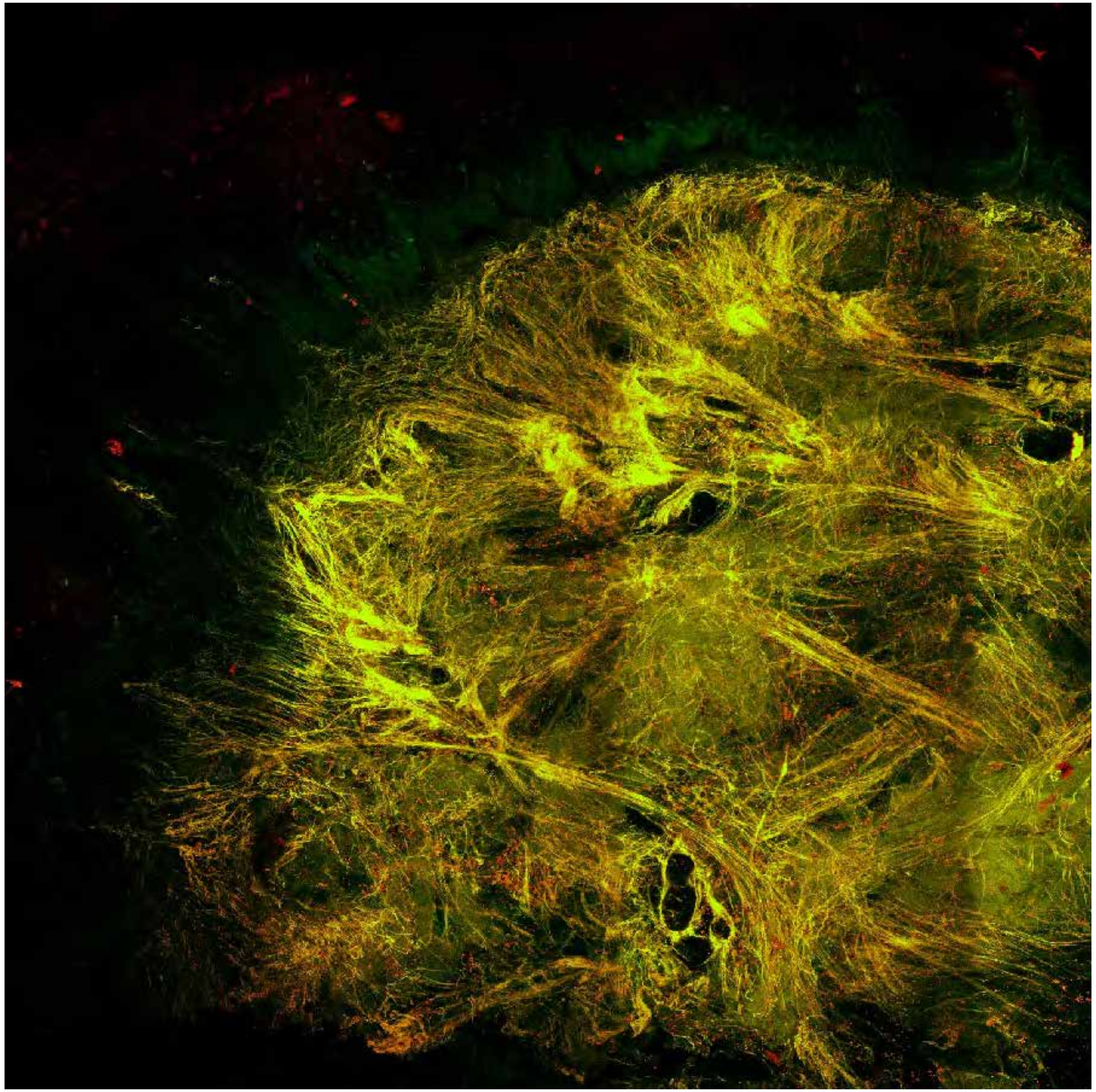

PARPi-FL (green) + Autofluorescence (red/yellow)

2154

### Close-up

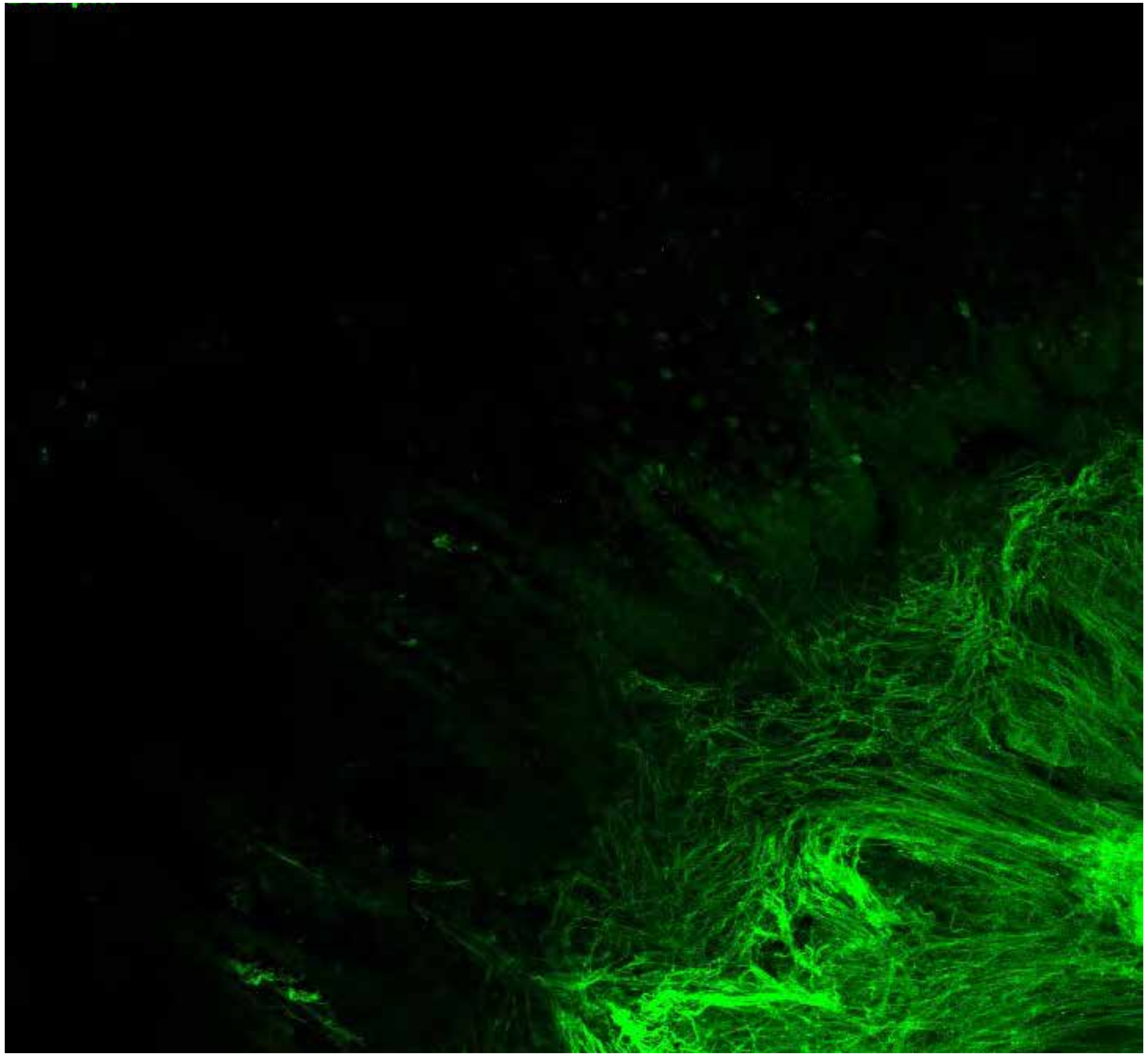

PARPi-FL only (green)

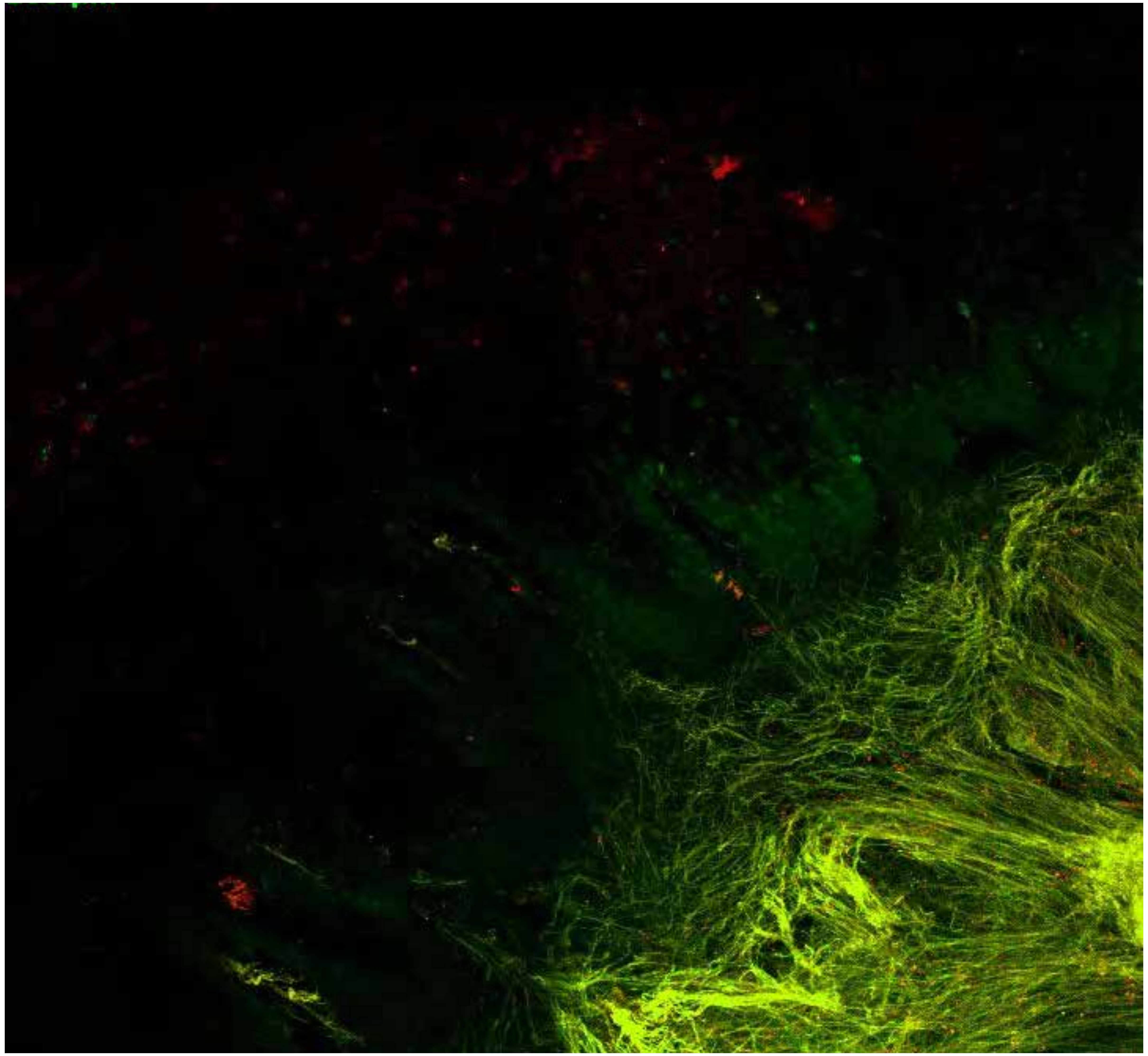

PARPi-FL (green) + Autofluorescence (red/yellow)

2154

### Close-up

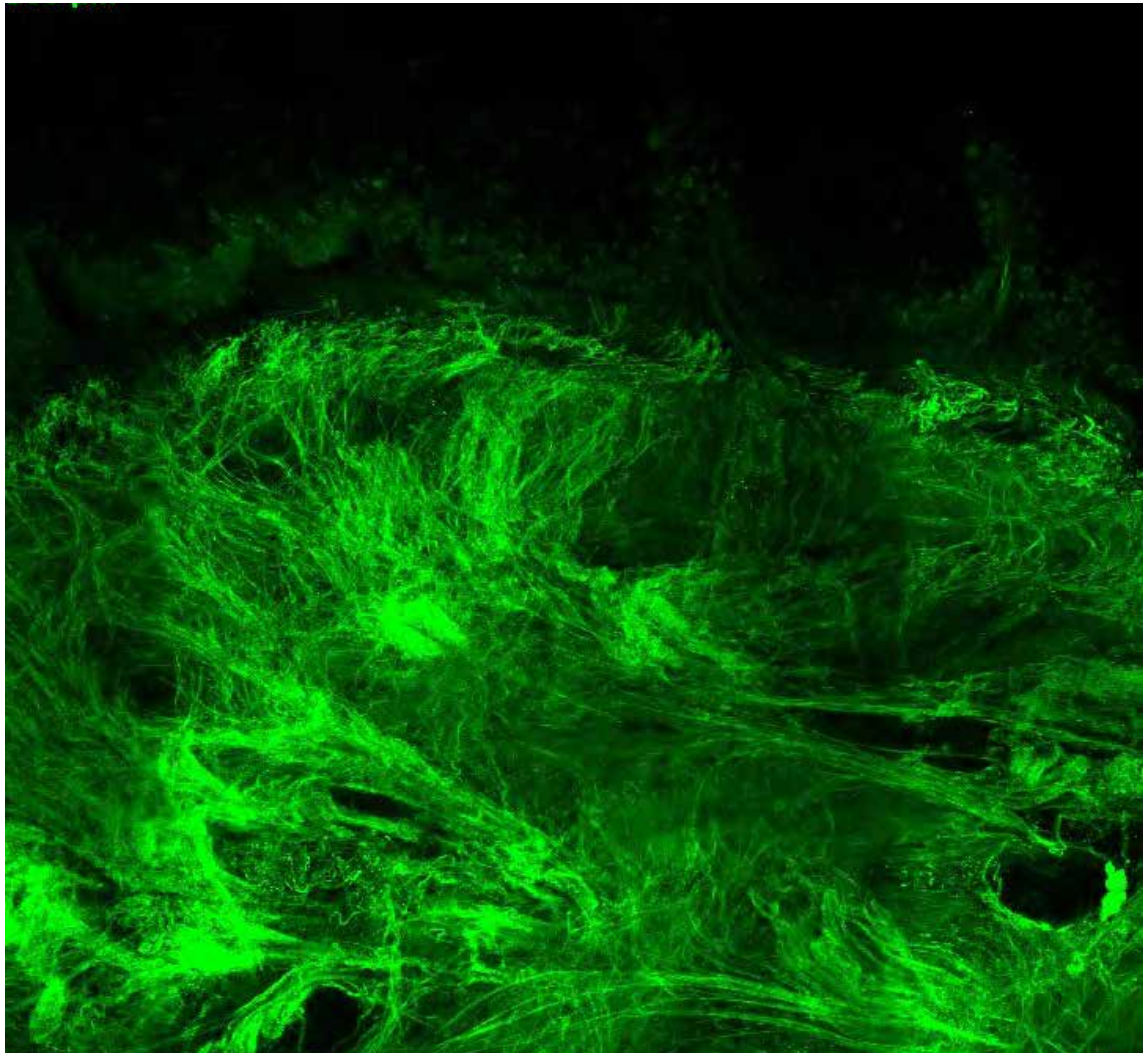

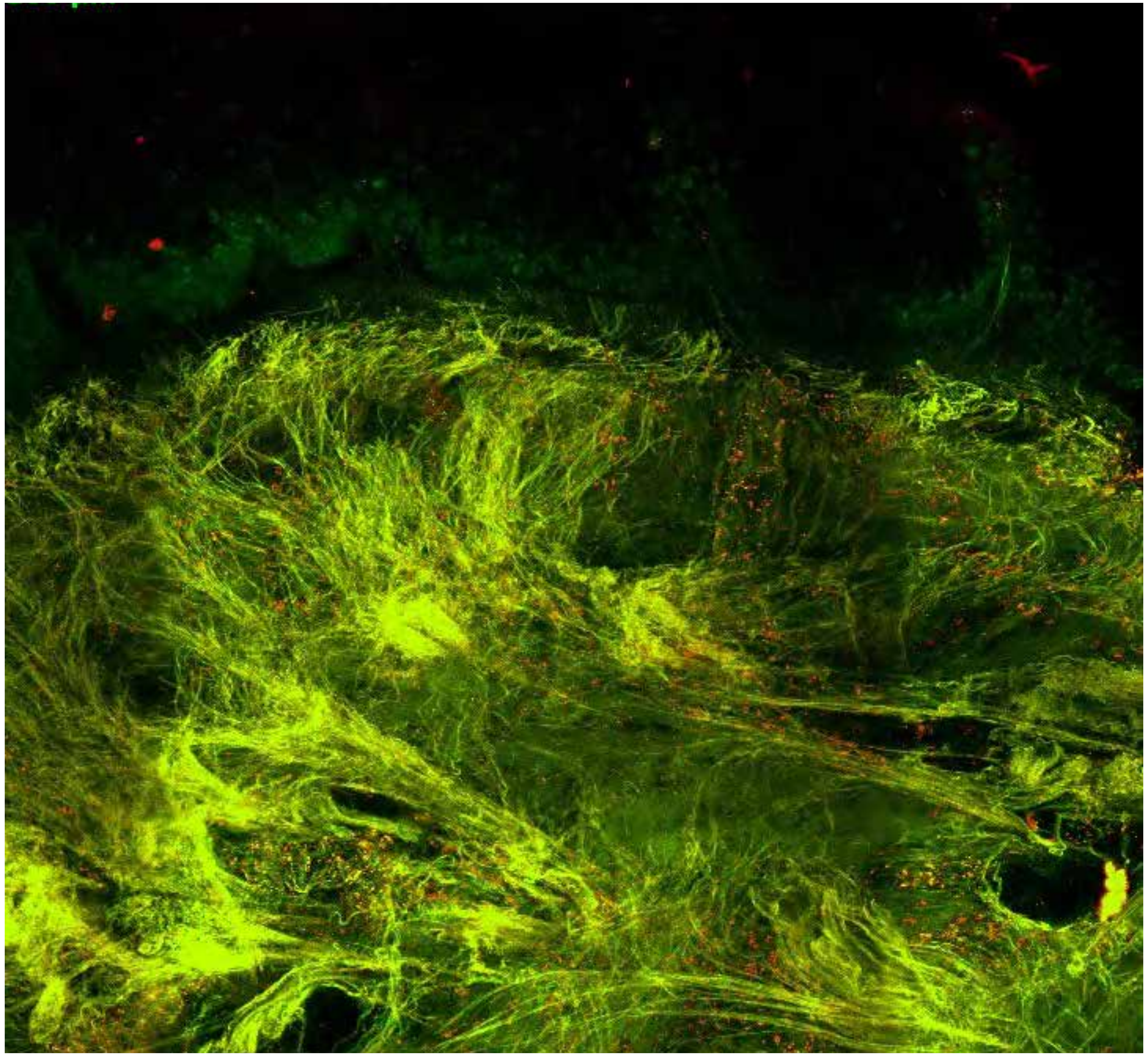

**PARPi-FL only (green)**

**PARPi-FL (green) + Autofluorescence (red/yellow)**

2154

### Close-up

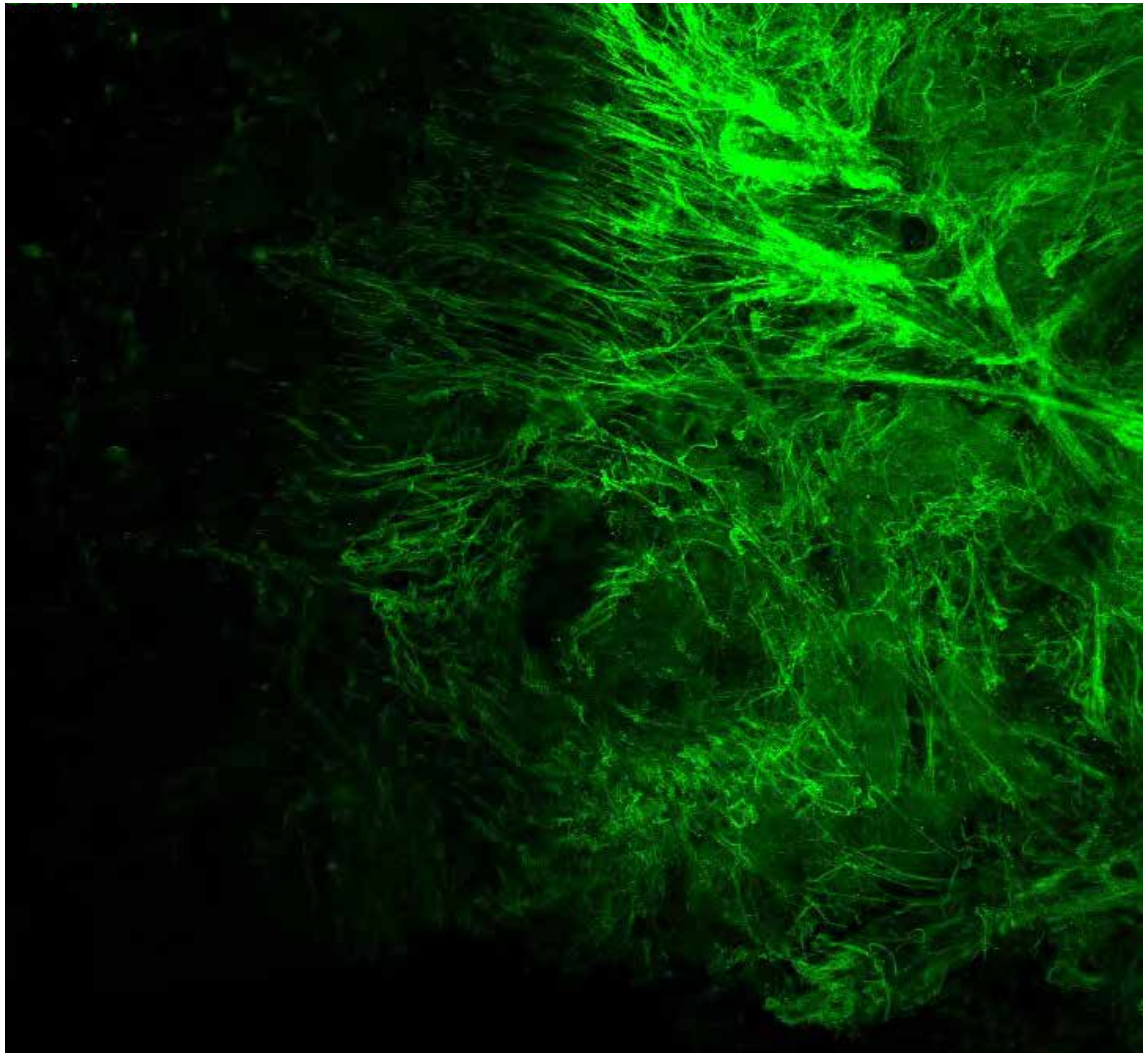

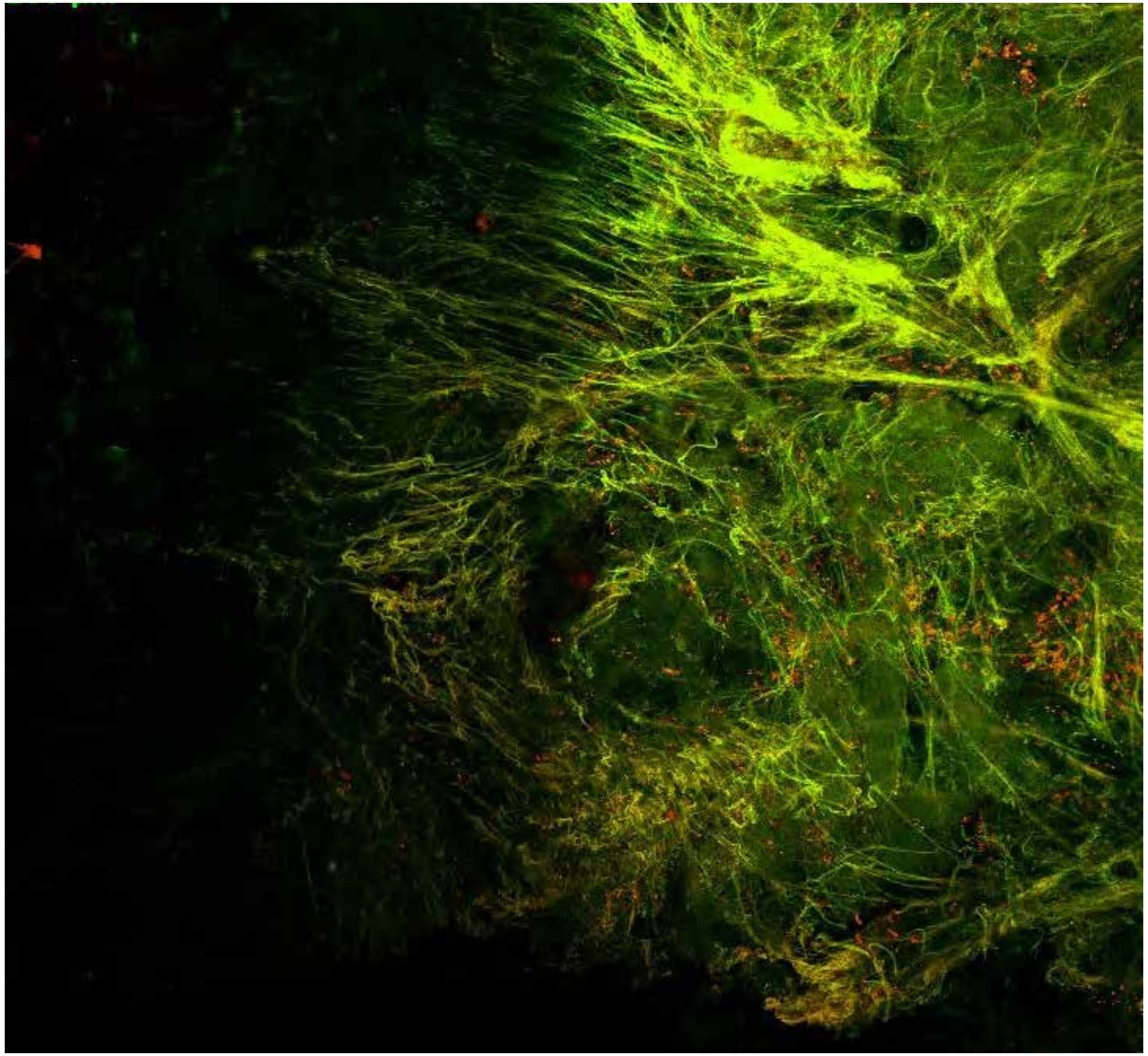

PARPi-FL only (green)

PARPi-FL (green) + Autofluorescence (red/yellow)

2154

### Close-up

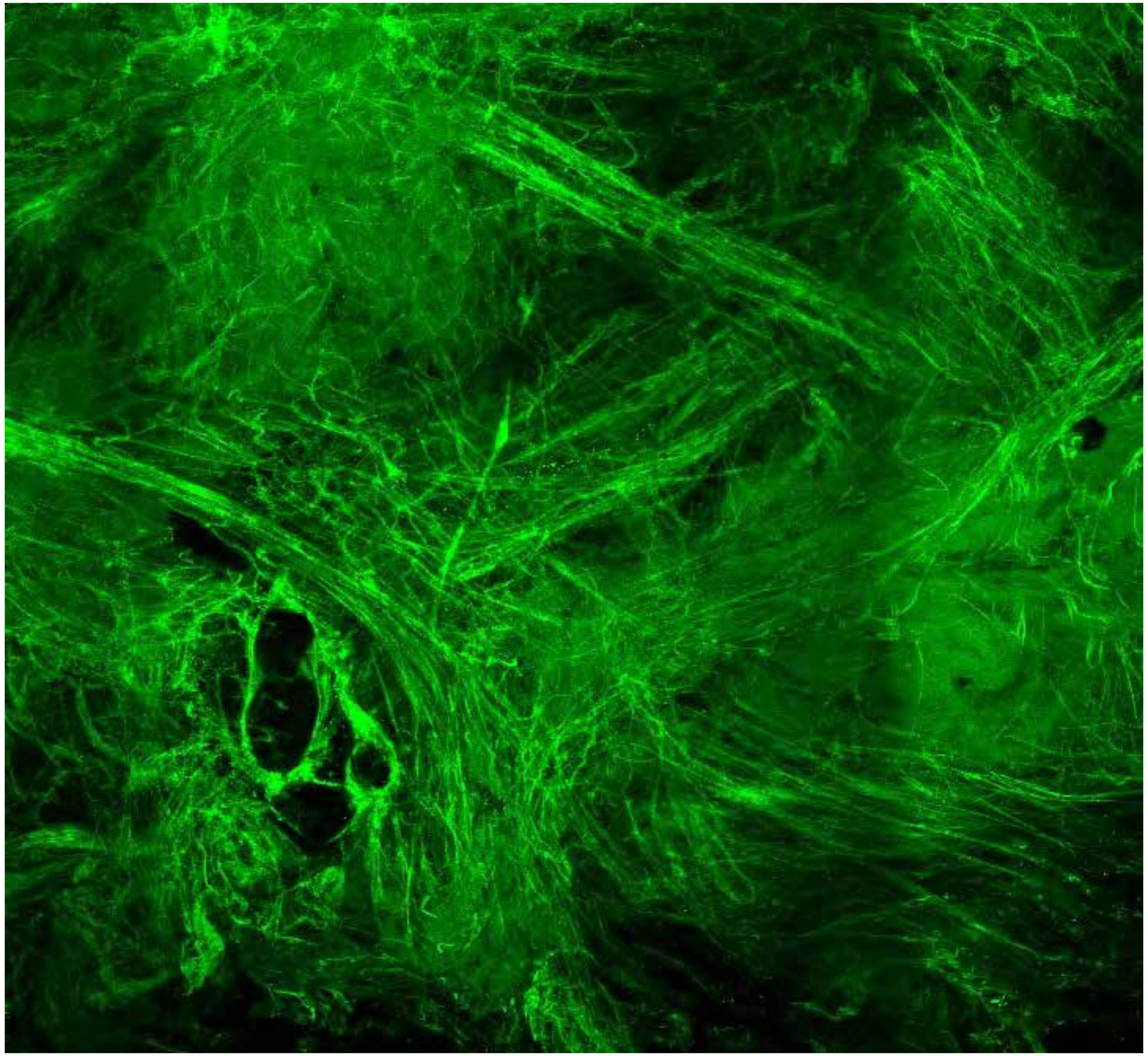

PARPi-FL only (green)

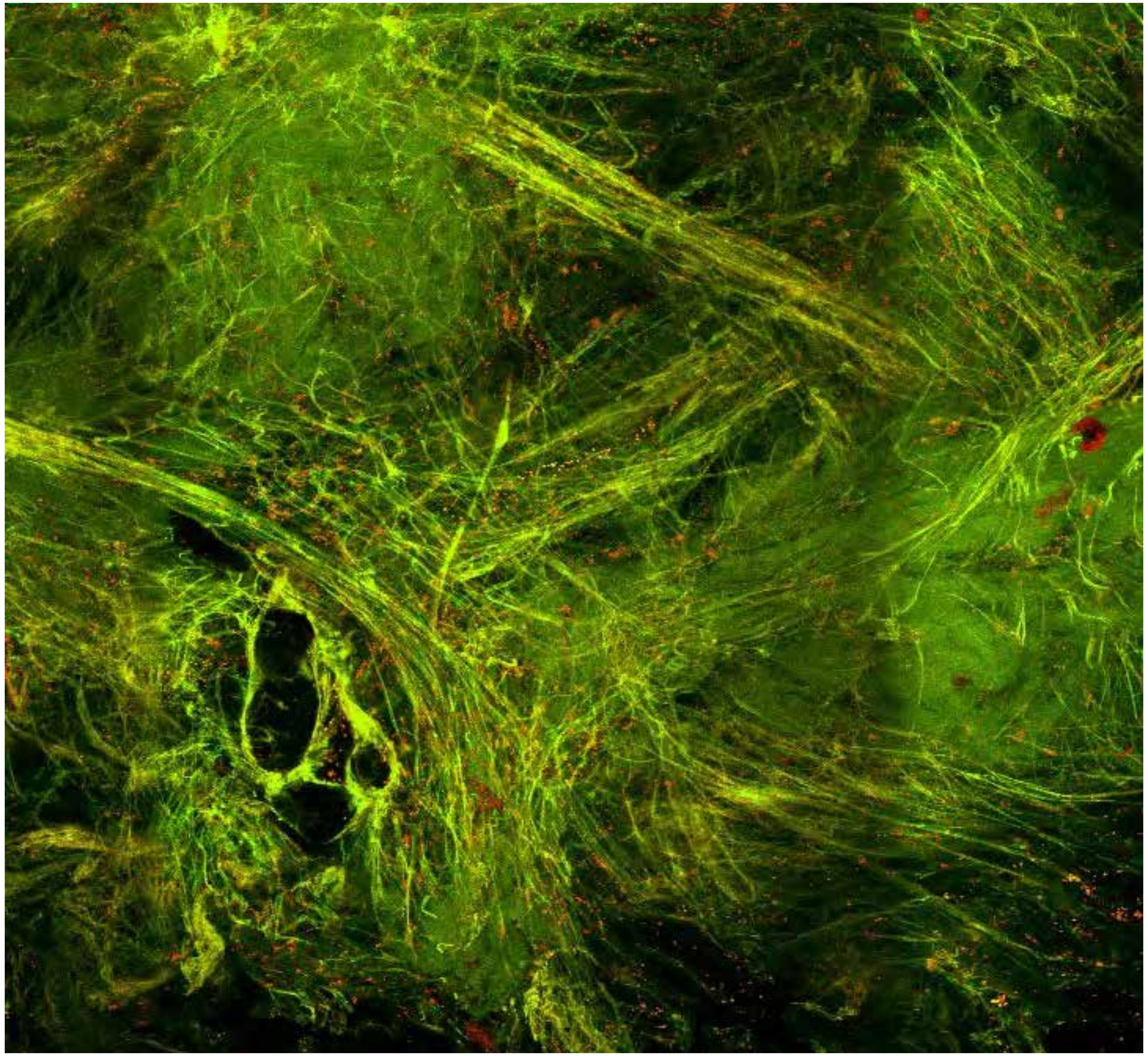

PARPi-FL (green) + Autofluorescence (red/yellow)

2154

10

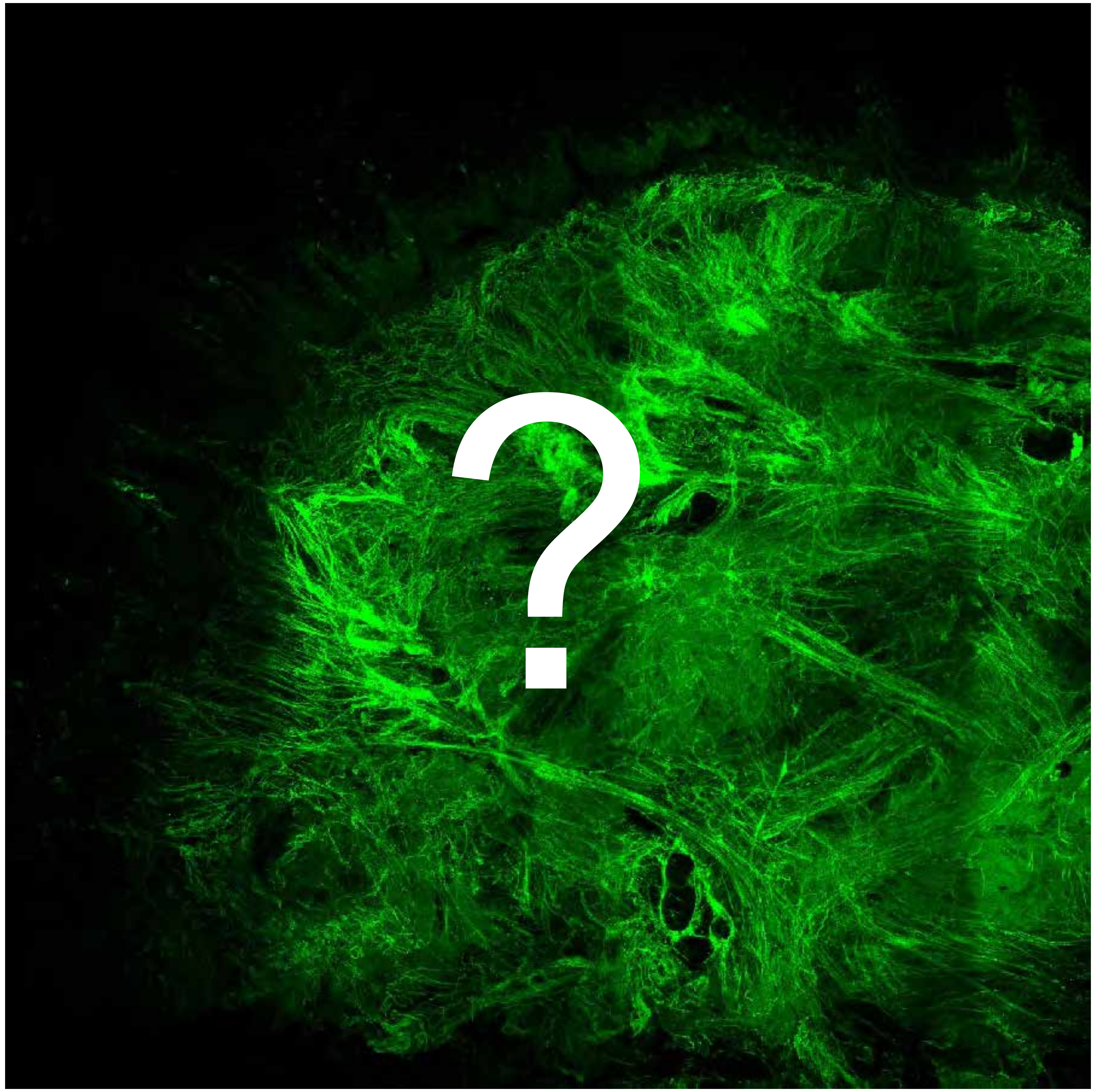


Tumor

Margin

2225

### Overview

Not available

Not available

**PARPi-FL only (green)**

**PARPi-FL (green) + Autofluorescence (red/yellow)**

2225

### Close-up

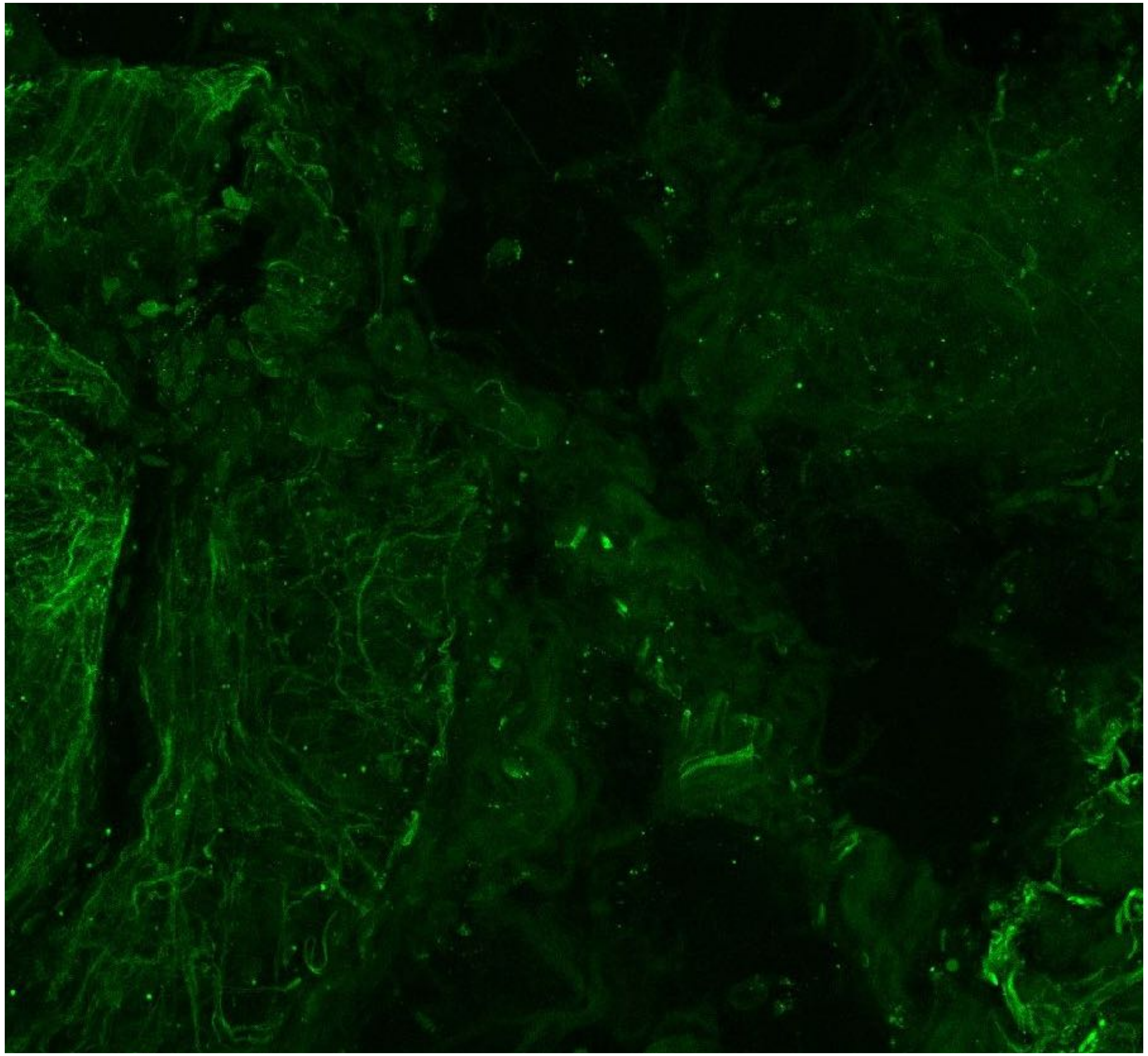

**PARPi-FL only (green)**

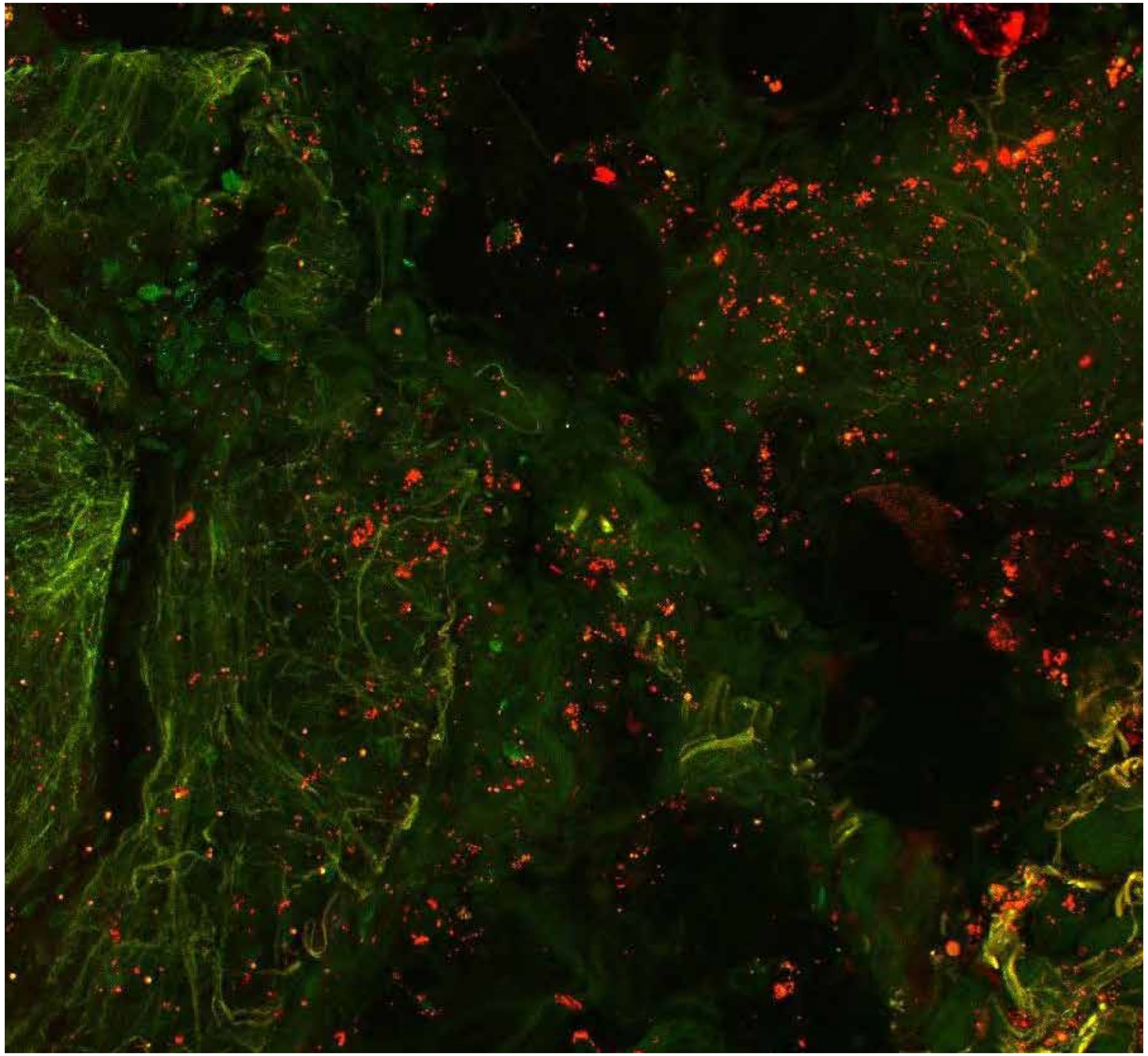

**PARPi-FL (green) + Autofluorescence (red/yellow)**

2225

### Close-up

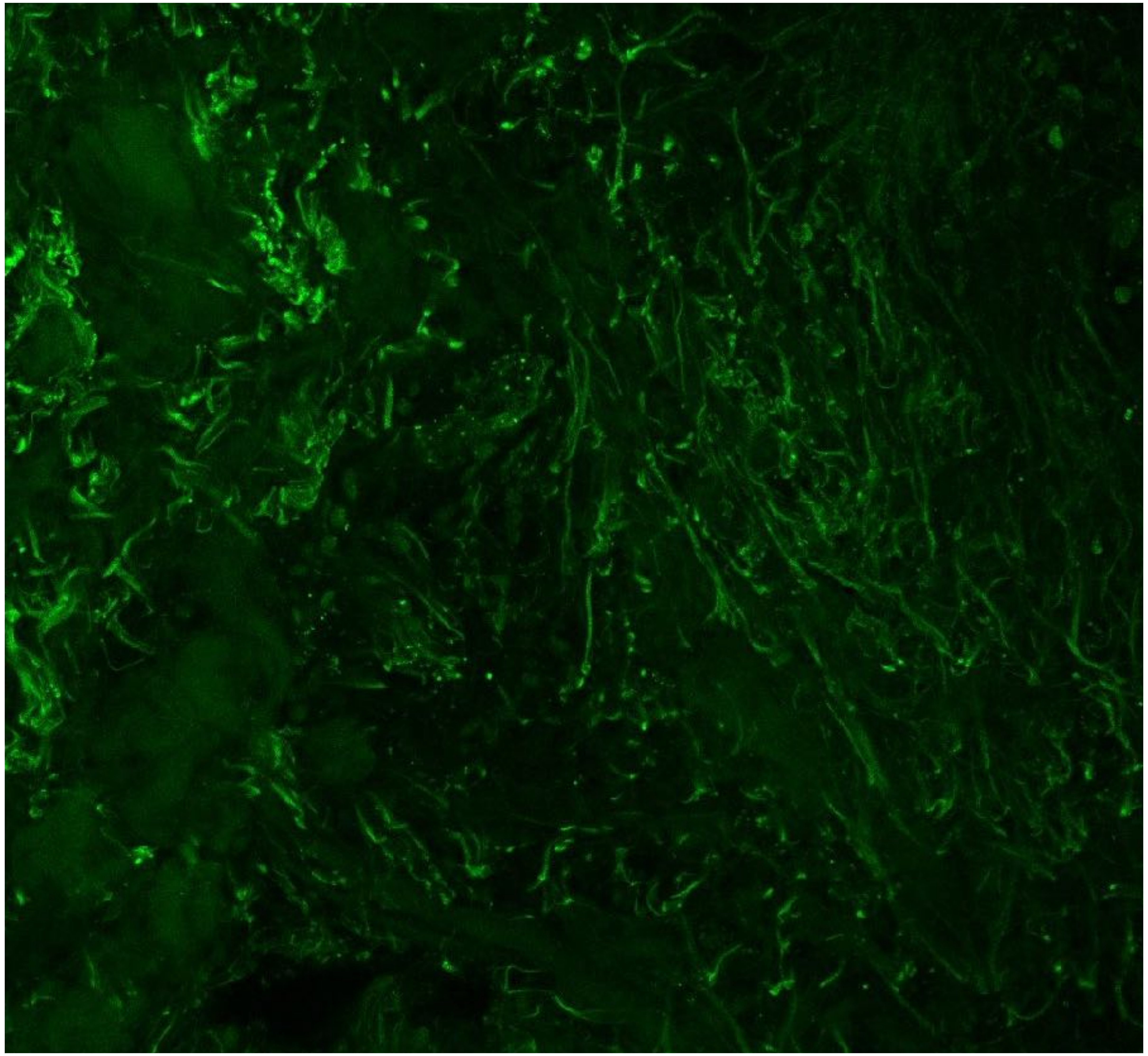

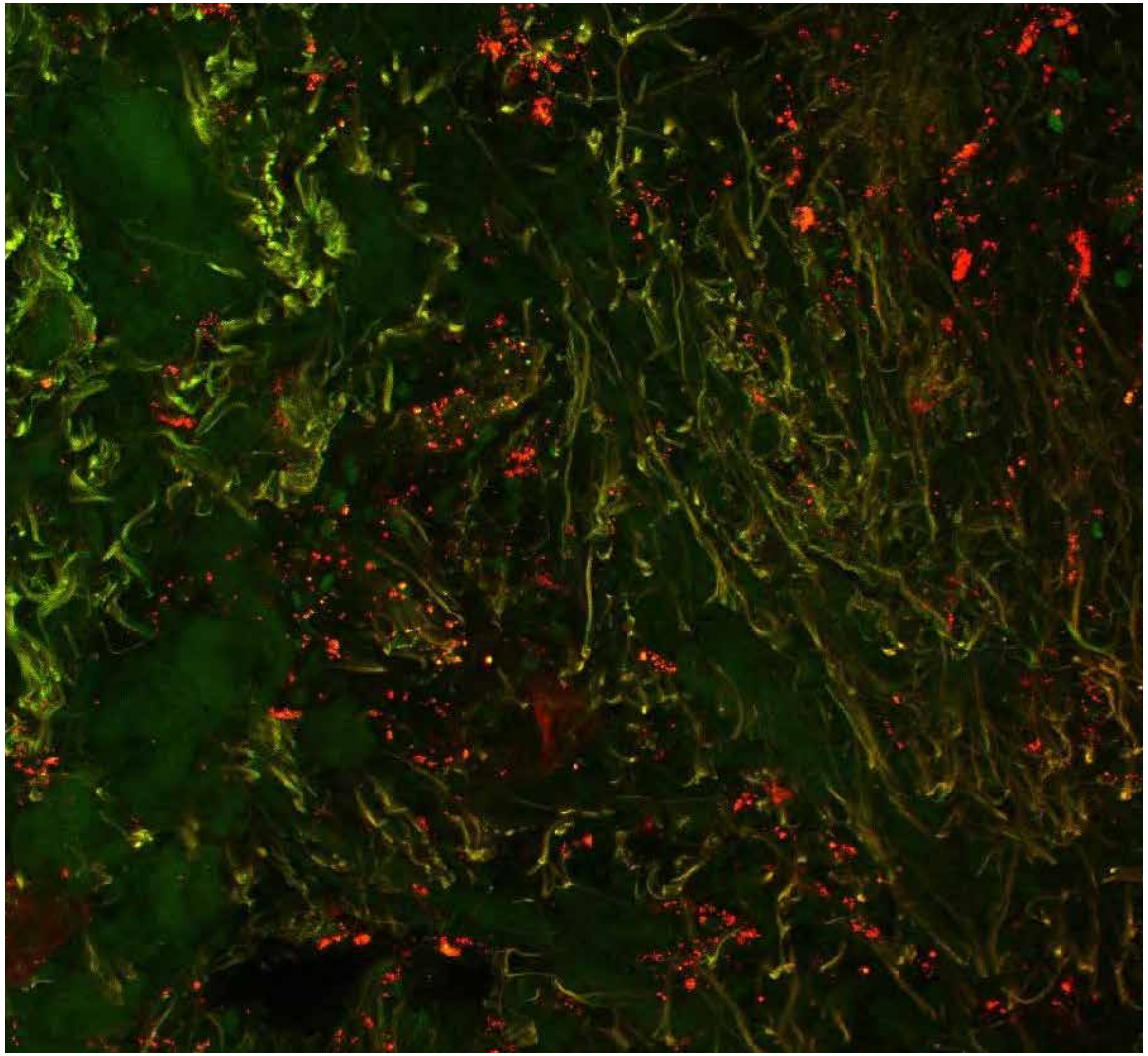

**PARPi-FL only (green)**

**PARPi-FL (green) + Autofluorescence (red/yellow)**

2225

### Close-up

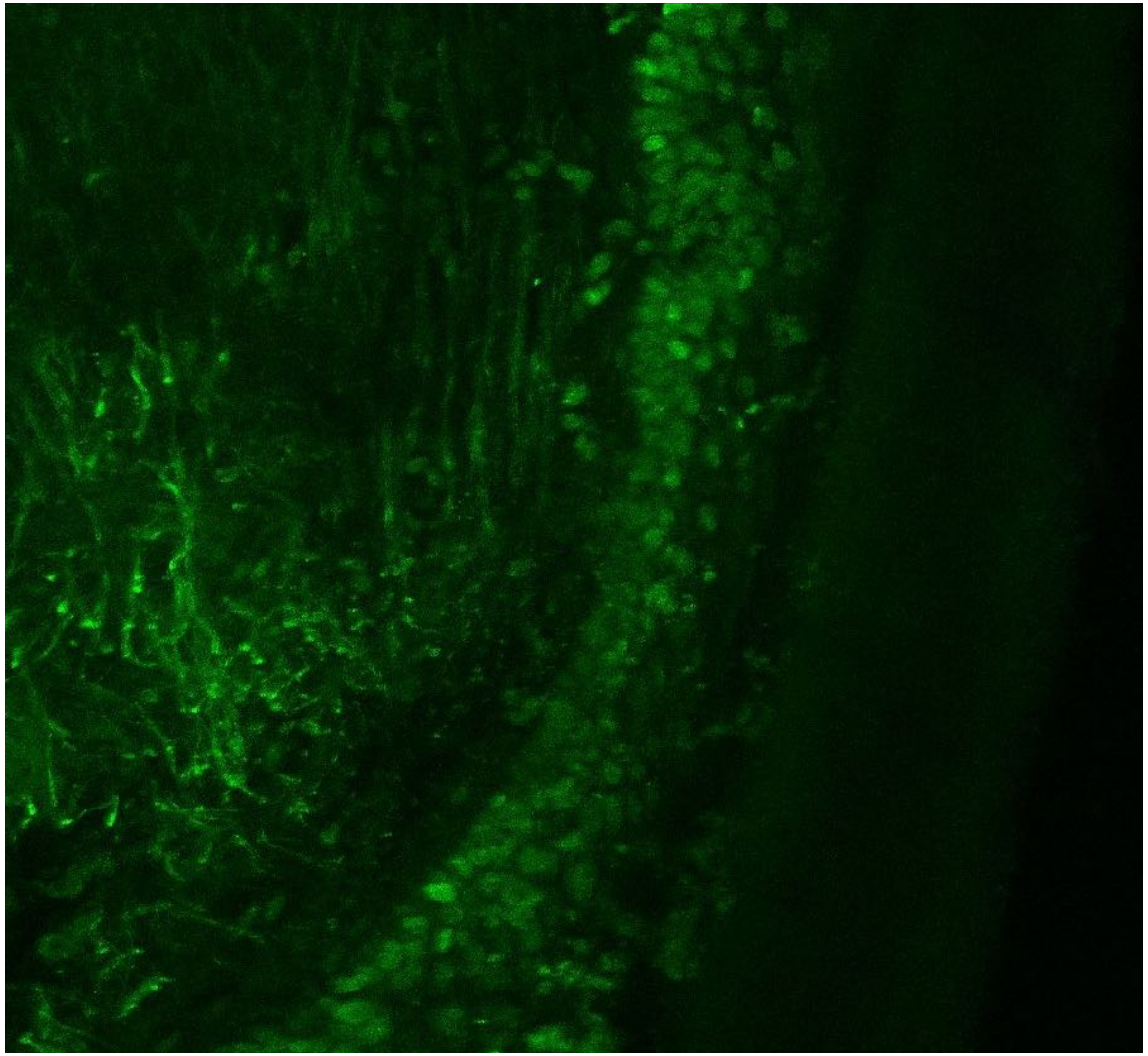

PARPi-FL only (green)

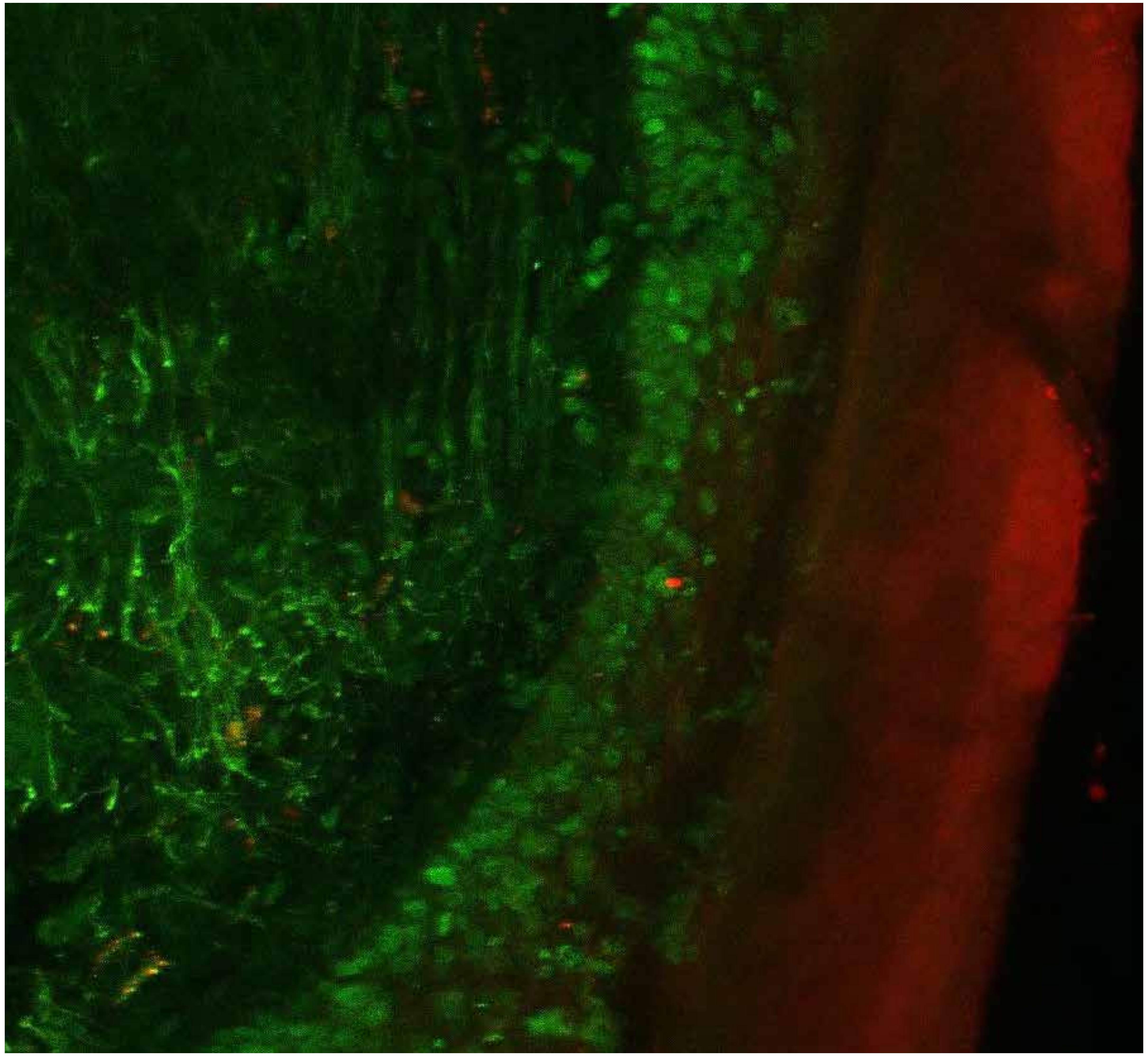

PARPi-FL (green) + Autofluorescence (red/yellow)

2225

10

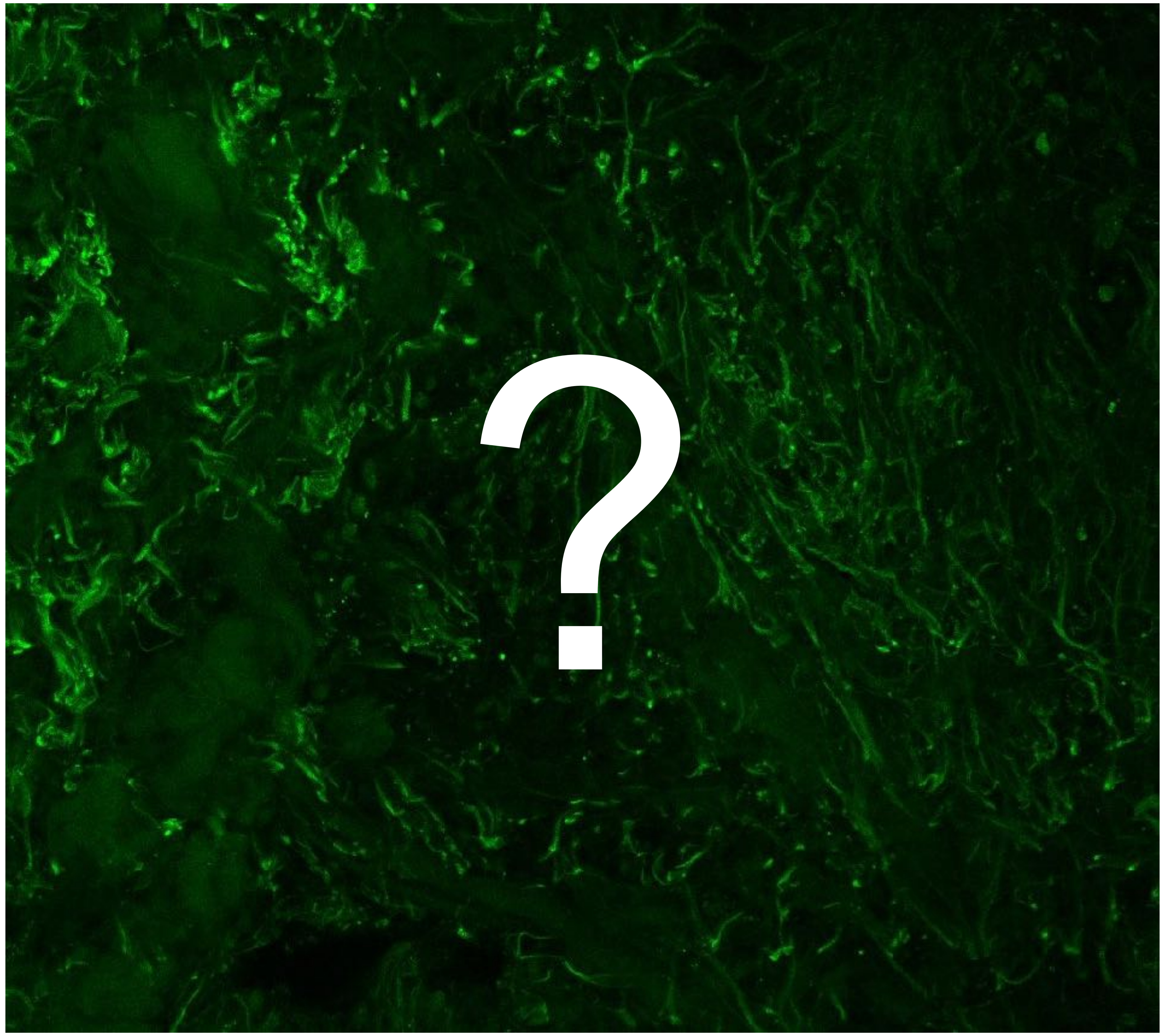


Tumor

Margin

2627

### Overview

Not available

Not available

**PARPi-FL only (green)**

**PARPi-FL (green) + Autofluorescence (red/yellow)**

2627

### Close-up

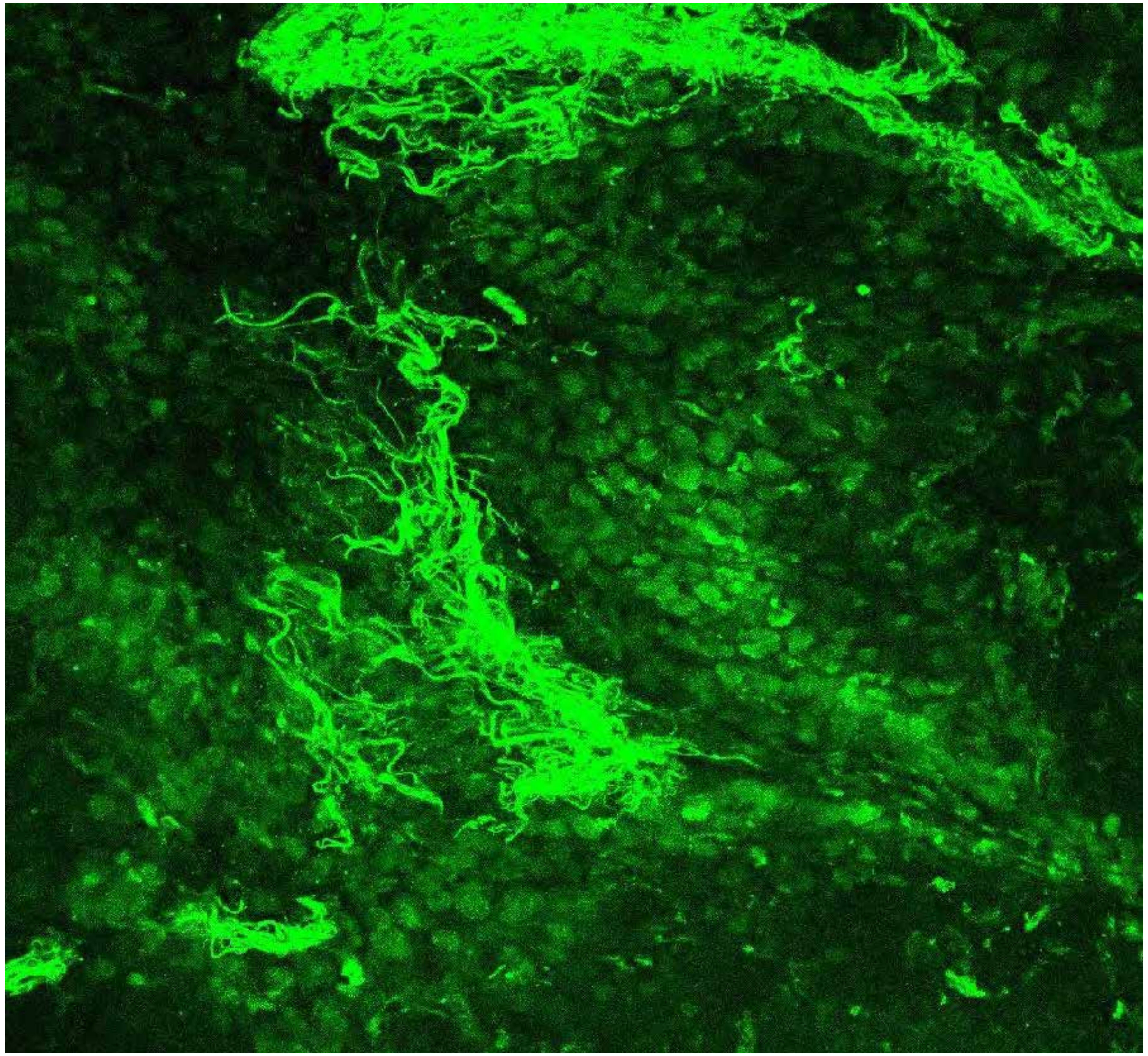

Not available

**PARPi-FL only (green)**

**PARPi-FL (green) + Autofluorescence (red/yellow)**

2627

### Close-up

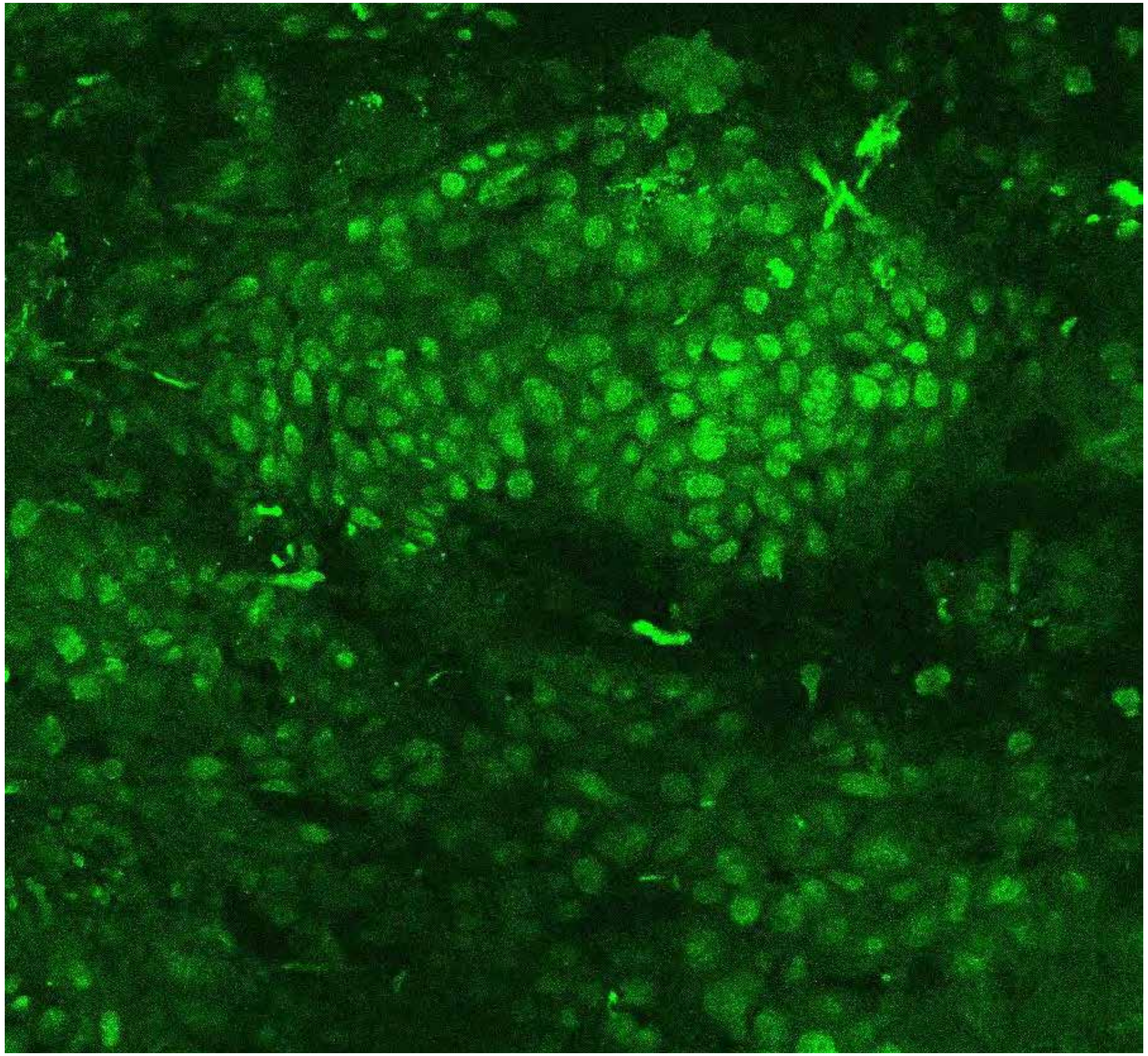

Not available

**PARPi-FL only (green)**

**PARPi-FL (green) + Autofluorescence (red/yellow)**

2627

### Close-up

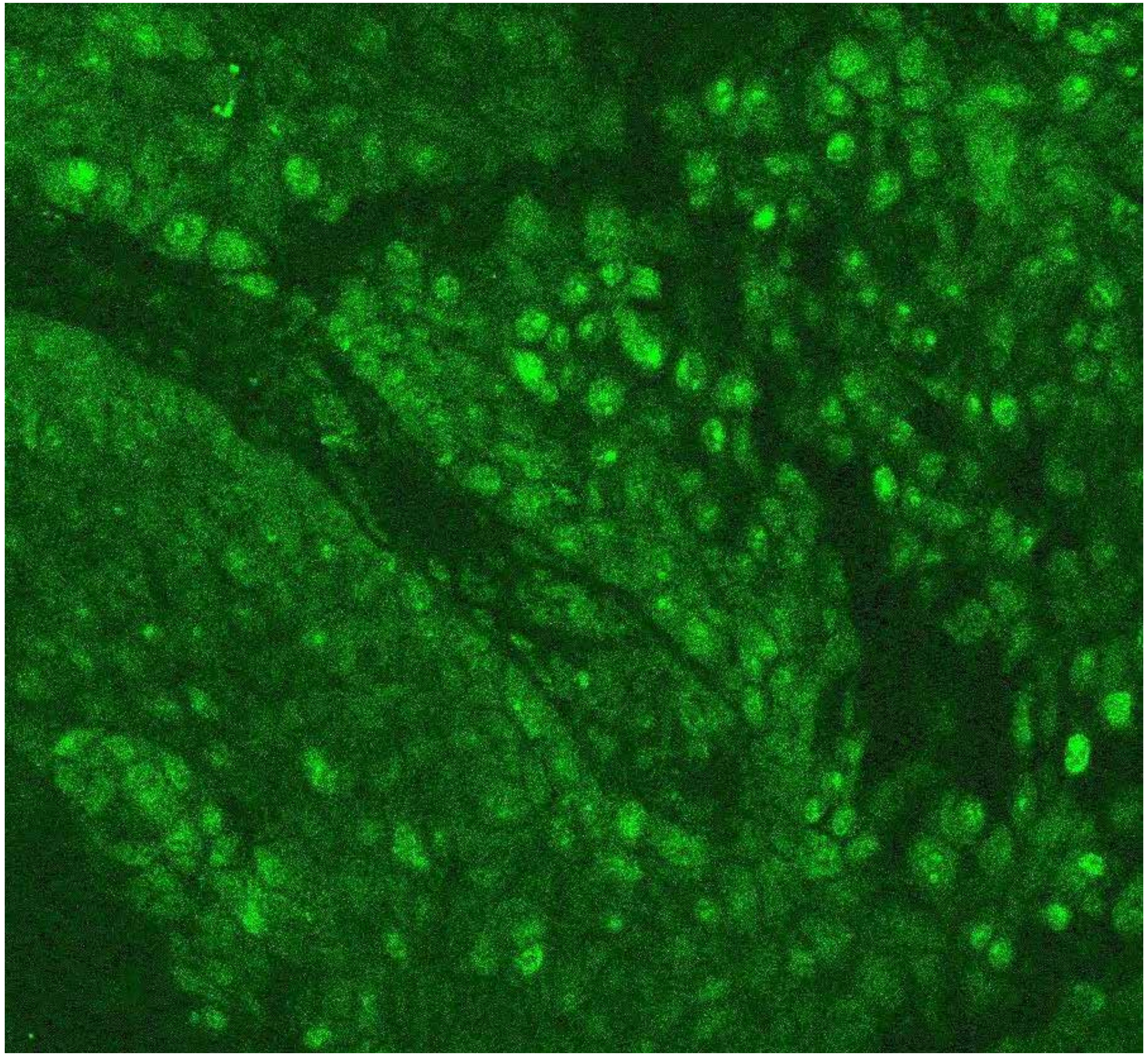

Not available

**PARPi-FL only (green)**

**PARPi-FL (green) + Autofluorescence (red/yellow)**

2627

### Close-up

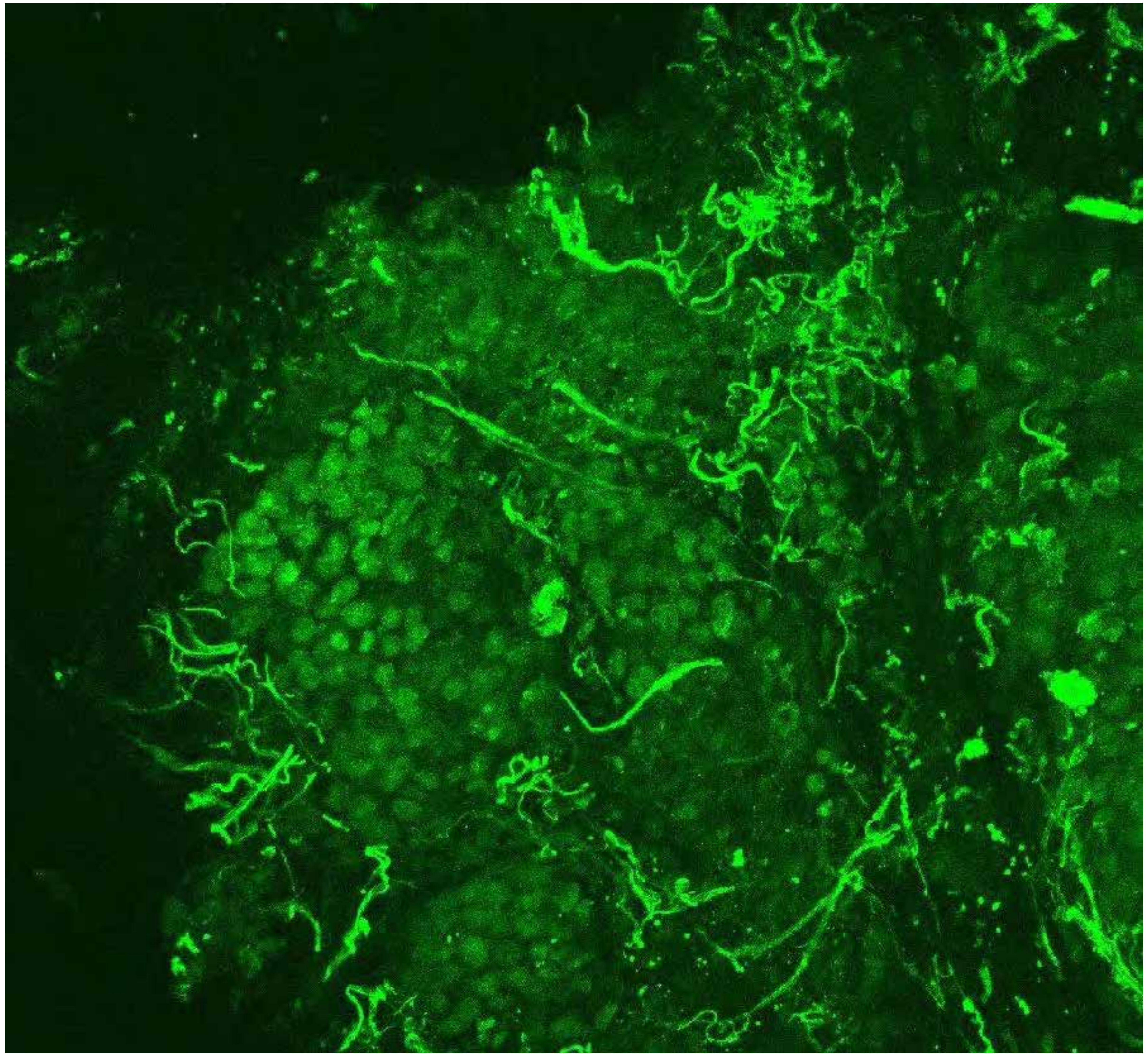

Not available

**PARPi-FL only (green)**

**PARPi-FL (green) + Autofluorescence (red/yellow)**

2627

### Close-up

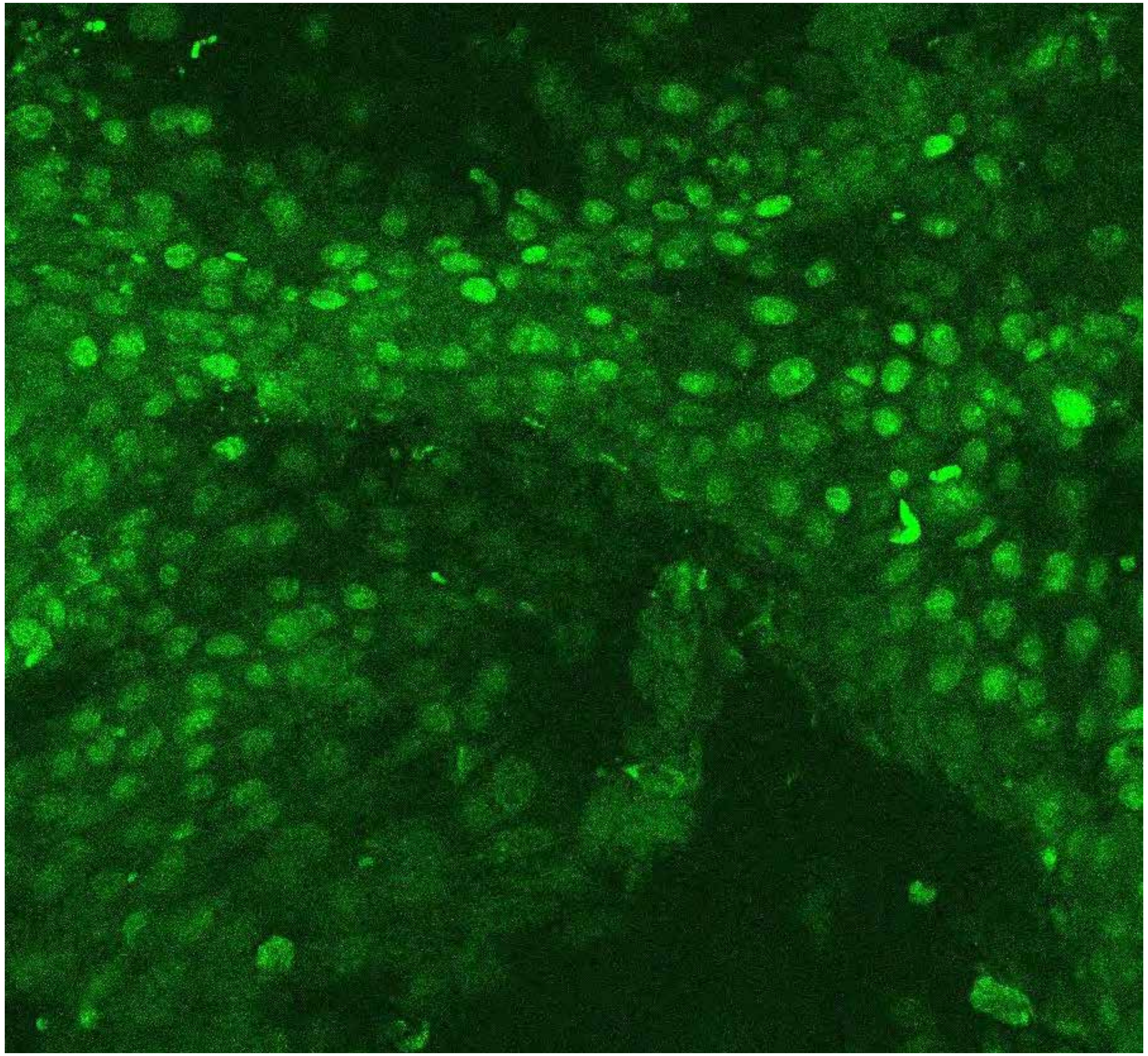

PARPi-FL only (green)

Not available

PARPi-FL (green) + Autofluorescence (red/yellow)

2627

10

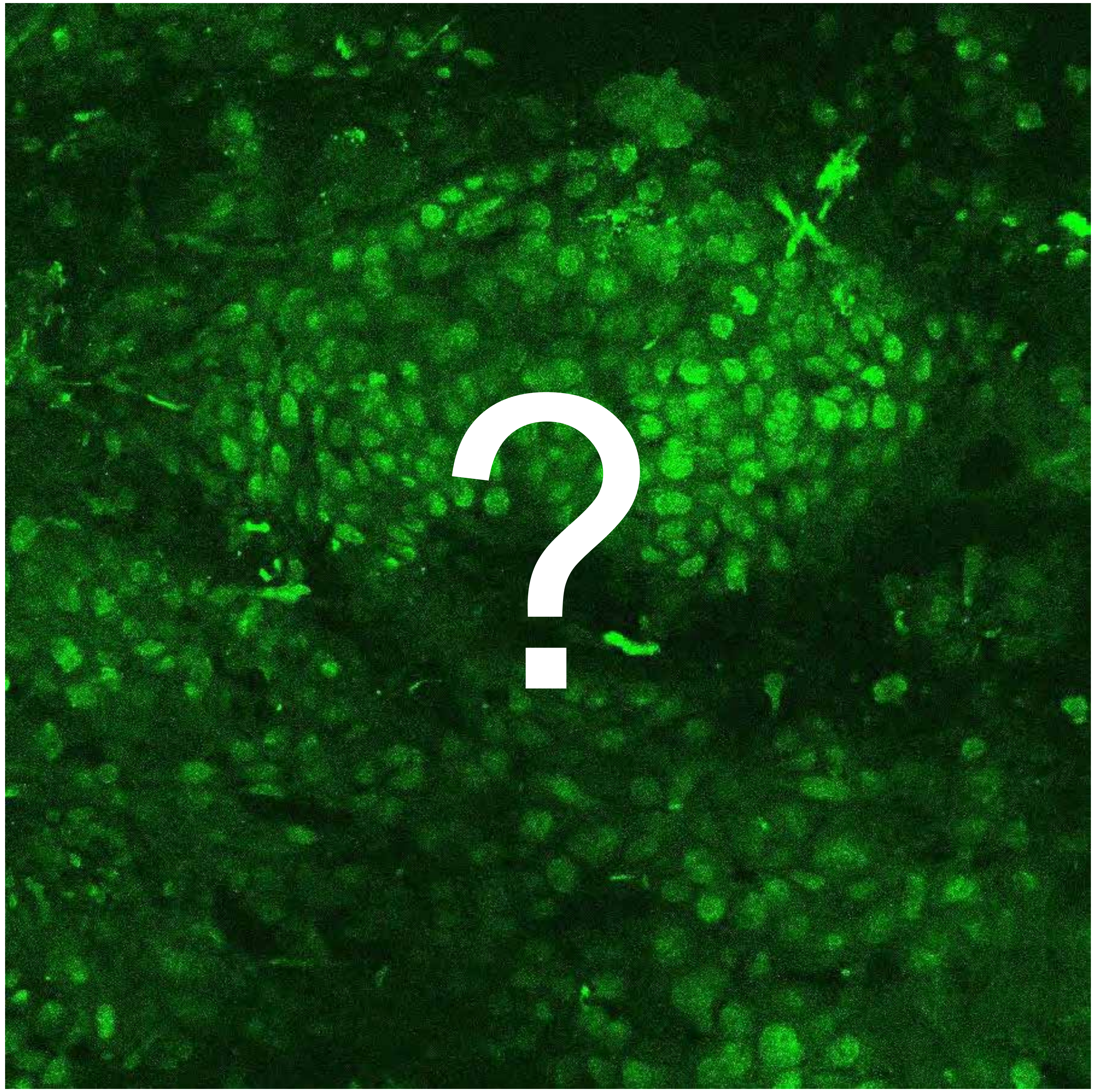


Tumor

Margin

2665

### Overview

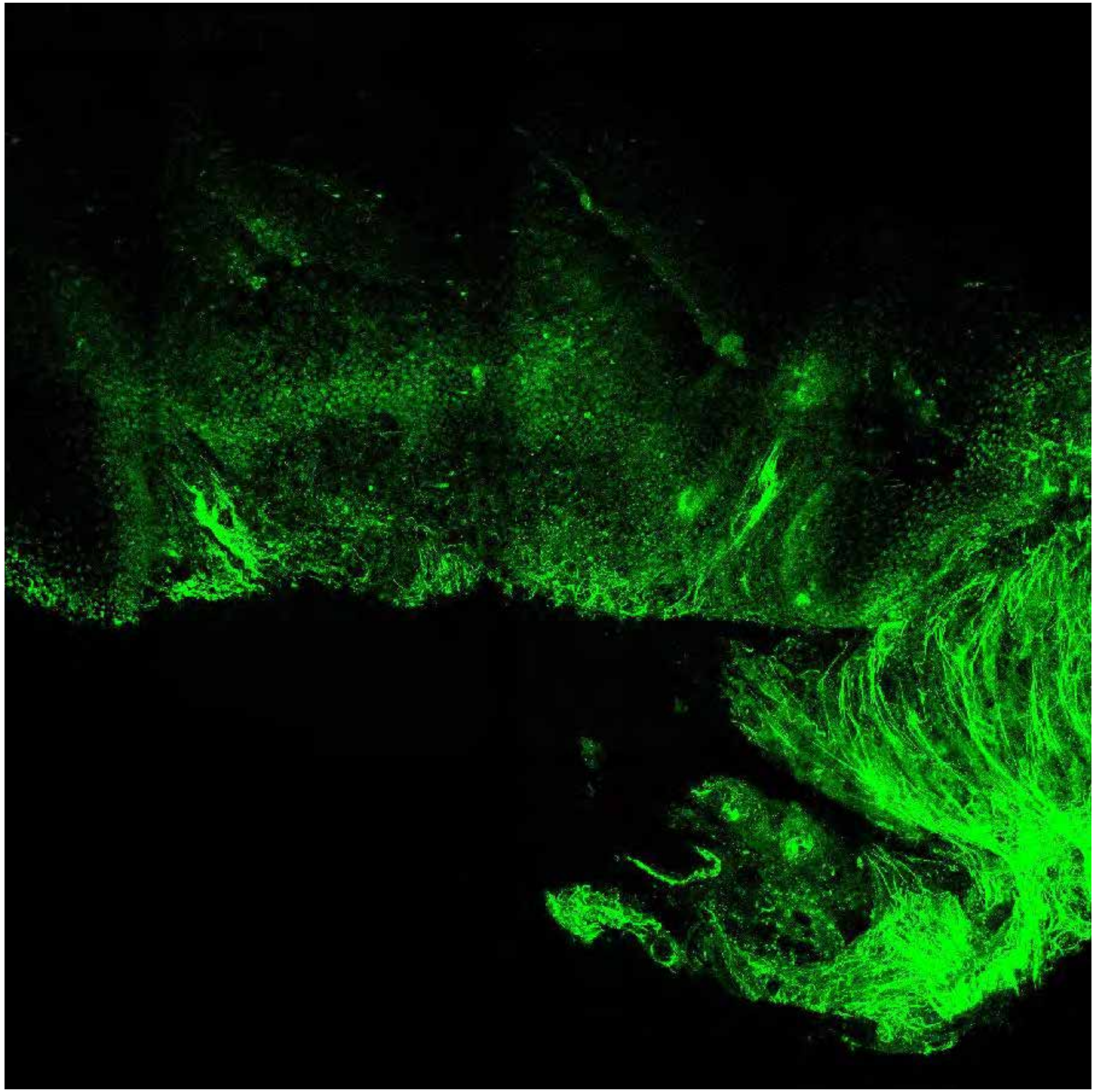

PARPi-FL only (green)

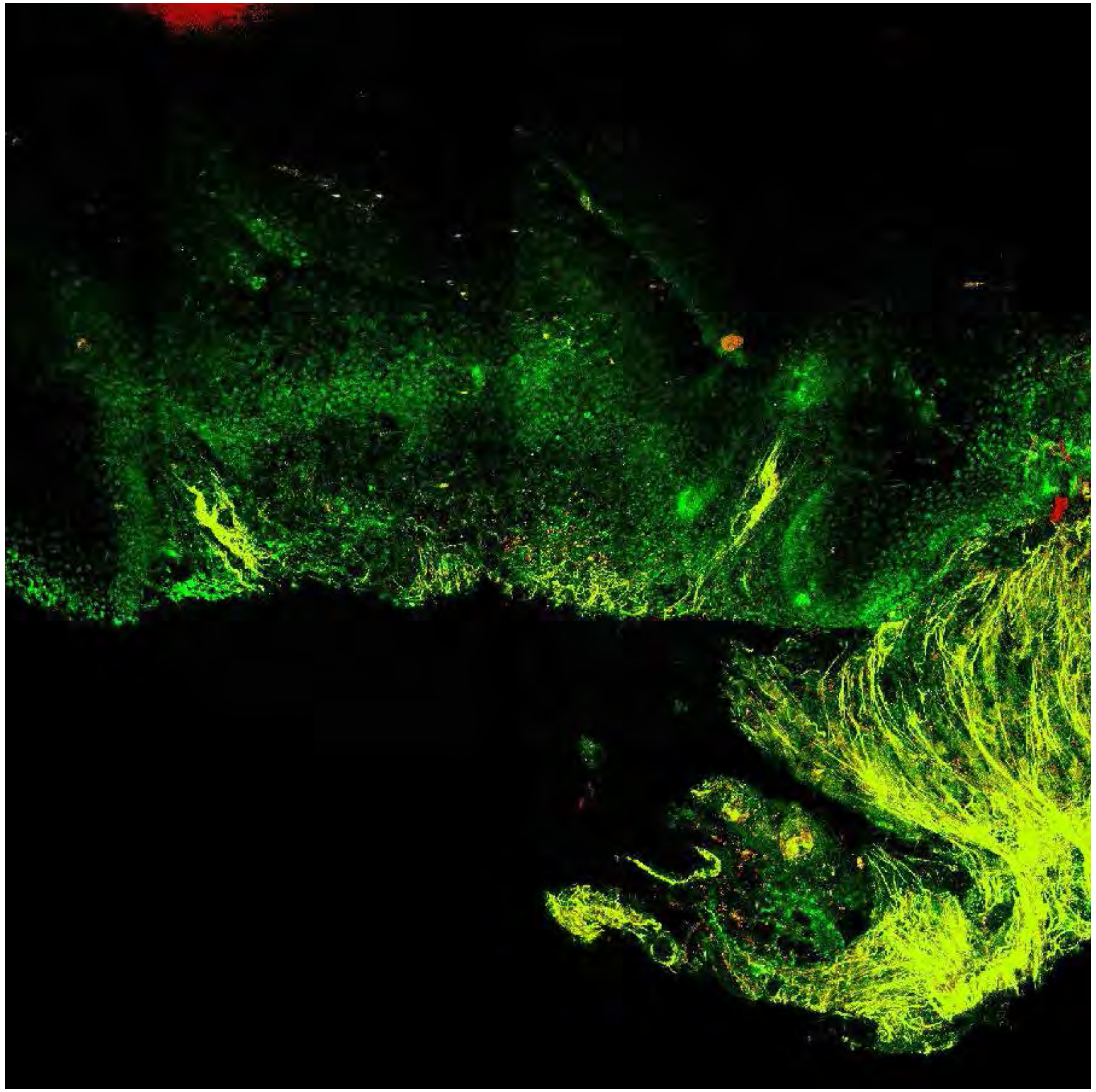

PARPi-FL (green) + Autofluorescence (red/yellow)

2665

### Close-up

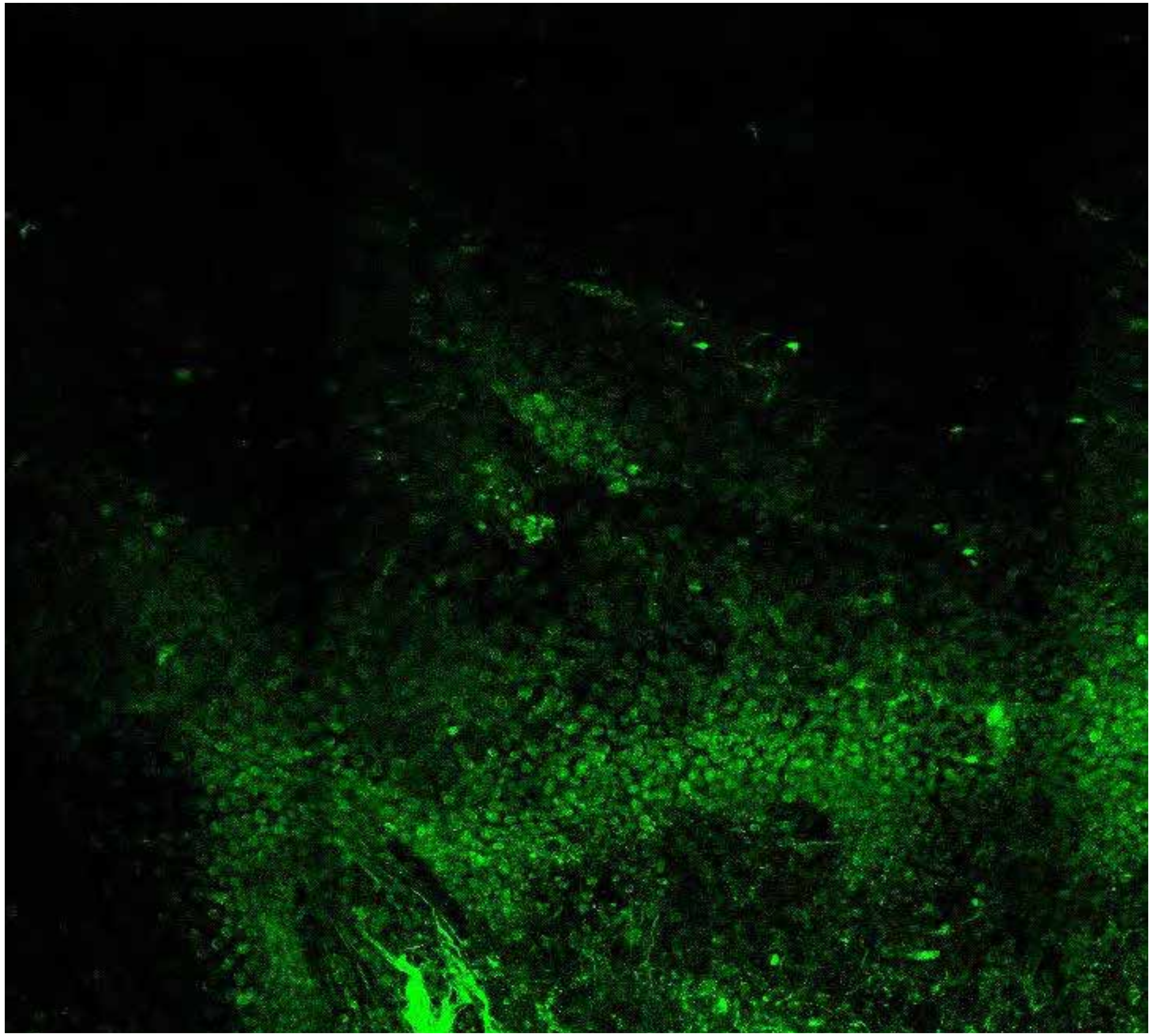

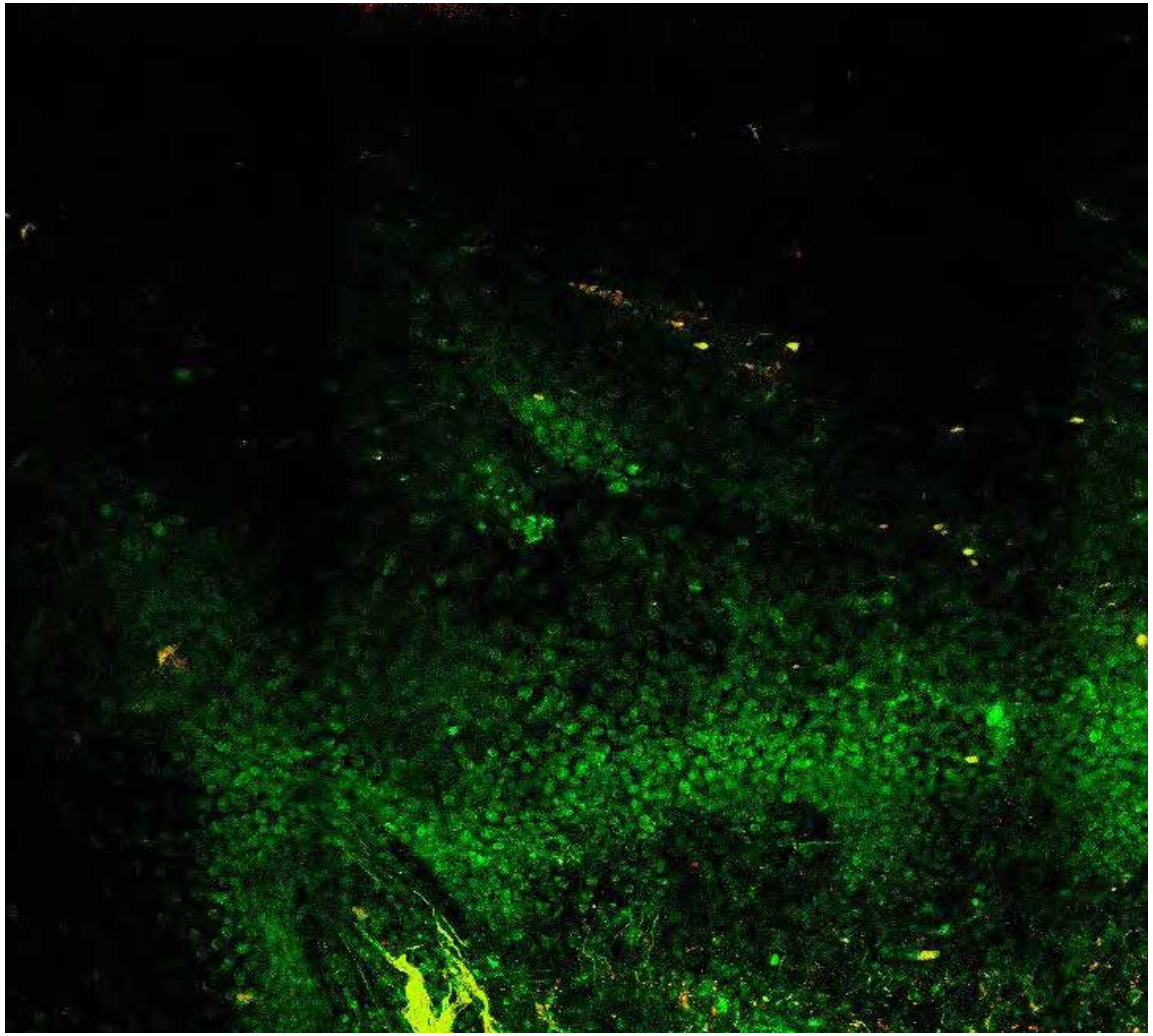

**PARPi-FL only (green)**

**PARPi-FL (green) + Autofluorescence (red/yellow)**

2665

### Close-up

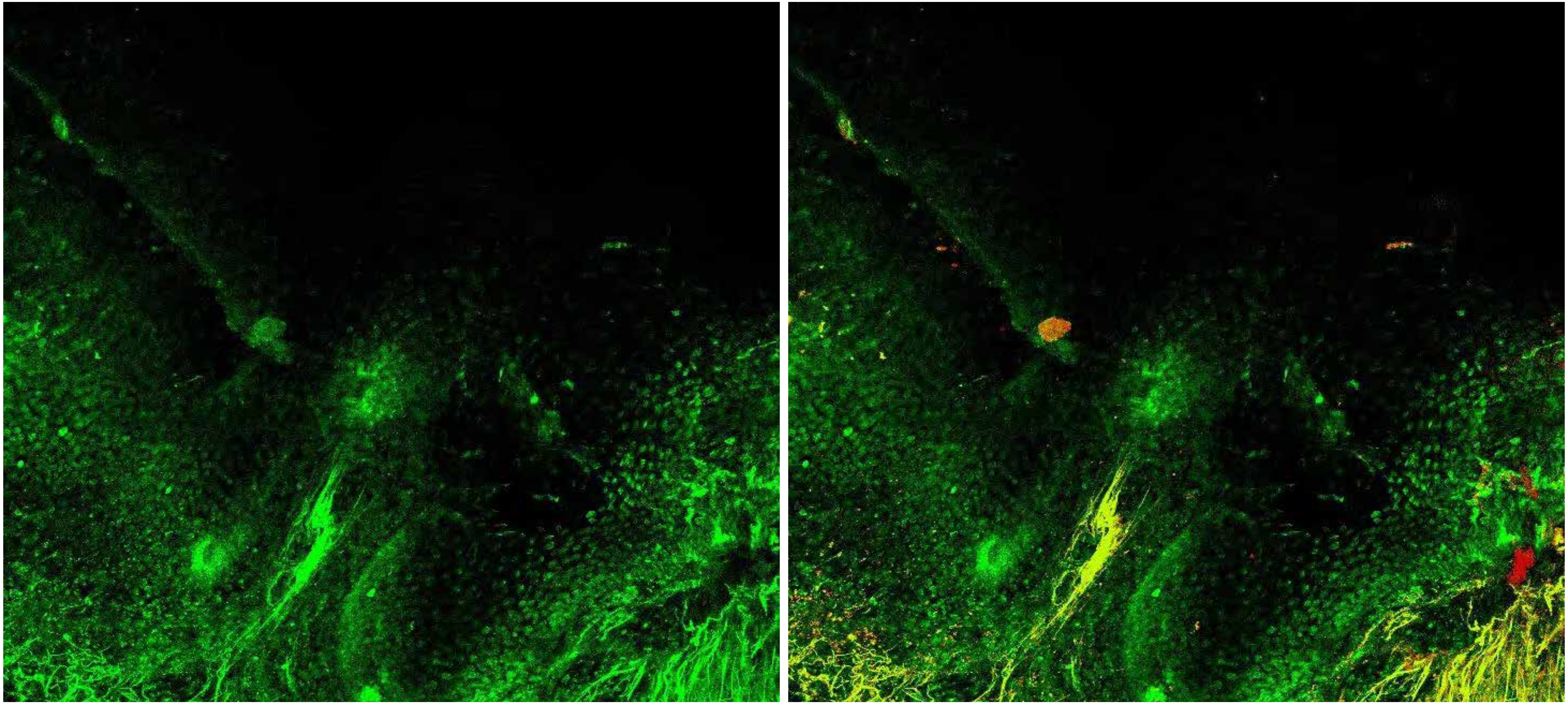

PARPi-FL only (green)

PARPi-FL (green) + Autofluorescence (red/yellow)

# 2665

### Close-up

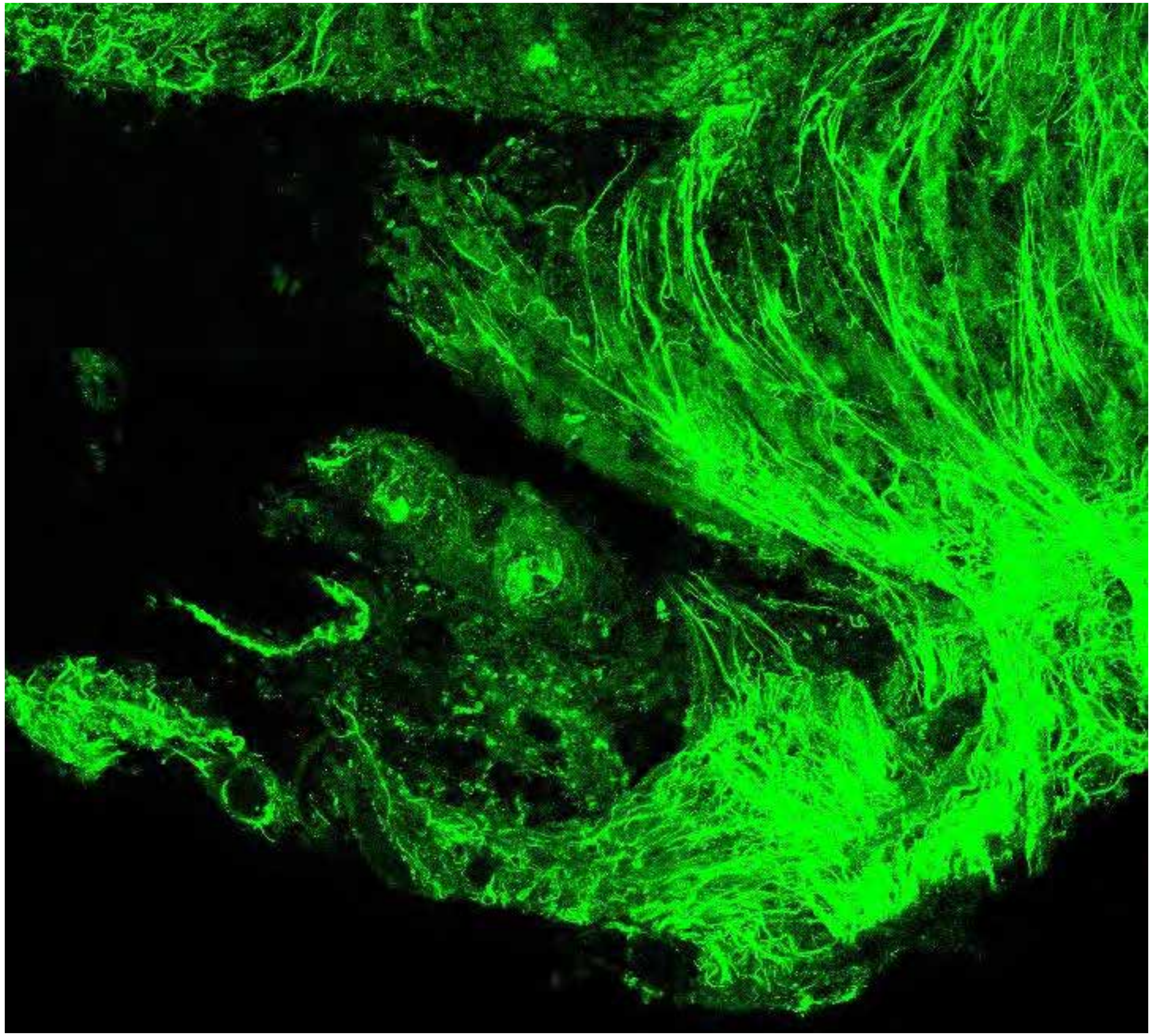

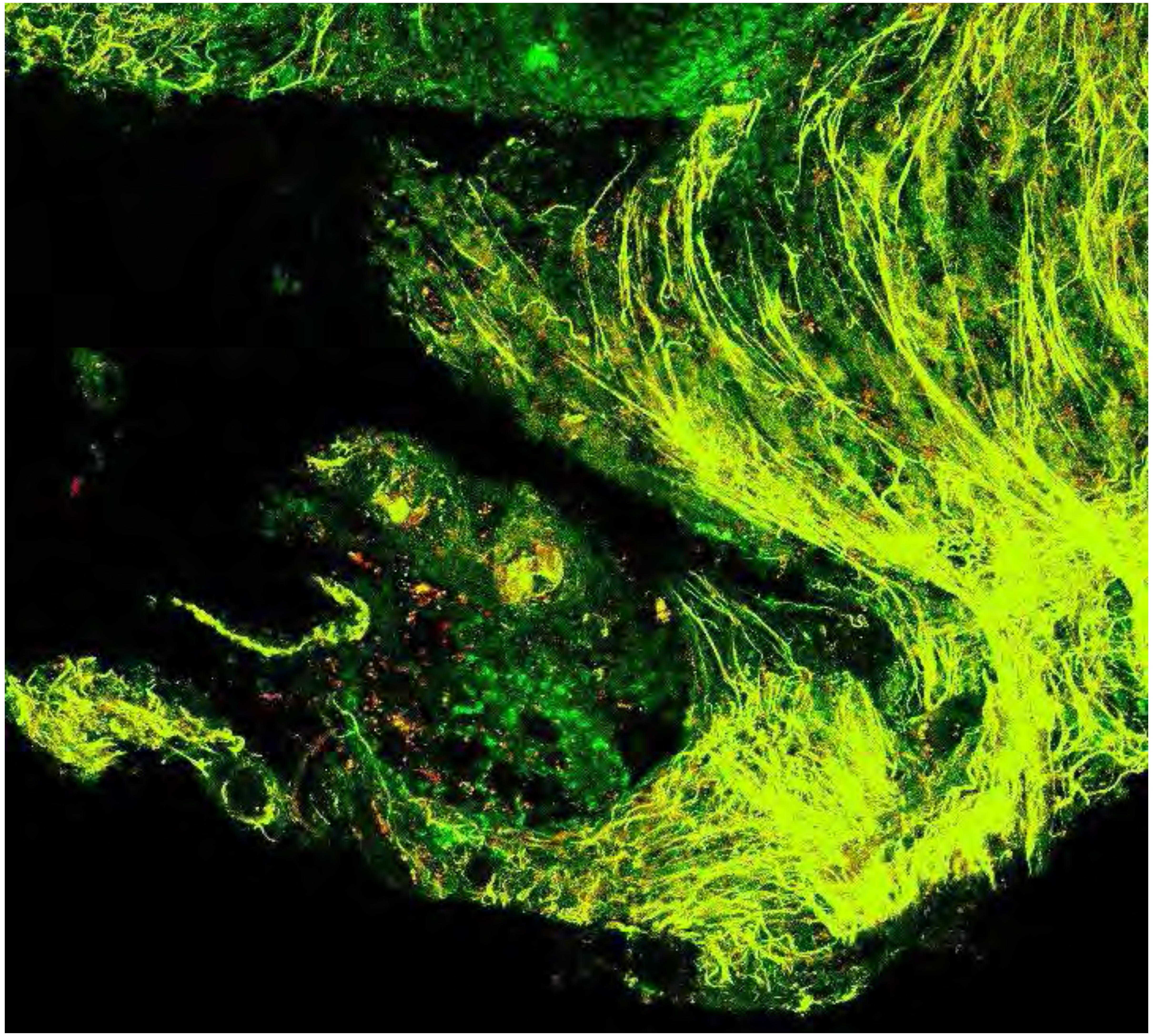

**PARPi-FL only (green)**

**PARPi-FL (green) + Autofluorescence (red/yellow)**

2665

10

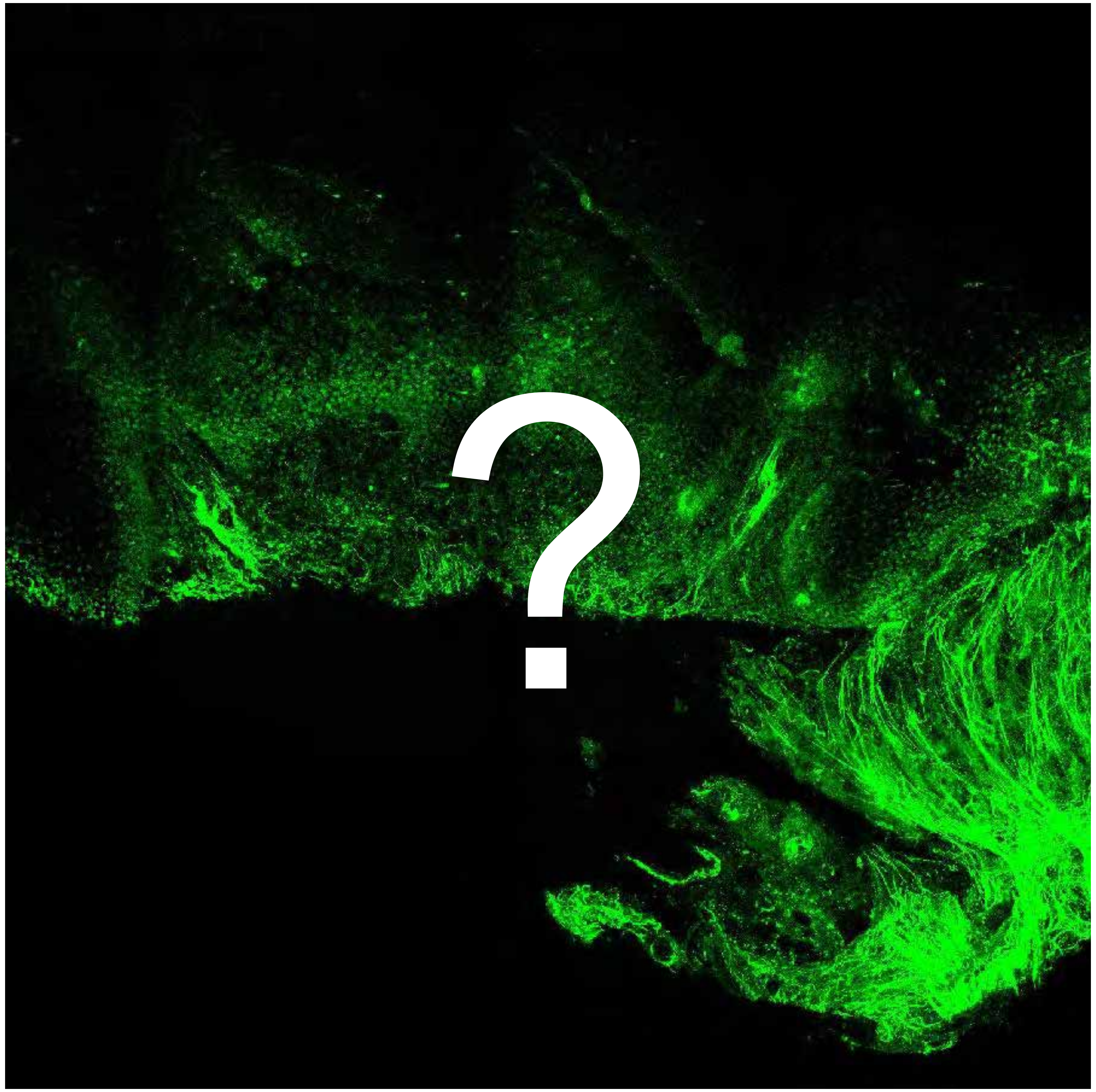


Tumor

Margin

3296

### Overview

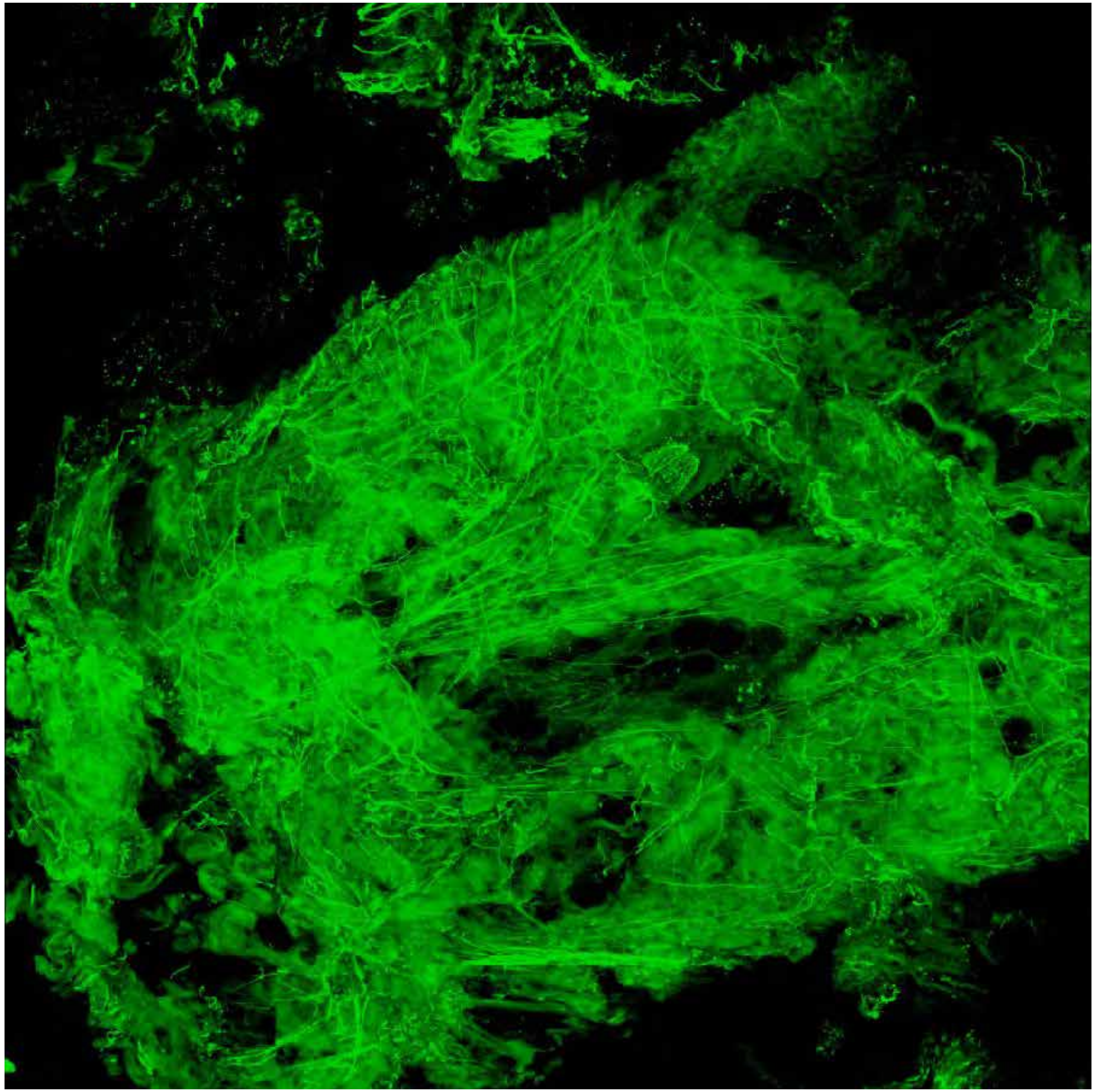

PARPi-FL only (green)

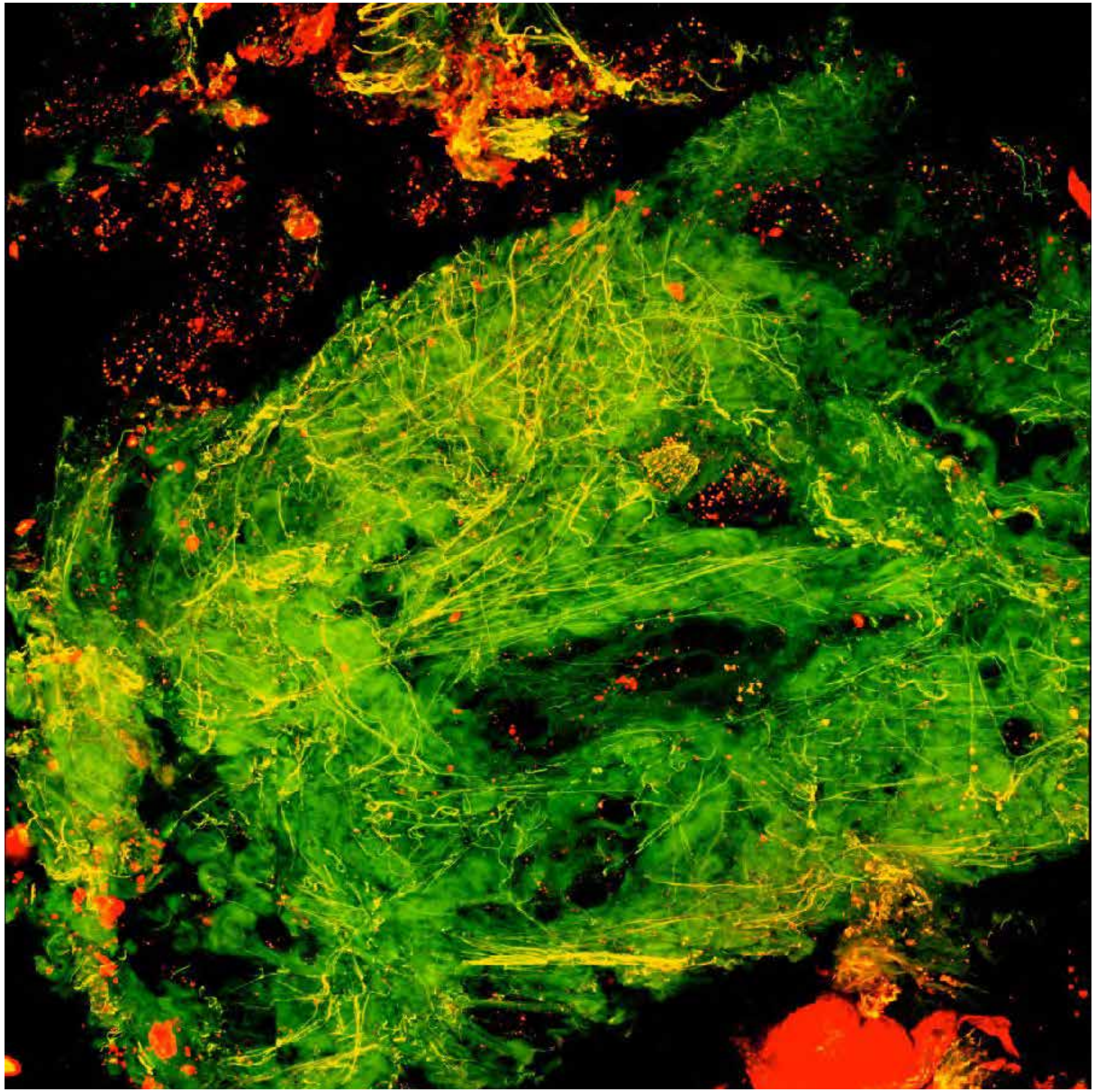

PARPi-FL (green) + Autofluorescence (red/yellow)

3296

### Close-up

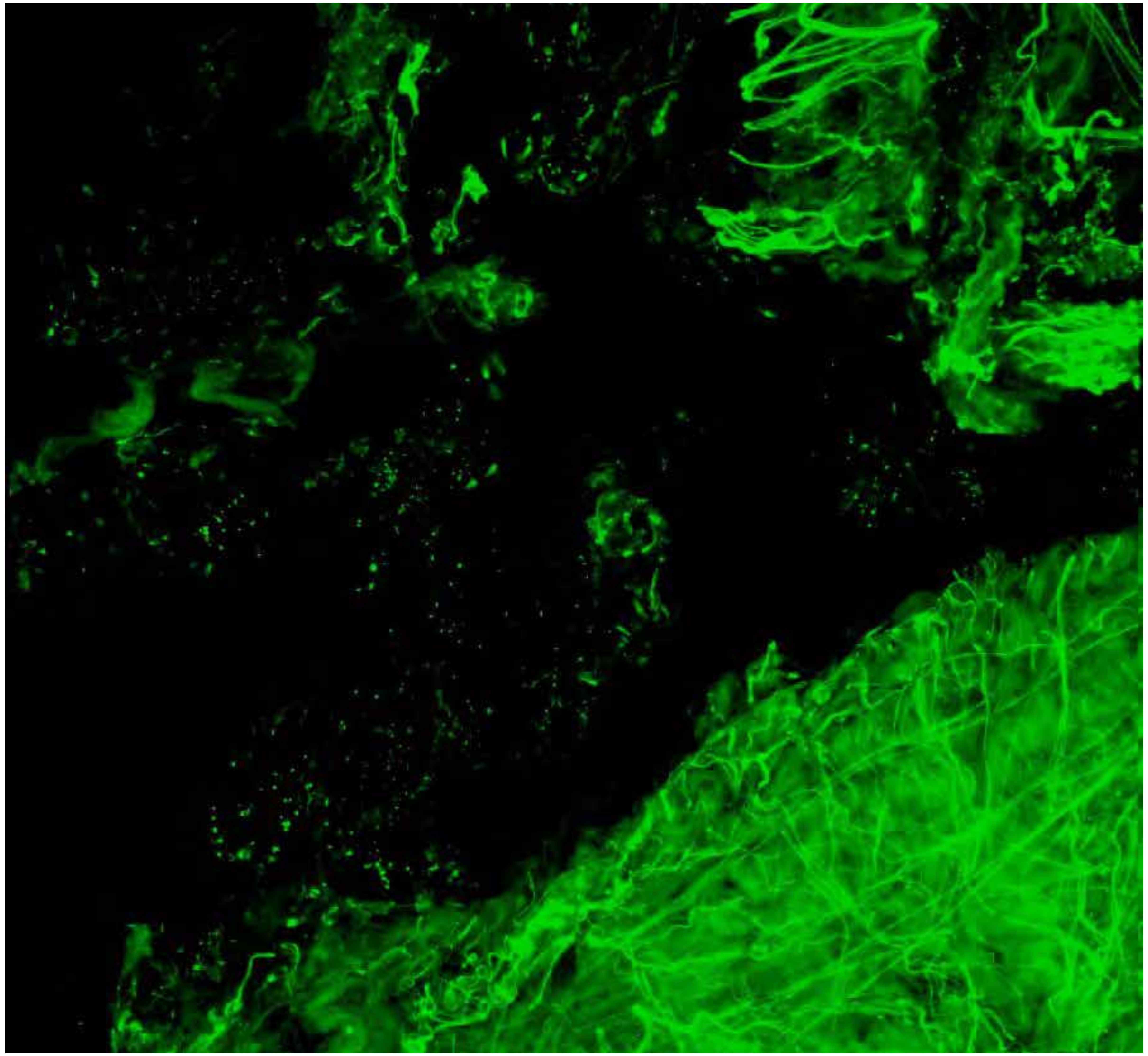

PARPi-FL only (green)

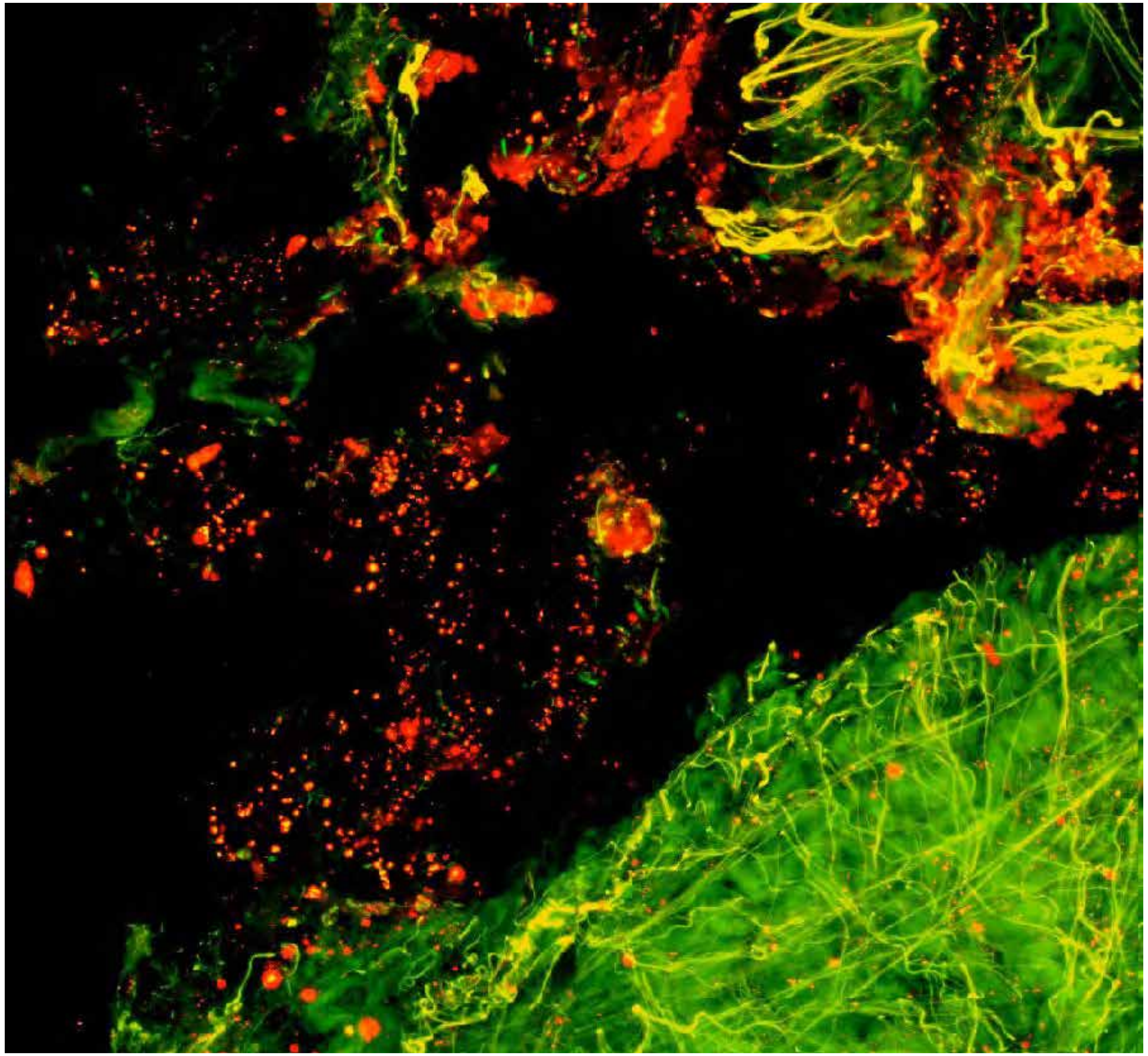

PARPi-FL (green) + Autofluorescence (red/yellow)

### Close-up

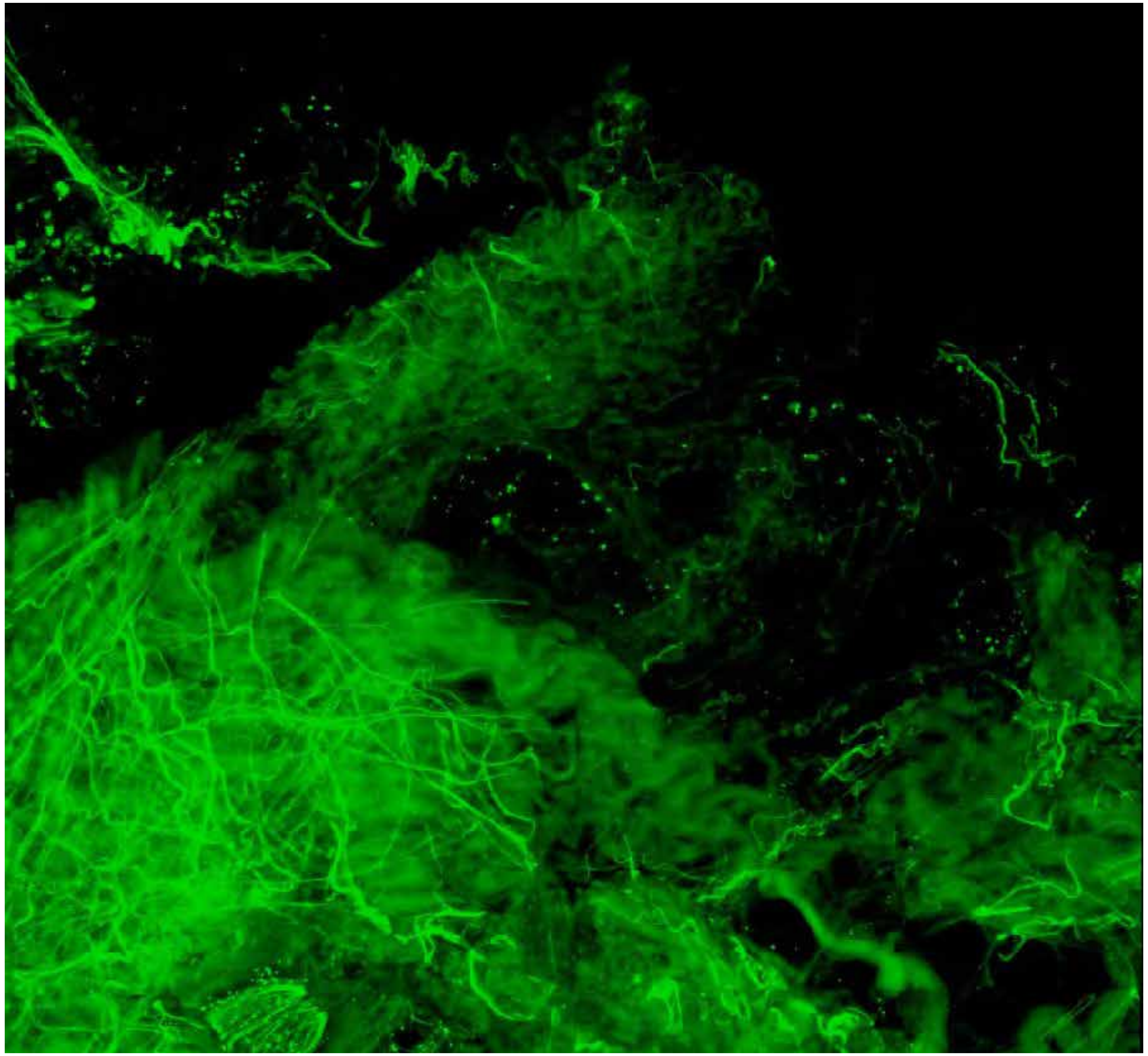

PARPi-FL only (green)

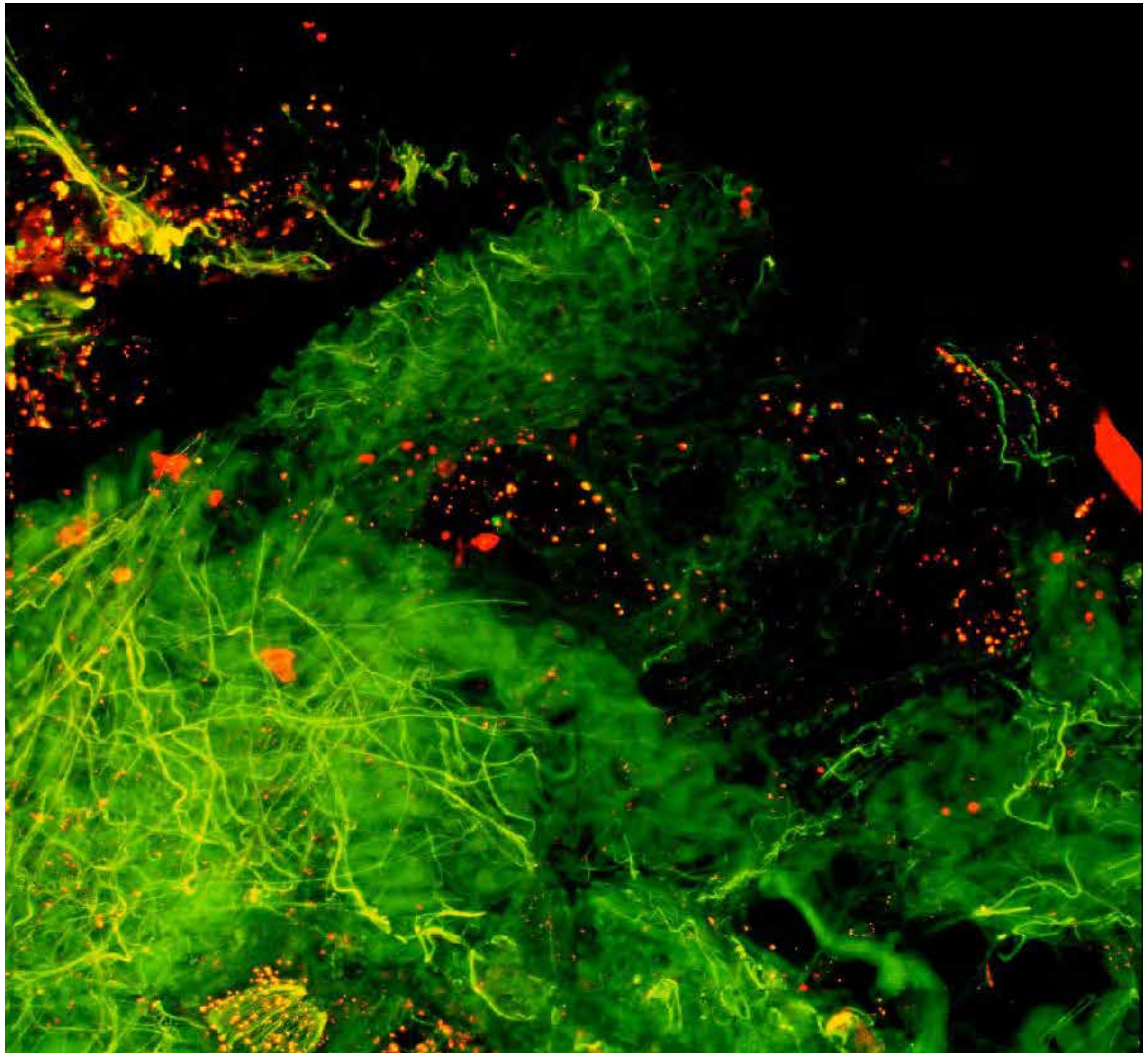

PARPi-FL (green) + Autofluorescence (red/yellow)

3296

### Close-up

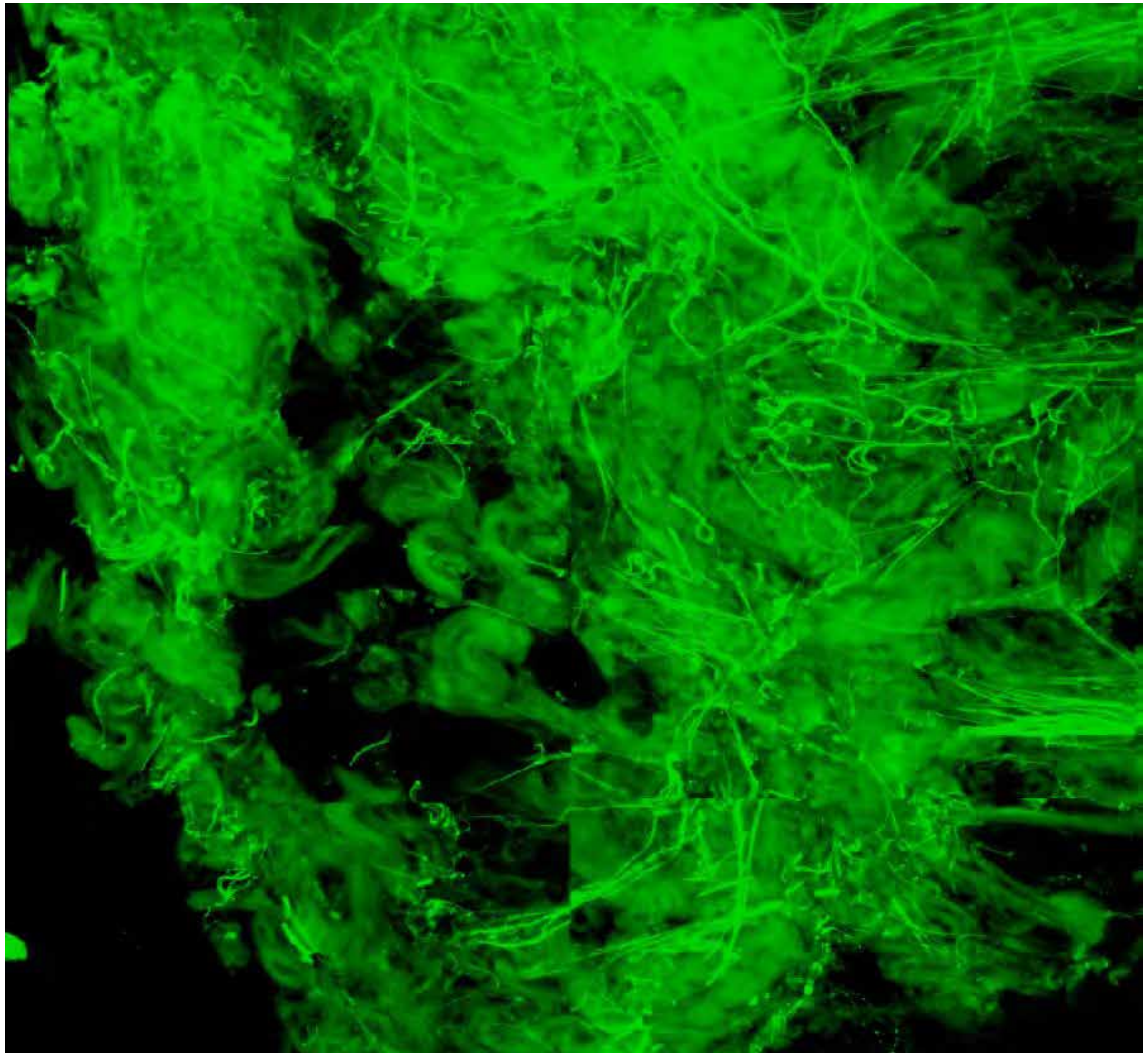

PARPi-FL only (green)

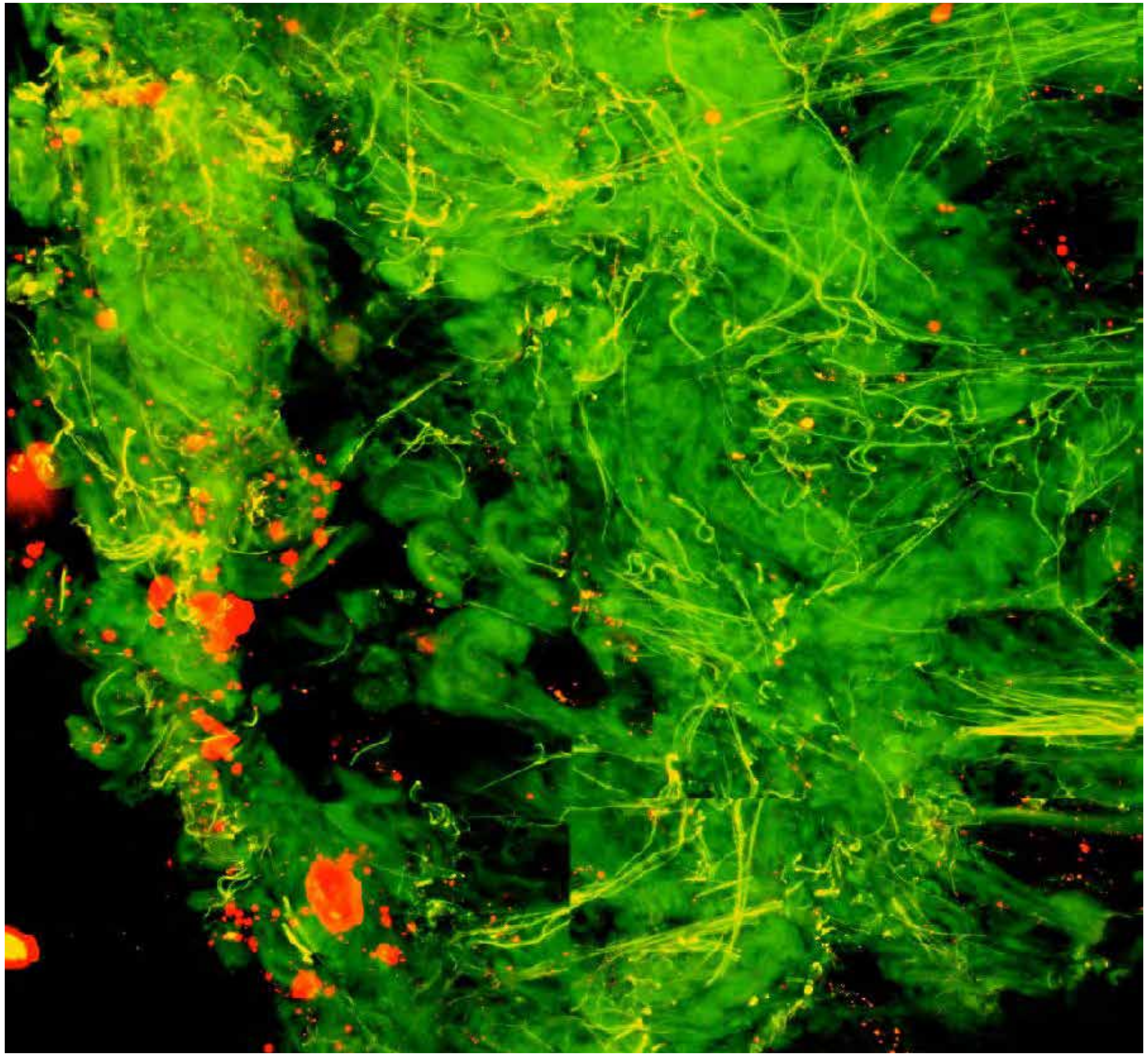

PARPi-FL (green) + Autofluorescence (red/yellow)

3296

### Close-up

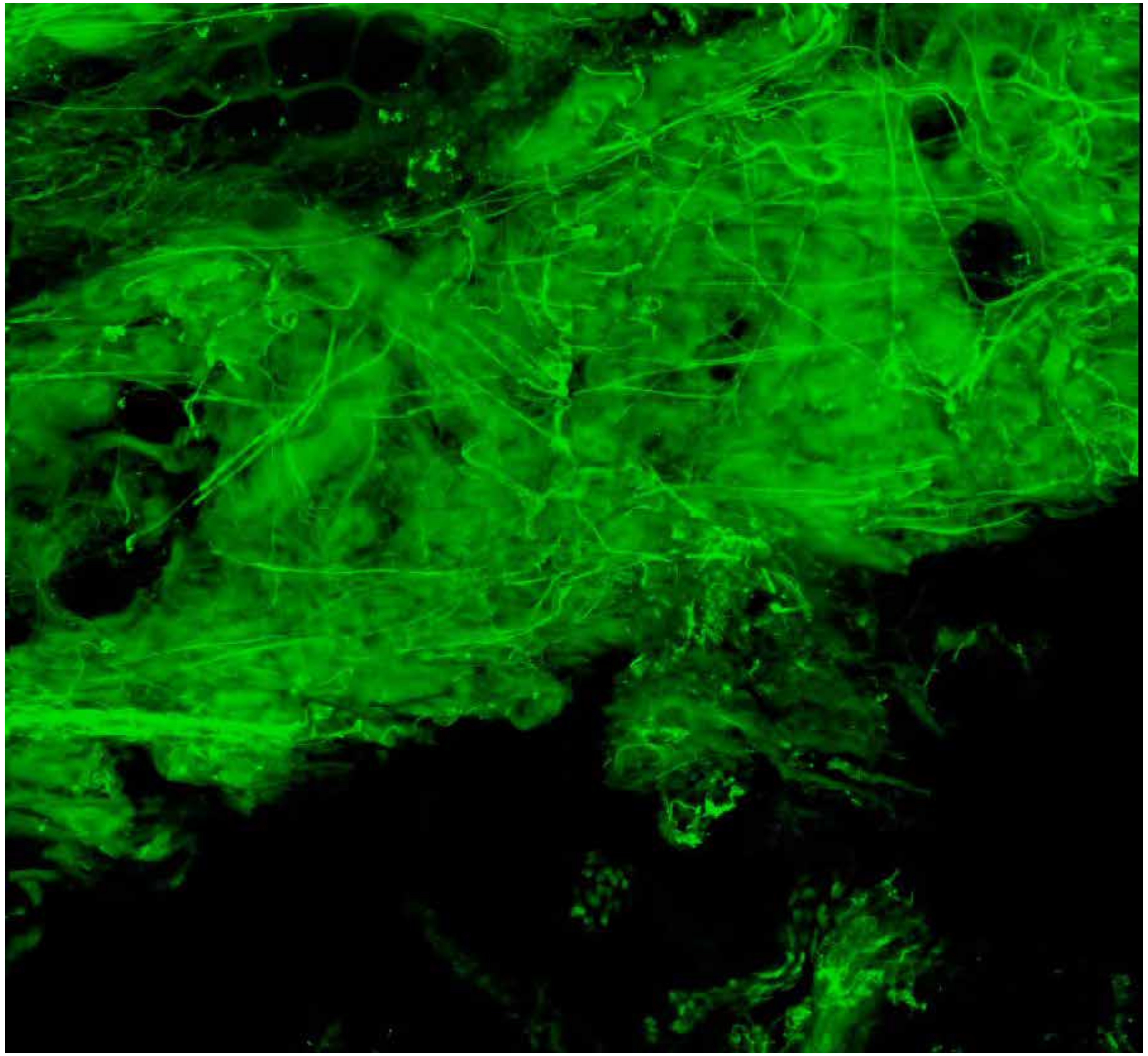

PARPi-FL only (green)

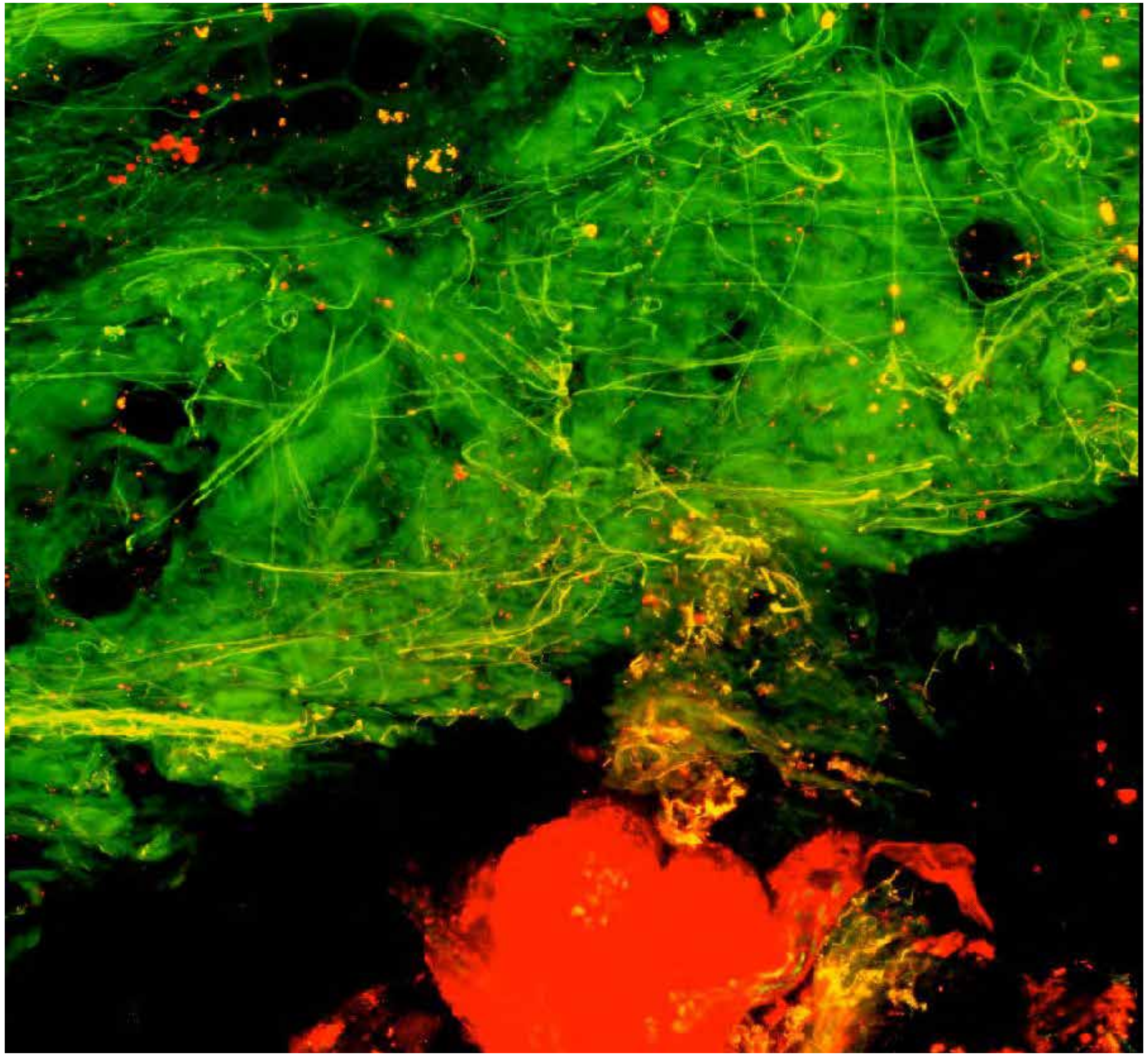

PARPi-FL (green) + Autofluorescence (red/yellow)

3296



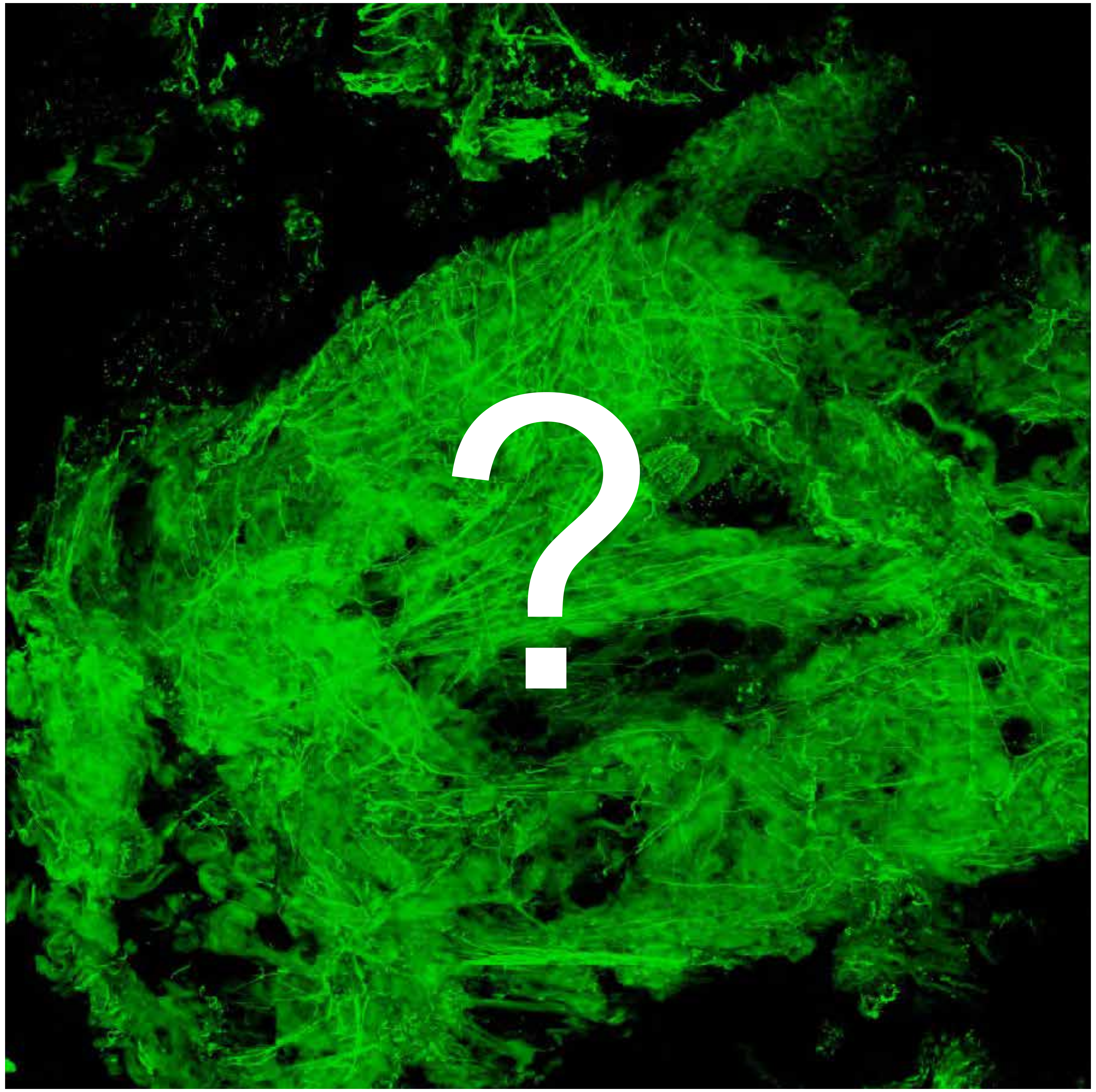


Tumor

Margin

3403

### Overview

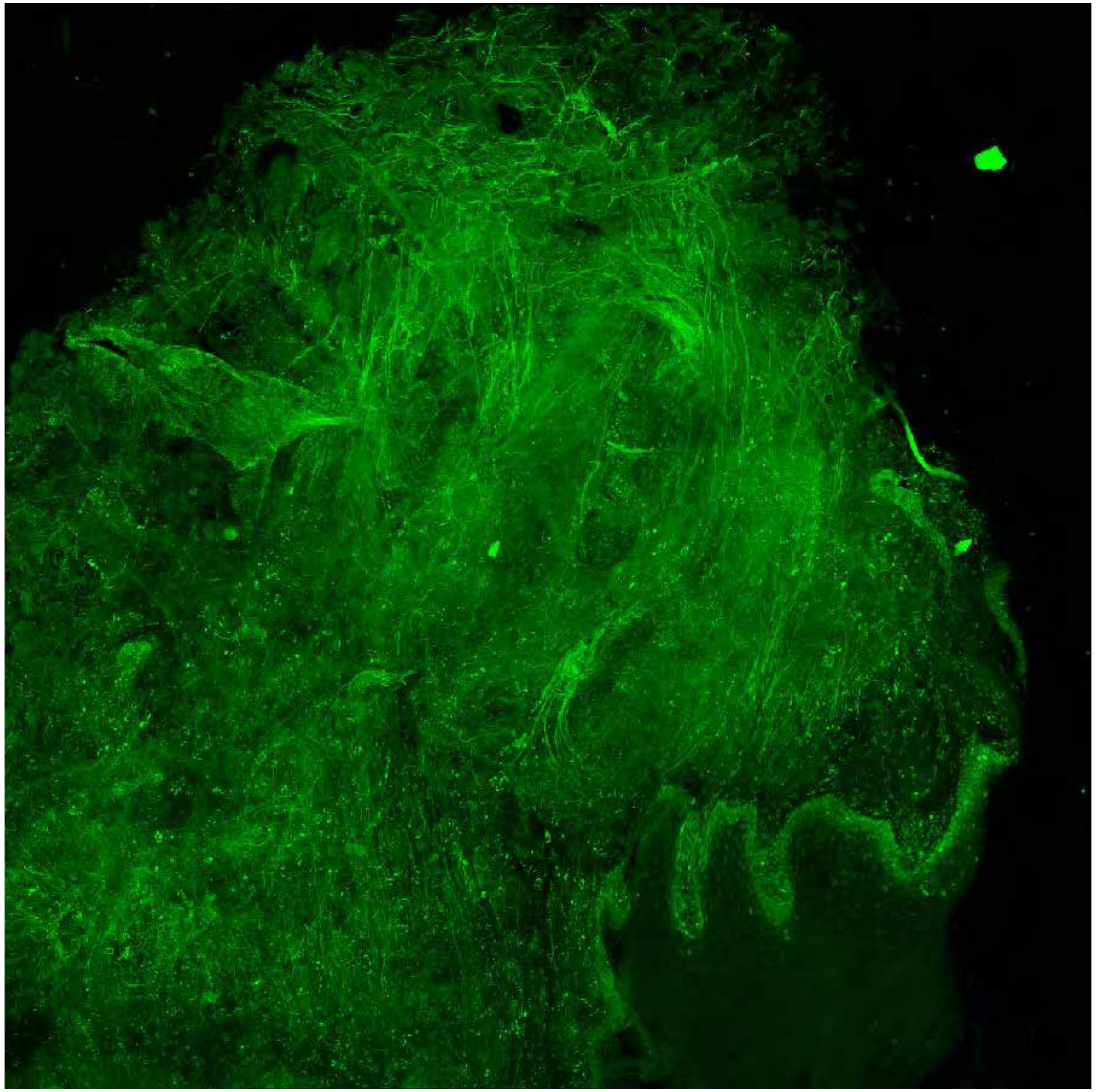

PARPi-FL only (green)

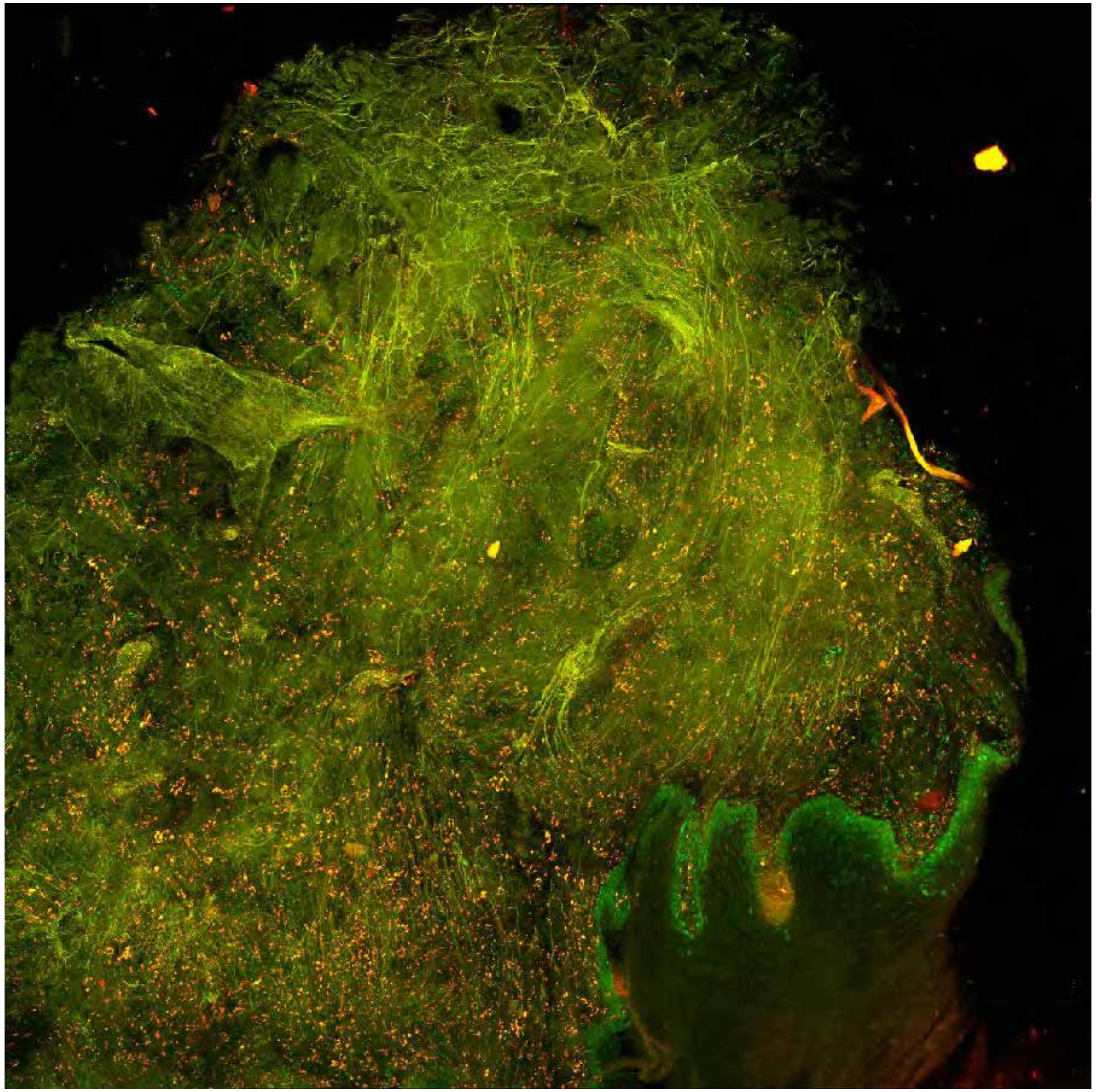

PARPi-FL (green) + Autofluorescence (red/yellow)

3403

### Close-up

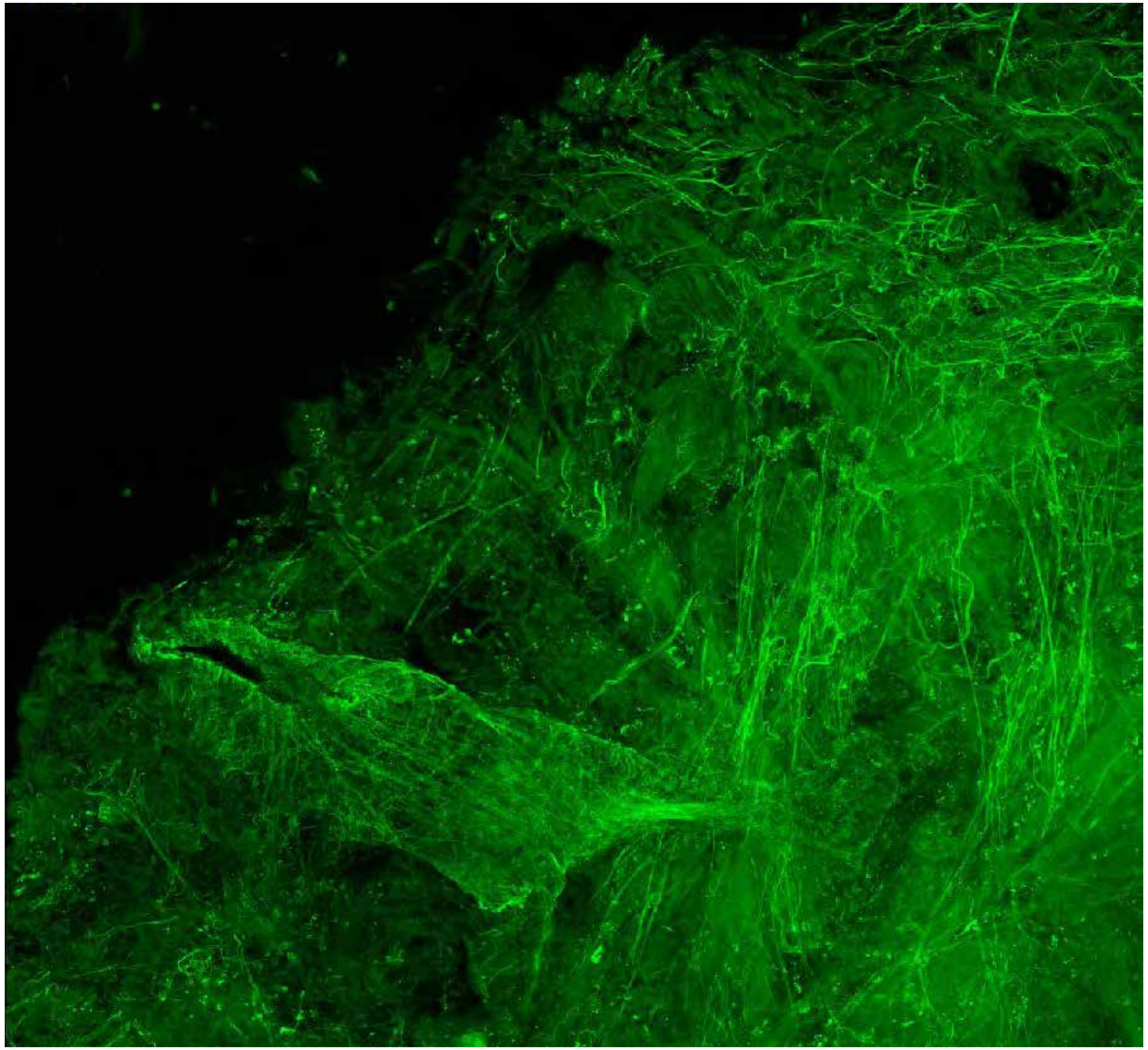

**PARPi-FL only (green)**

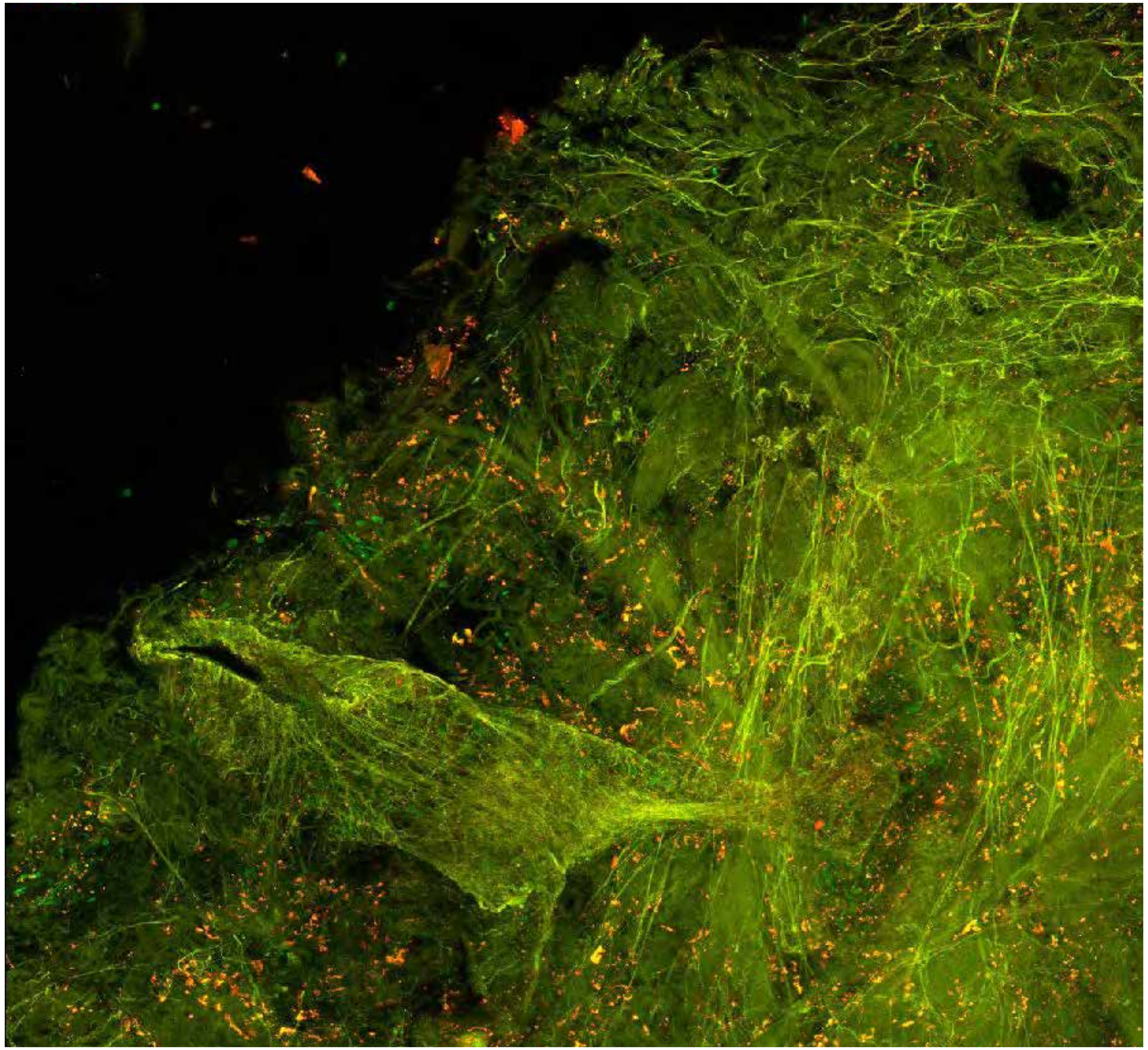

**PARPi-FL (green) + Autofluorescence (red/yellow)**

3403

### Close-up

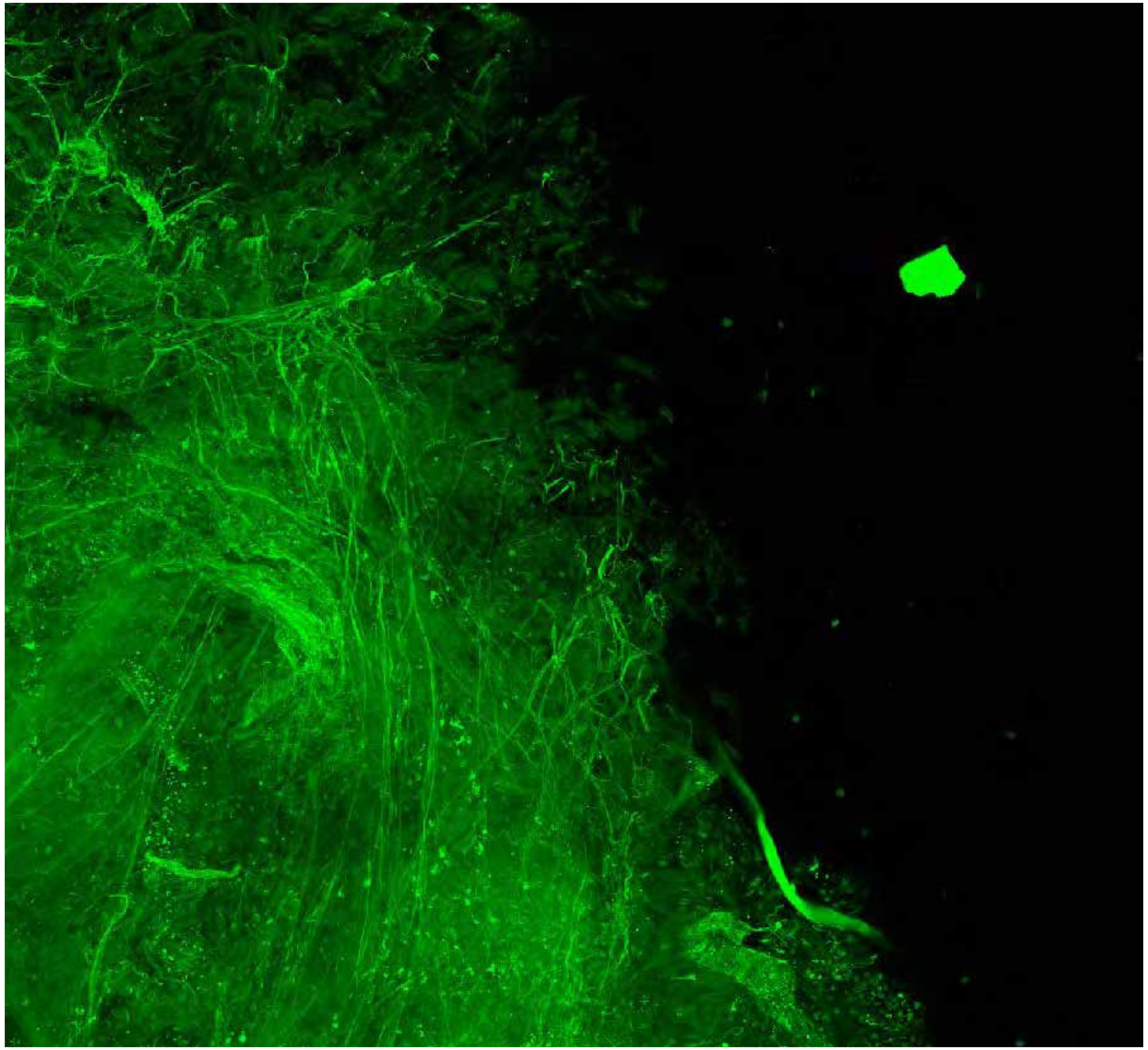

**PARPi-FL only (green)**

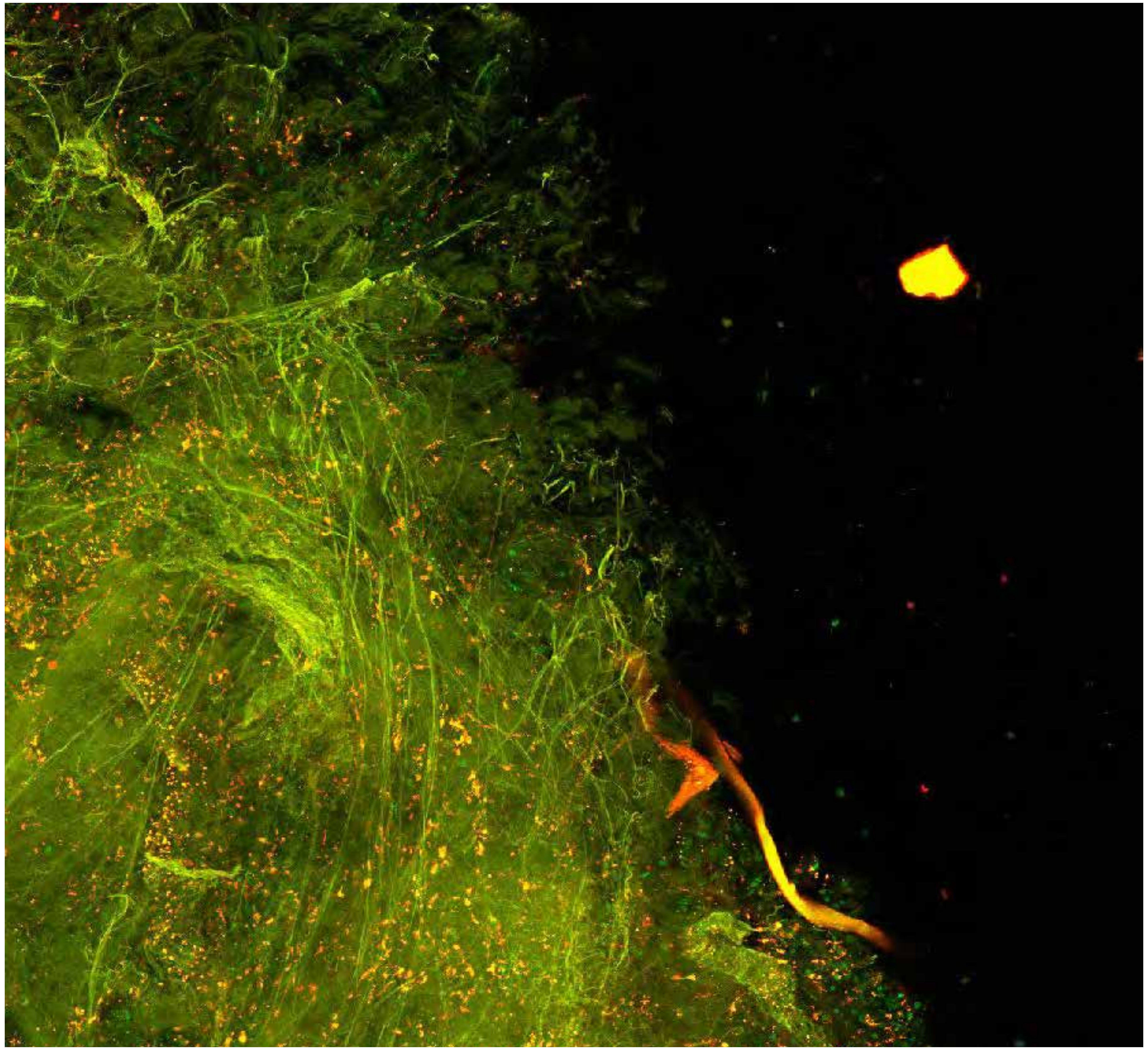

**PARPi-FL (green) + Autofluorescence (red/yellow)**

3403

### Close-up

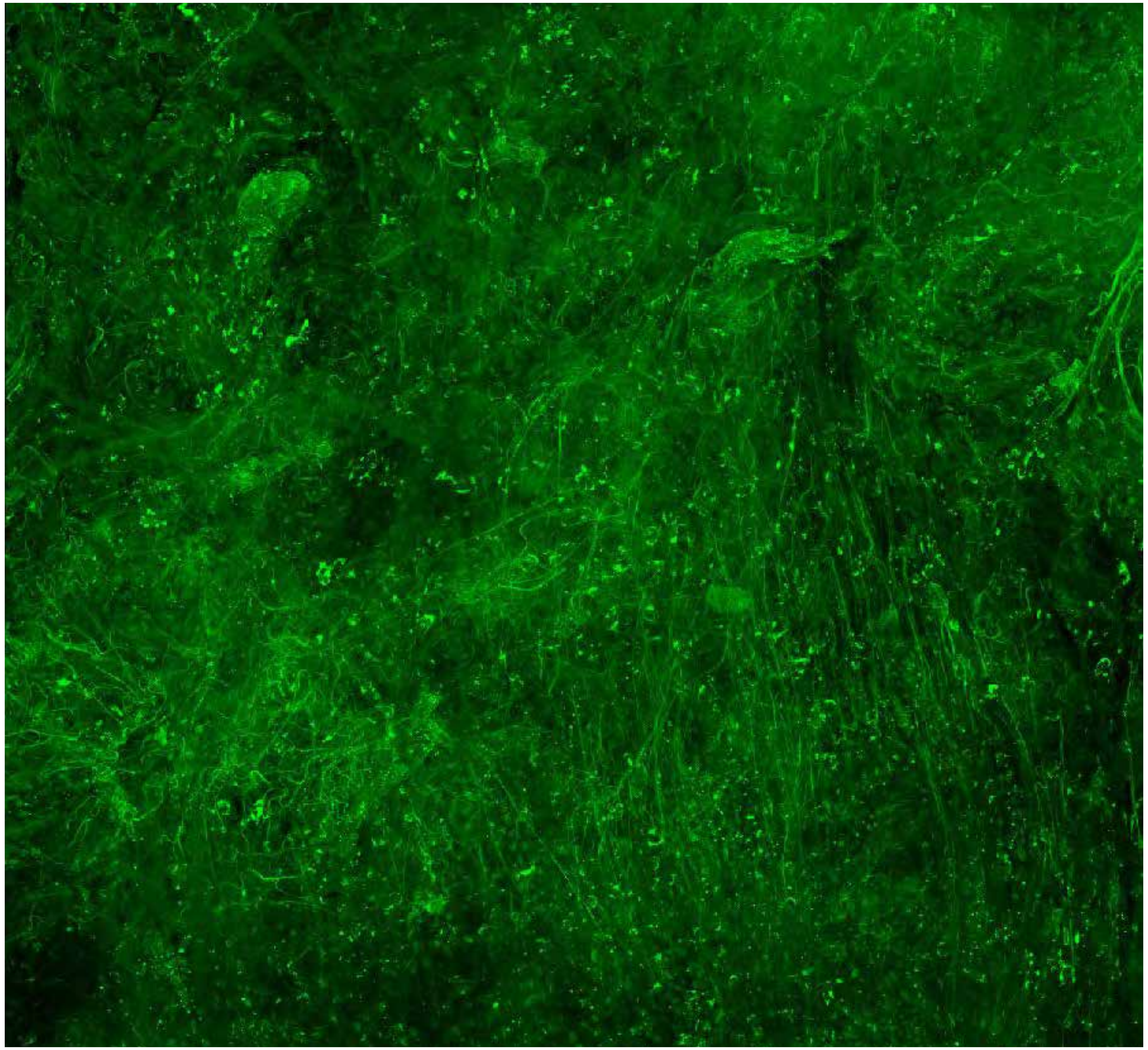

PARPi-FL only (green)

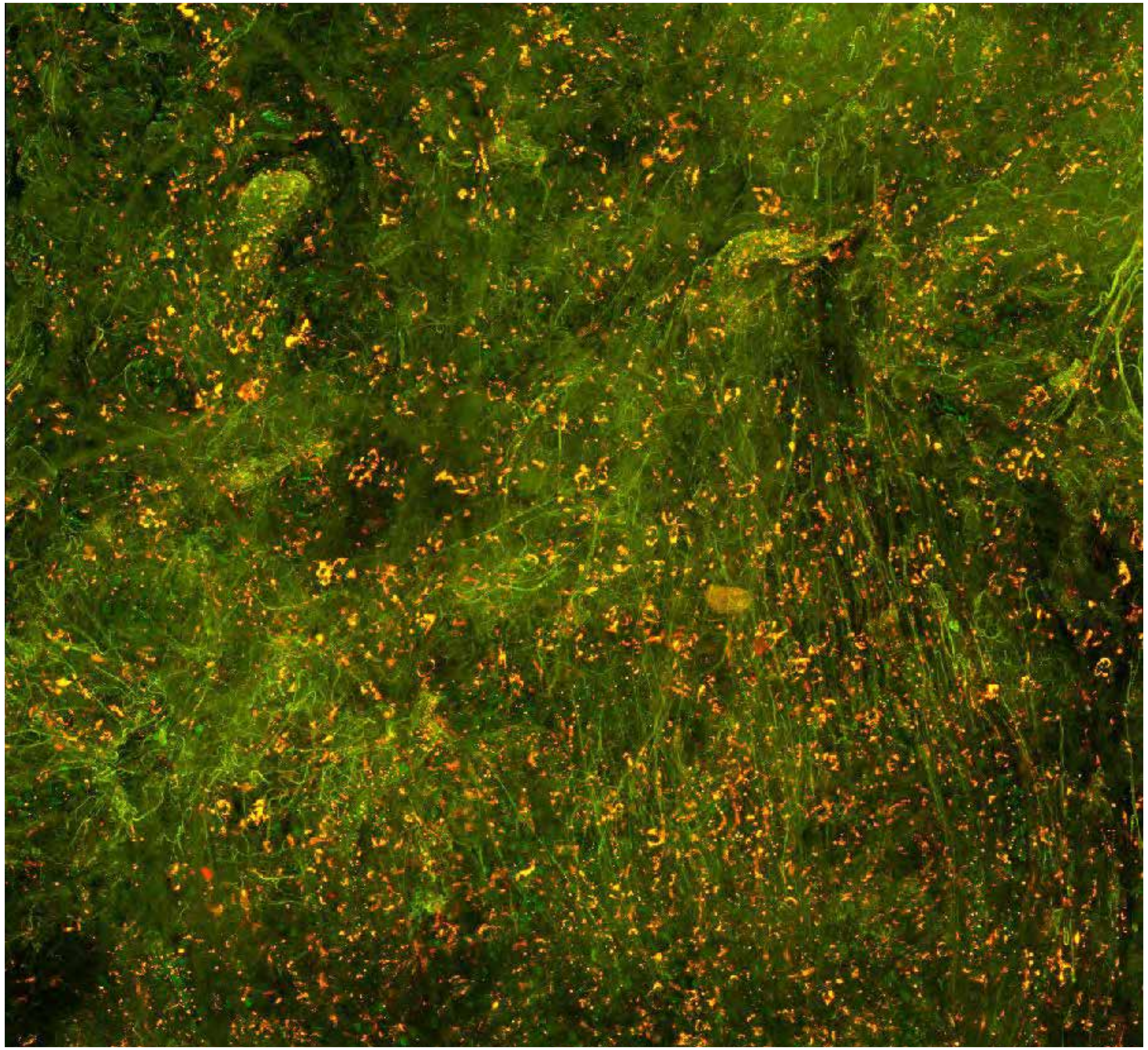

PARPi-FL (green) + Autofluorescence (red/yellow)

3403

### Close-up

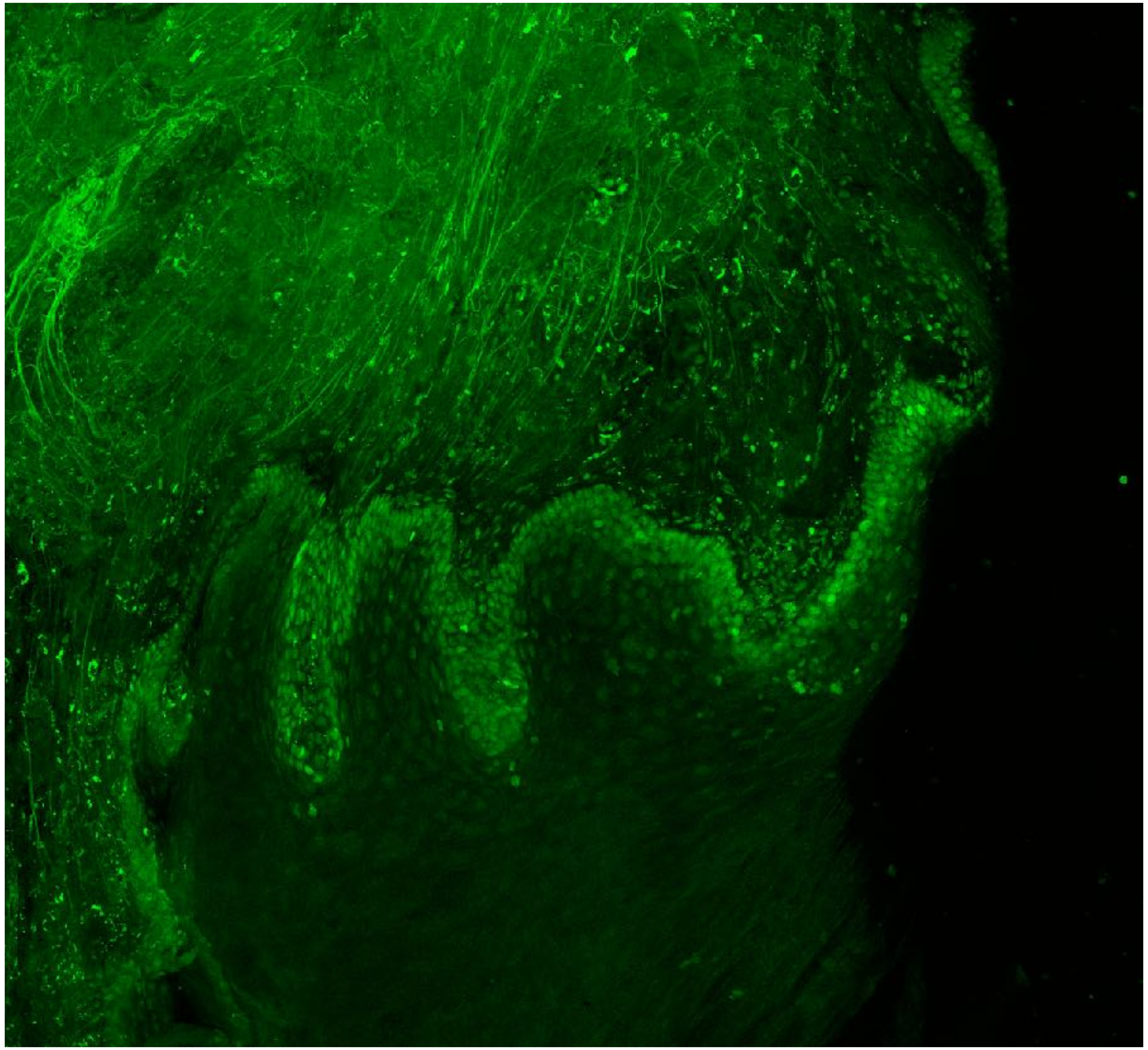

PARPi-FL only (green)

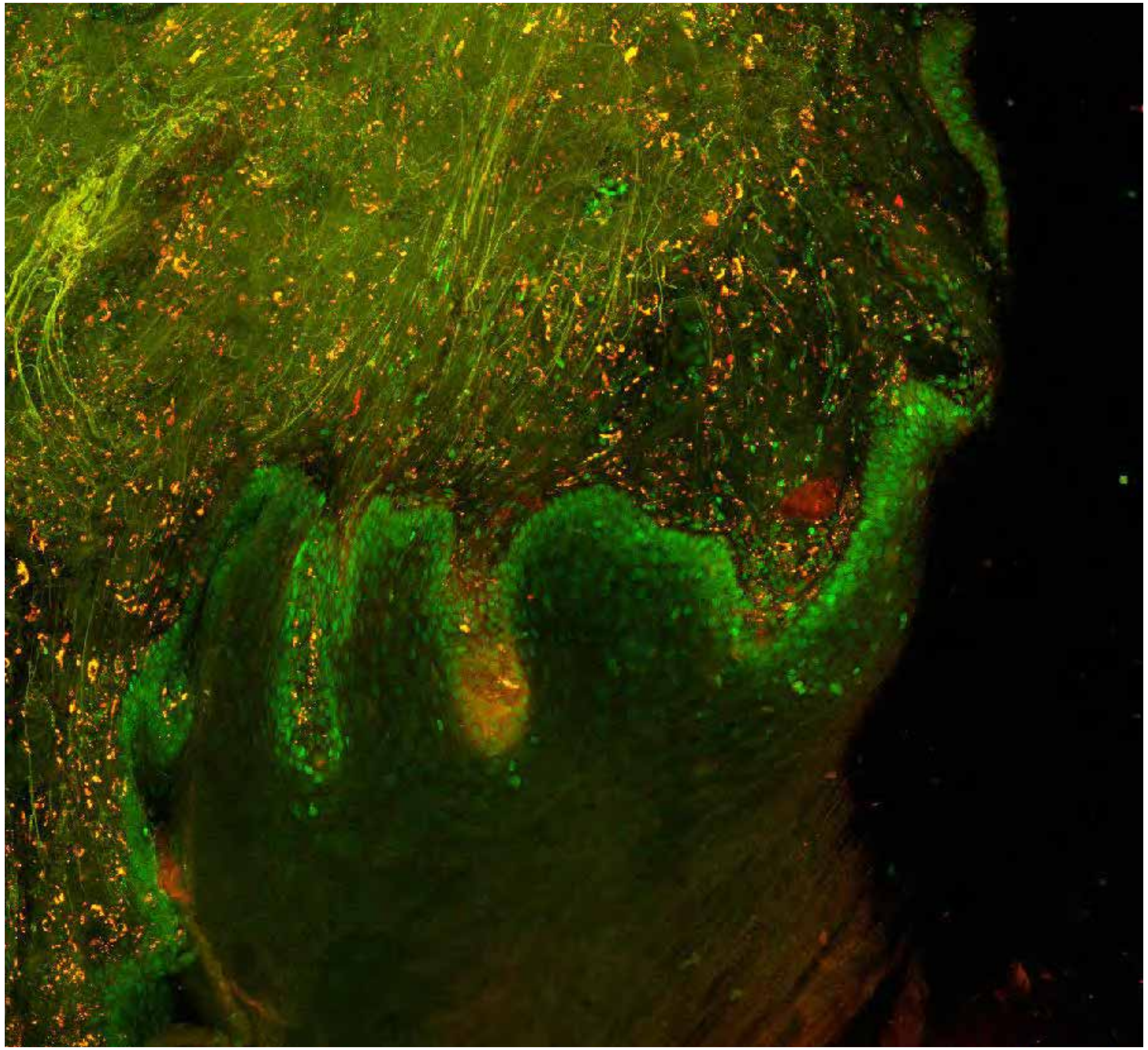

PARPi-FL (green) + Autofluorescence (red/yellow)

3403

10

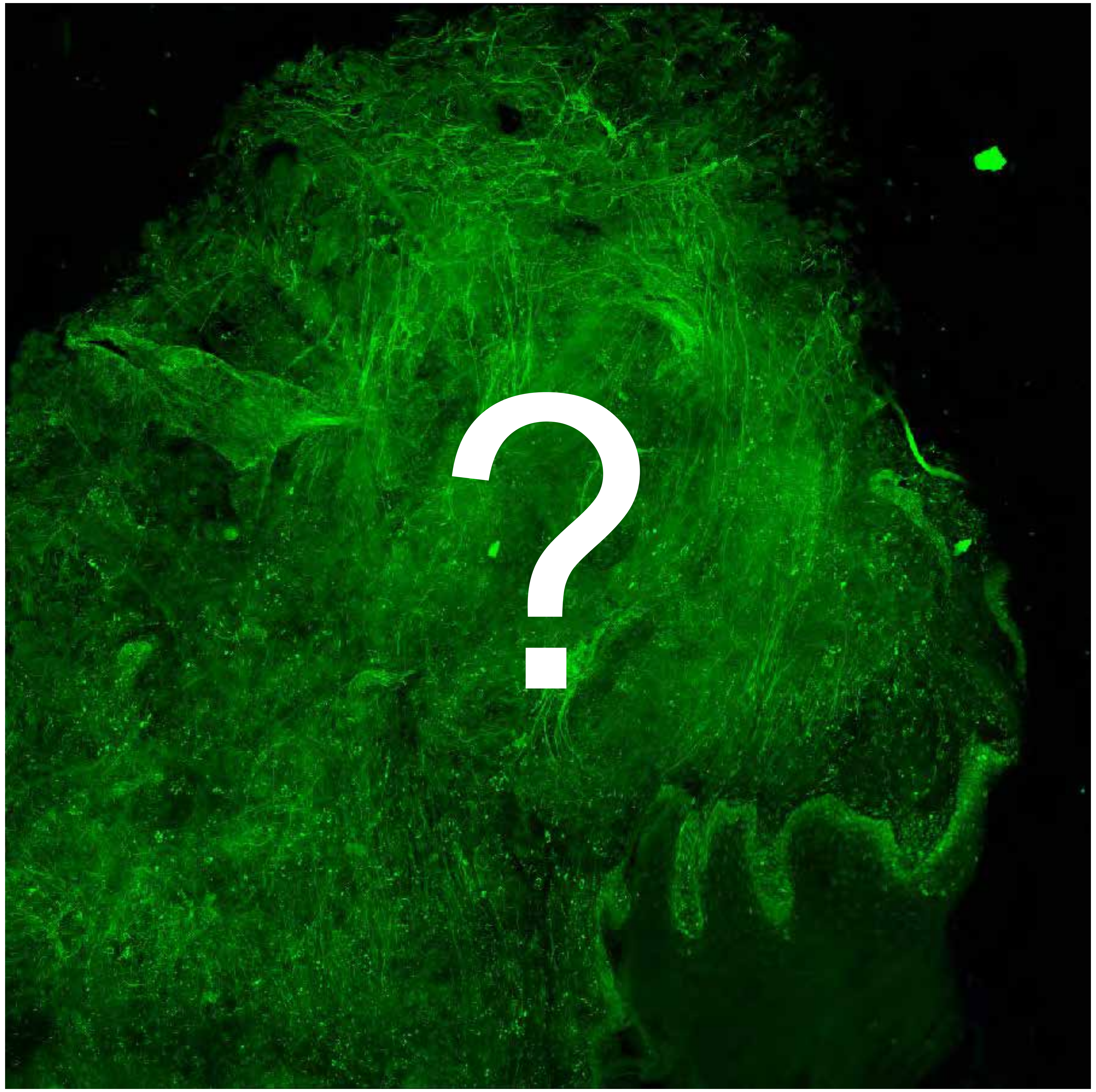


Tumor

Margin

3460

### Overview

Not available

Not available

**PARPi-FL only (green)**

**PARPi-FL (green) + Autofluorescence (red/yellow)**

3460

### Close-up

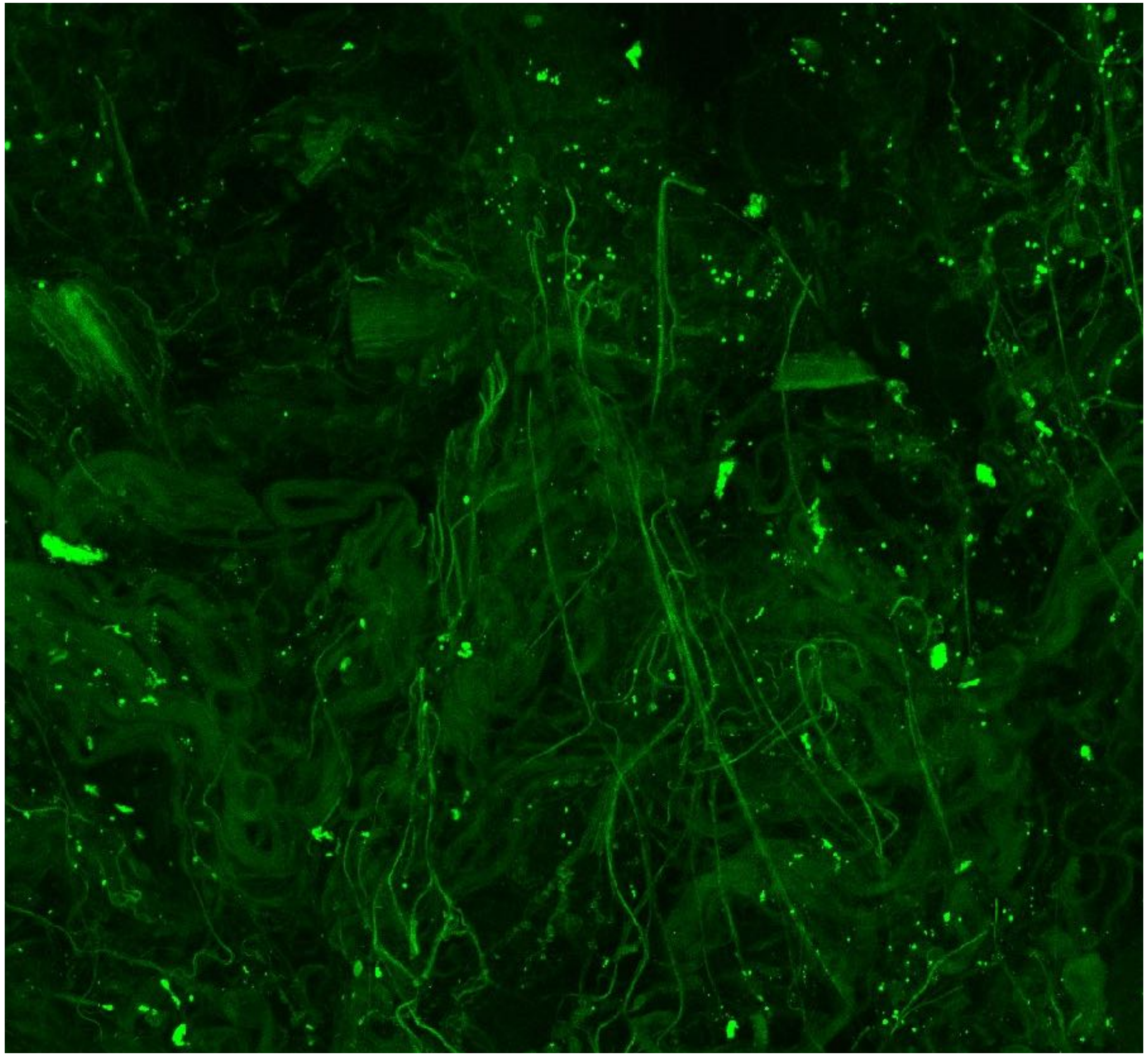

Not available

**PARPi-FL only (green)**

**PARPi-FL (green) + Autofluorescence (red/yellow)**

3460

10

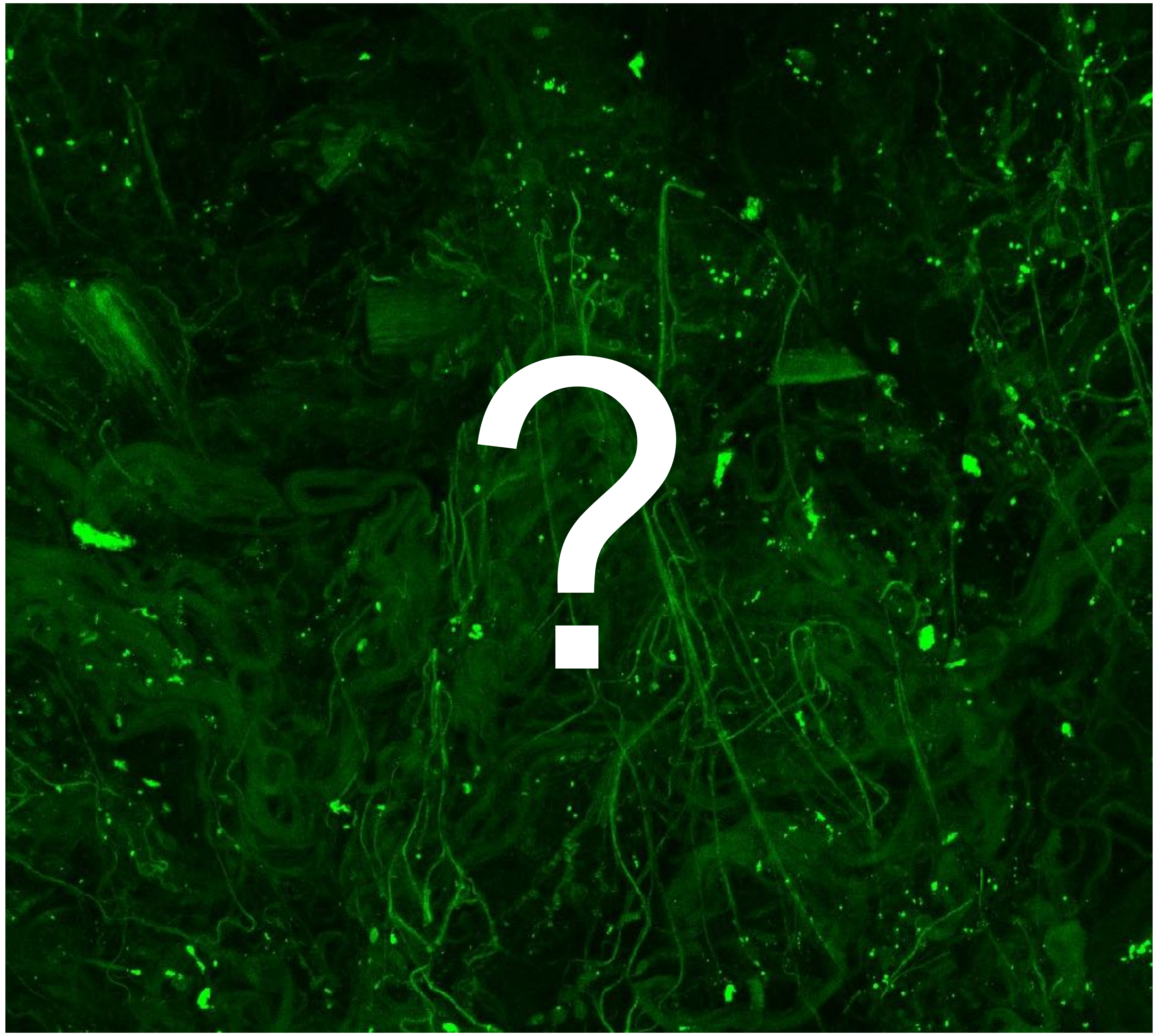

Tumor

Margin

4341

### Overview

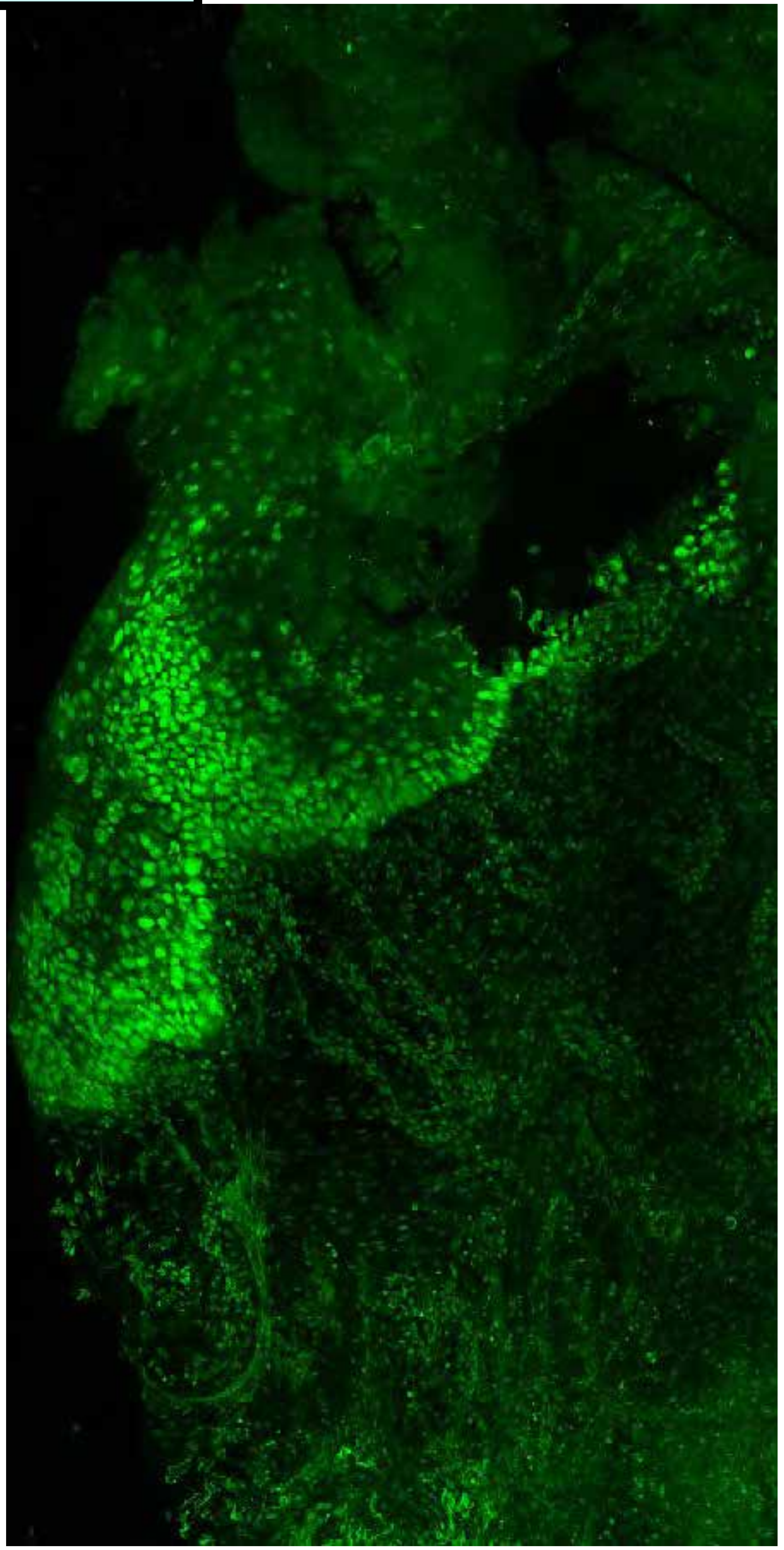

PARPi-FL only (green)

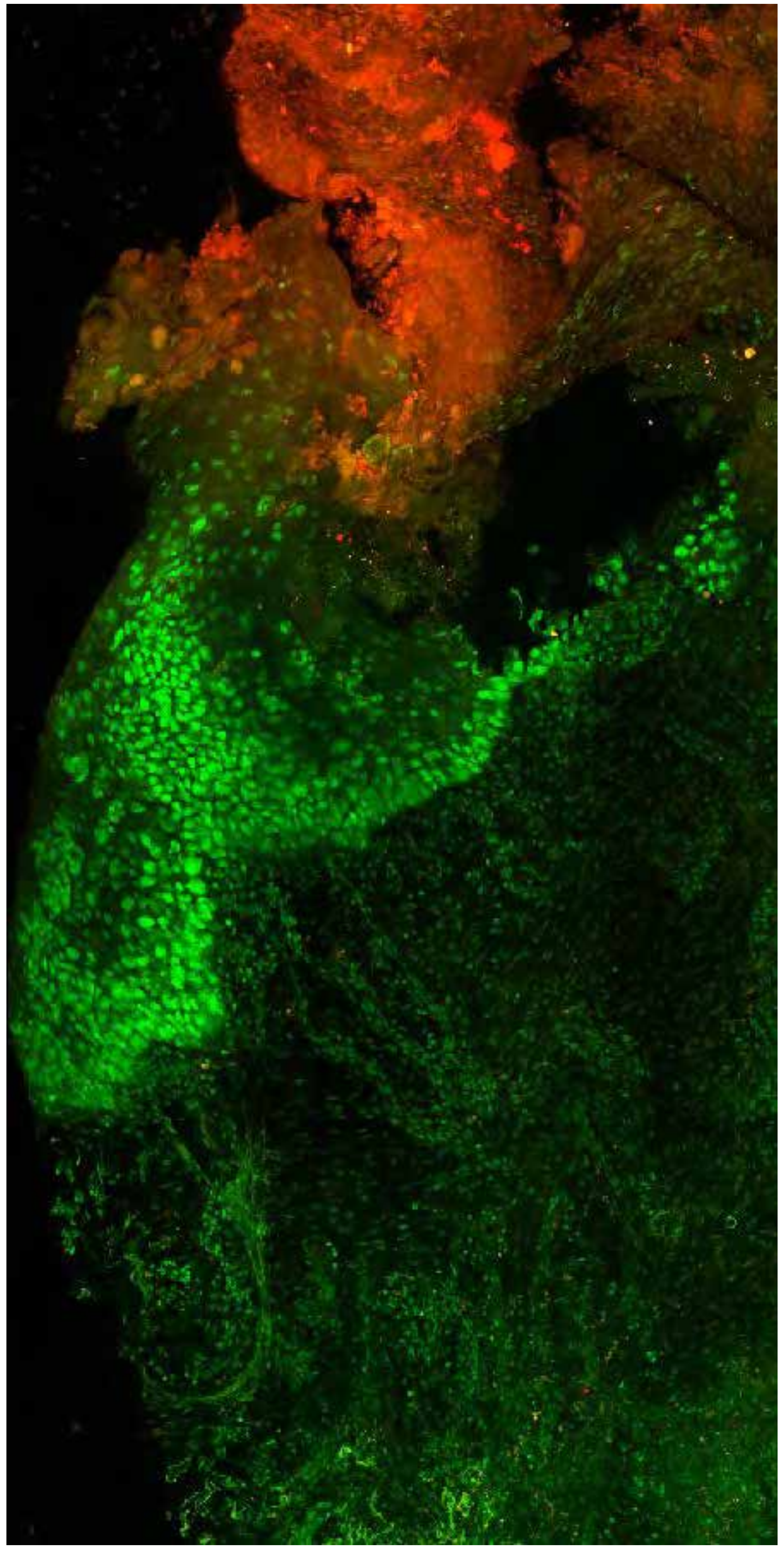

PARPi-FL (green) + Autofluorescence (red/yellow)

4341

### Close-up

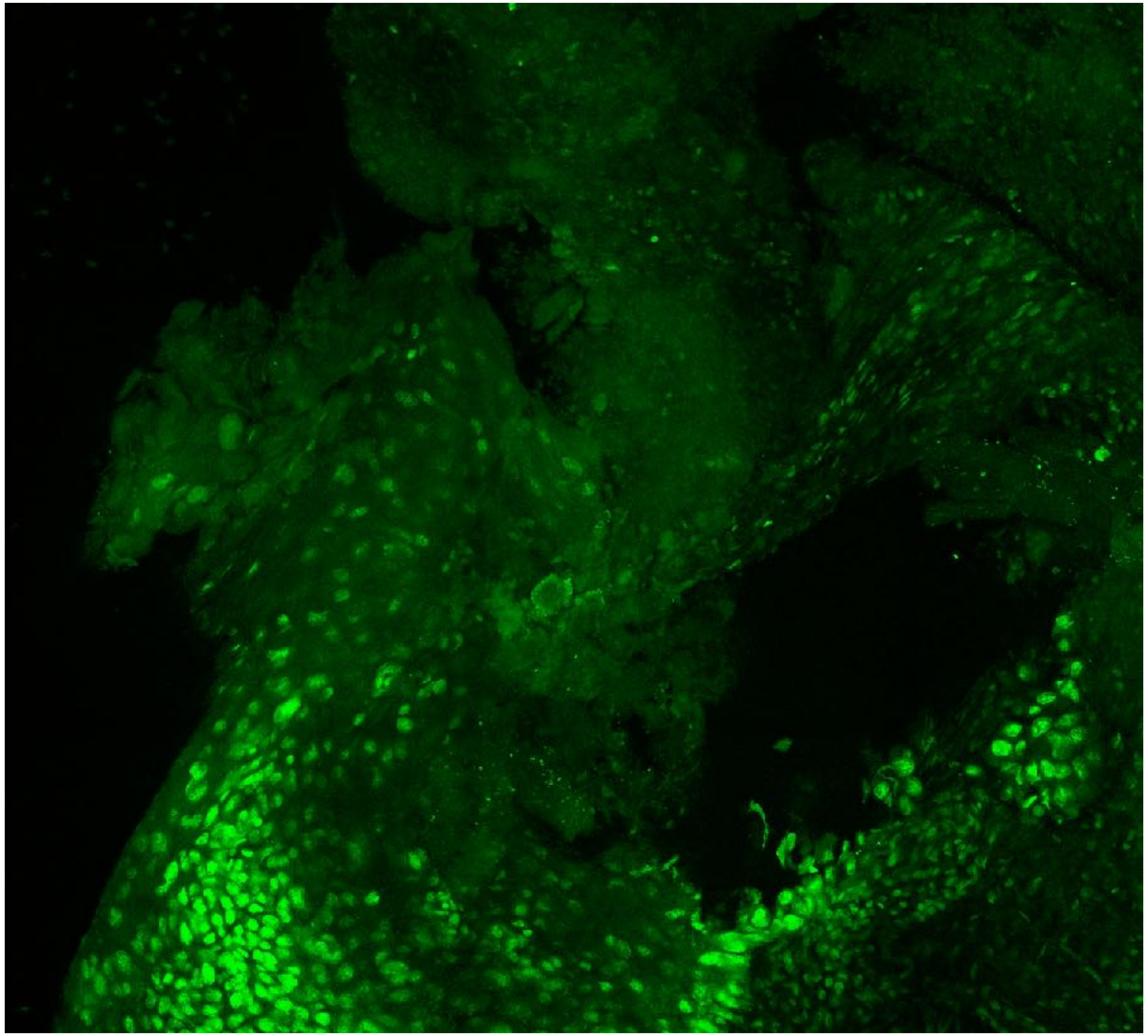

PARPi-FL only (green)

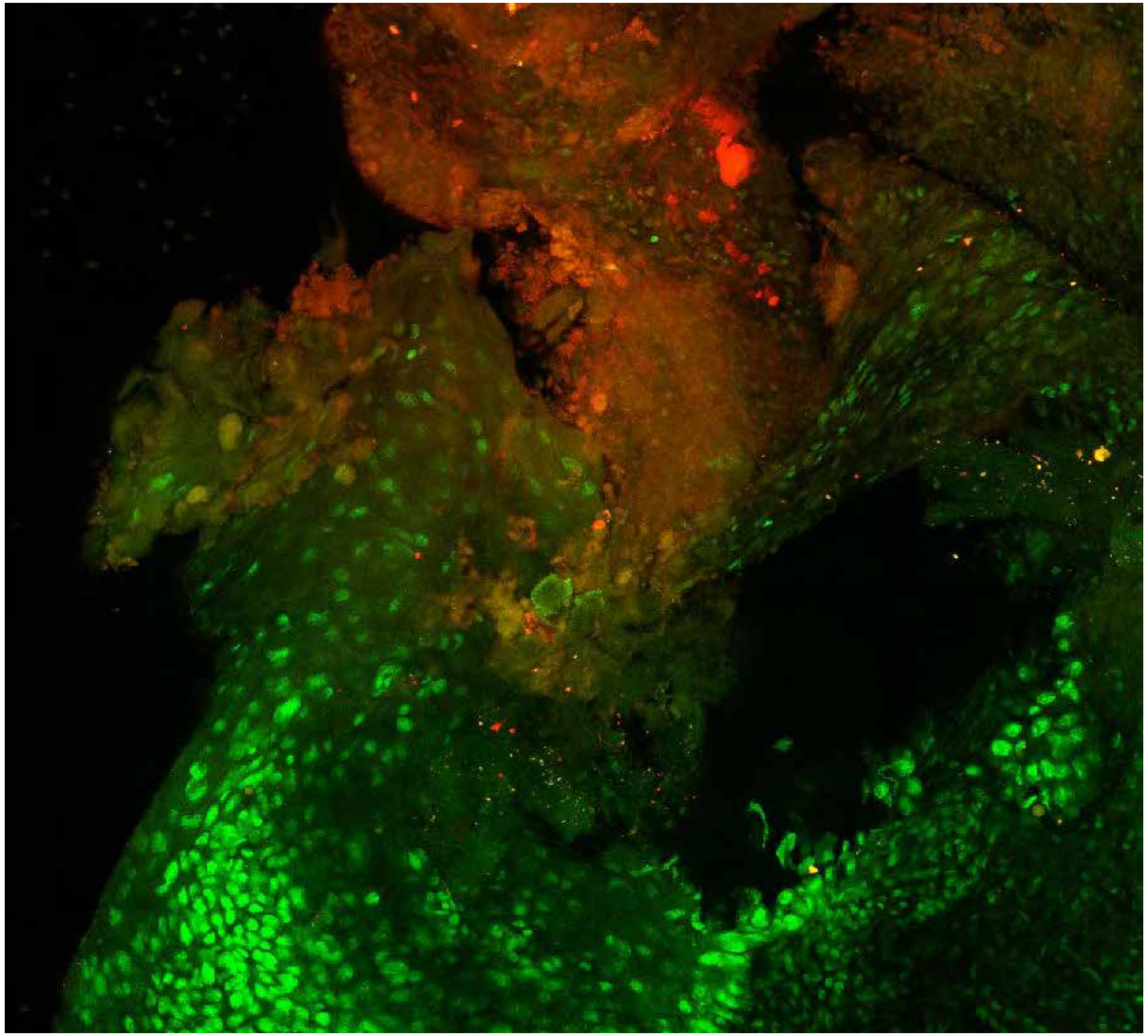

PARPi-FL (green) + Autofluorescence (red/yellow)

4341

### Close-up

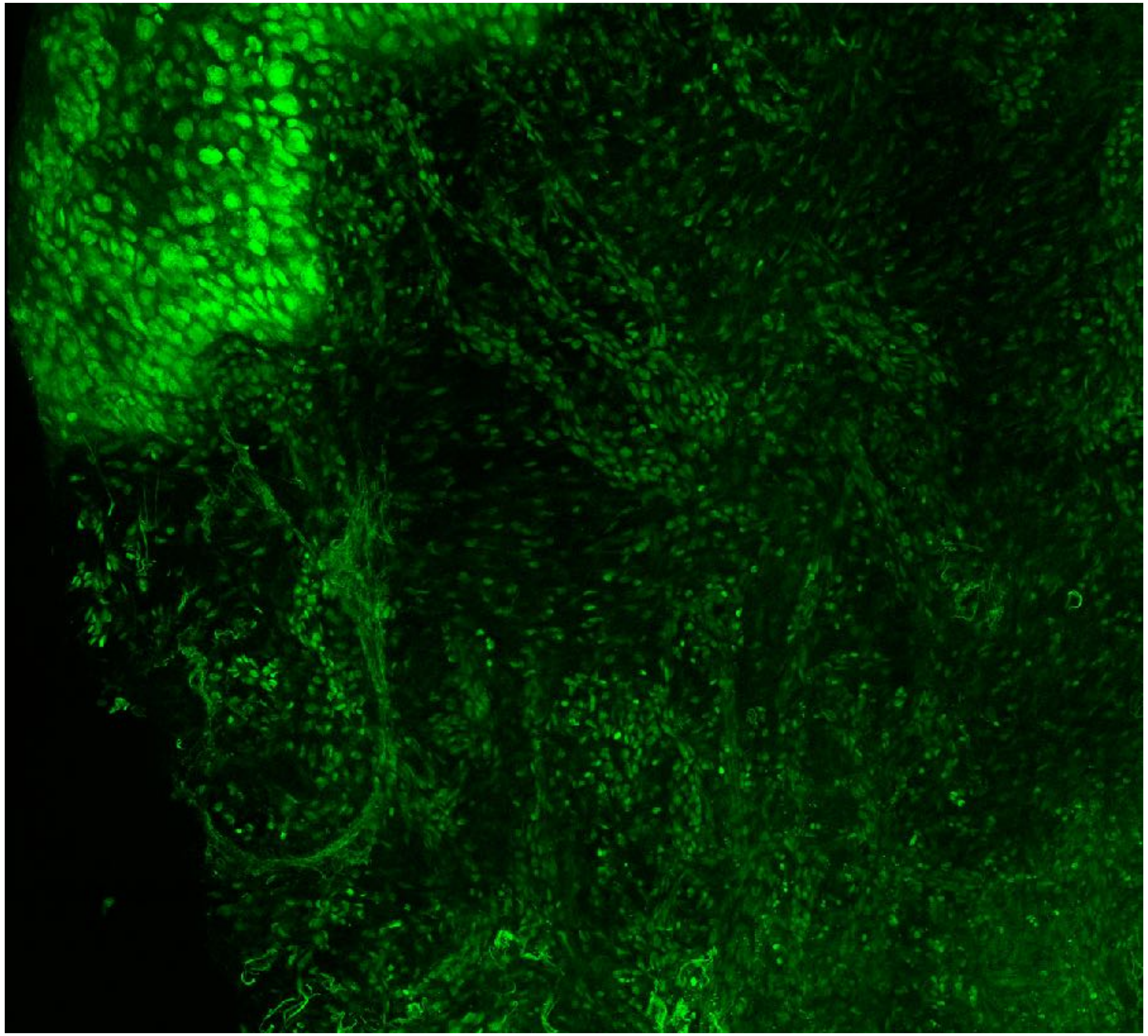

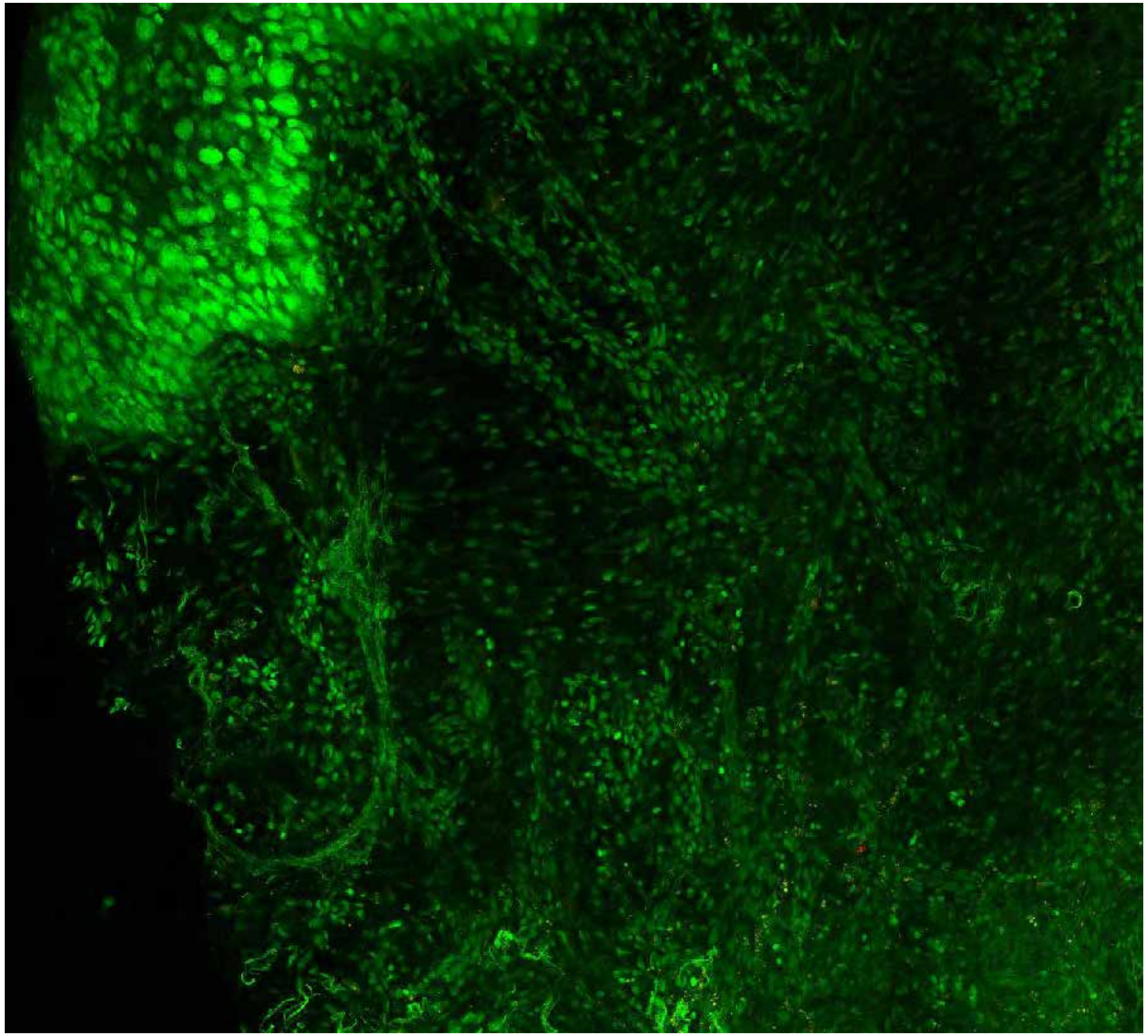

**PARPi-FL only (green)** **PARPi-FL (green) + Autofluorescence (red/yellow)**

4341

### Close-up

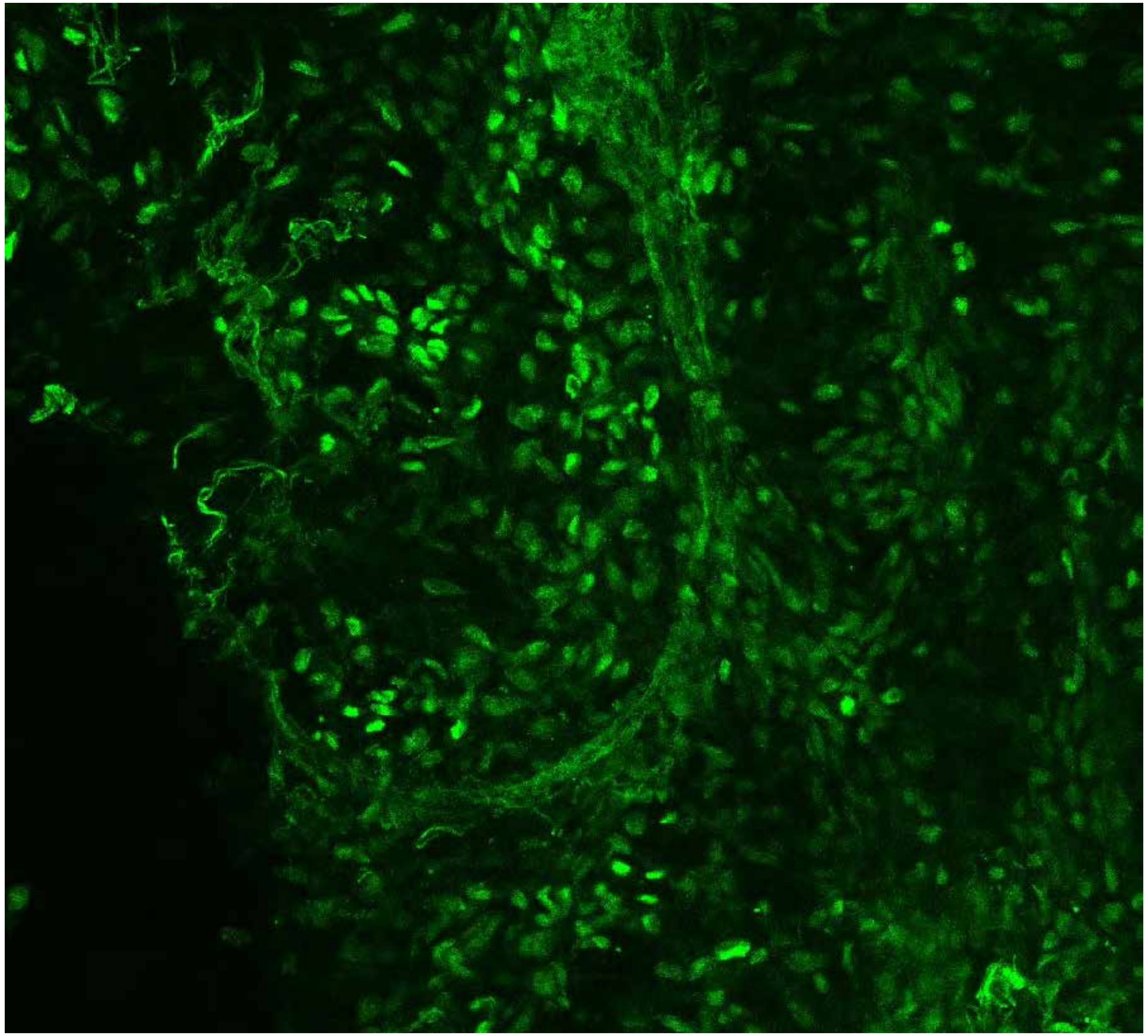

PARPi-FL only (green)

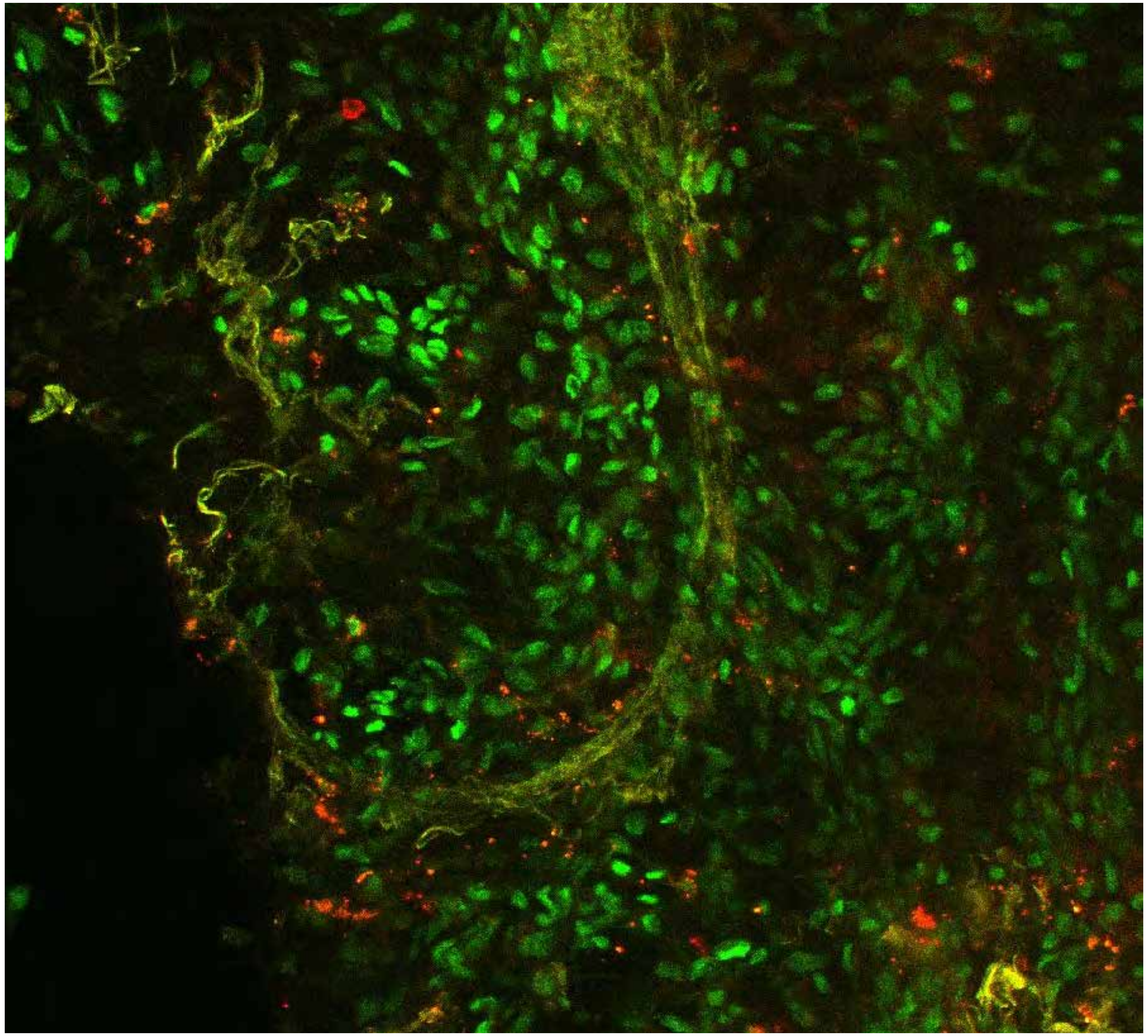

PARPi-FL (green) + Autofluorescence (red/yellow)

4341

10

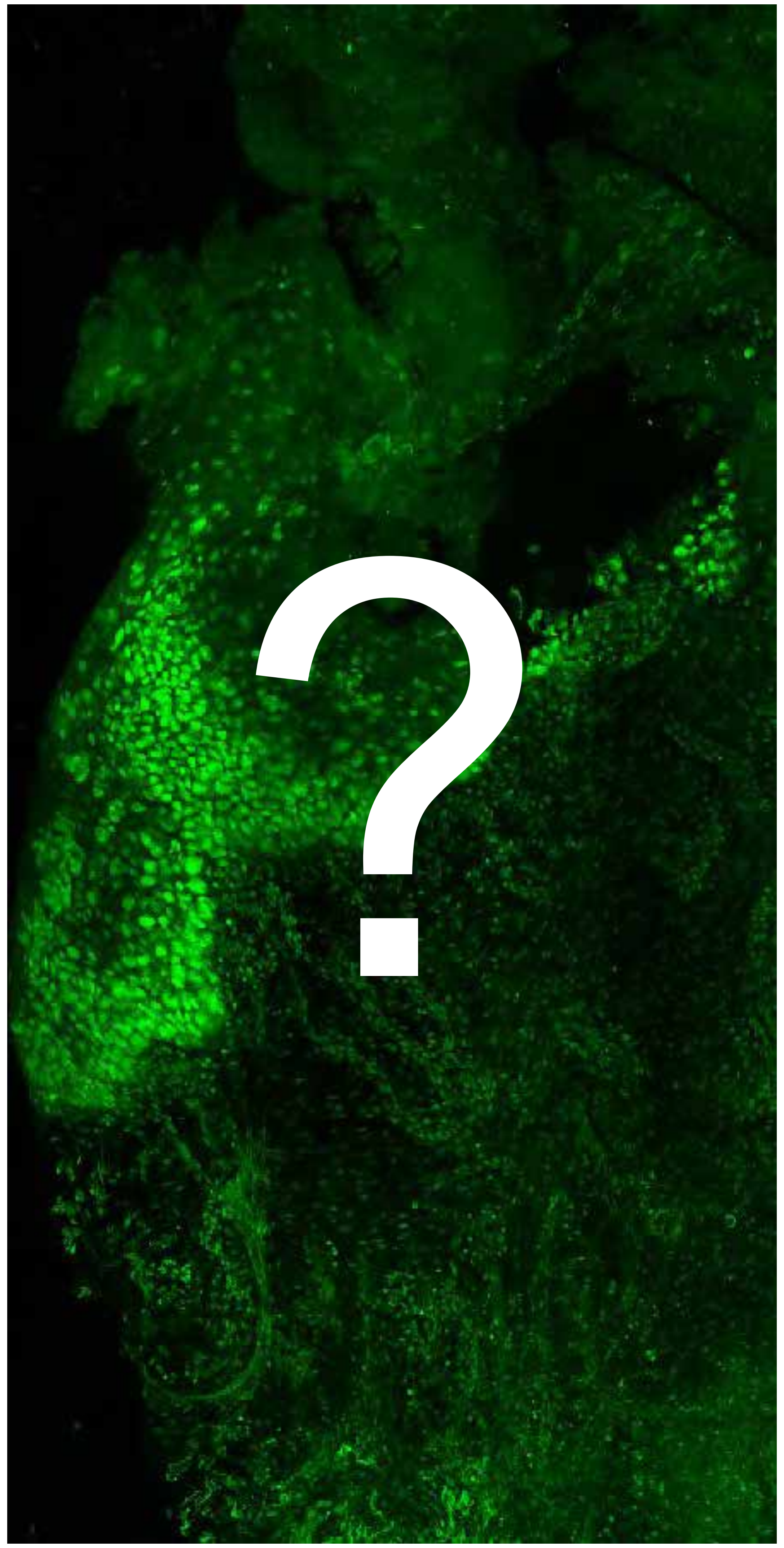


Tumor

Margin

4570

### Overview

Not available

Not available

**PARPi-FL only (green)**

**PARPi-FL (green) + Autofluorescence (red/yellow)**

4570

### Close-up

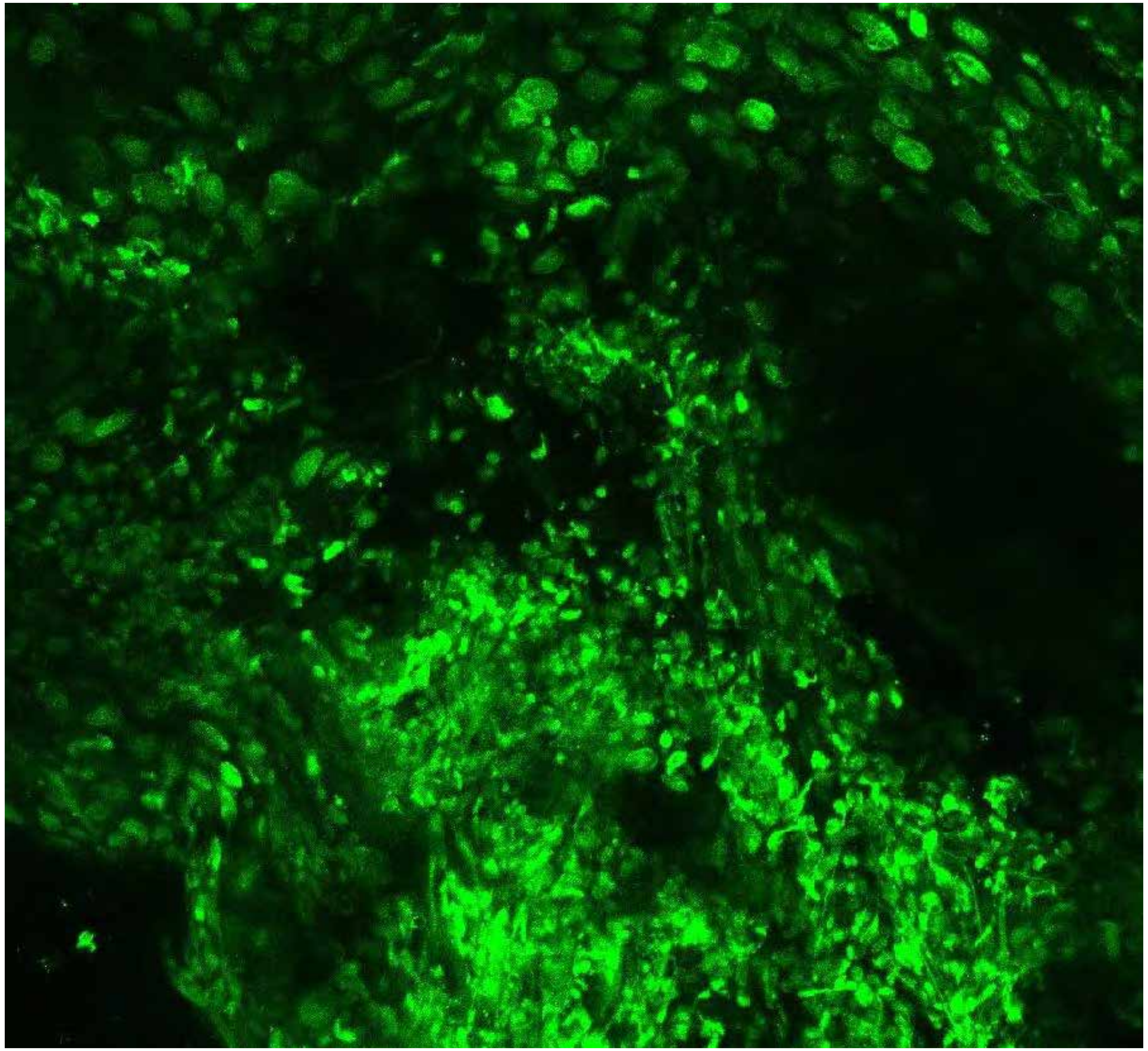

PARPi-FL only (green)

Not available

PARPi-FL (green) + Autofluorescence (red/yellow)

4570

### Close-up

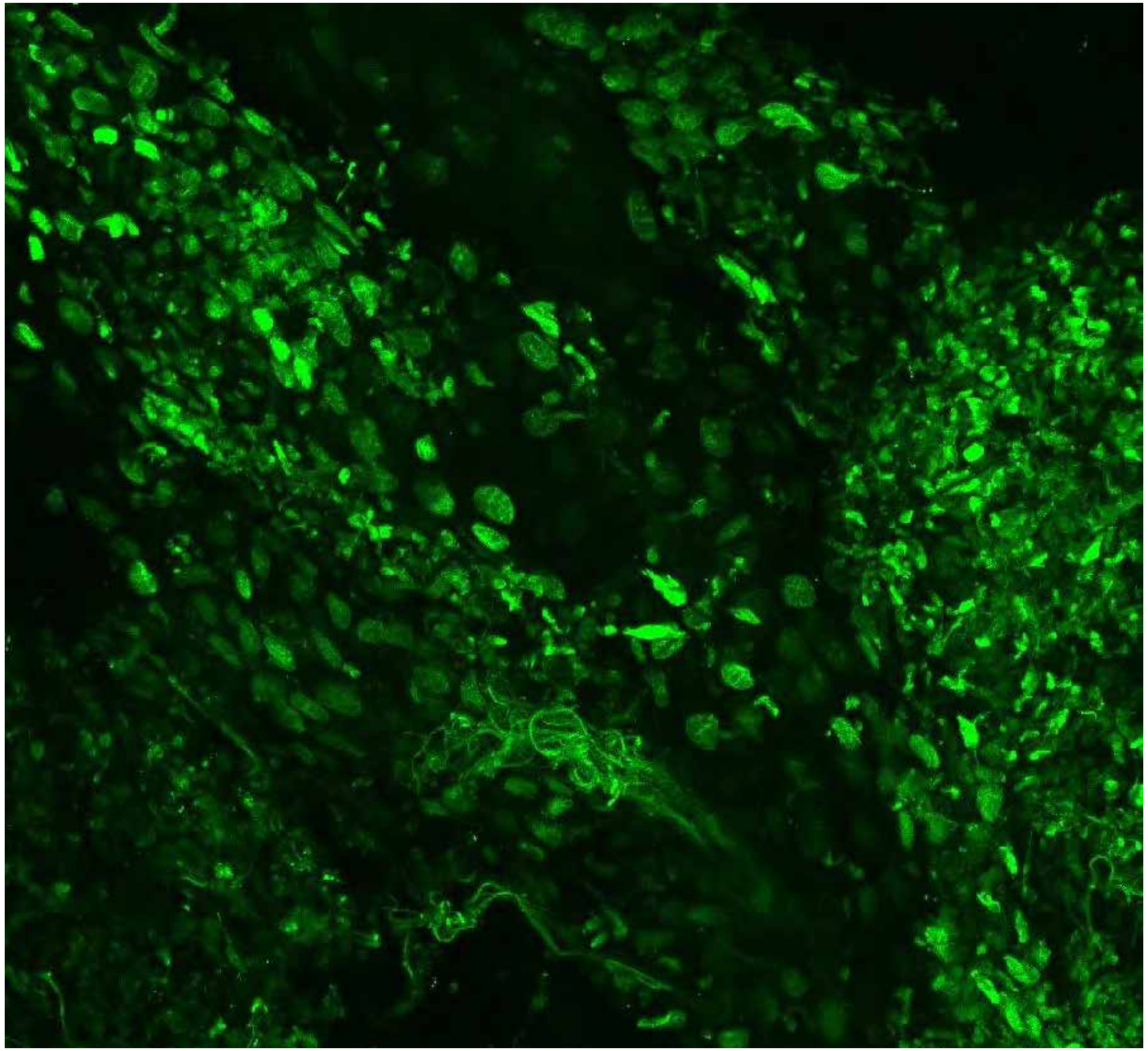

Not available

**PARPi-FL only (green)**

**PARPi-FL (green) + Autofluorescence (red/yellow)**

4570

### Close-up

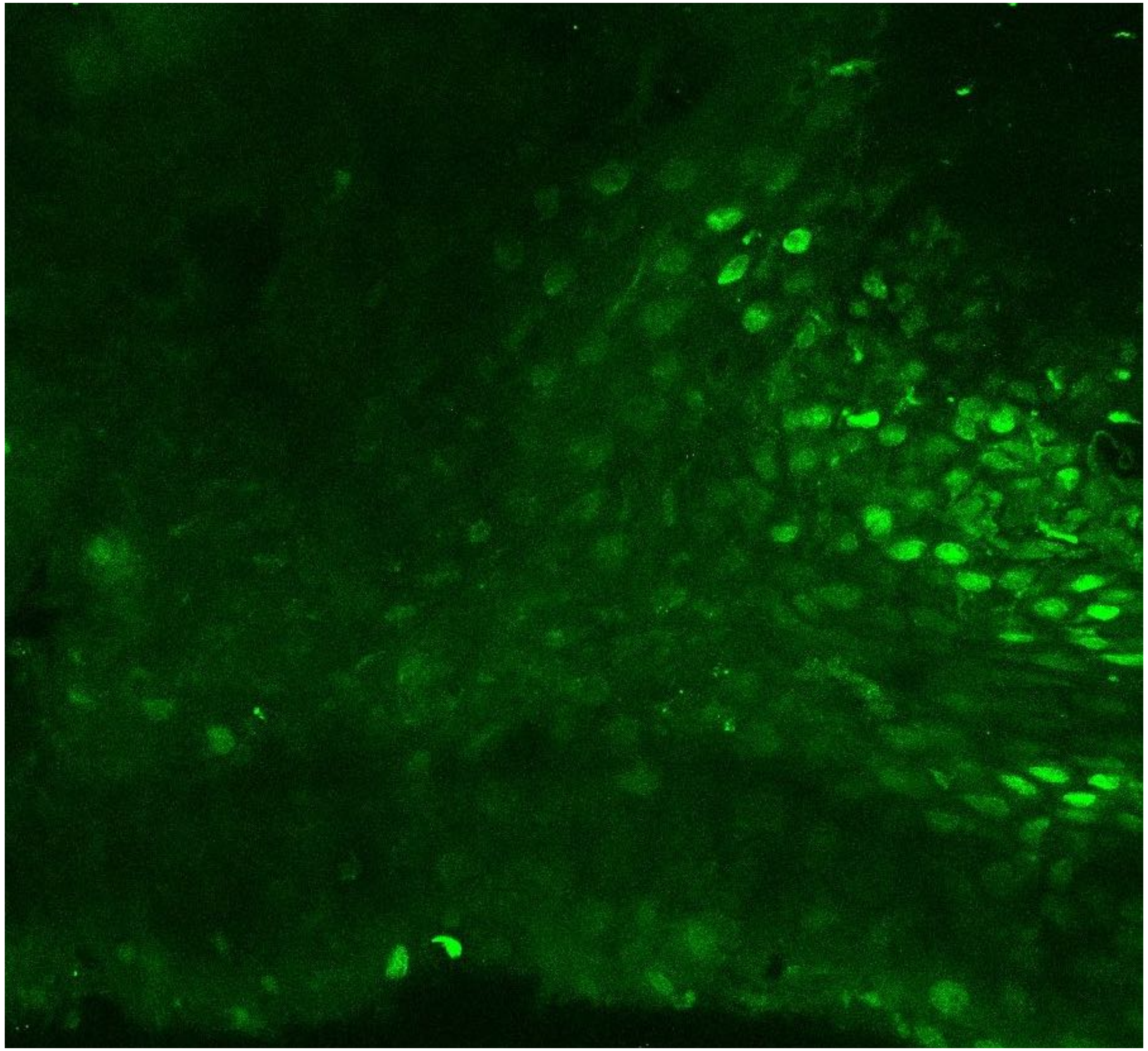

PARPi-FL only (green)

Not available

PARPi-FL (green) + Autofluorescence (red/yellow)

4570

### Close-up

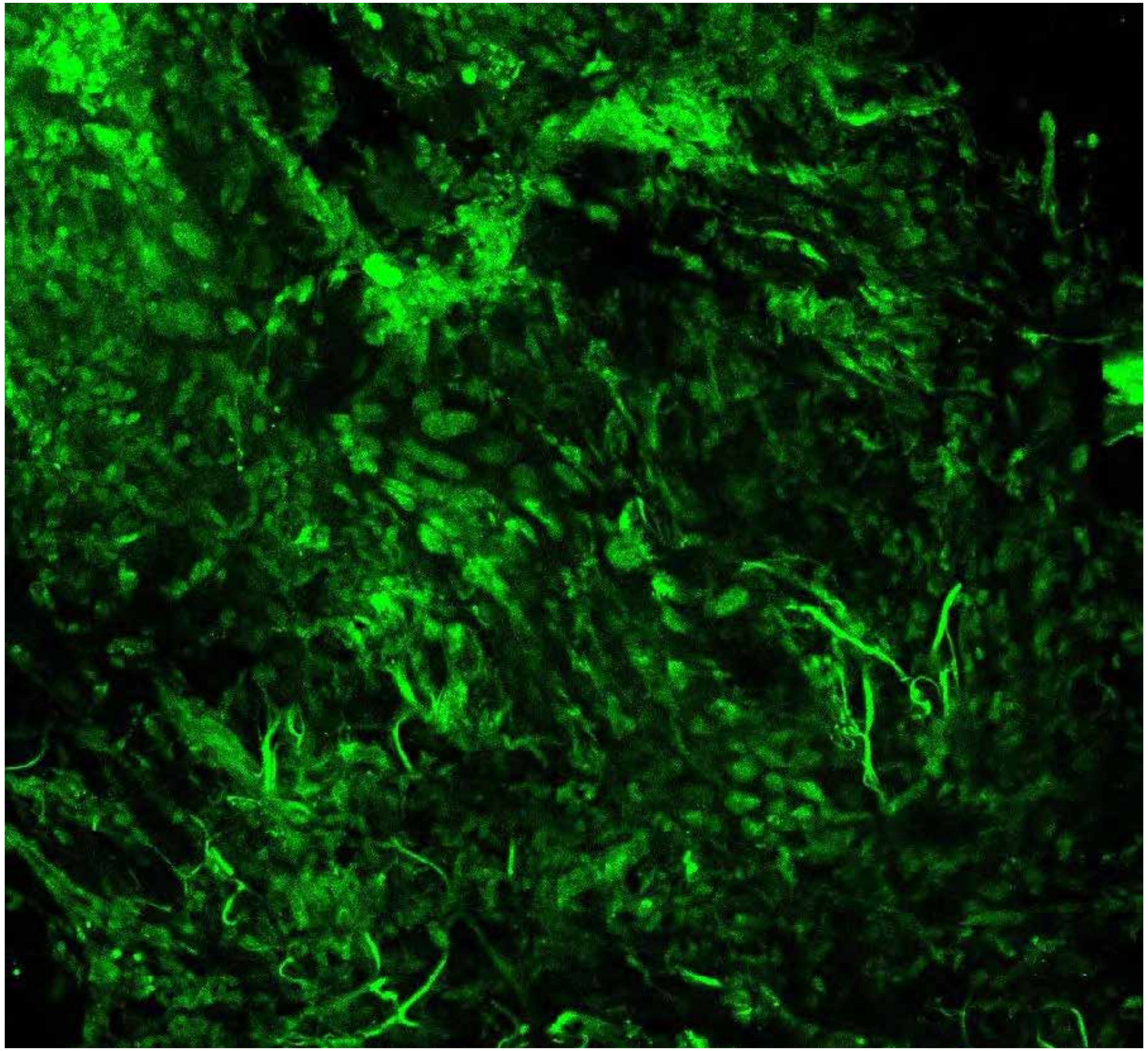

Not available

**PARPi-FL only (green)**

**PARPi-FL (green) + Autofluorescence (red/yellow)**

4570

10

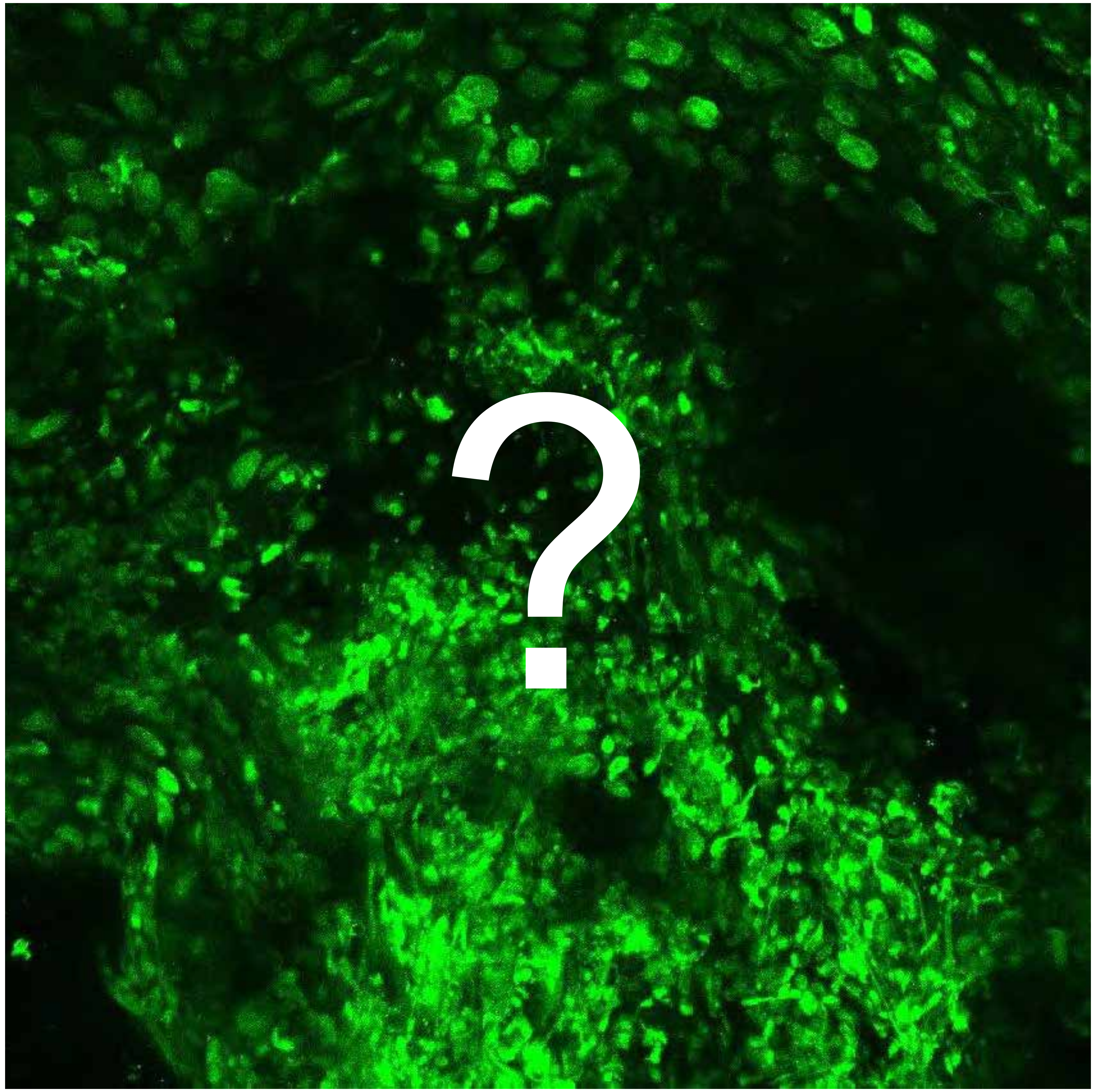


Tumor

Margin

4574

### Overview

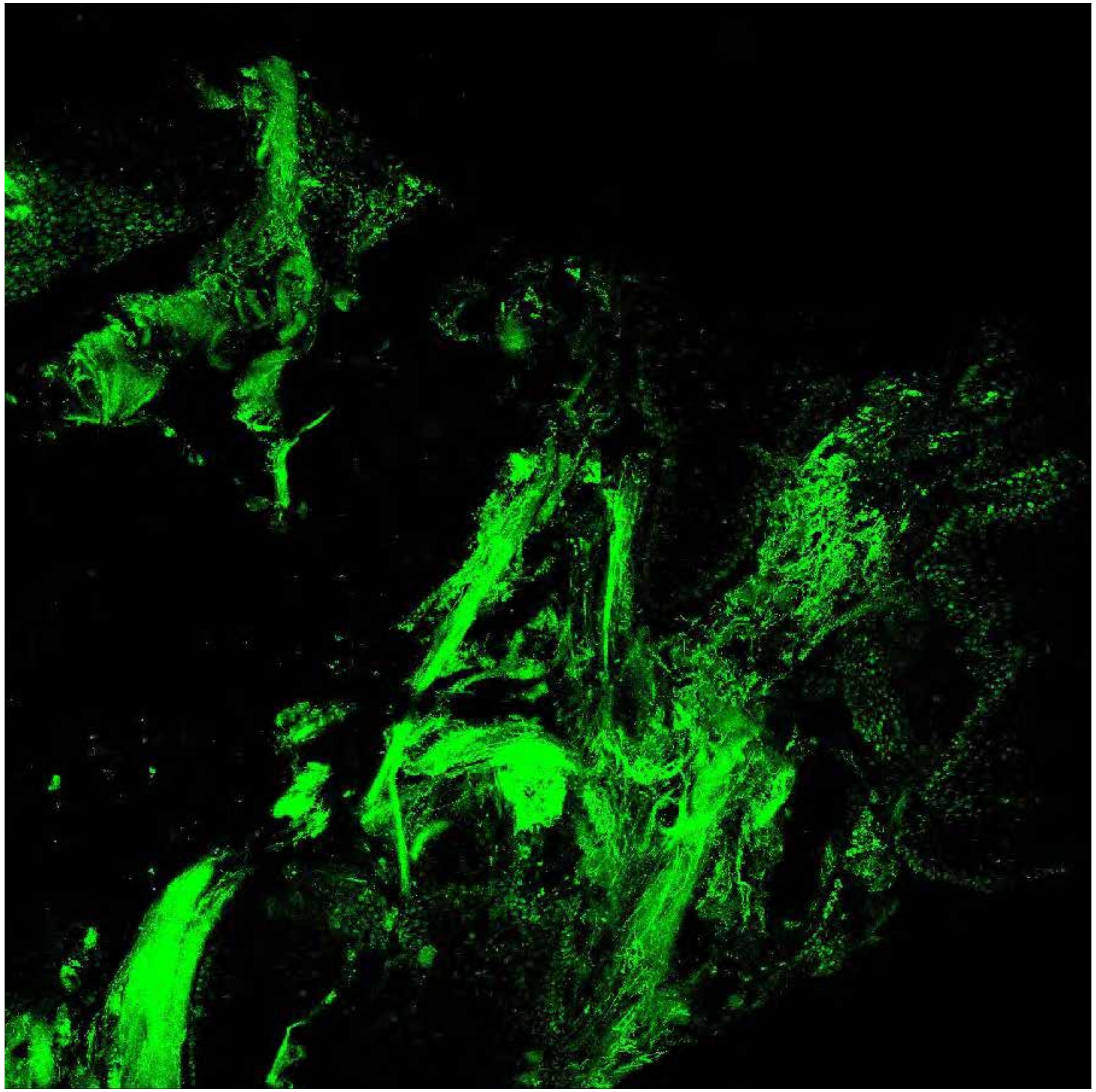

**PARPi-FL only (green)**

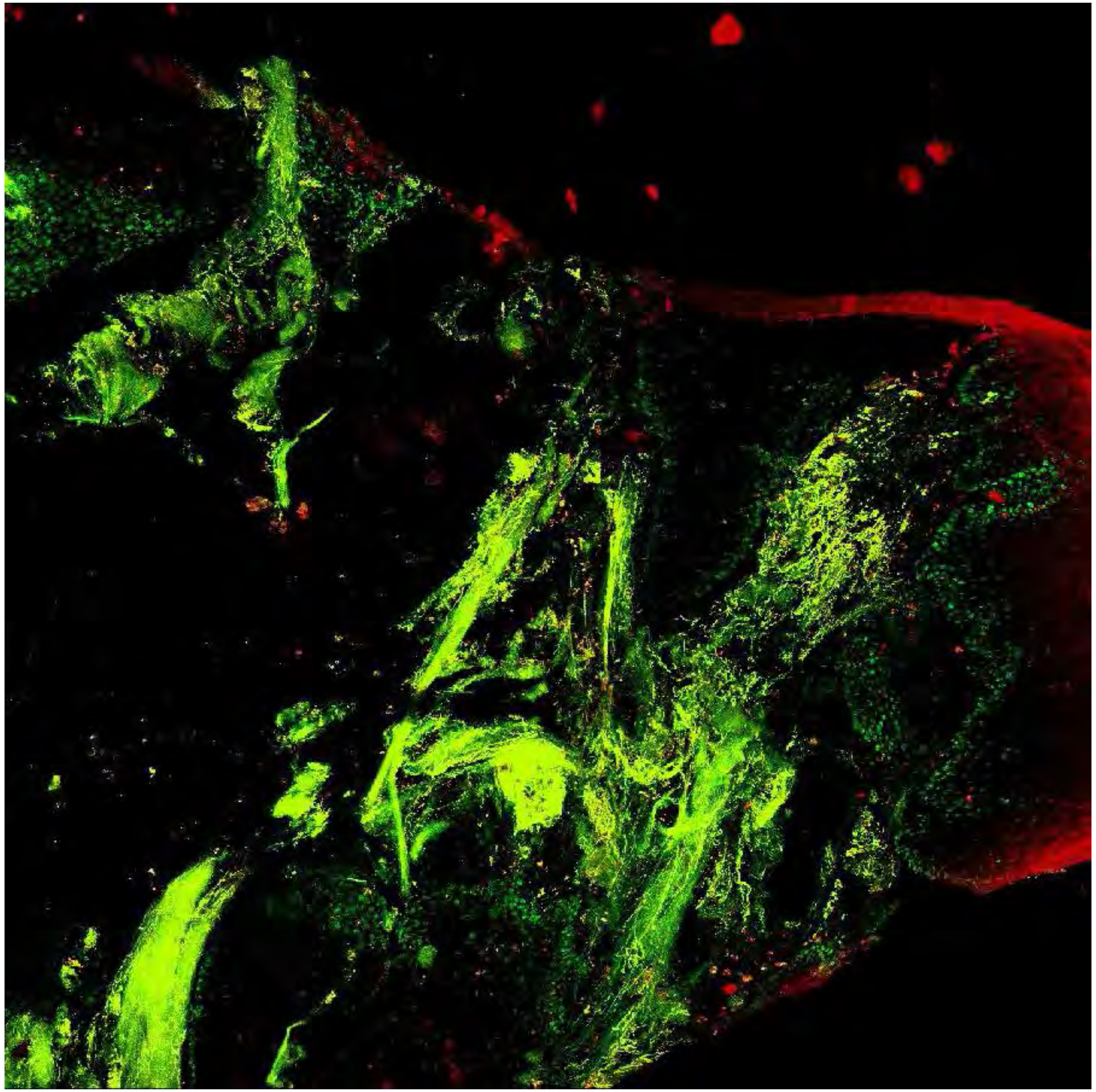

**PARPi-FL (green) + Autofluorescence (red/yellow)**

4574

### Close-up

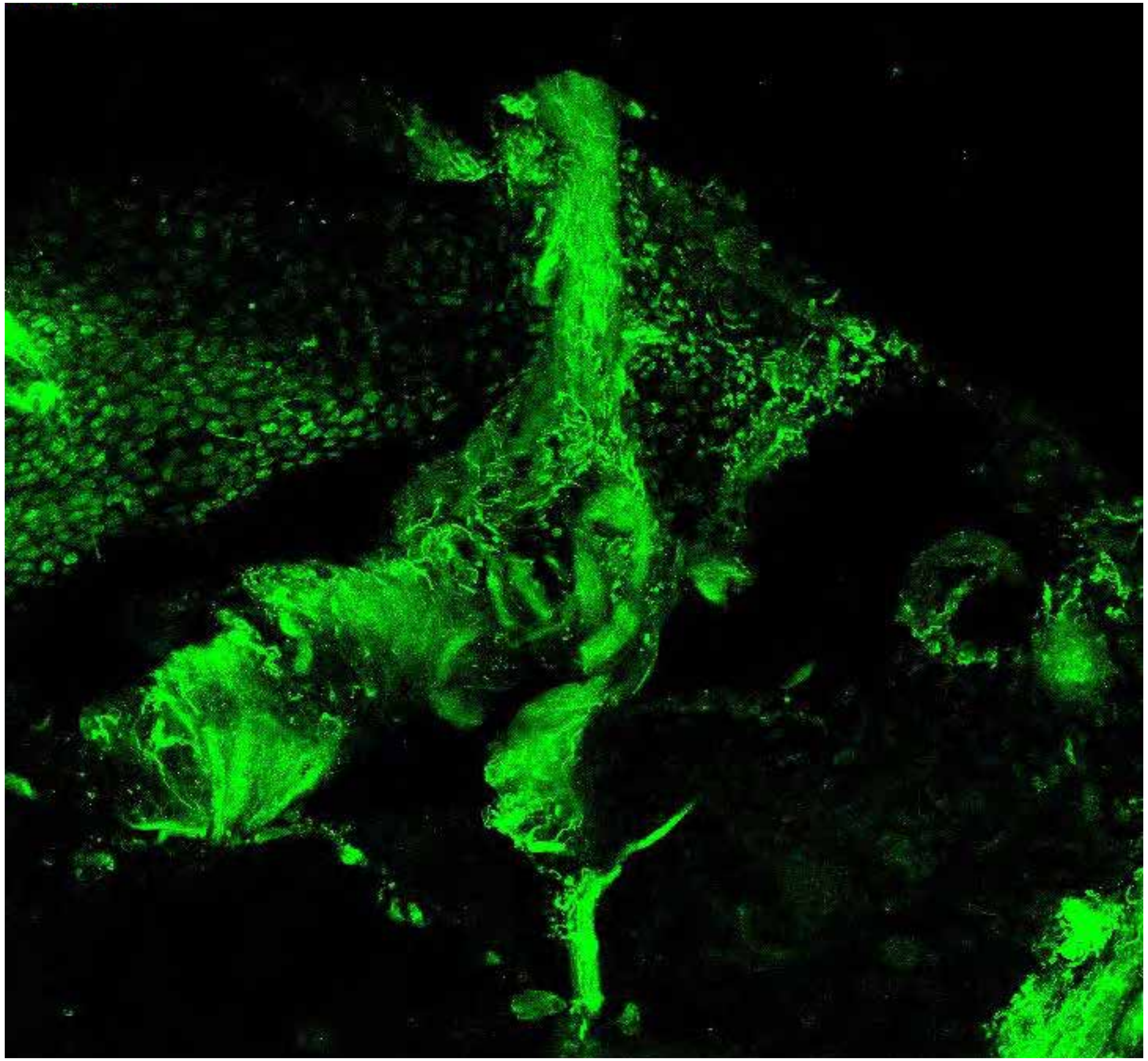

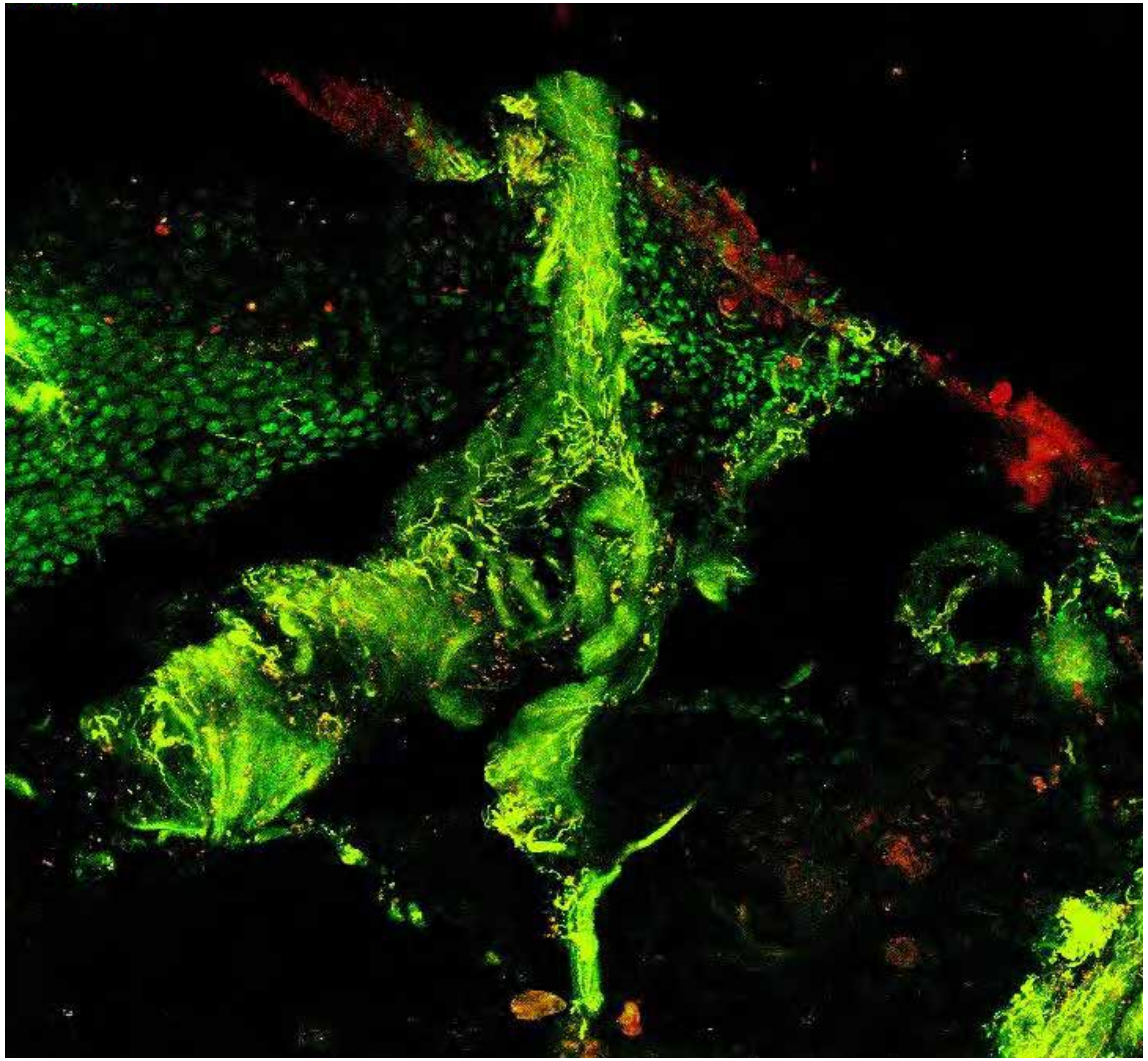

**PARPi-FL only (green)**

**PARPi-FL (green) + Autofluorescence (red/yellow)**

4574

### Close-up

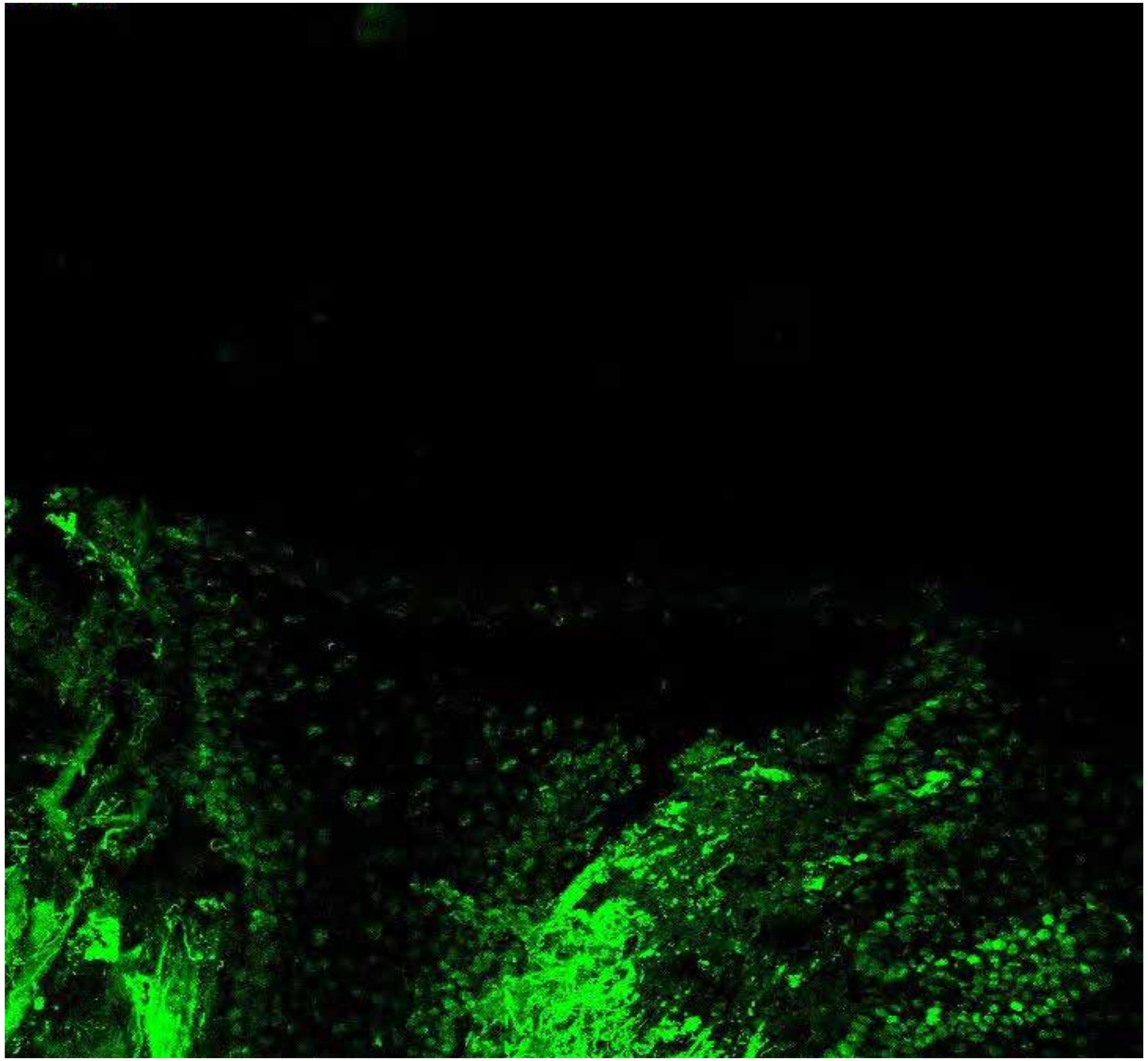

PARPi-FL only (green)

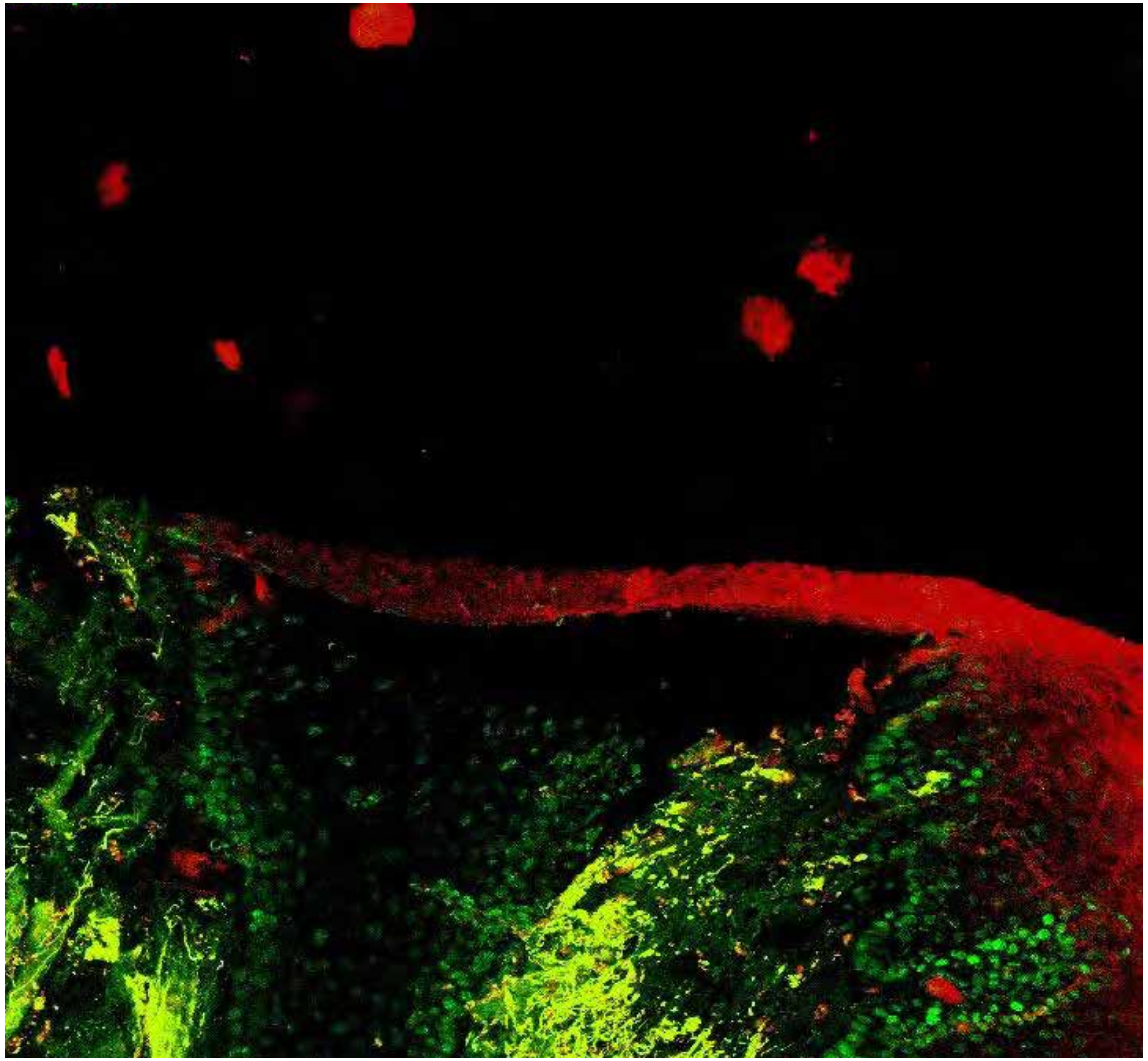

PARPi-FL (green) + Autofluorescence (red/yellow)

4574

### Close-up

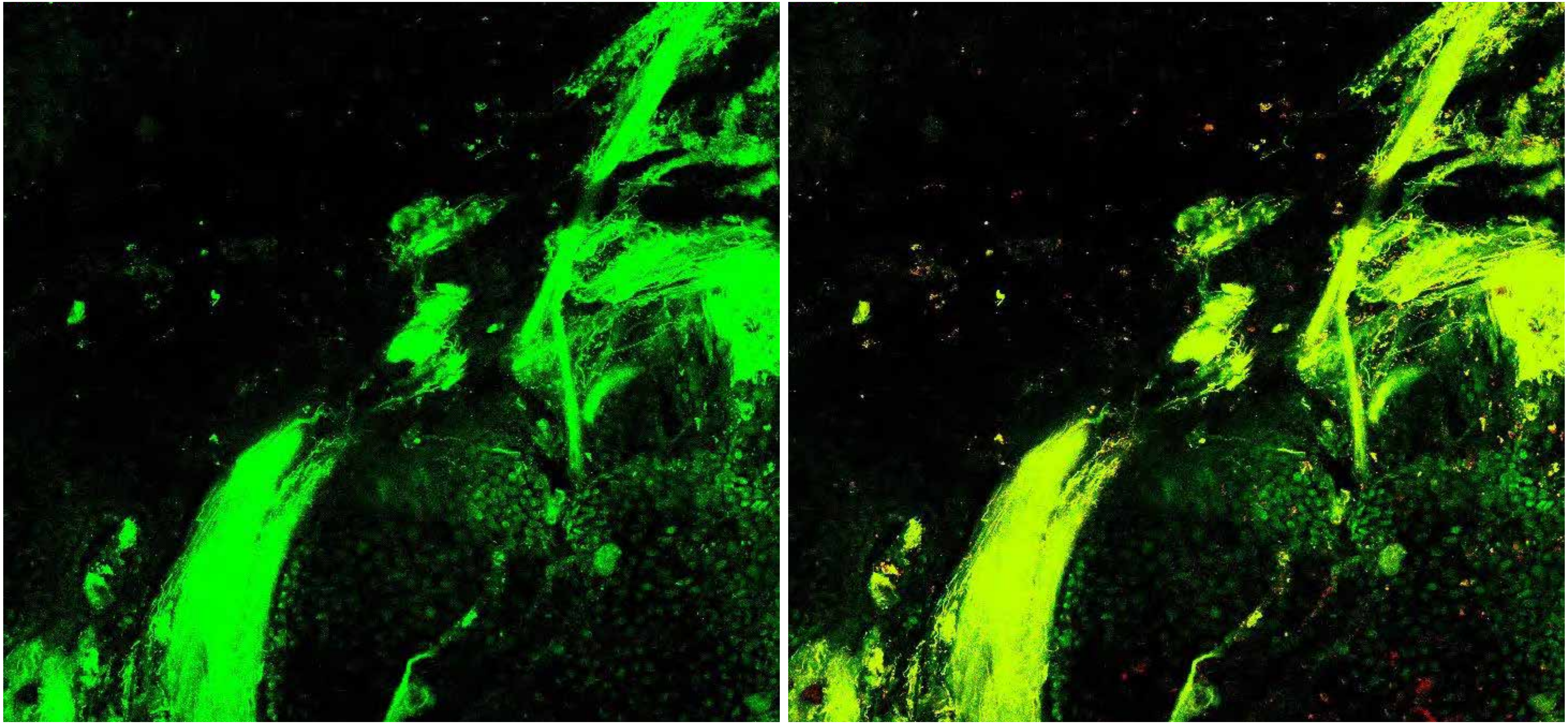

PARPi-FL only (green)

PARPi-FL (green) + Autofluorescence (red/yellow)

4574

### Close-up

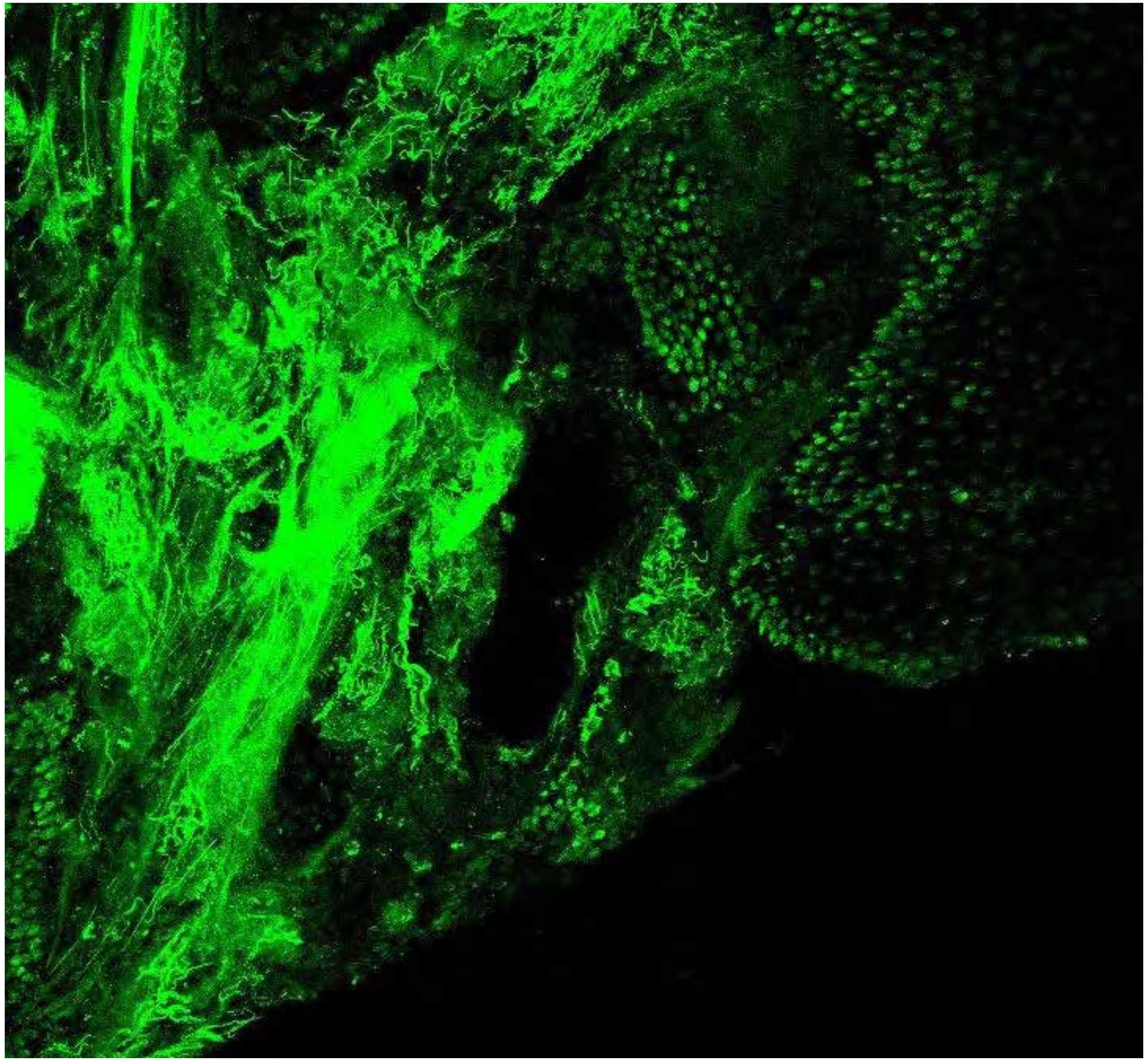

PARPi-FL only (green)

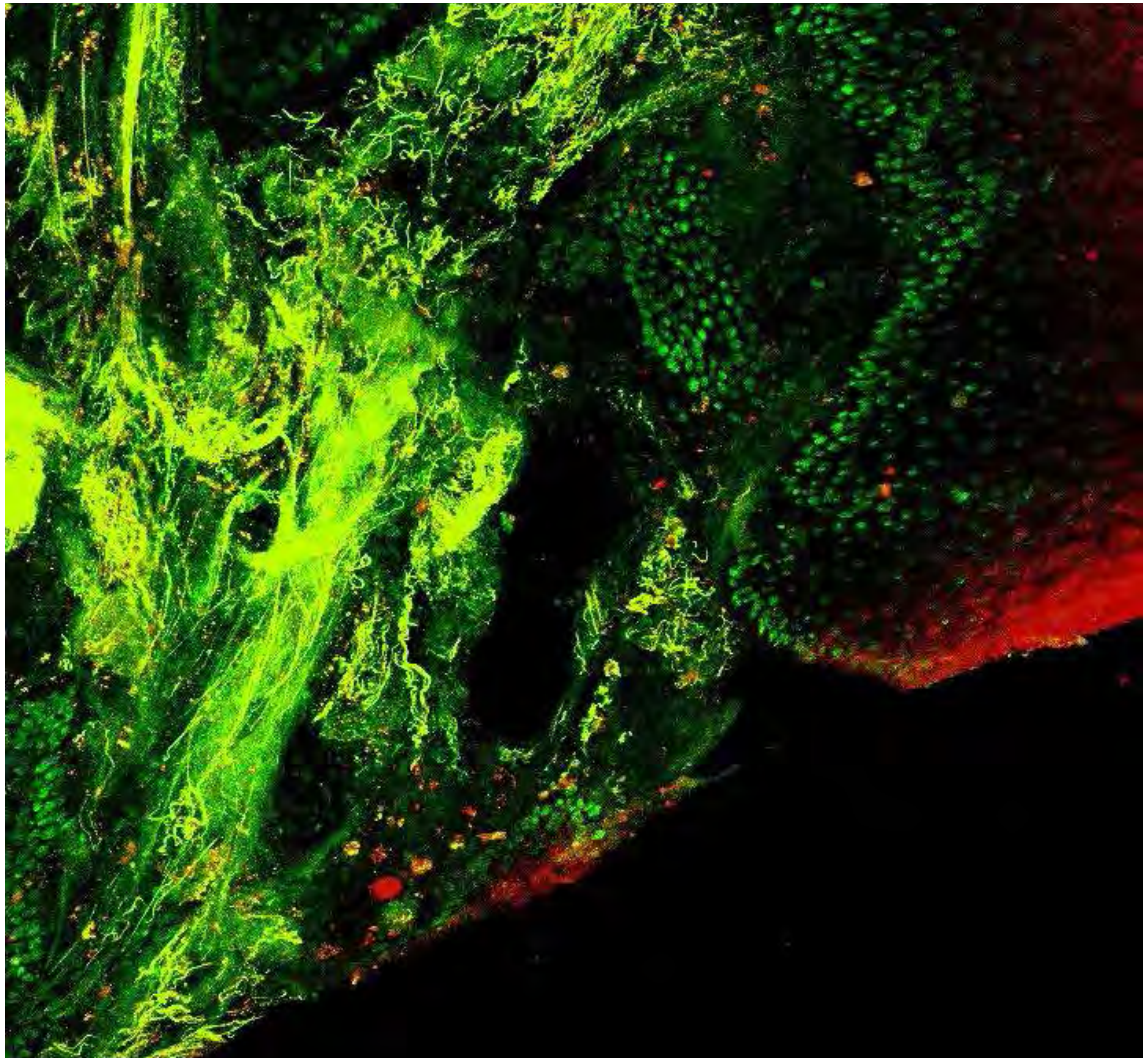

PARPi-FL (green) + Autofluorescence (red/yellow)

4574

10

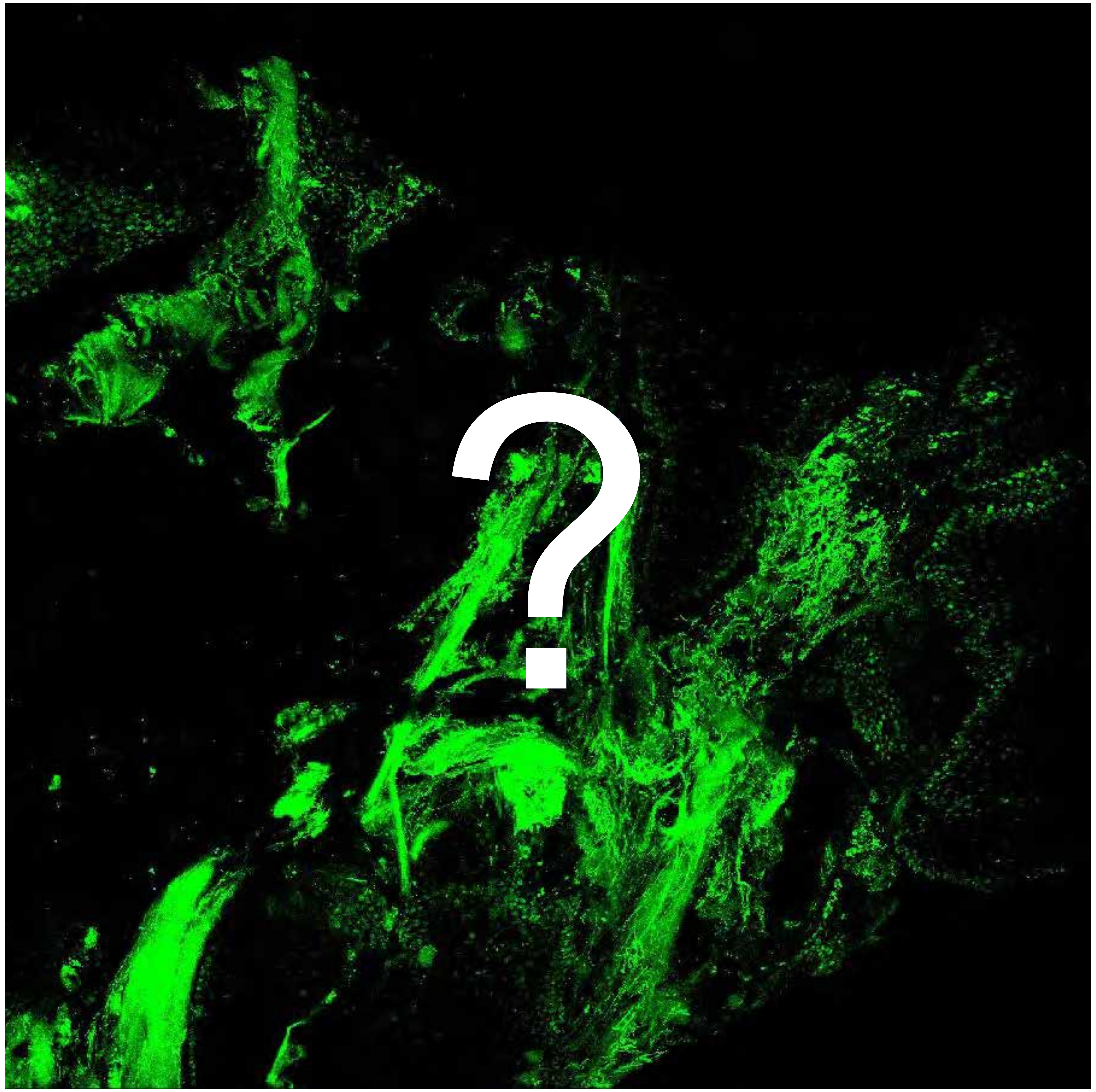


Tumor

Margin

5347

### Overview

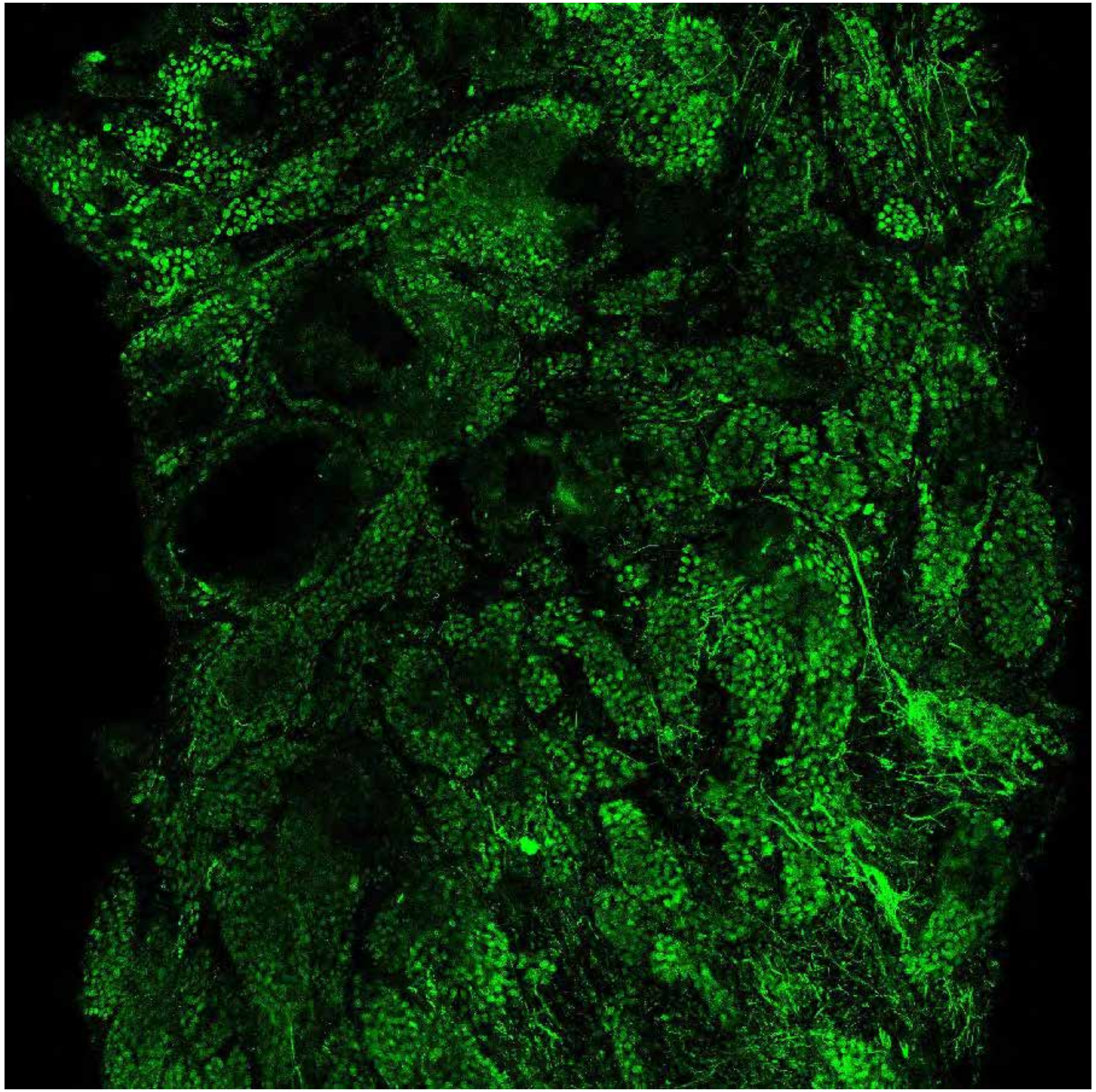

PARPi-FL only (green)

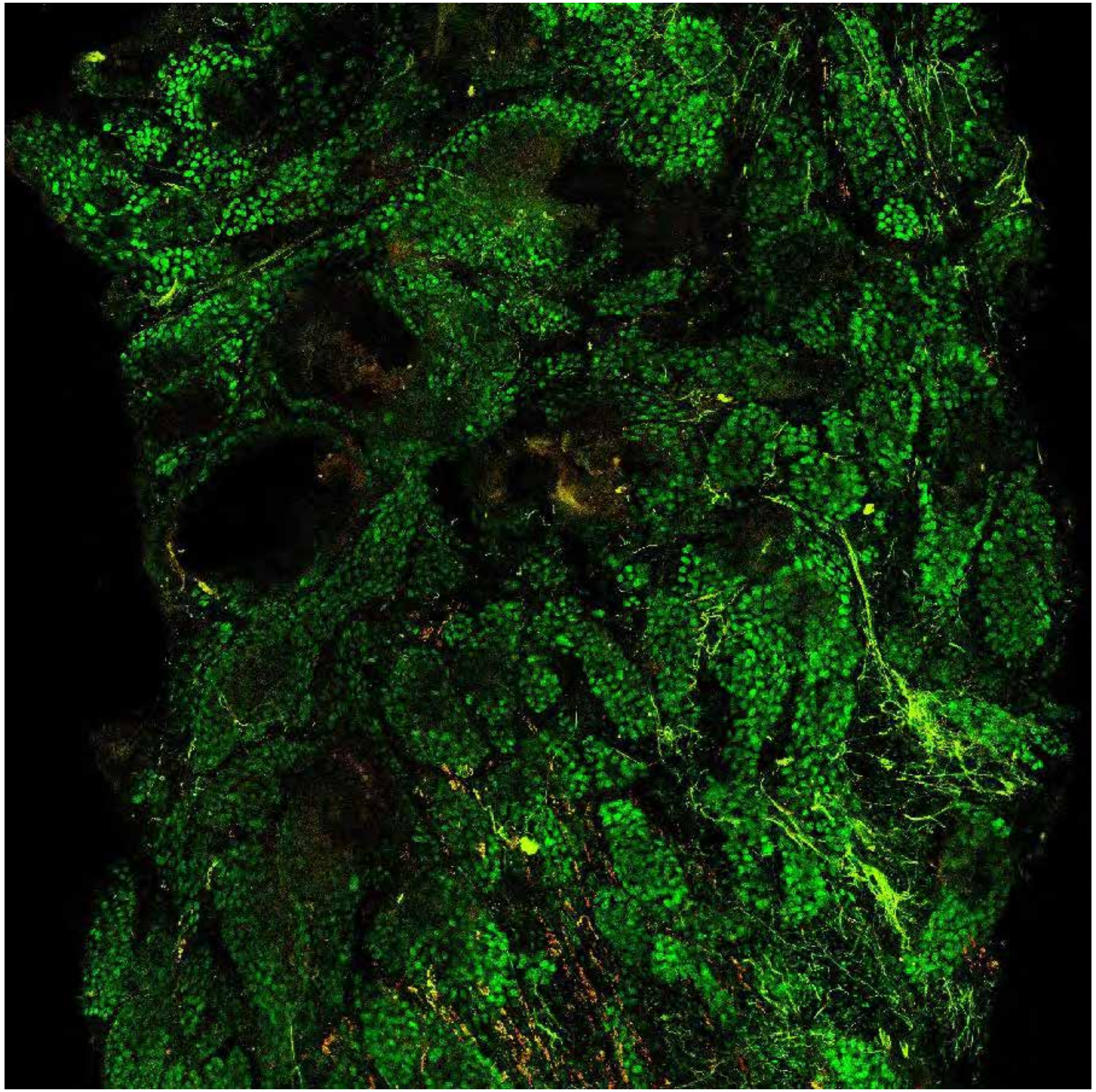

PARPi-FL (green) + Autofluorescence (red/yellow)

5347

### Close-up

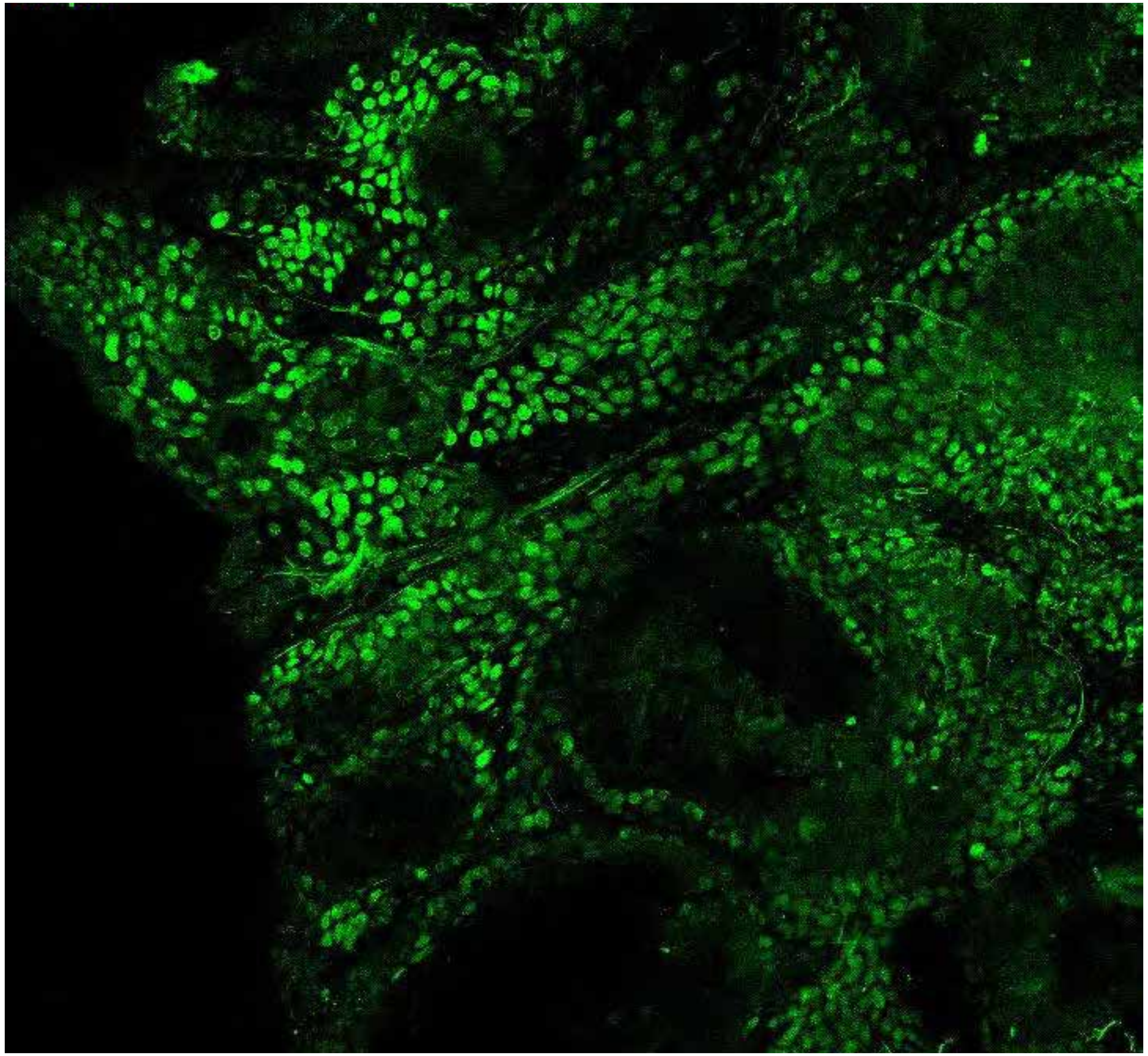

**PARPi-FL only (green)**

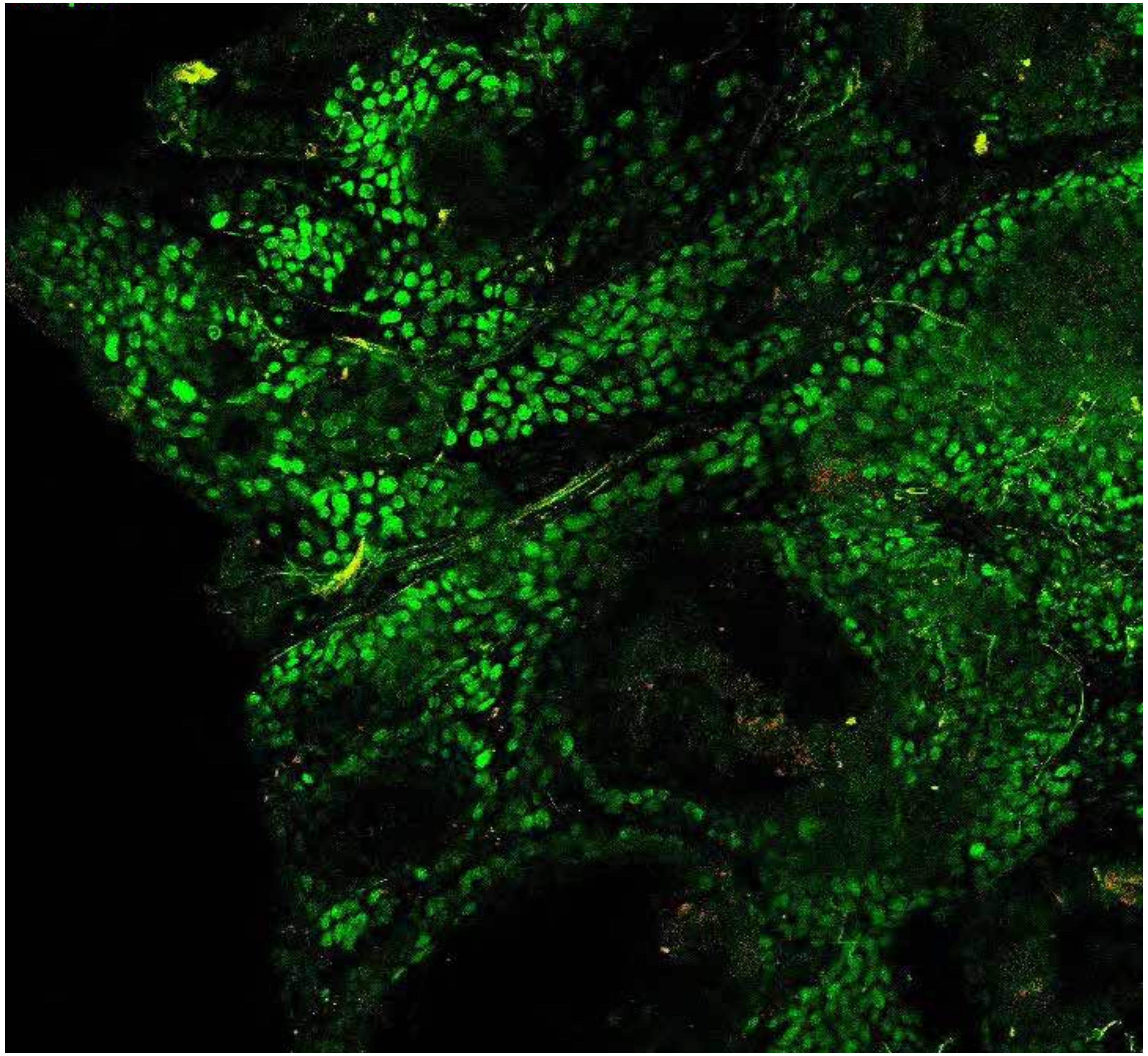

**PARPi-FL (green) + Autofluorescence (red/yellow)**

5347

### Close-up

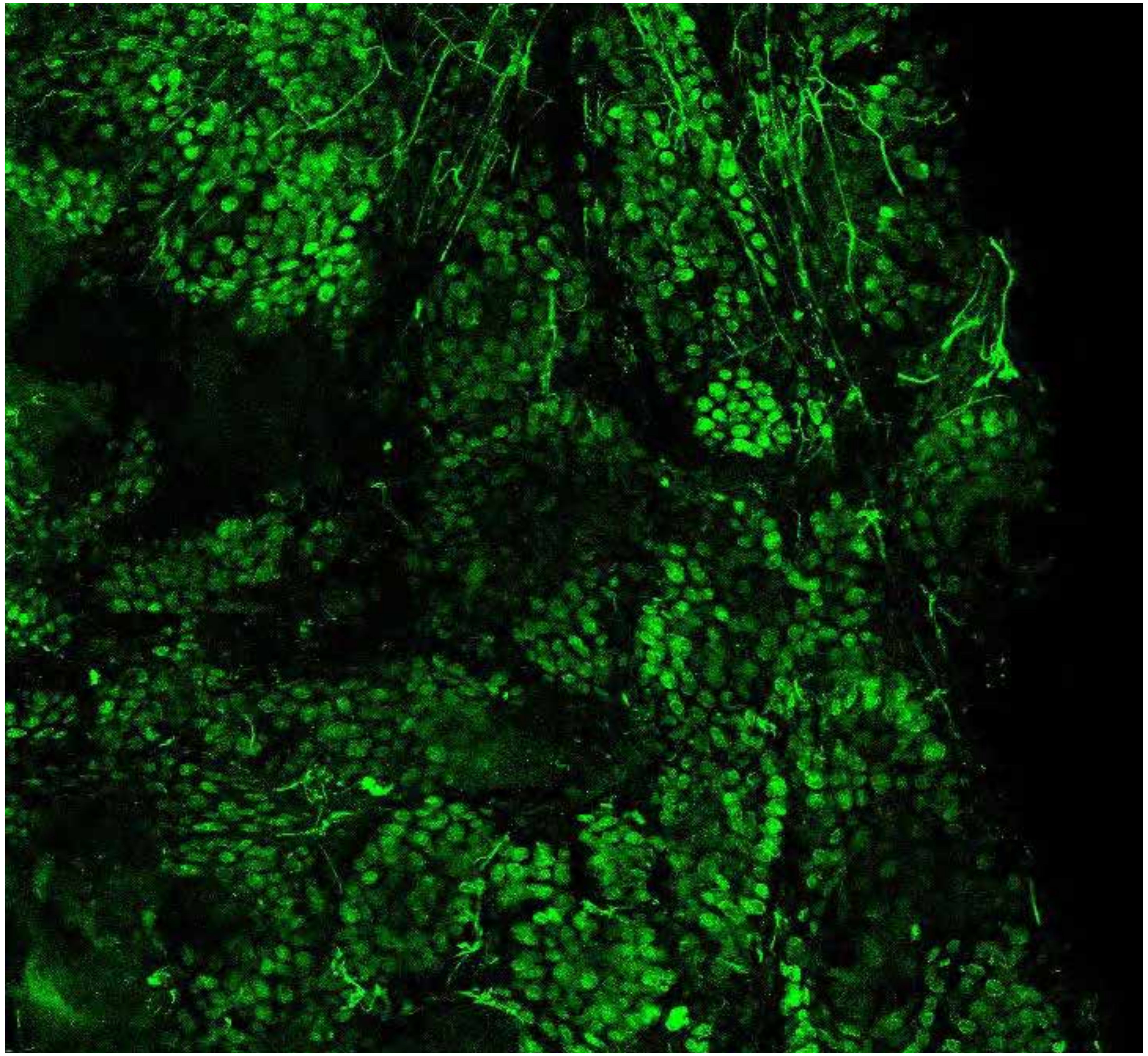

PARPi-FL only (green)

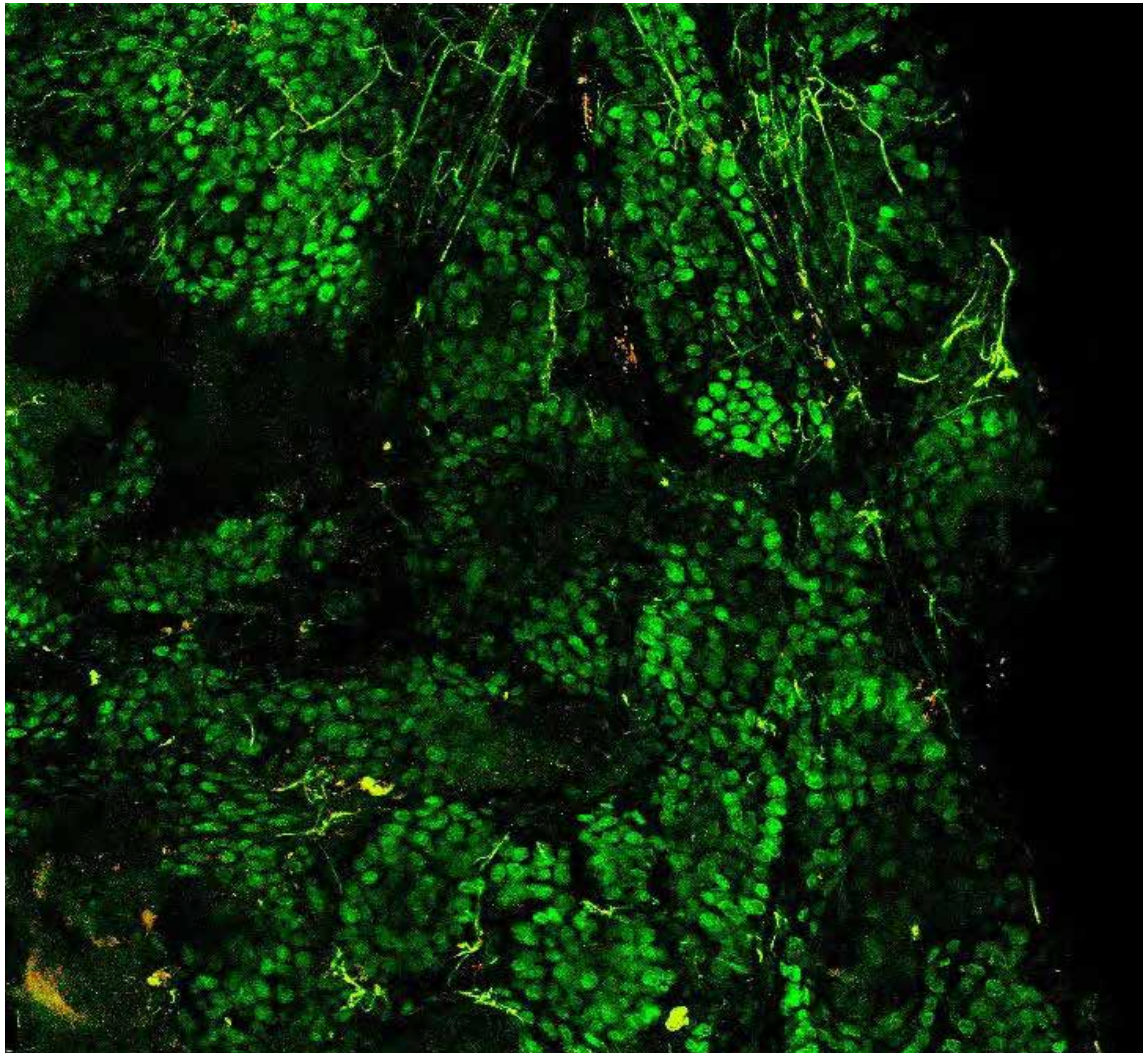

PARPi-FL (green) + Autofluorescence (red/yellow)

5347

### Close-up

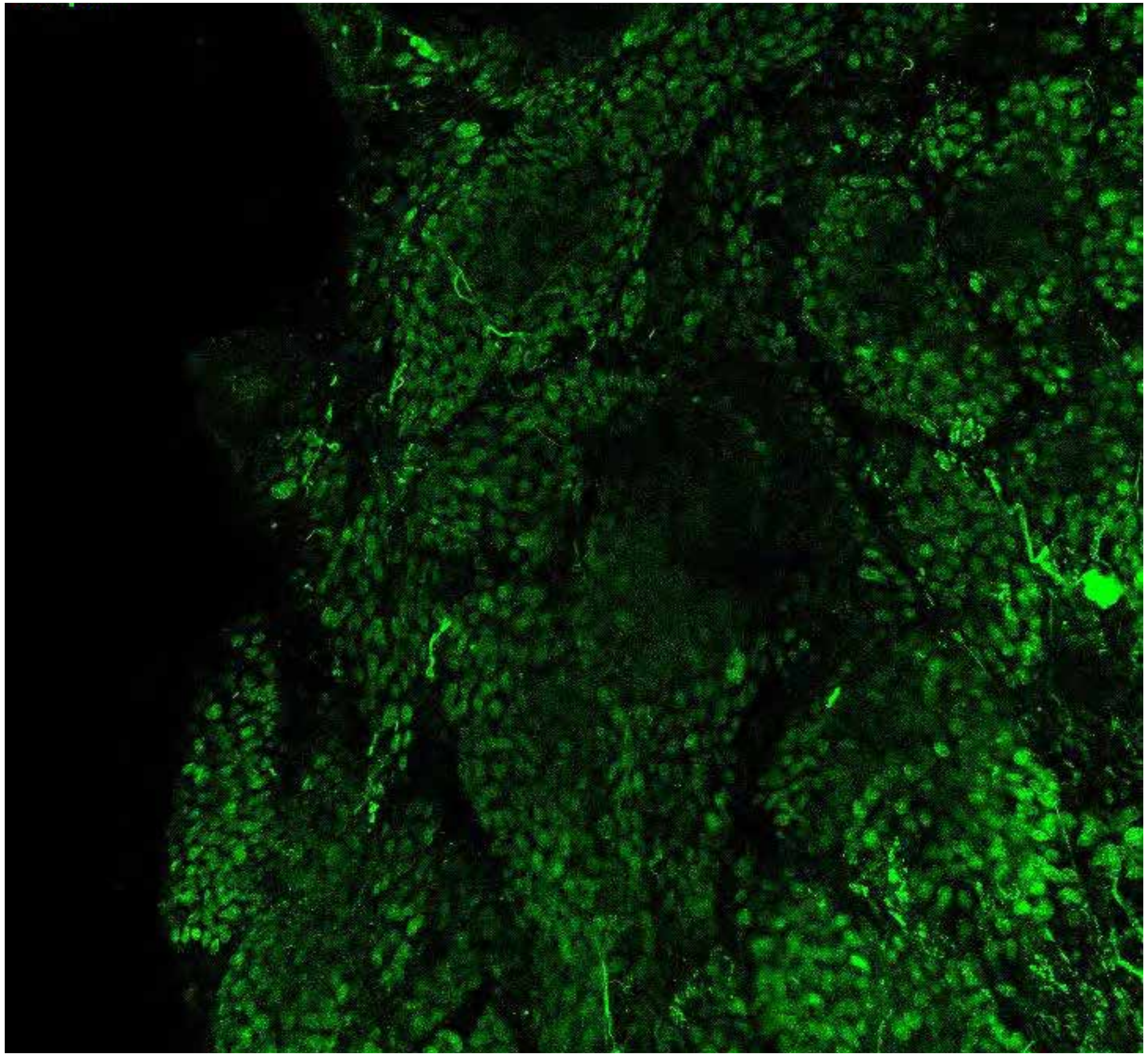

PARPi-FL only (green)

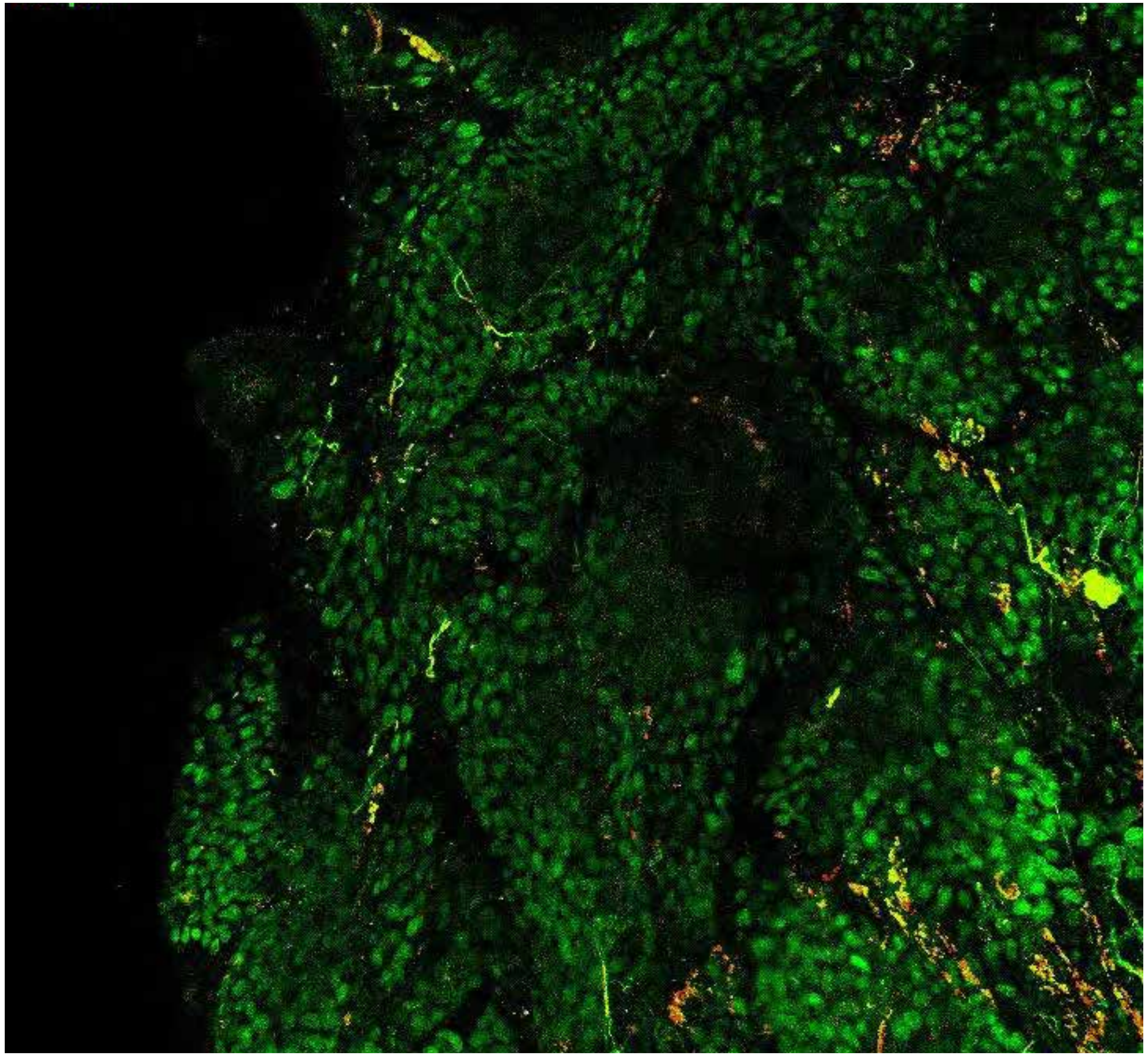

PARPi-FL (green) + Autofluorescence (red/yellow)

5347

### Close-up

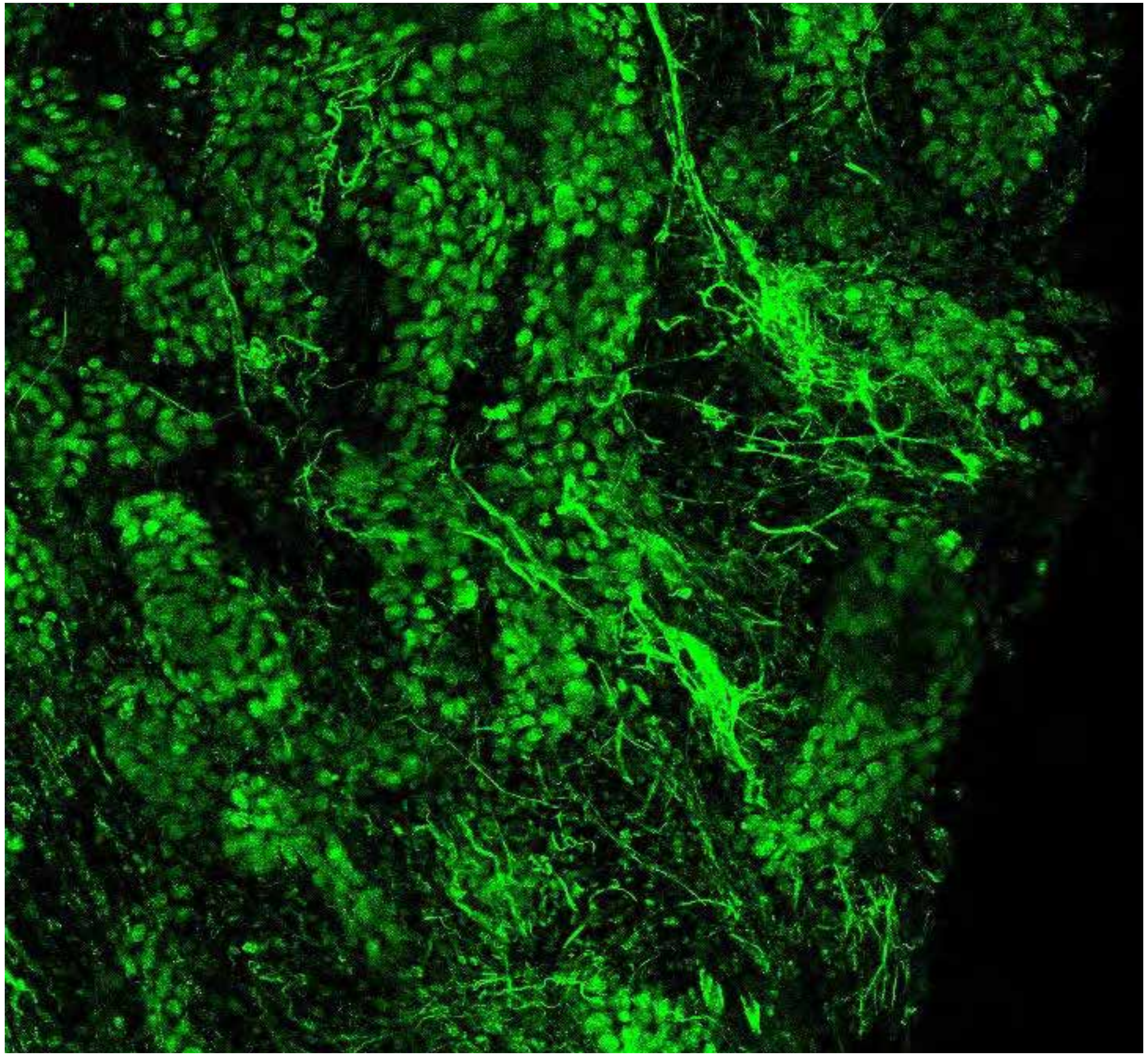

PARPi-FL only (green)

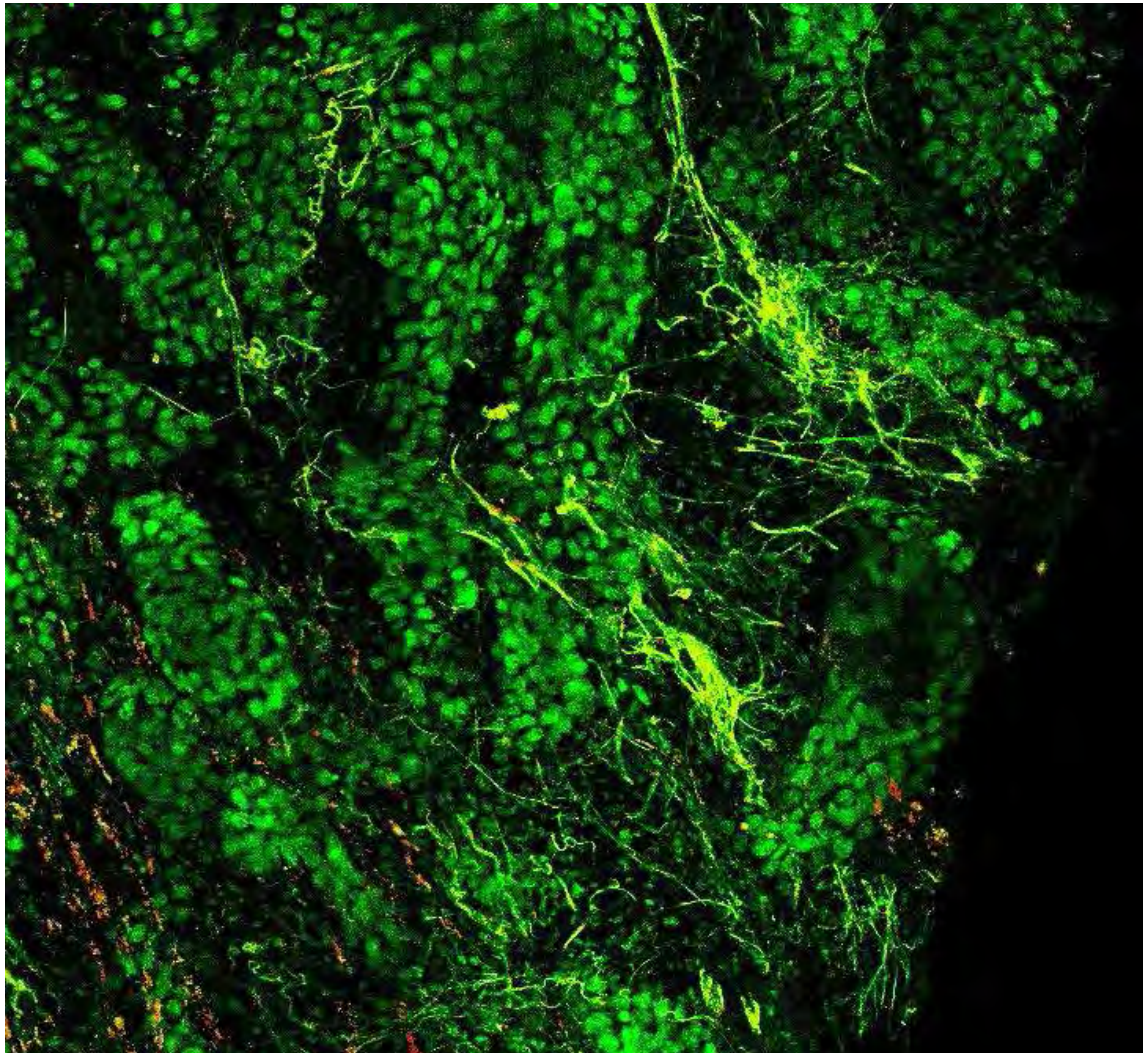

PARPi-FL (green) + Autofluorescence (red/yellow)

5347



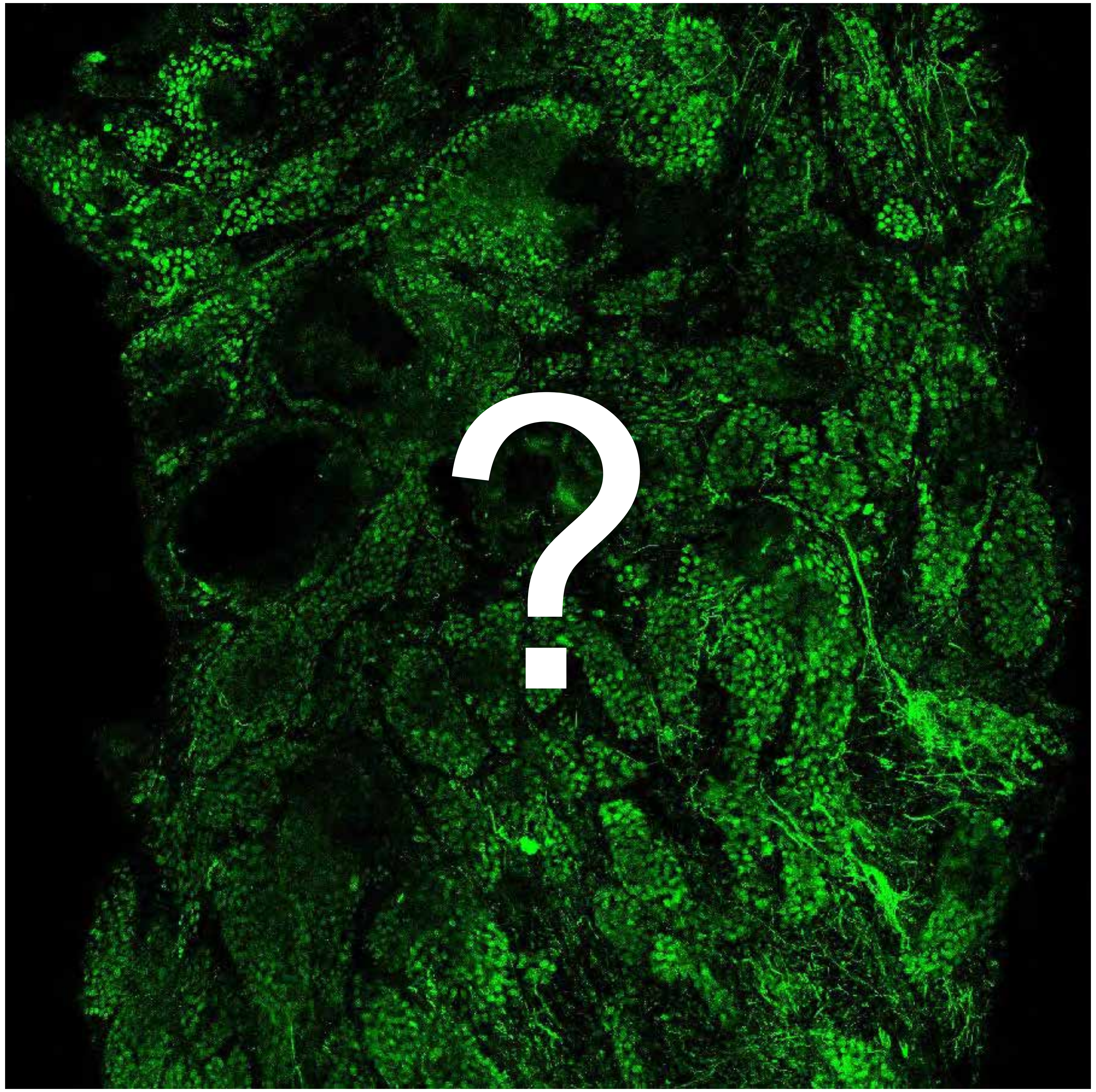


Tumor

Margin

5489

10

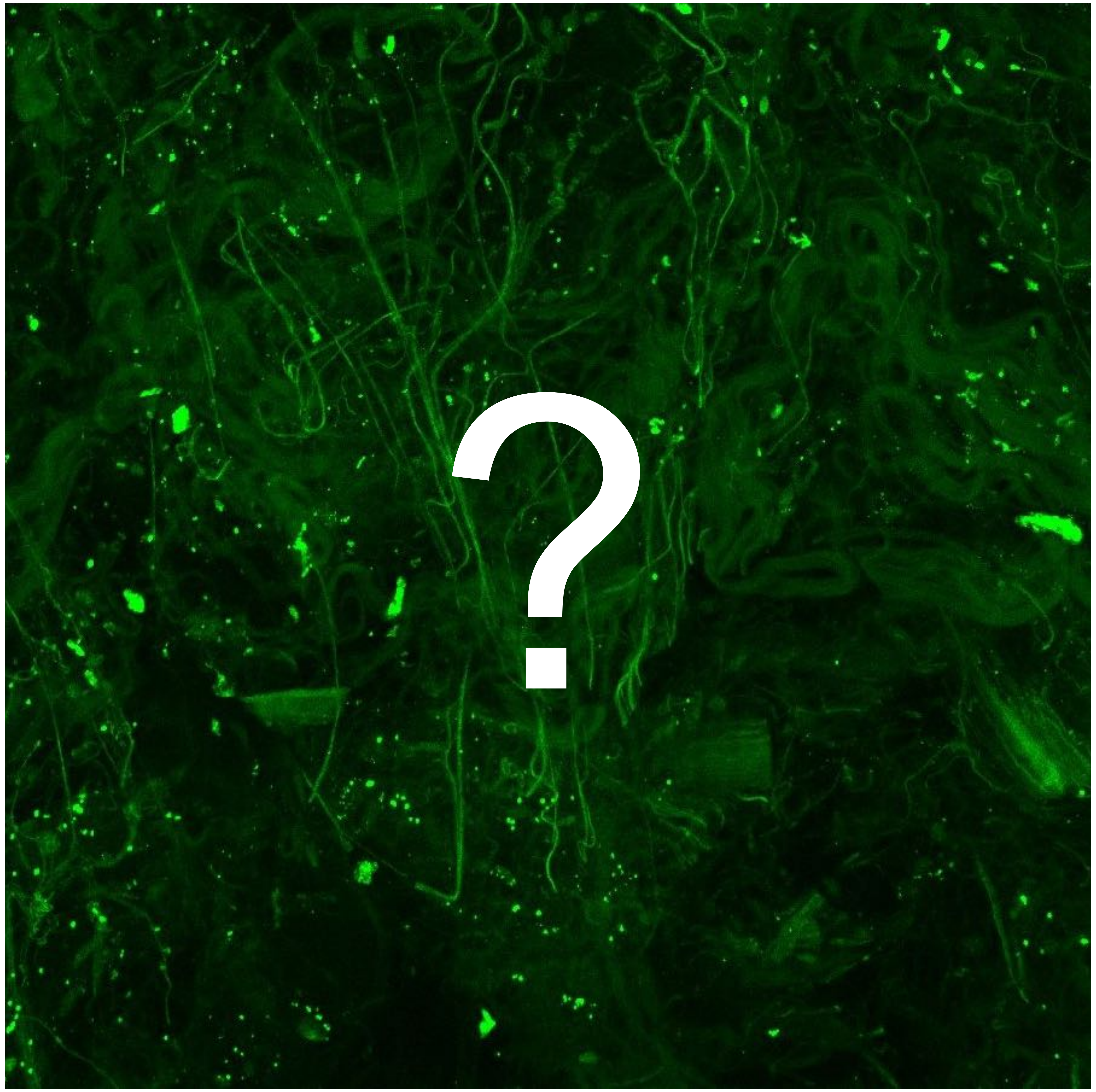


Tumor

Margin

6076

### Overview

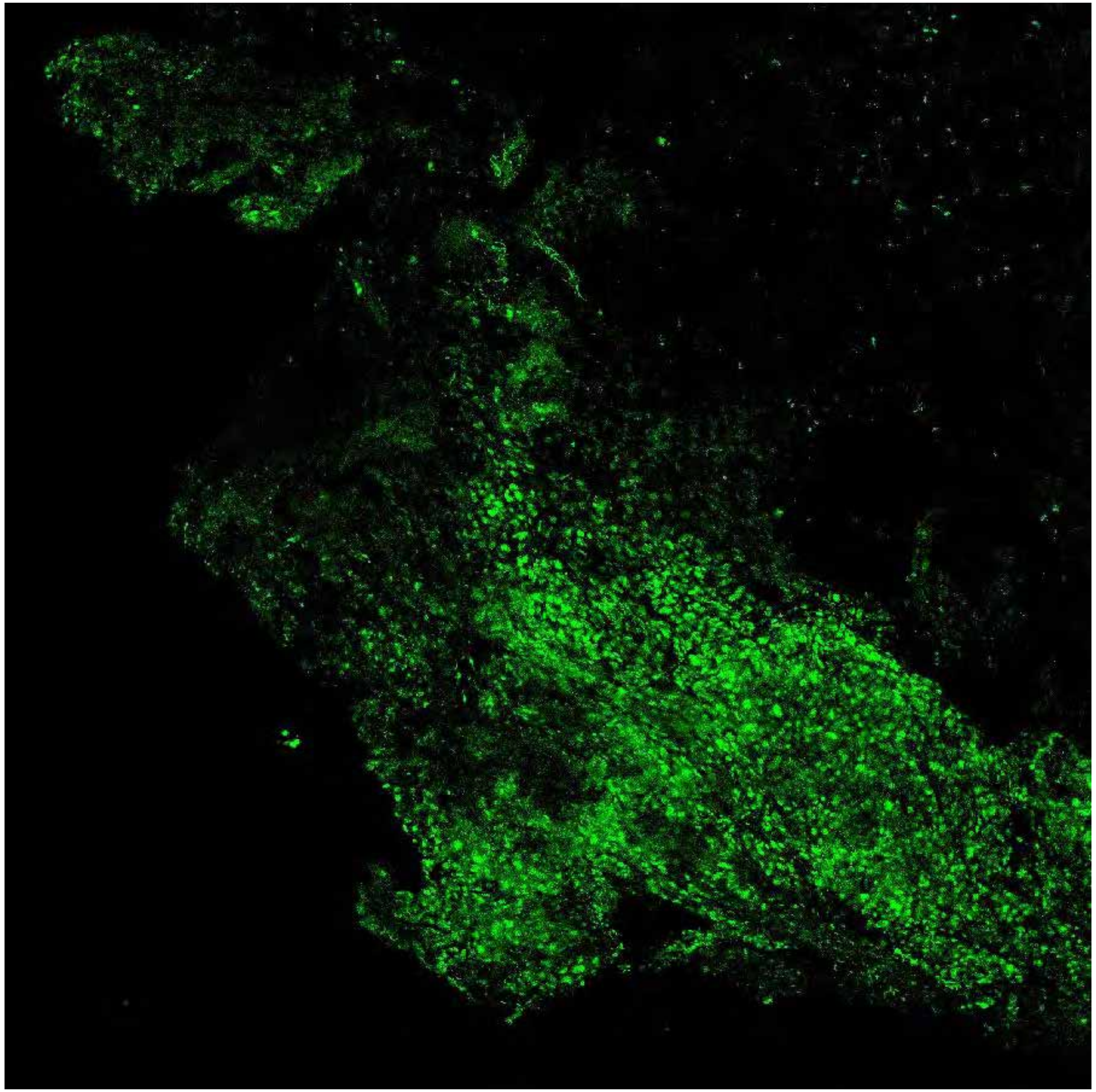

PARPi-FL only (green)

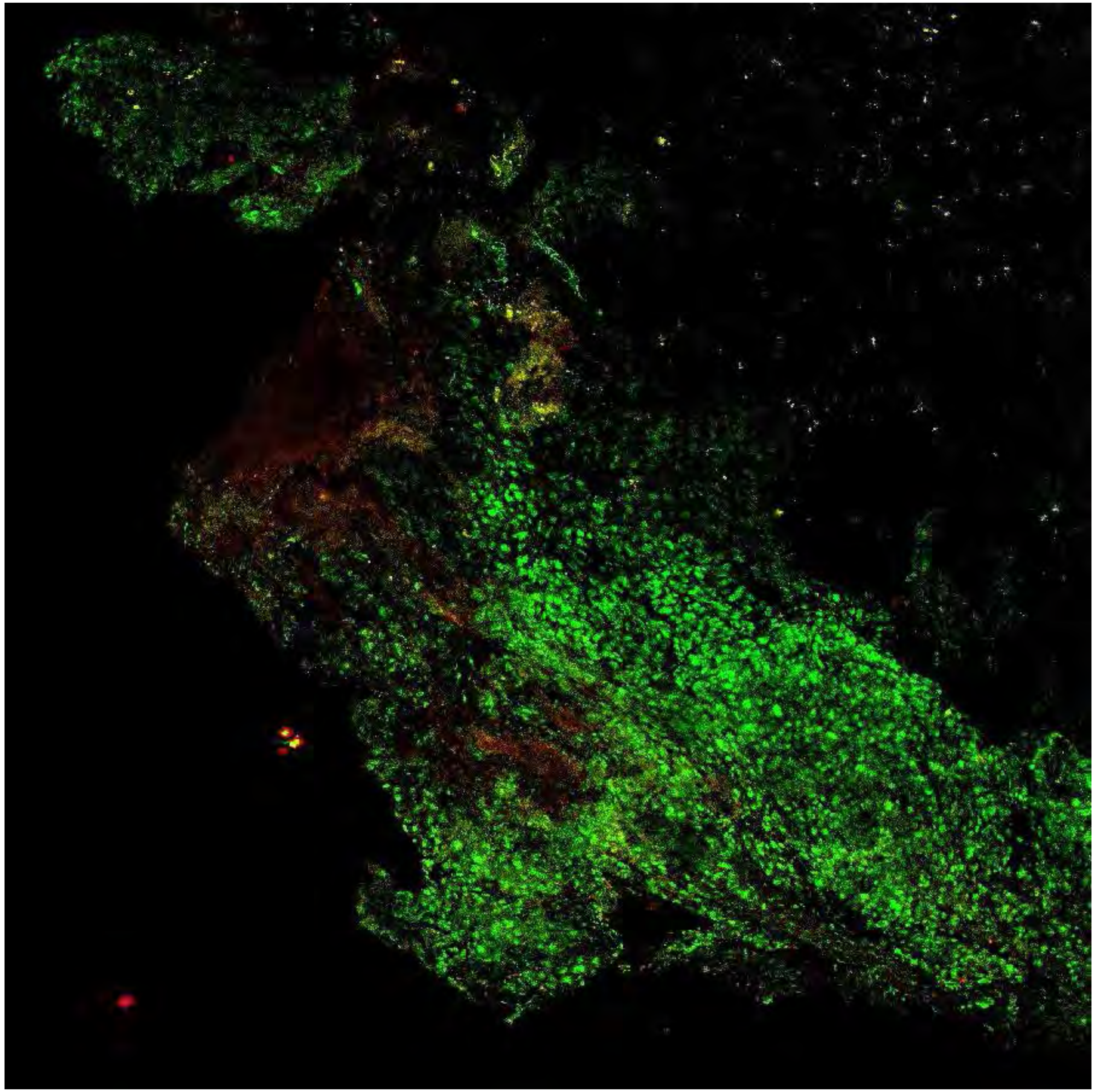

PARPi-FL (green) + Autofluorescence (red/yellow)

6076

### Close-up

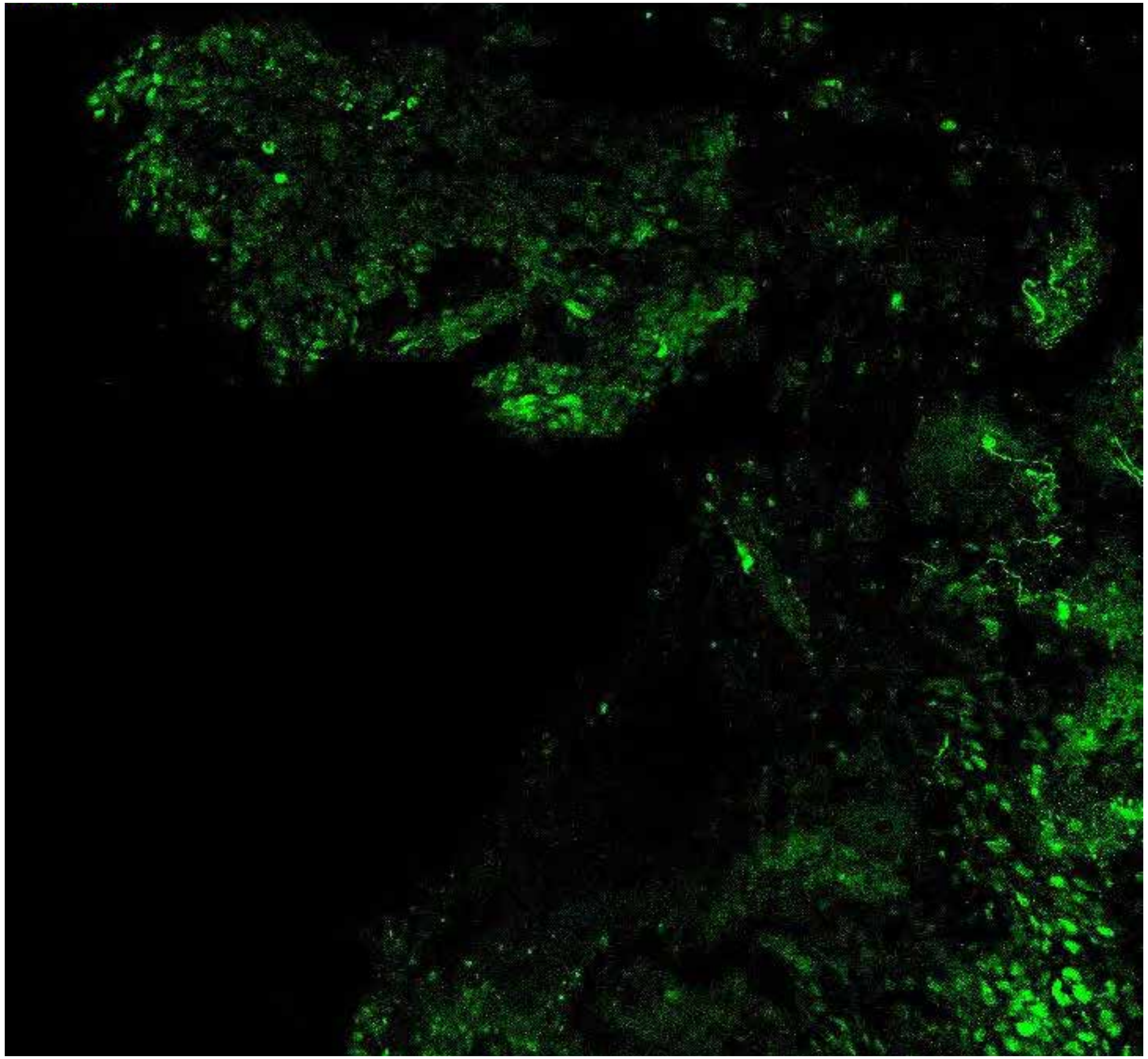

PARPi-FL only (green)

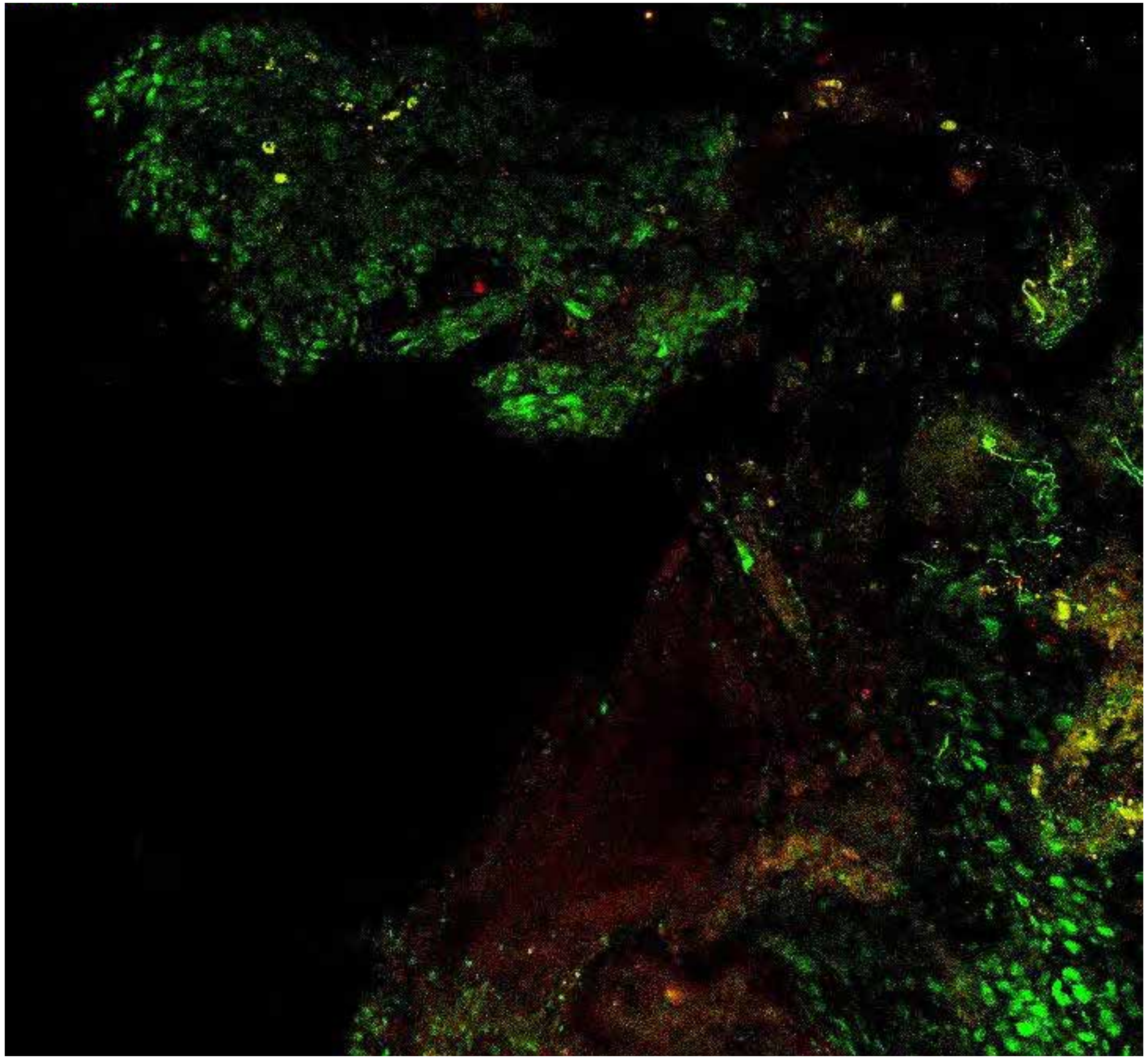

PARPi-FL (green) + Autofluorescence (red/yellow)

6076

### Close-up

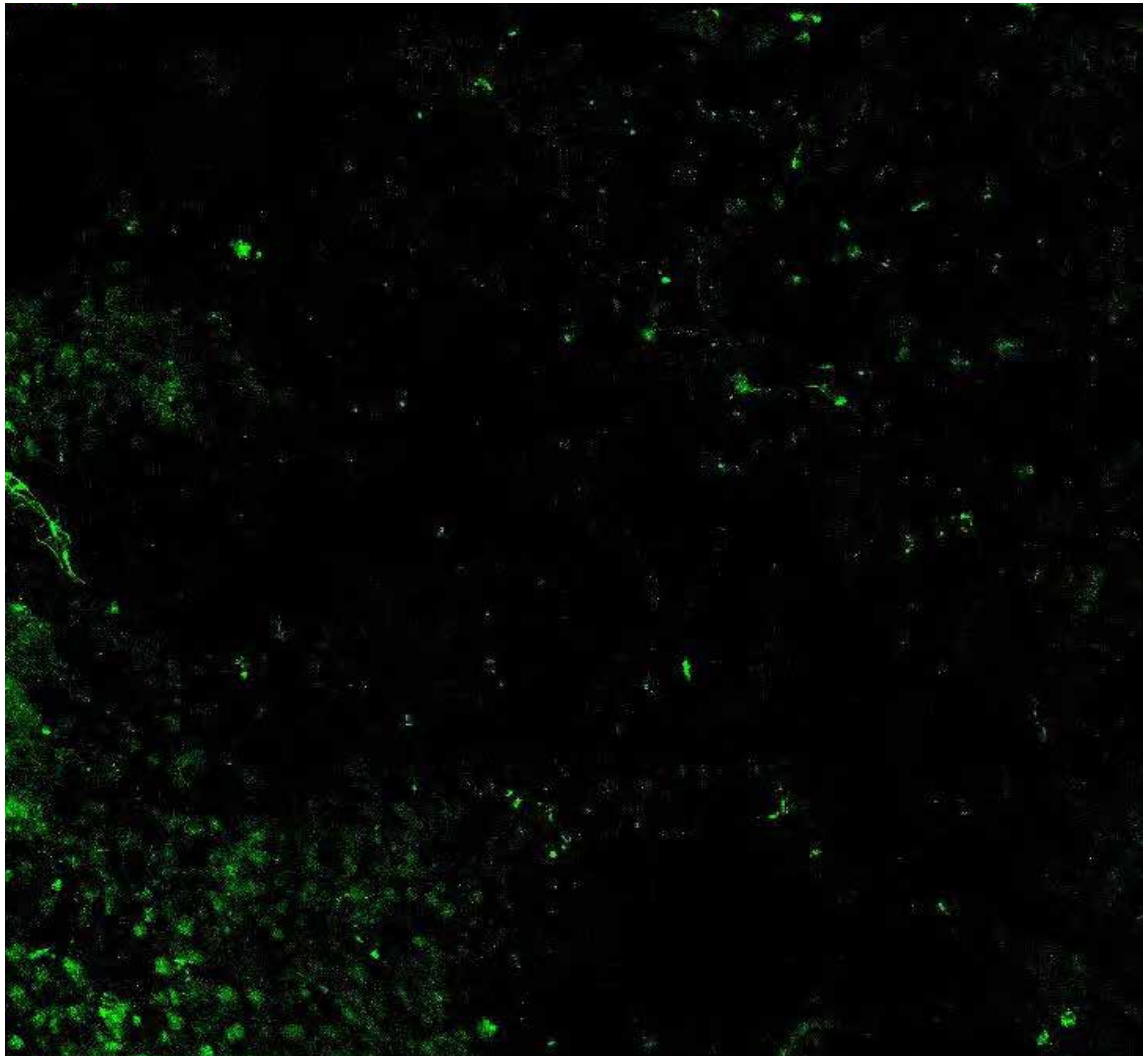

**PARPi-FL only (green)**

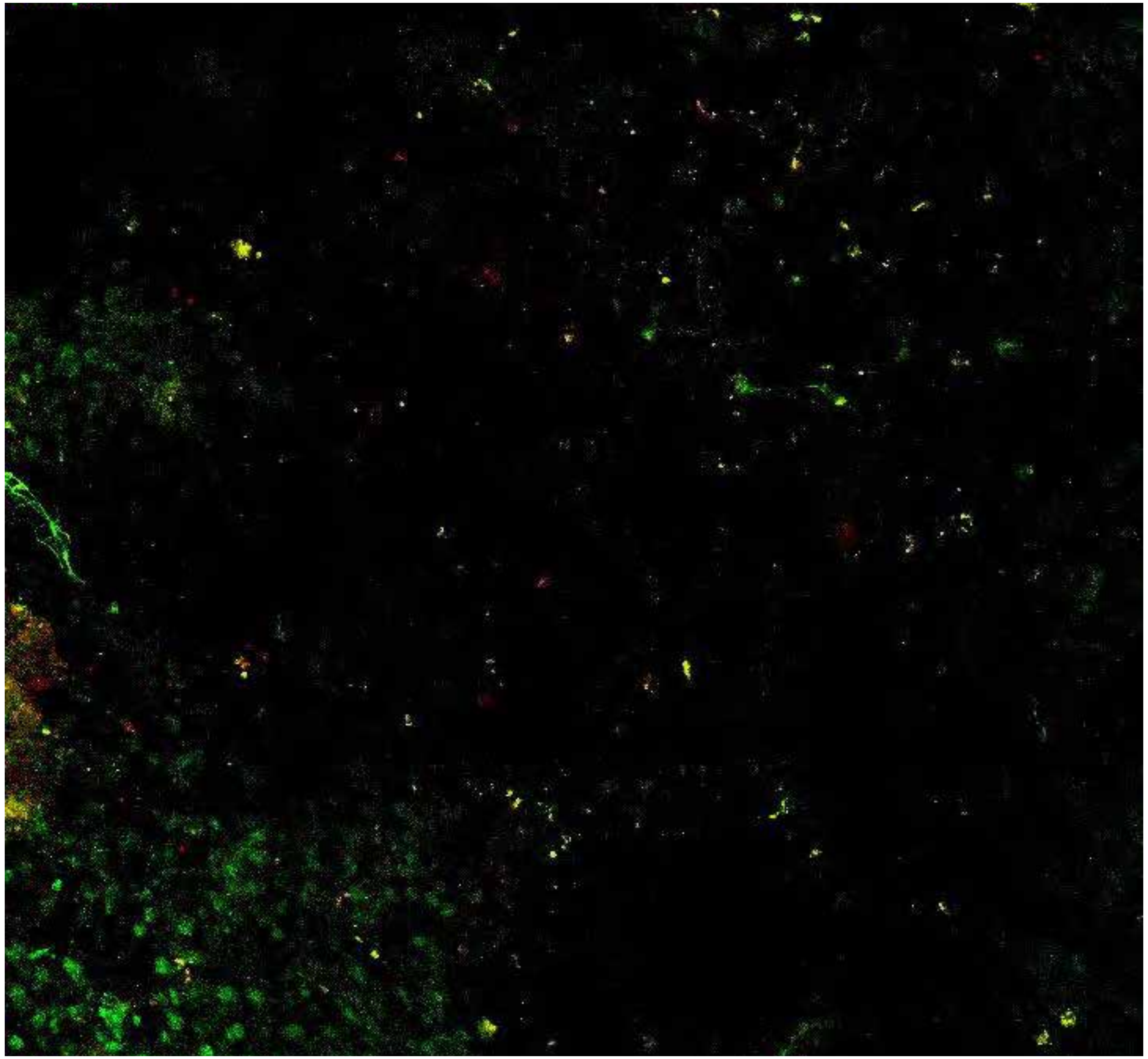

**PARPi-FL (green) + Autofluorescence (red/yellow)**

6076

### Close-up

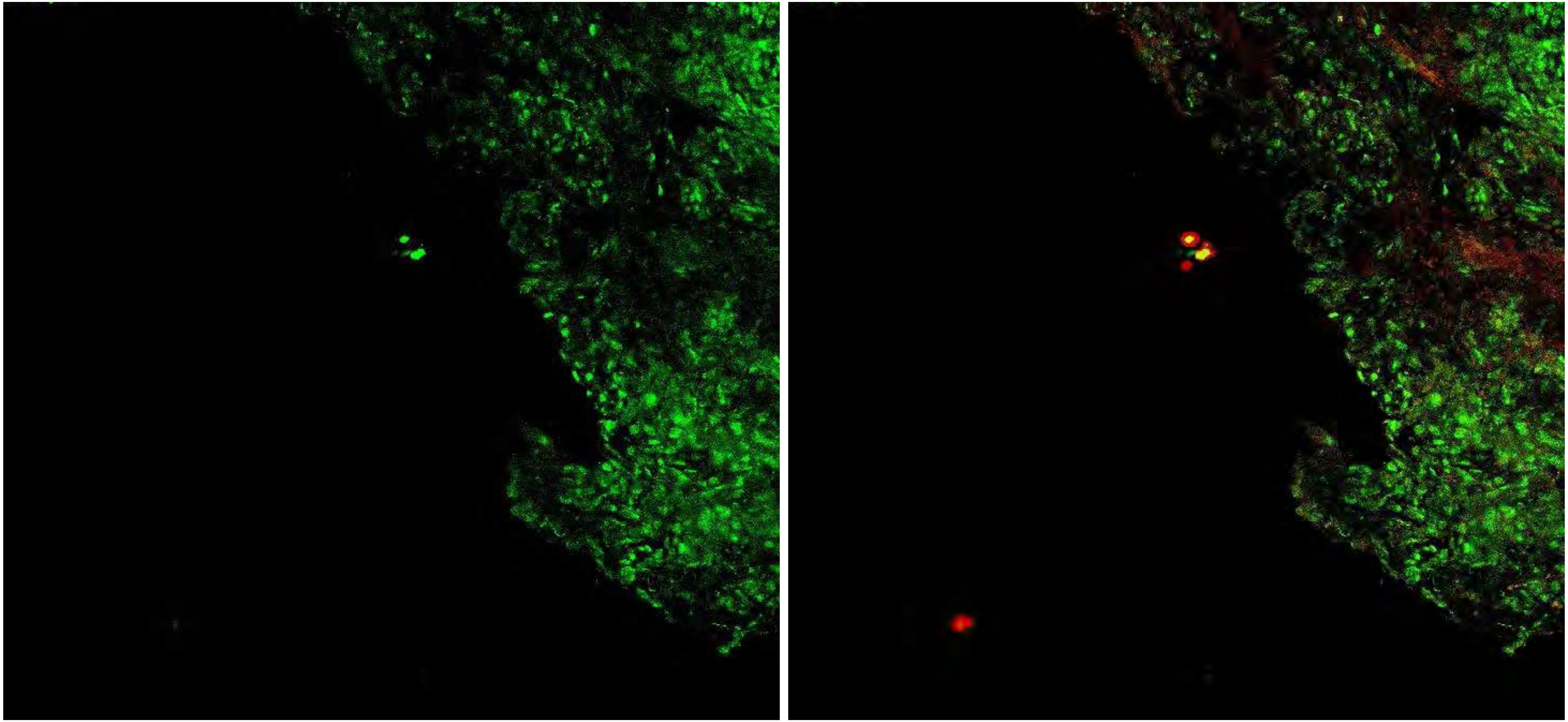

PARPi-FL only (green)    PARPi-FL (green) + Autofluorescence (red/yellow)

6076

### Close-up

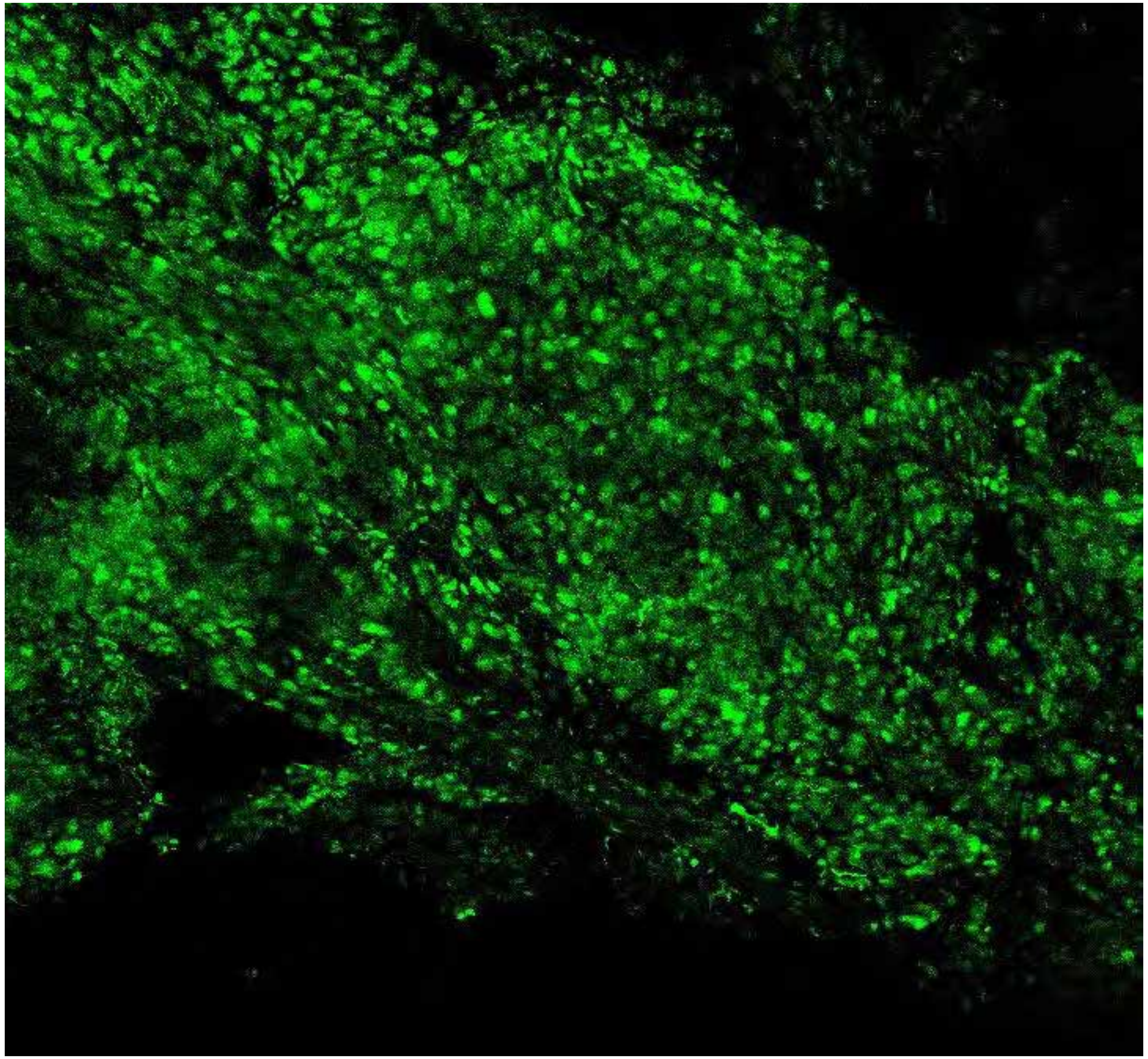

PARPi-FL only (green)

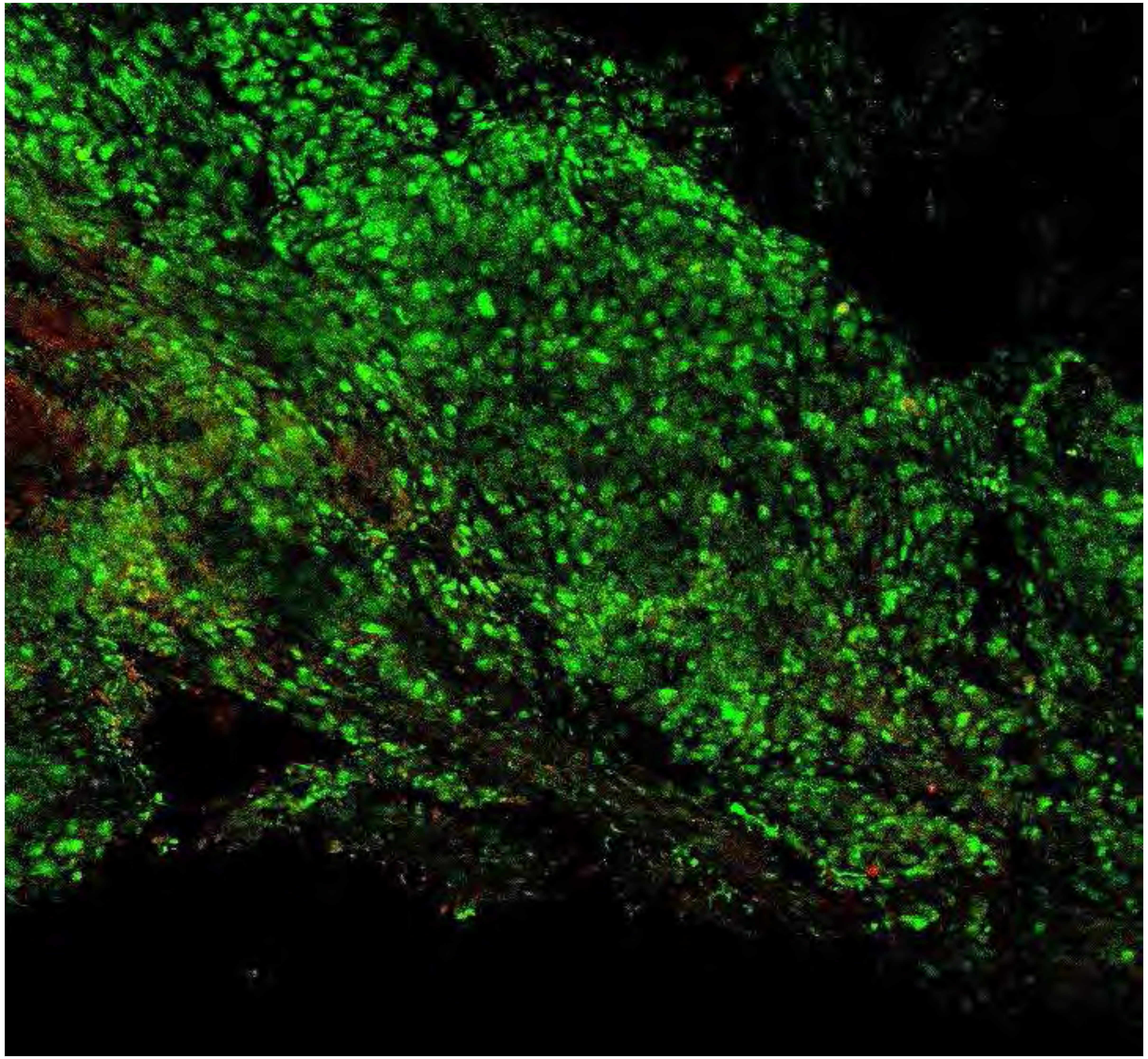

PARPi-FL (green) + Autofluorescence (red/yellow)

6076



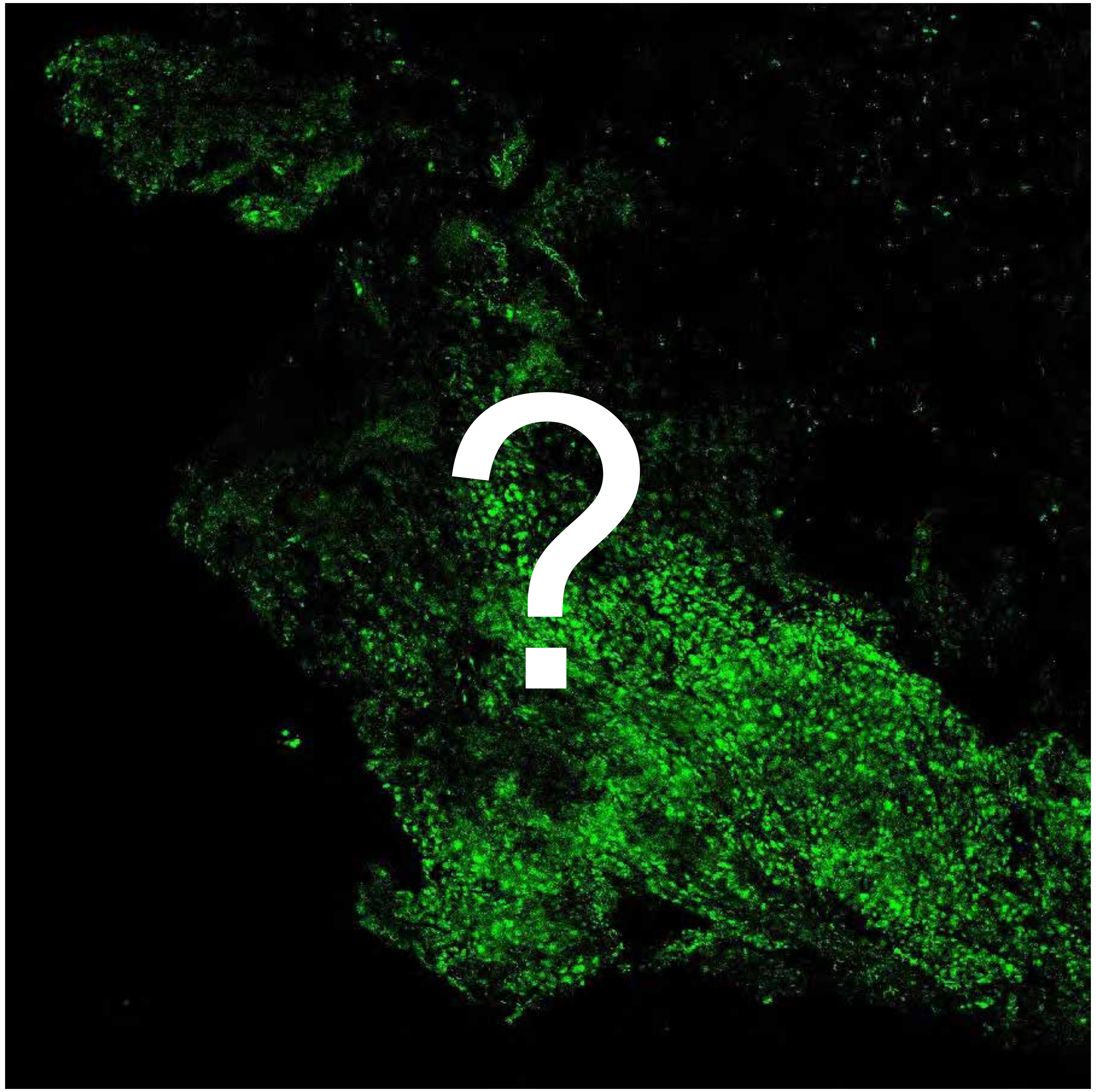


Tumor

Margin

# 6344

### Overview

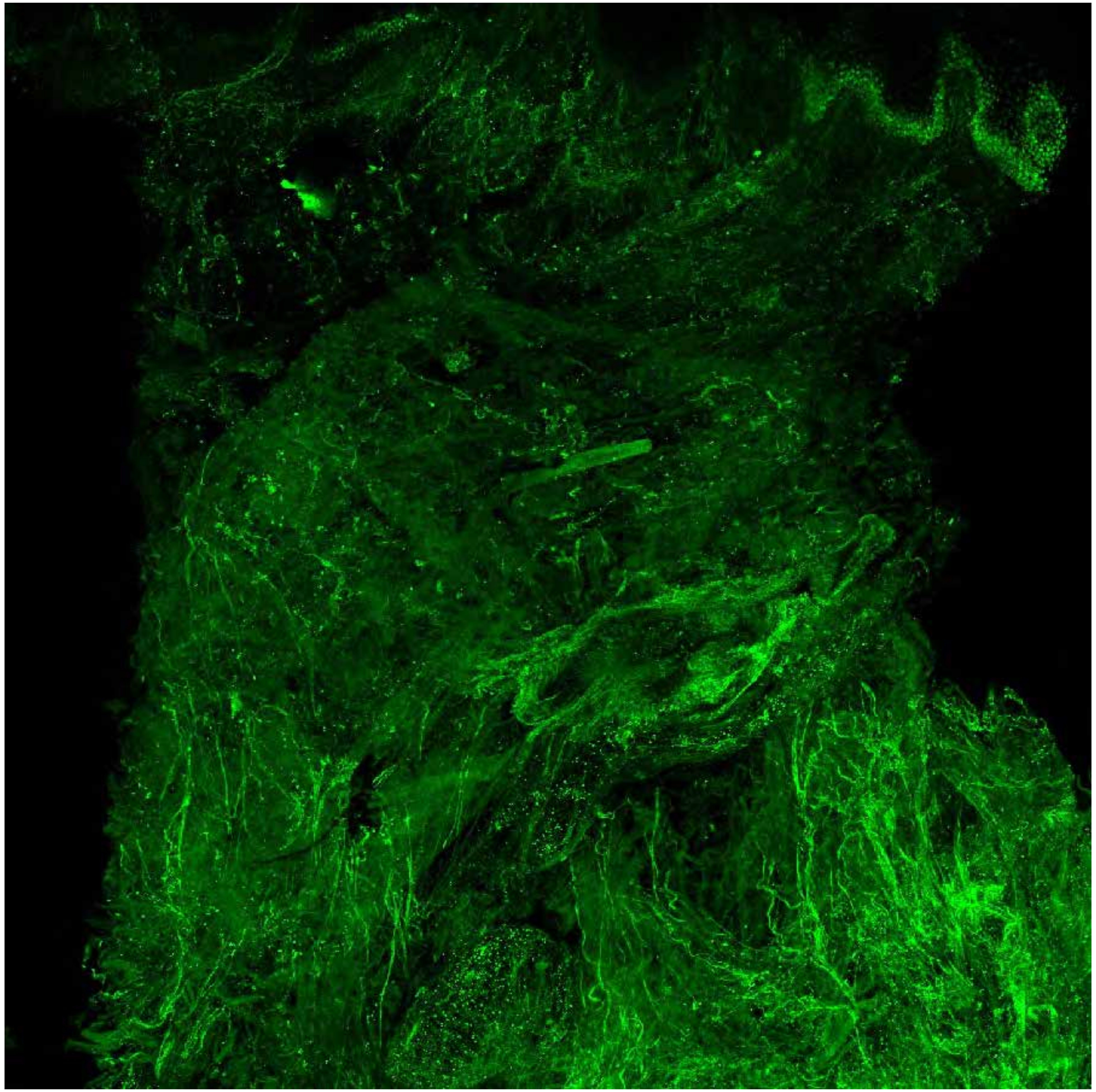

PARPi-FL only (green)

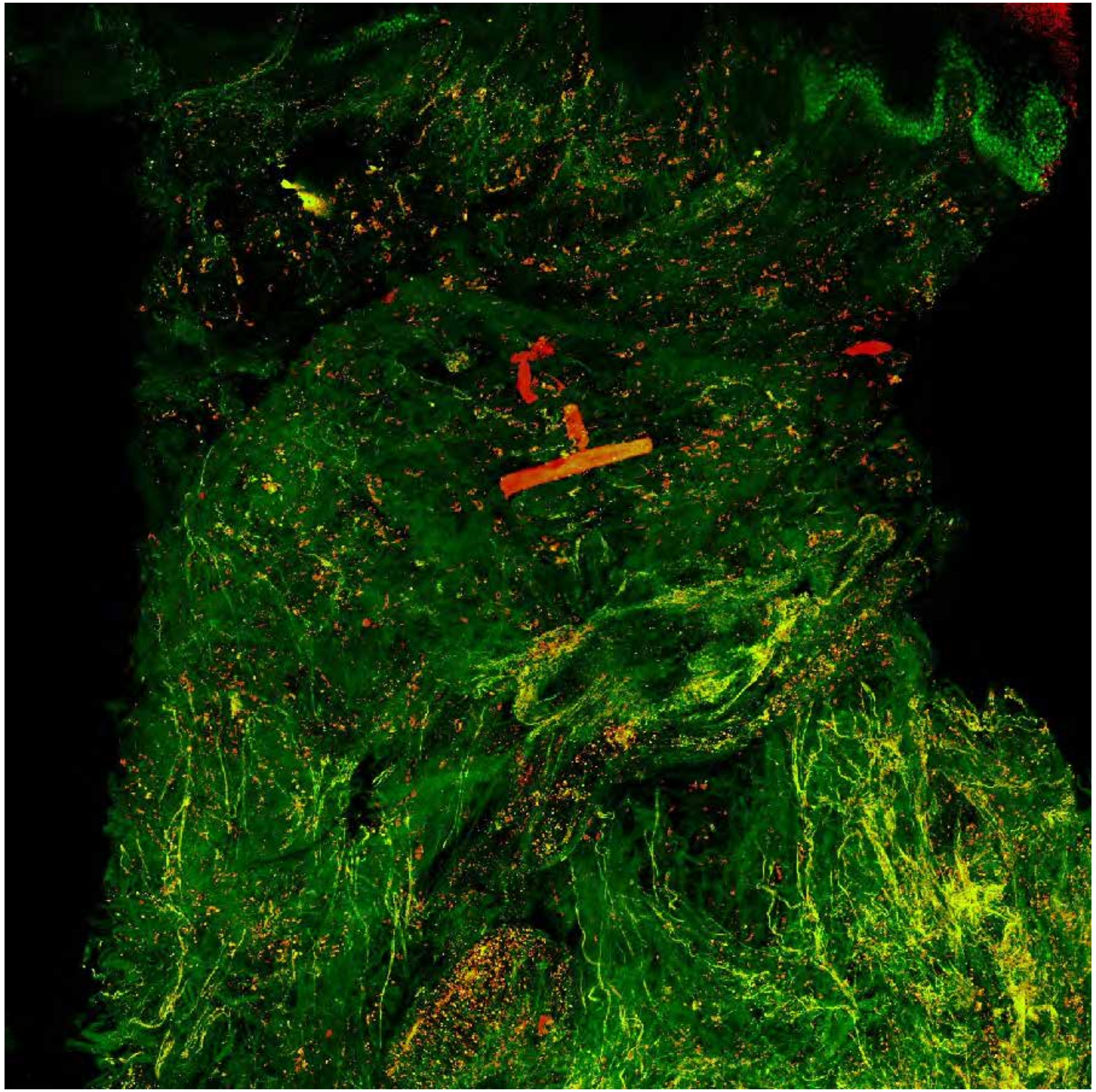

PARPi-FL (green) + Autofluorescence (red/yellow)

# 6344

### Close-up

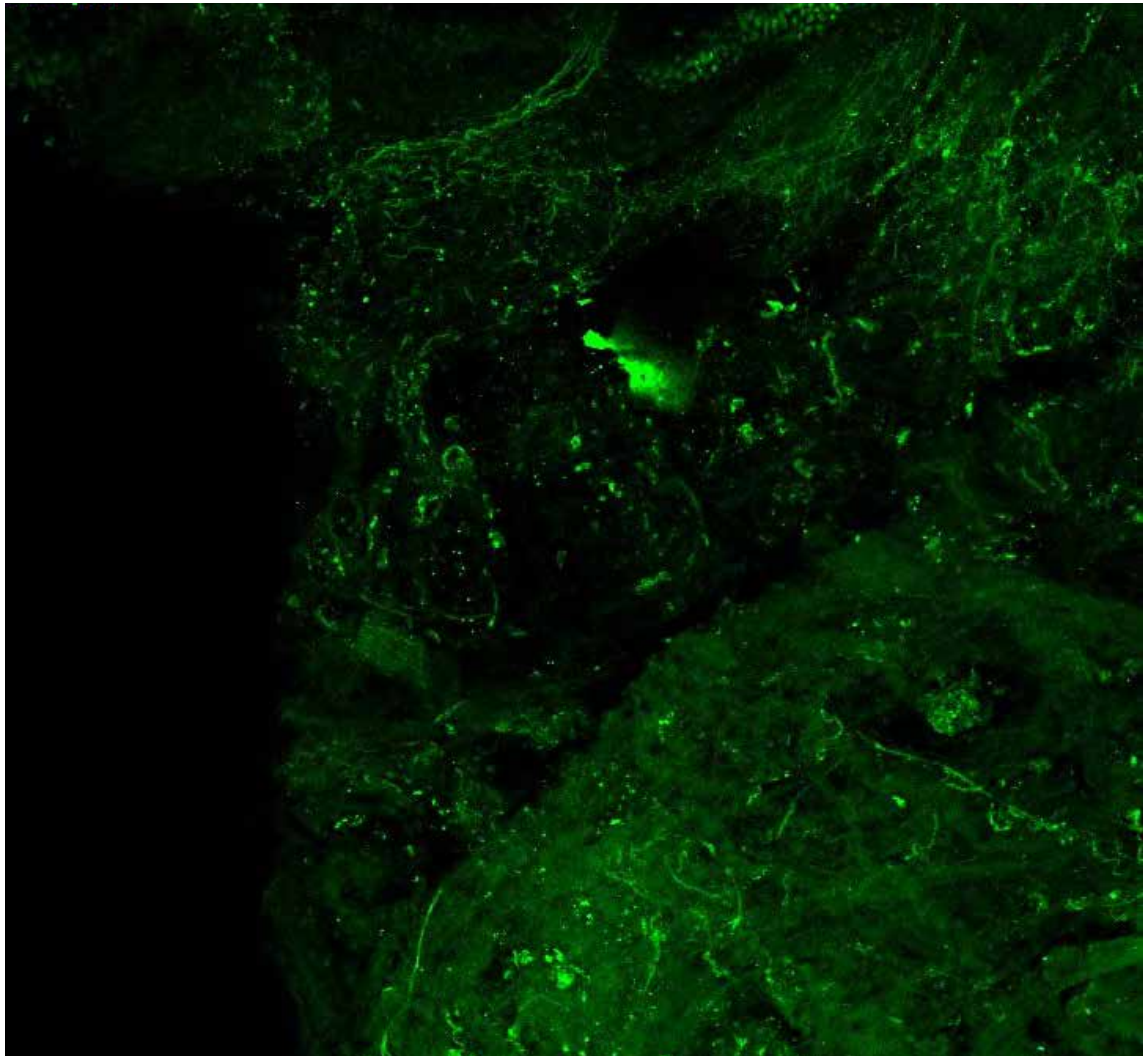

PARPi-FL only (green)

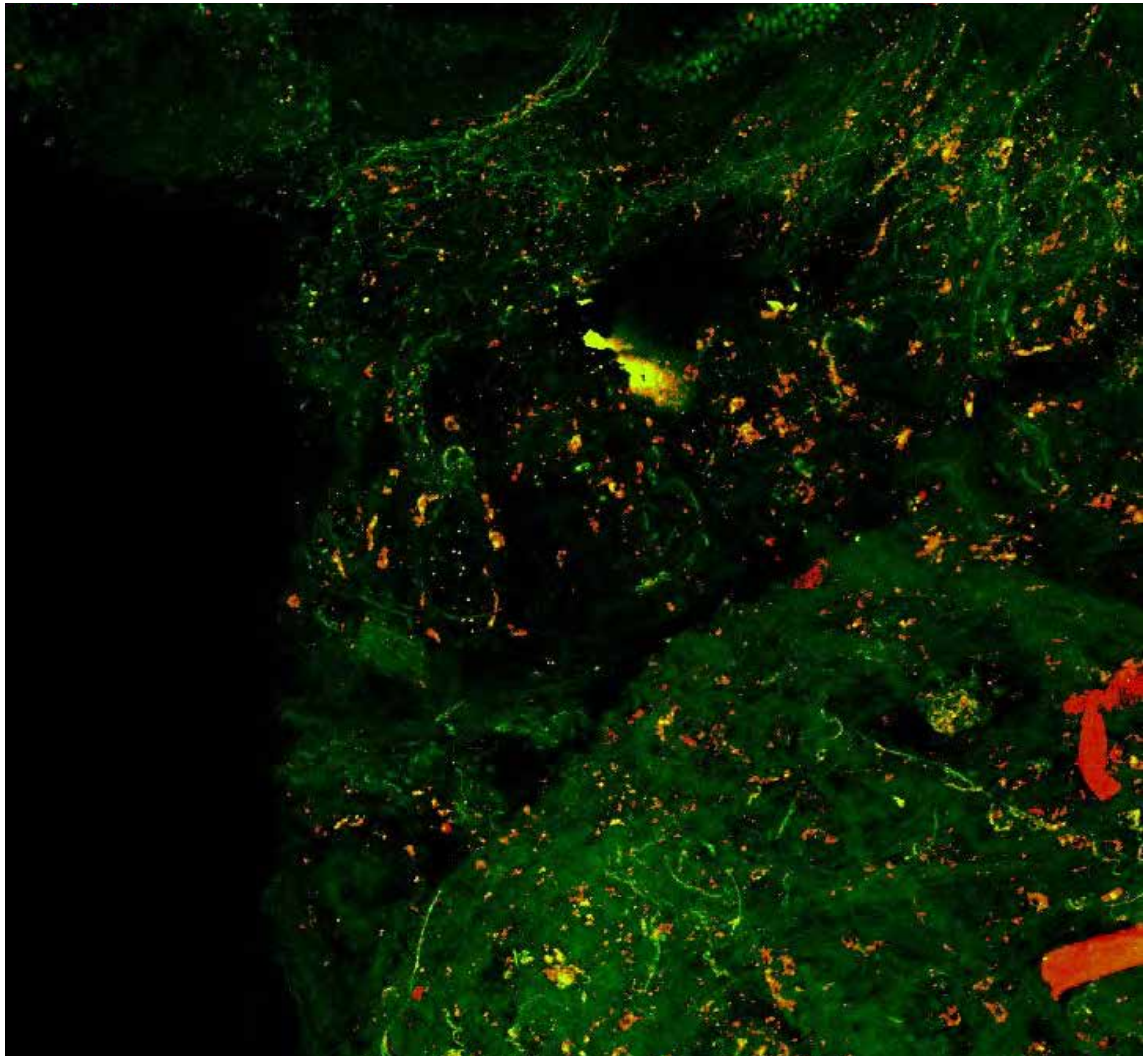

PARPi-FL (green) + Autofluorescence (red/yellow)

6344

### Close-up

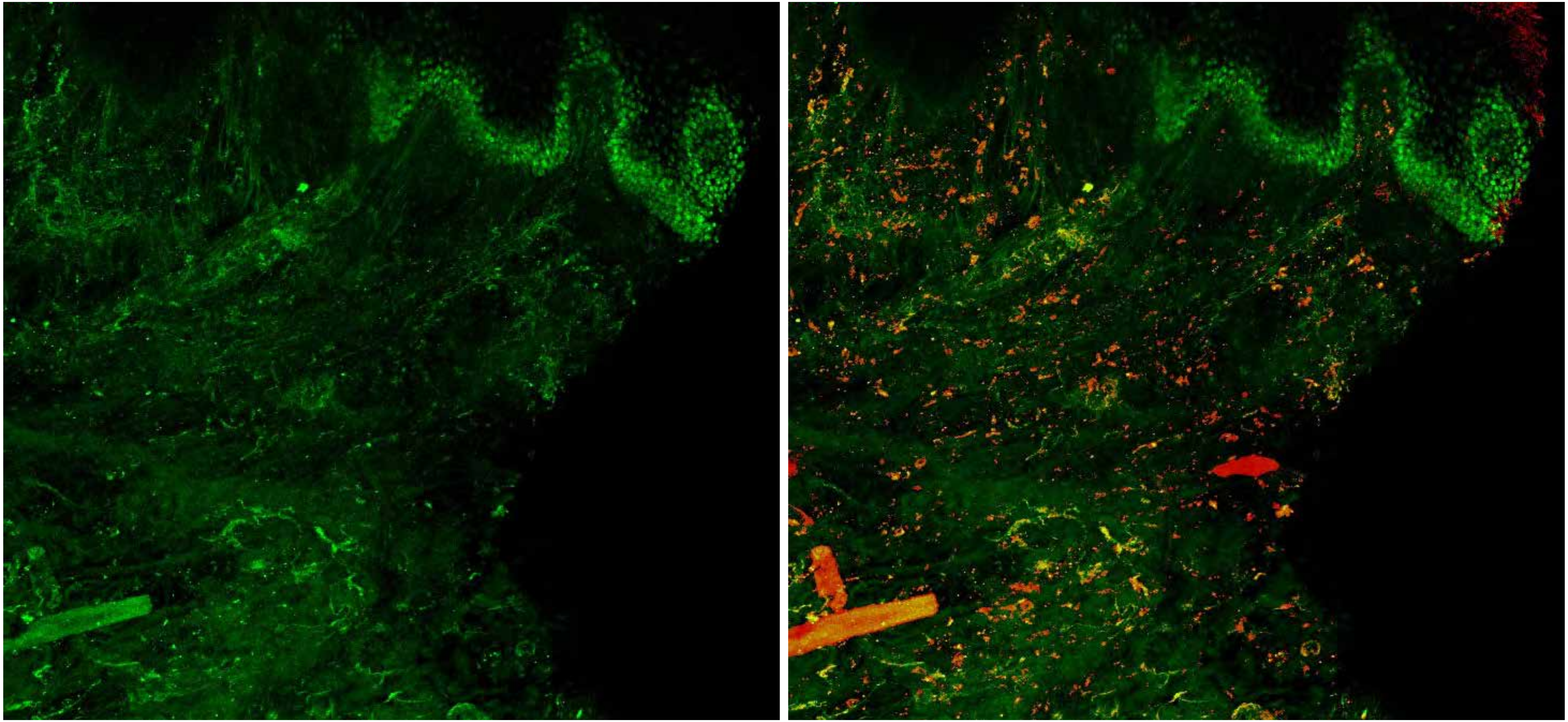

PARPi-FL only (green)

PARPi-FL (green) + Autofluorescence (red/yellow)

6344

### Close-up

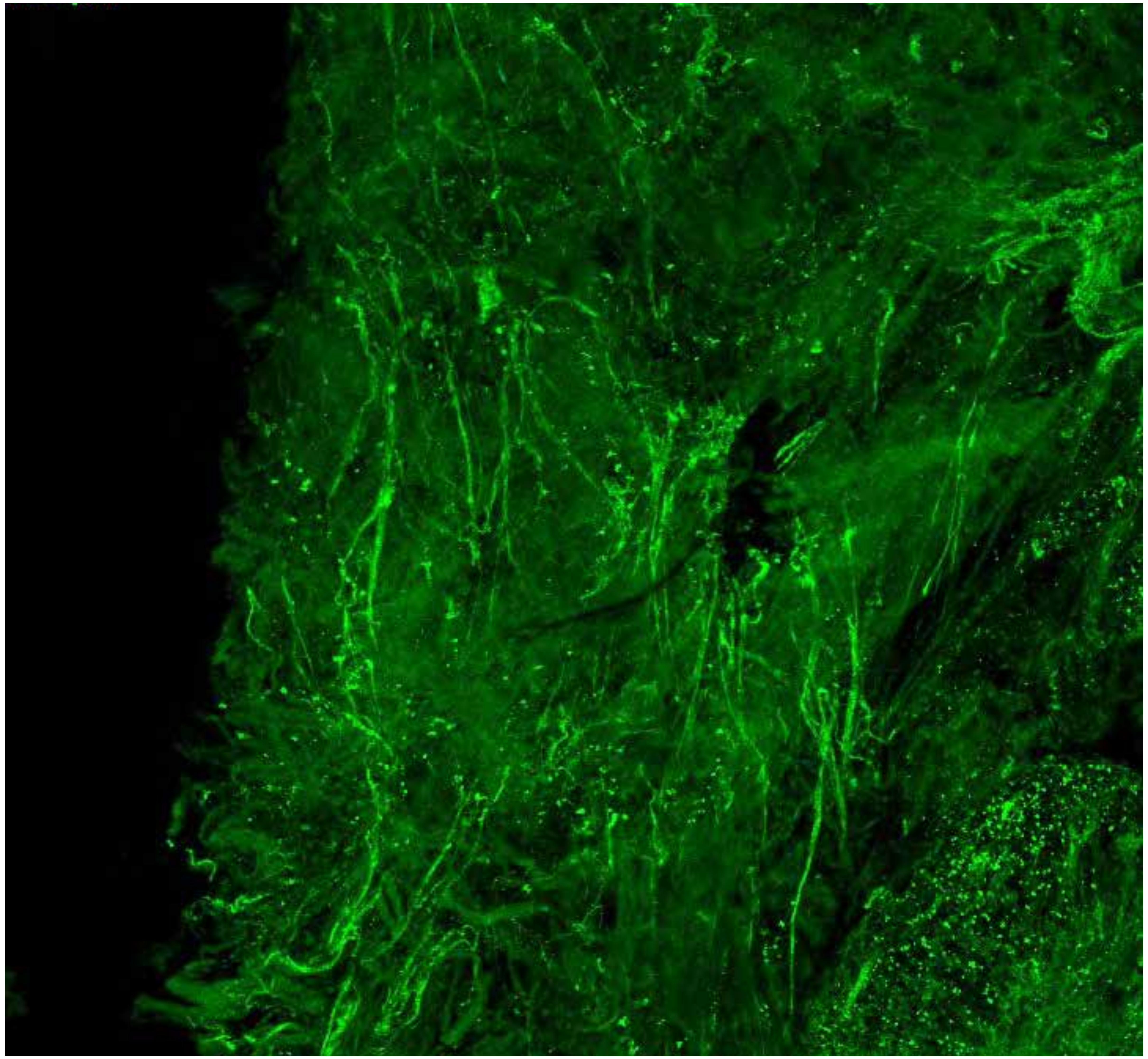

PARPi-FL only (green)

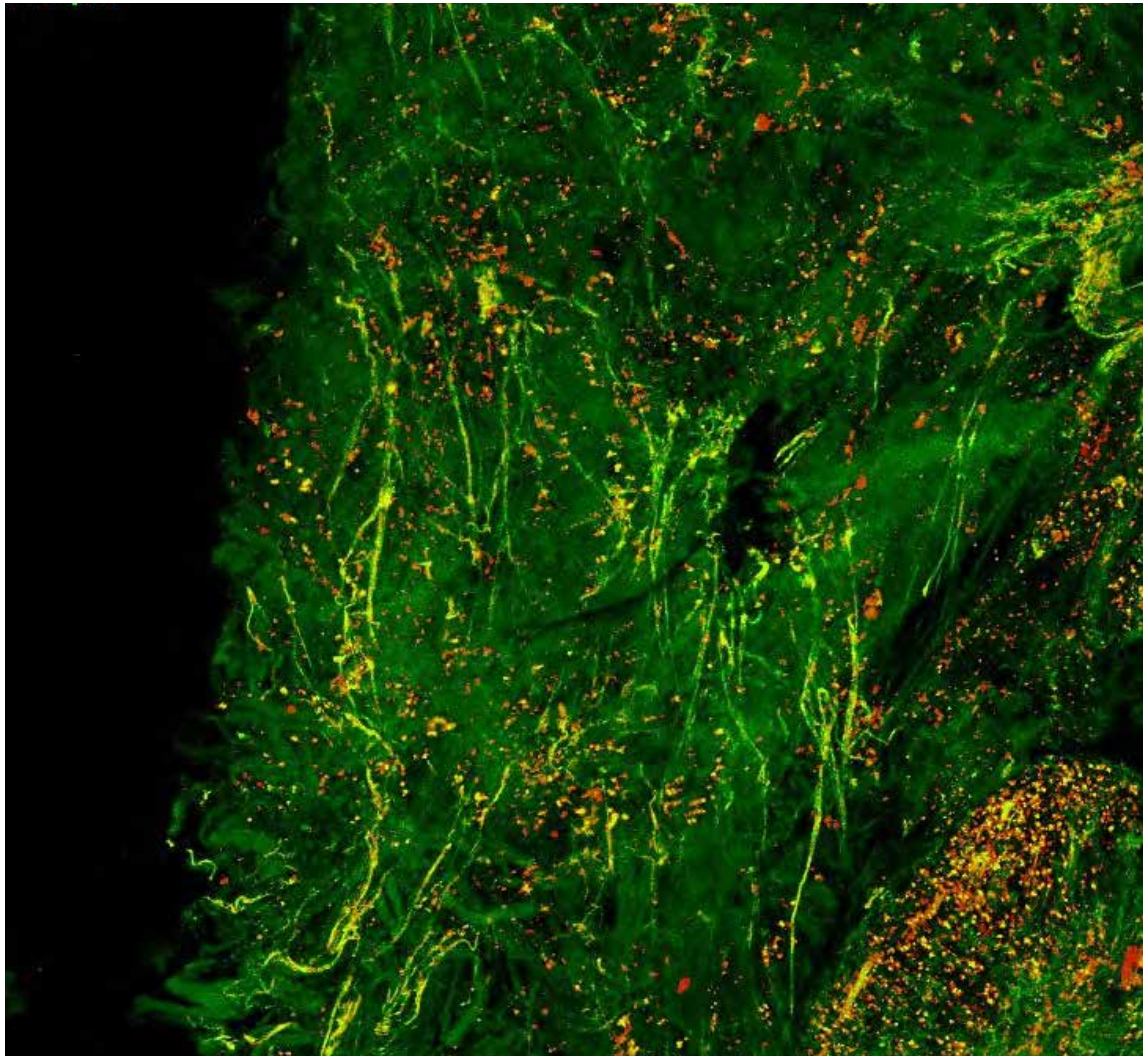

PARPi-FL (green) + Autofluorescence (red/yellow)

6344

### Close-up

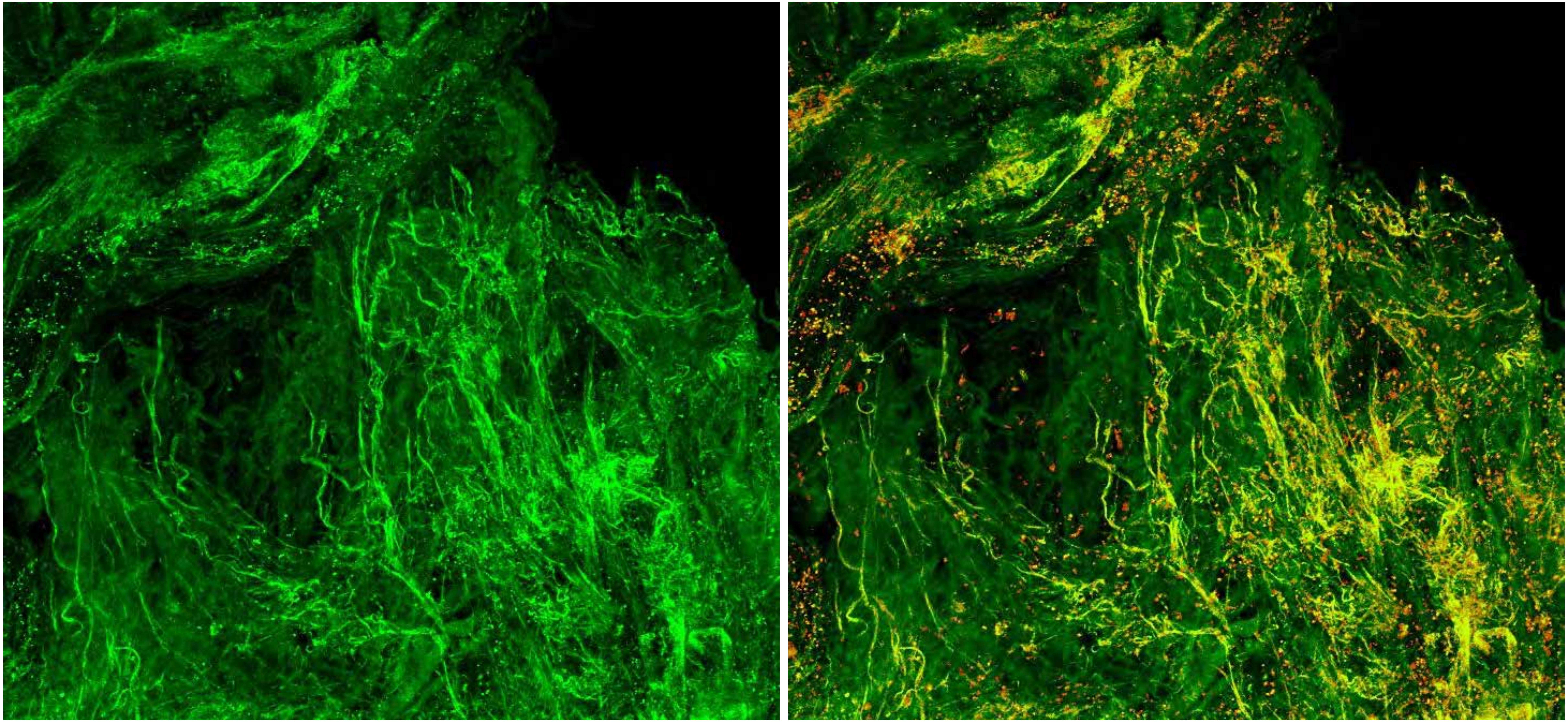

PARPi-FL only (green)　　PARPi-FL (green) + Autofluorescence (red/yellow)

6344



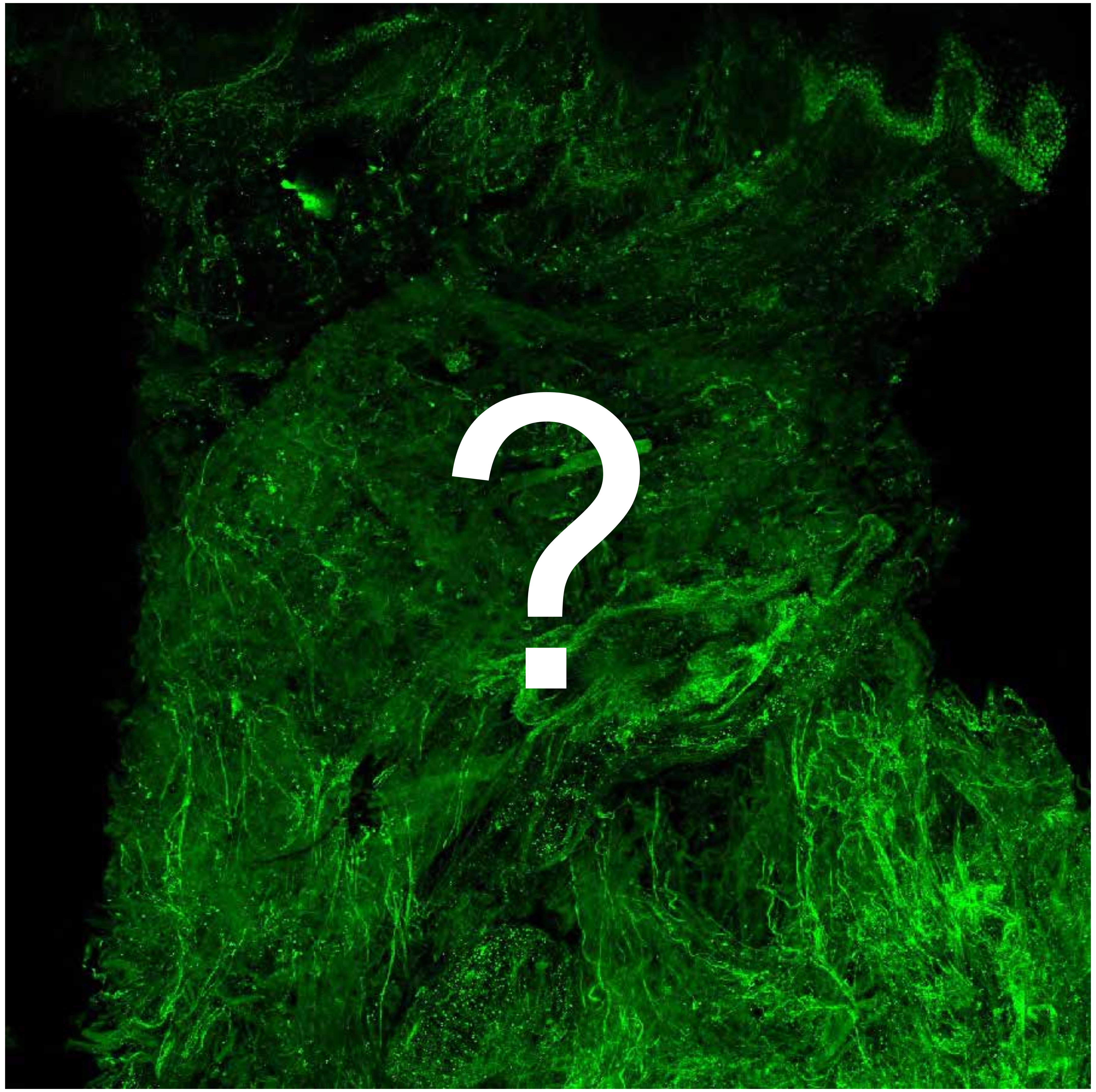


Tumor

Margin

9813

### Overview

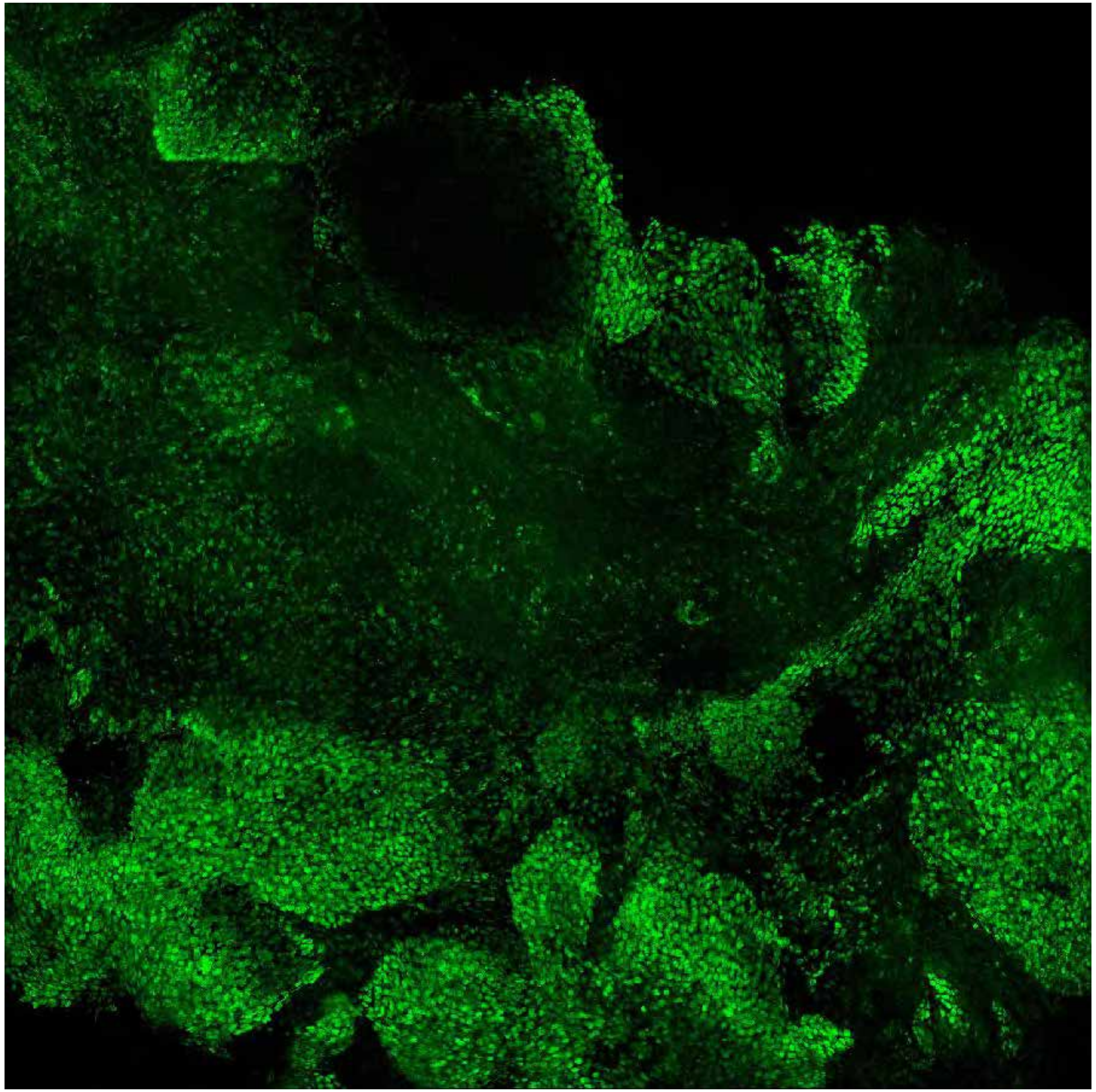

**PARPi-FL only (green)**

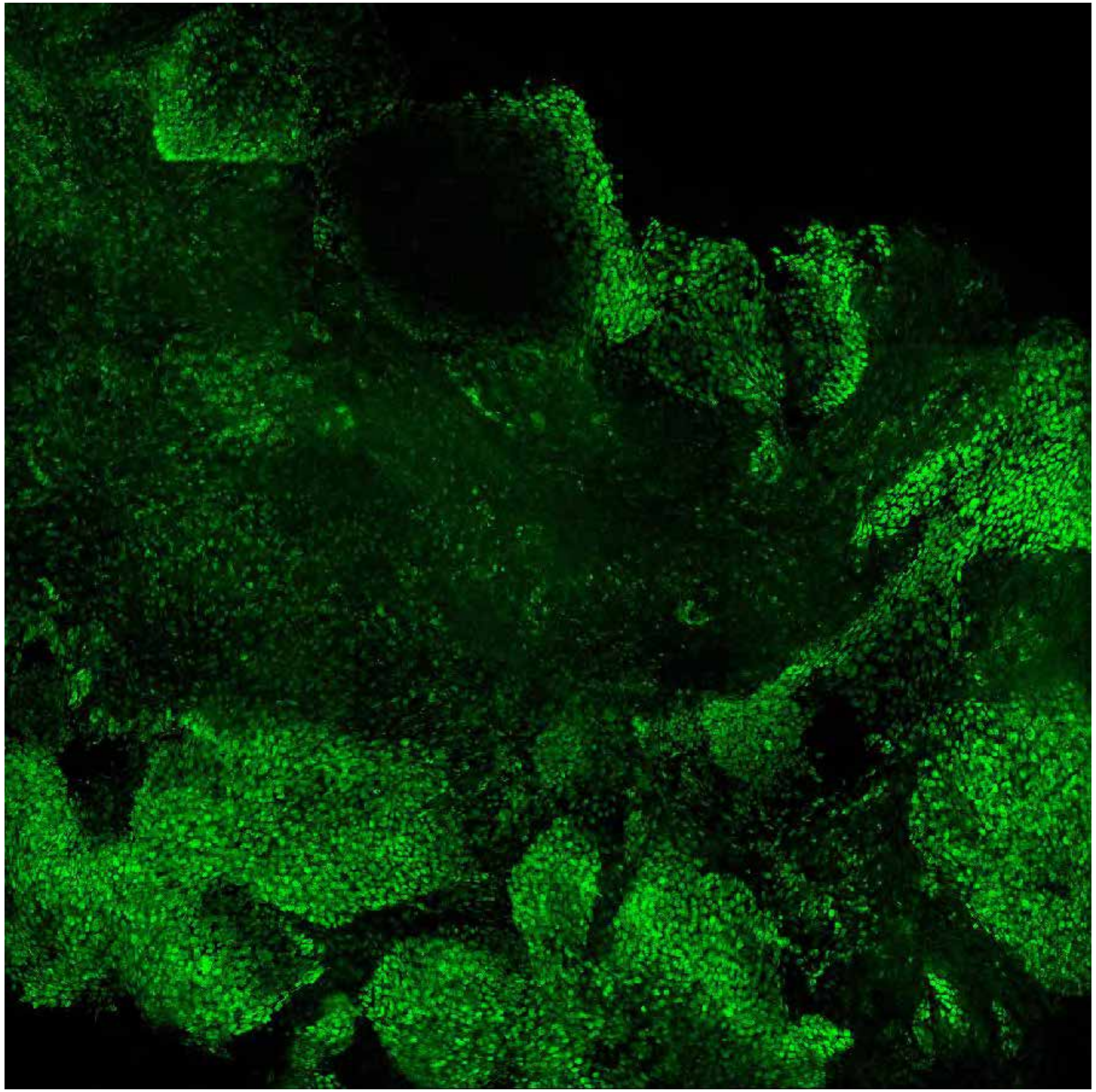

**PARPi-FL (green) + Autofluorescence (red/yellow)**

9813

### Close-up

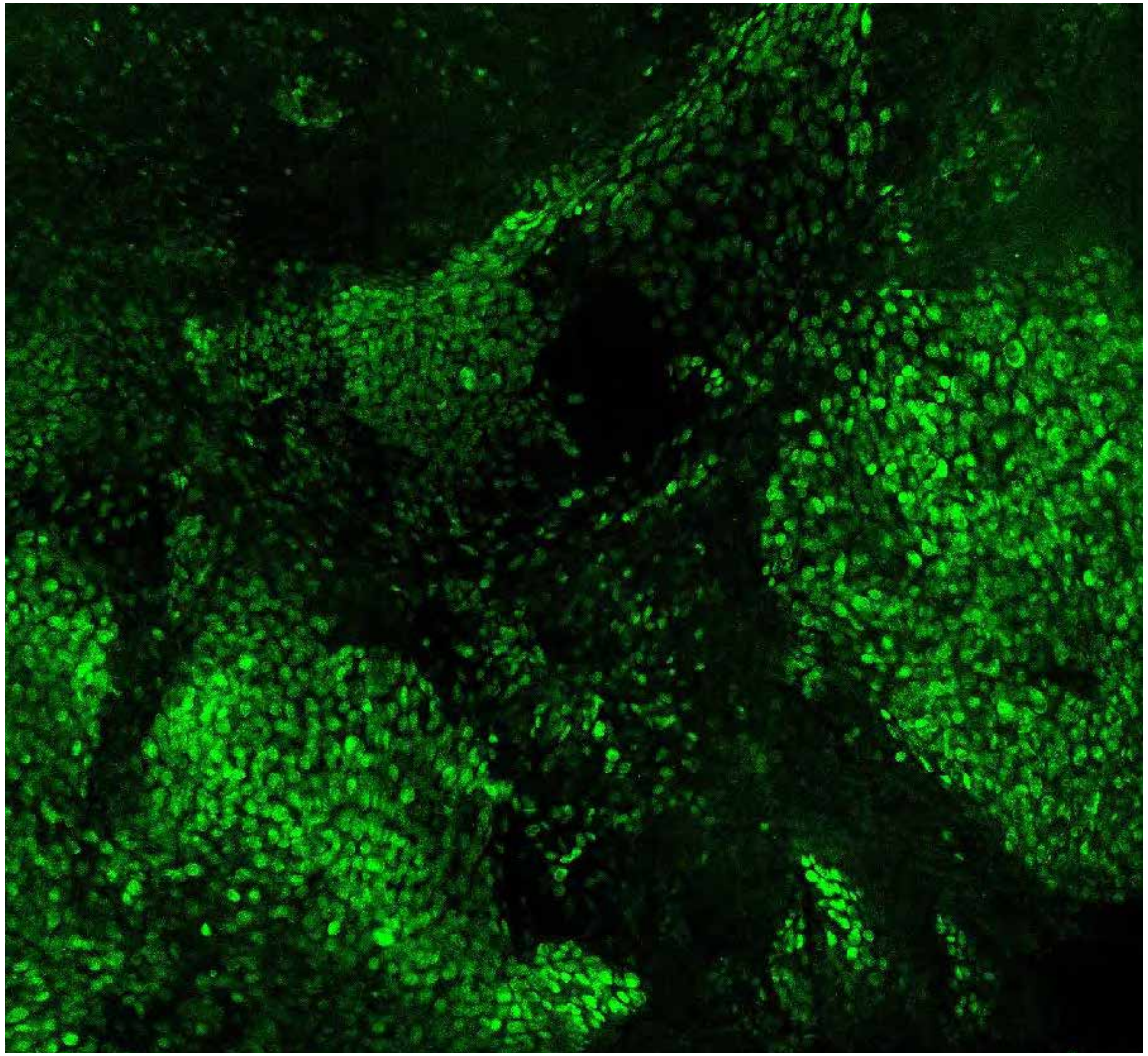

PARPi-FL only (green)

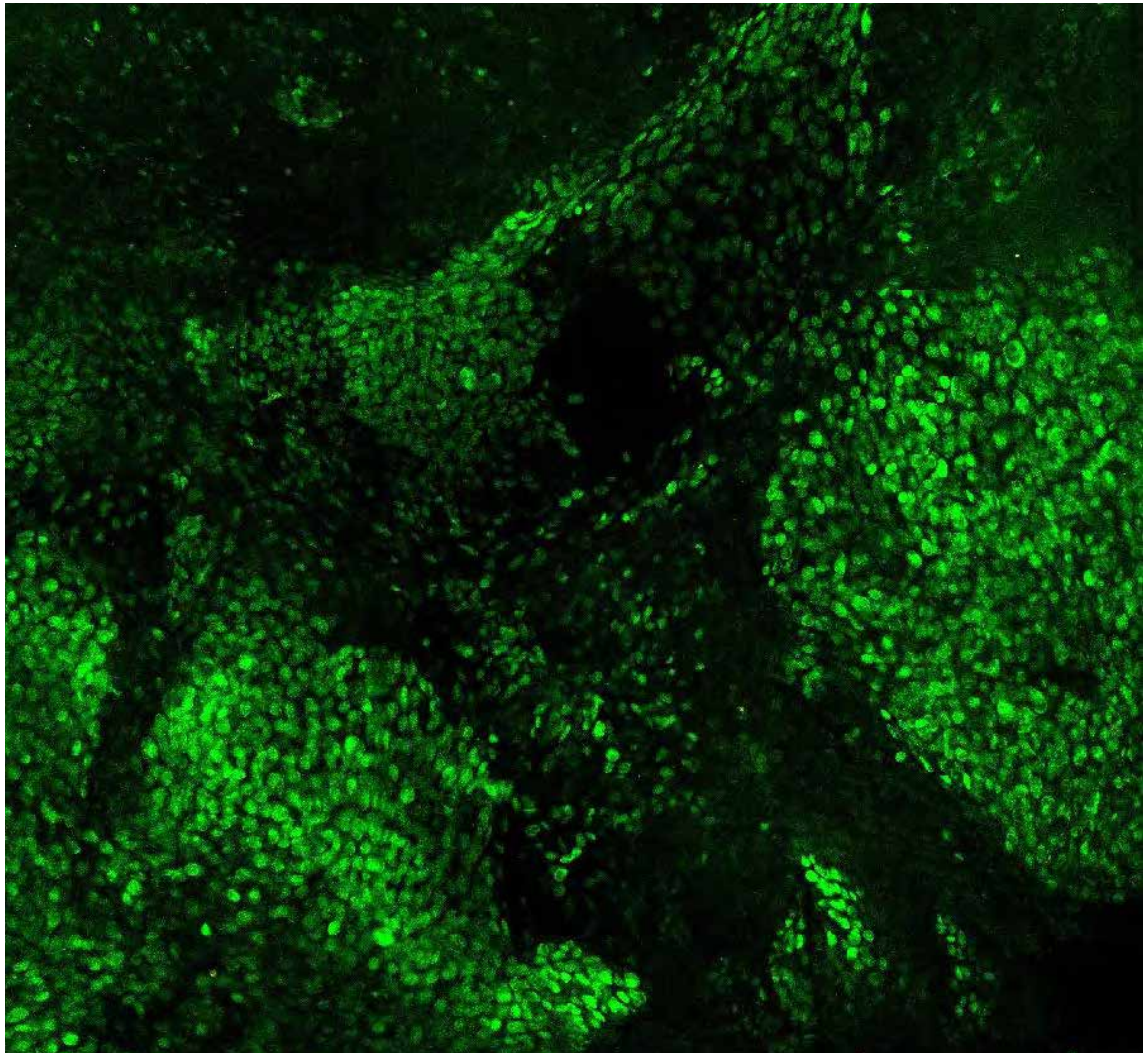

PARPi-FL (green) + Autofluorescence (red/yellow)

9813

### Close-up

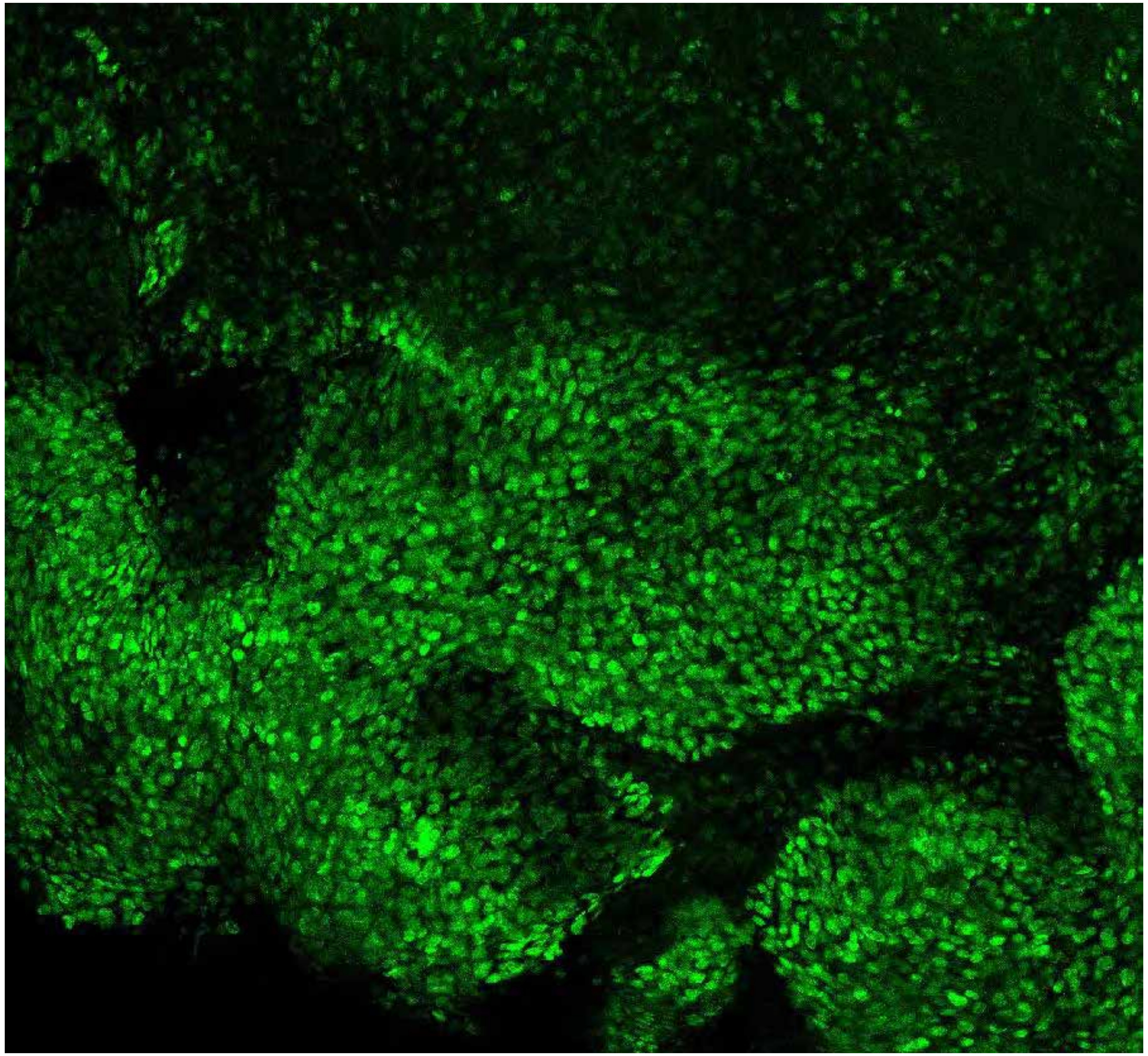

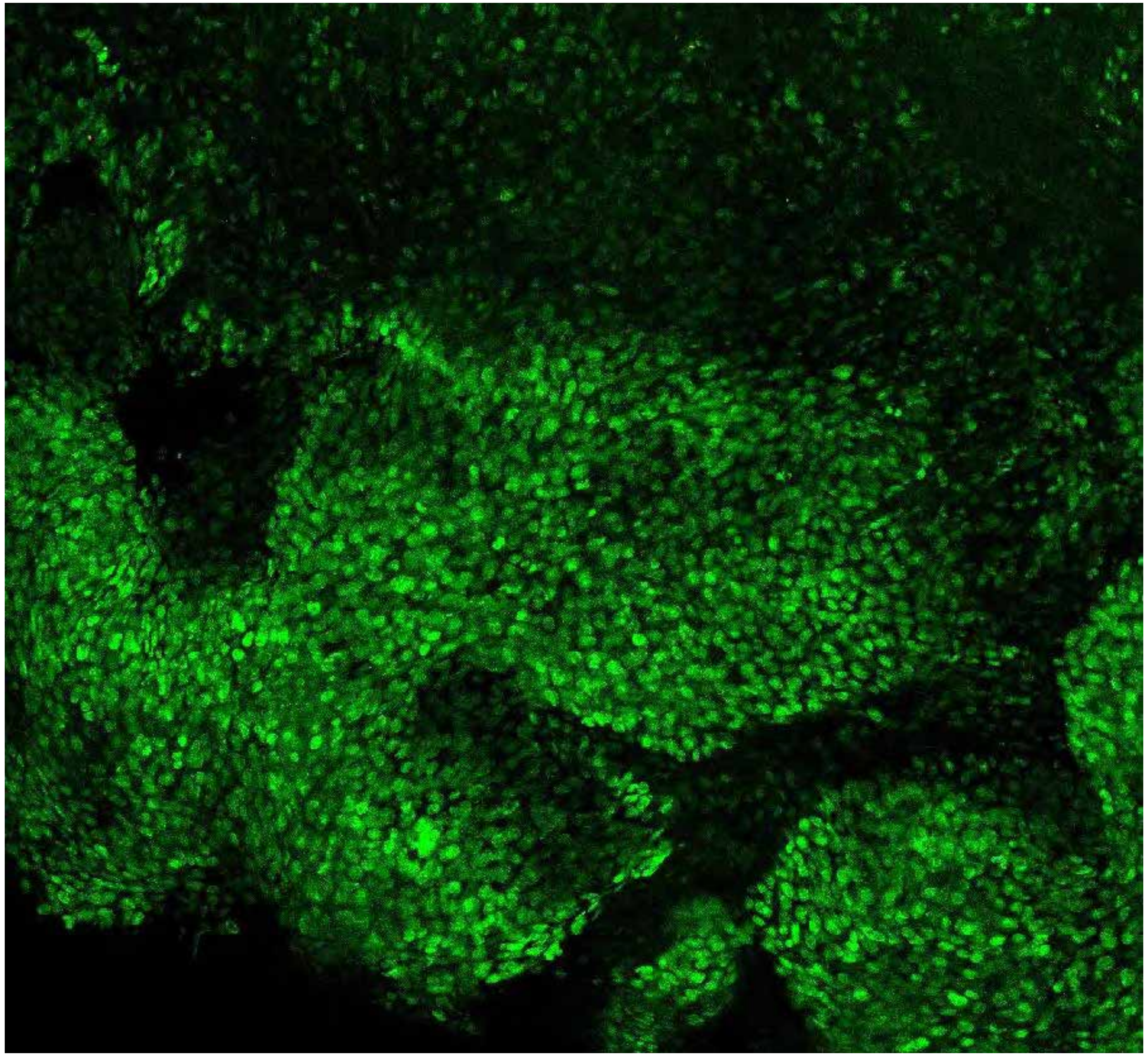

**PARPi-FL only (green)**

**PARPi-FL (green) + Autofluorescence (red/yellow)**

9813

### Close-up

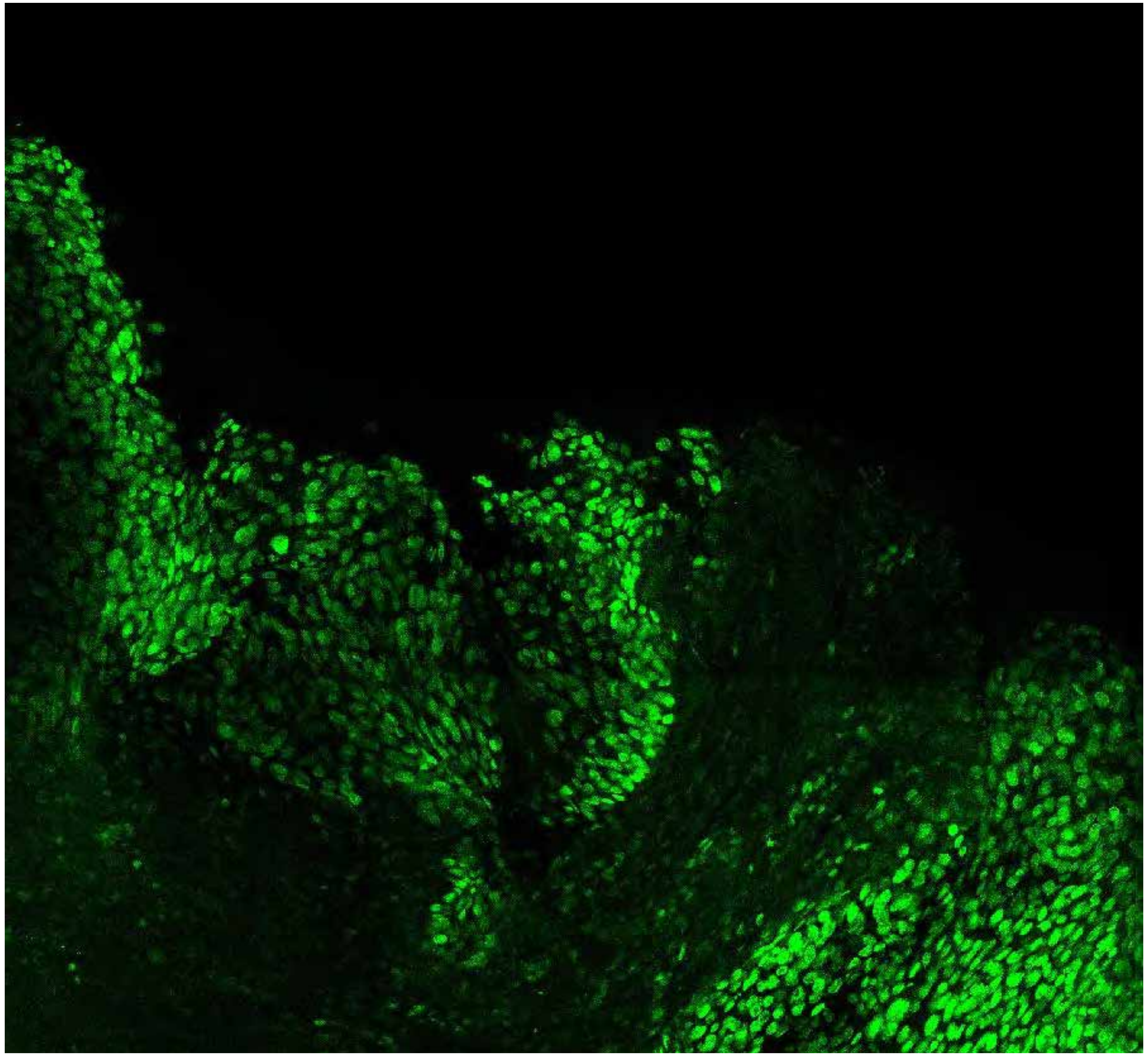

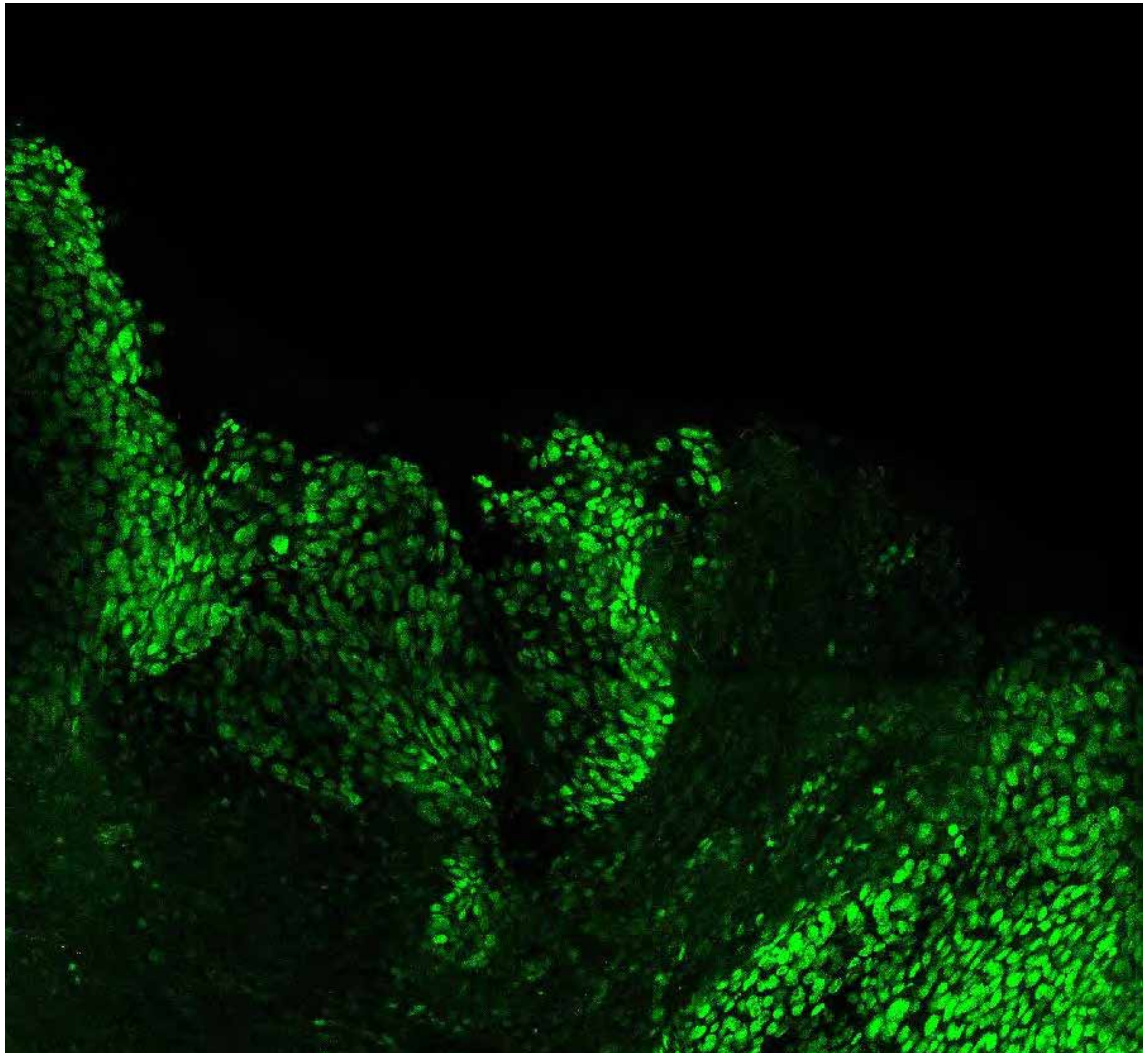

**PARPi-FL only (green)**

**PARPi-FL (green) + Autofluorescence (red/yellow)**

9813

### Close-up

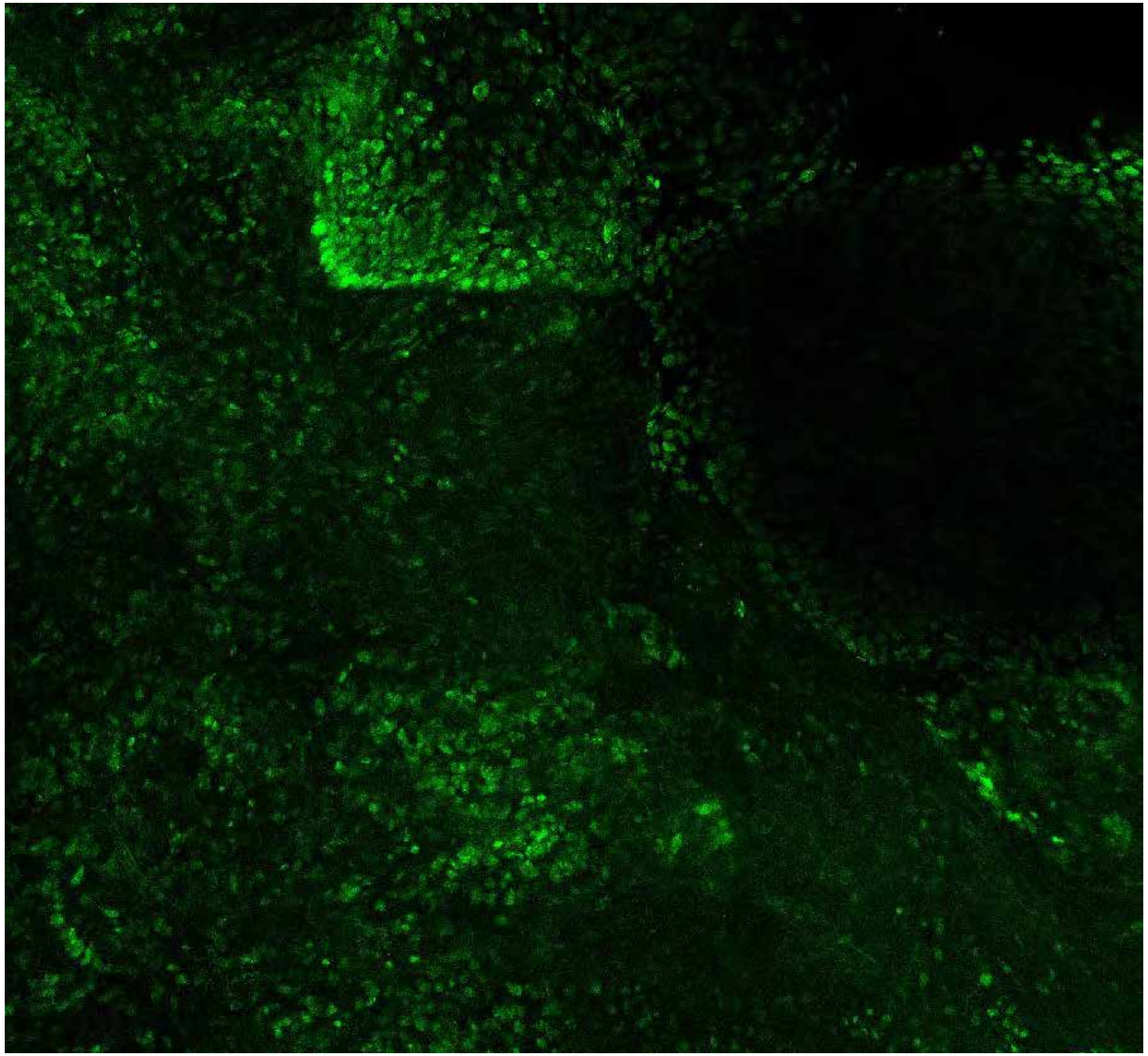

PARPi-FL only (green)

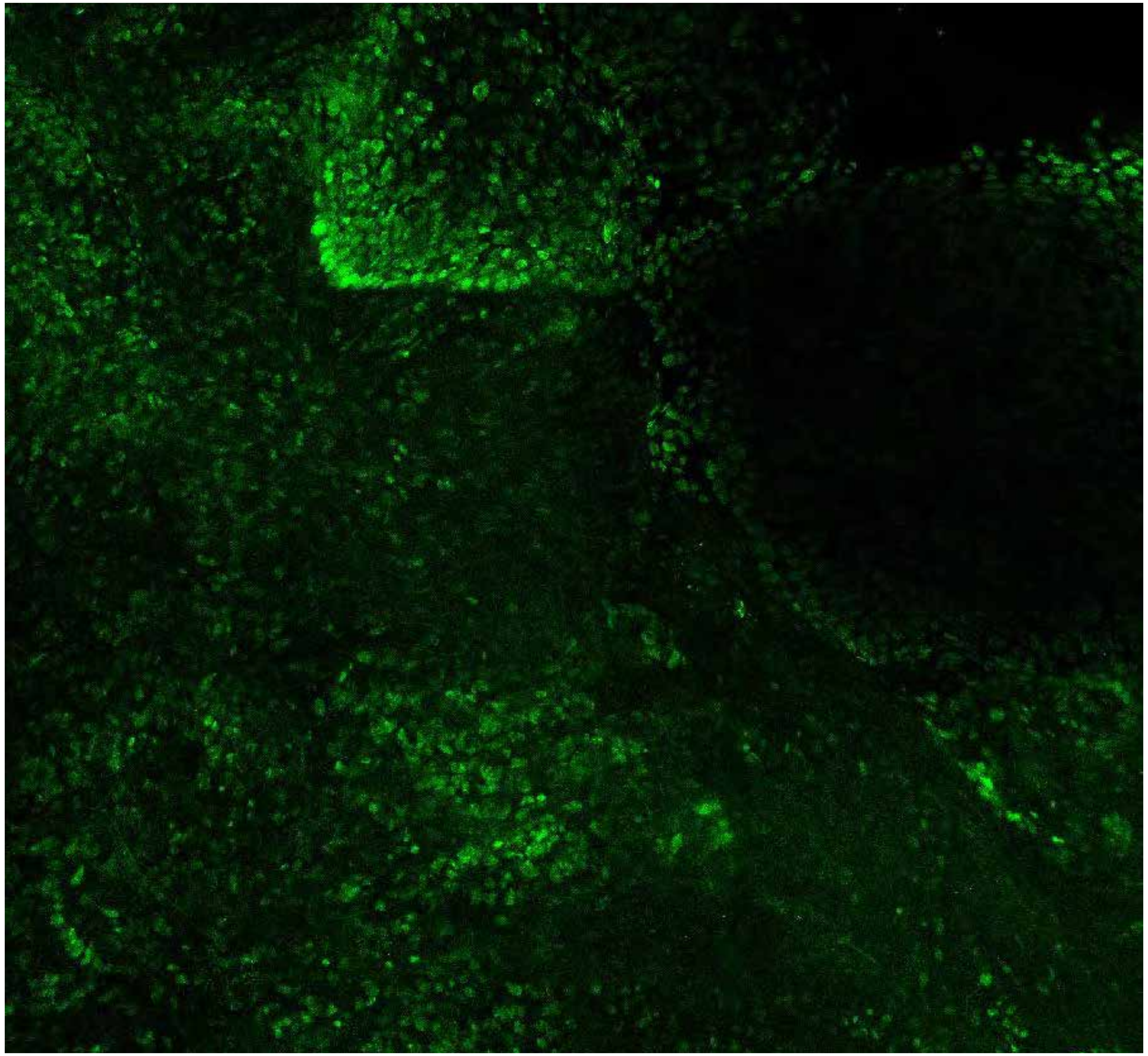

PARPi-FL (green) + Autofluorescence (red/yellow)

9813



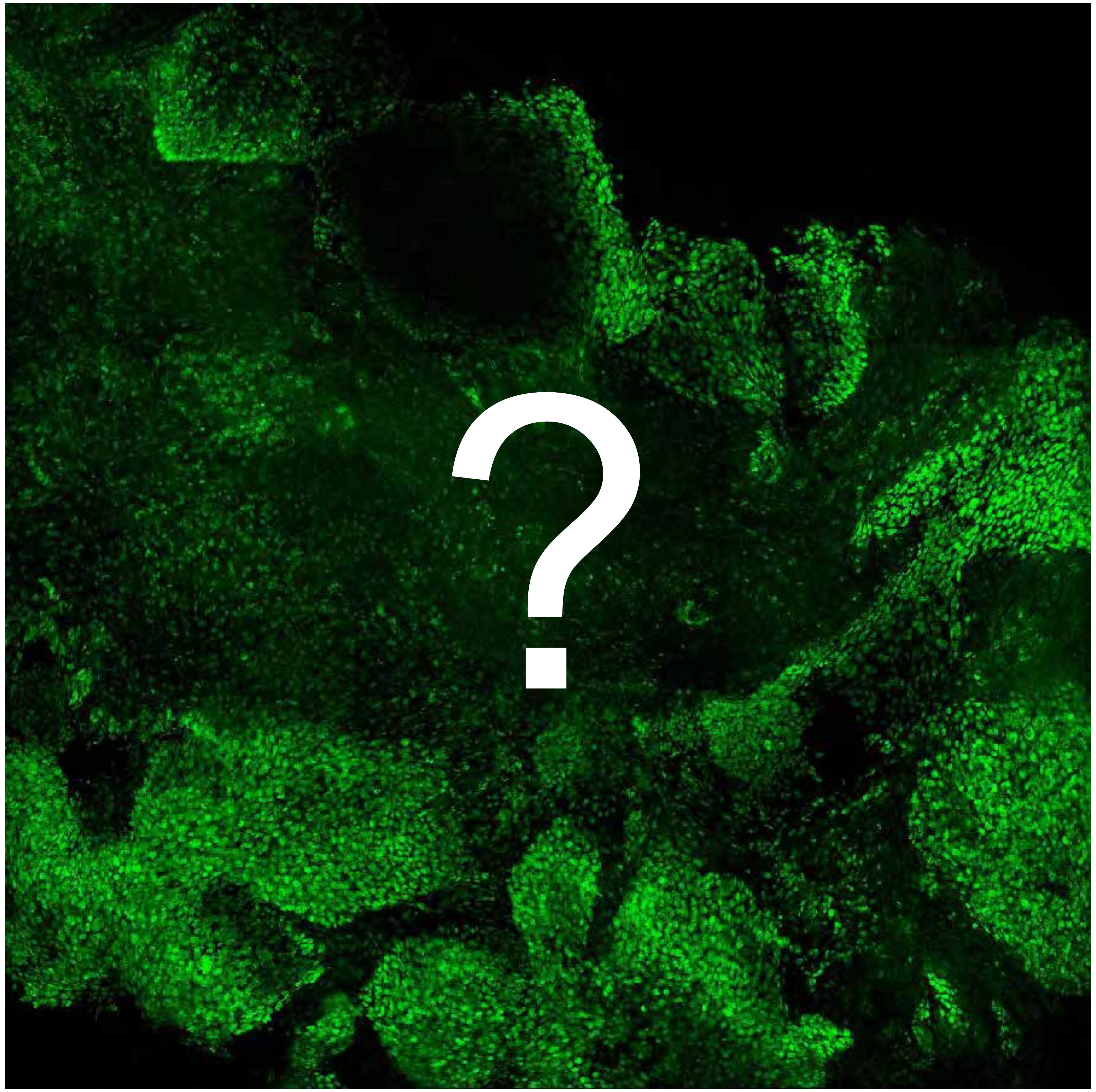


Tumor

Margin

### Congratulations, you are done!

Please hand your scoring sheet to the study supervisor and have a nice day.
